# Thousandfold Cell-Specific Pharmacology of Neurotransmission

**DOI:** 10.1101/2022.10.18.512779

**Authors:** Brenda C. Shields, Haidun Yan, Shaun S.X. Lim, Sasha C. Burwell, Elizabeth A. Fleming, Celine M. Cammarata, Elizabeth W. Kahuno, Purav P. Vagadia, Marie H. Loughran, Lei Zhiquan, Mark E. McDonnell, Miranda L. Scalabrino, Mishek Thapa, Tammy M. Hawley, Allen B. Reitz, Gary E. Schiltz, Court Hull, Greg D. Field, Lindsey L. Glickfeld, Michael R. Tadross

## Abstract

Cell-specific pharmaceutical technologies promise mechanistic insight into clinical drugs―those that treat, and often define, human disease. In particular, **DART** (drug acutely restricted by tethering) achieves genetically programmable control of drug concentration over cellular dimensions. The method is compatible with clinical pharmaceuticals and amenable to studies in behaving animals. Here, we describe **DART.2**, comprising three advances. First, we improve the efficiency of chemical capture, enabling cell-specific accumulation of drug to ∼3,000-times the ambient concentration in 15 min. Second, we develop tracer reagents, providing a behavior-independent measure of cellular target engagement in each animal. Third, we extend the method to positive allosteric modulators and outline design principles for this clinically significant class. We showcase the platform with four pharmaceuticals―two that weaken excitatory (AMPAR) or inhibitory (GABA_A_R) chemical neurotransmission, and two that strengthen these forms of synaptic communication. Across four labs, we tested reagents in the mouse cerebellum, basal ganglia, visual cortex, and retina. Collectively, we demonstrate robust, bidirectional editing of chemical neurotransmission. We provide for distribution of validated reagents, community design principles, and synthetic building blocks for application to diverse pharmaceuticals.

## INTRODUCTION

Pharmaceutical mechanisms are difficult to resolve in complex systems, despite knowledge at the molecular scale. The challenge is pronounced in the brain, wherein molecular drug interactions are transformed―by brain circuits―to produce effects on complex behavioral dimensions including mood, anxiety, psychosis, addiction, attention, and movement (Hyman, 2014). Cell-specific technologies offer new hope in overcoming this ‘circuit-pharmacology’ gap by making it possible to deliver a drug to genetically defined cell types, individually or in combination, all while observing effects on brain dynamics and animal behavior. **DART** (drug acutely restricted by tethering) is a cell-specific technology that genetically programs cells to express the HaloTag Protein (**HTP**), which in turn captures the chemical HaloTag Ligand (**HTL**) along with its attached drug (**Rx**). Thus, brain cells are genetically co-opted to boost **Rx** concentration over their own cellular dimensions. The approach is particularly amenable to surface-receptor drugs, as we have shown that the **HTP** can traffick efficiently to the cell surface, where it has unimpeded access to ambient chemical reagents (Shields et al., 2017).

Many pharmaceuticals act at the cell surface. In particular, synaptic chemical transmission between brain cells is predominantly mediated by two varieties of postsynaptic receptor. Each is attuned to a specific chemical and acts within milliseconds to open an ion-selective pore, which is the conduit of biological electricity. In particular, the GABA_A_R (γ-aminobutyric acid receptor) transduces the chemical GABA into electrical inhibition. Conversely, the AMPAR (α-amino-3-hydroxy-5-methylisoxazole-4-propionic acid receptor) converts glutamate into electrical excitation. In this way, brain cells communicate to one another in milliseconds, empowering animals to navigate the world in real time. We previously developed **YM90K**^**DART**^, a cell-specific AMPAR antagonist, and showcased the approach in a mouse model of Parkinson’s disease (Shields et al., 2017). Clinical trials had failed to show efficacy of this drug class in humans with Parkinson’s Disease (Eggert et al., 2010; Lees et al., 2012; Rascol et al., 2012). We likewise saw no efficacy in parkinsonian mice upon delivering **YM90K**^**DART**^ to all cells of a malfunctioning brain region―even side-effects were absent, suggesting therapeutic irrelevance. It was thus surprising that **YM90K**^**DART**^ revitalized healthy motor function when delivered to *fewer* cells in the same brain region (Shields et al., 2017). While counterintuitive, the findings suggest a conserved circuit-pharmacology principle, whereby restricting drug action may unlock a substantial reservoir of untapped efficacy.

To extend and diversify experimental opportunities, **DART.2** offers three new features. The first and most vital is improved cell specificity (up to 3,000-fold in 15 min), enabling an otherwise epileptogenic GABA_A_R antagonist to safely reach desired cells in behaving mice. The second feature improves rigor with a set of multicolor fluorescent tracer reagents, which provide a behavior-independent metric of technical success in each animal. The third and final advance is extension to PAM (positive allosteric modulator) drugs for the GABA_A_R and AMPAR, altogether enabling cell-specific, bidirectional editing of these two forms of chemical neurotransmission. We provide for distribution of validated reagents, along with chemical building blocks and community guidelines for extension to diverse pharmaceuticals.

## RESULTS

### Principles of DART

An **Rx**^**DART**^ is a two-headed chemical with an **Rx** (drug) and **HTL** (HaloTag Ligand) on opposite ends of a flexible linker. DART exploits differences in **Rx** vs **HTL** characteristics to impact ^**+**^**HTP** (genetically programmed, on-target) cells while sparing ^**-**^**HTP** (non-programmed, off-target) cells. A key difference is that ^**-**^**HTP** cells can only interact with ambient **Rx** via low-affinity reversible binding to native receptors. In contrast, ^**+**^**HTP** cells covalently bind the **HTL** moiety to irreversibly accumulate tethered **Rx** as time proceeds. This difference is manifested by the units of dosing, with *concentration* (nM) used for ^**-**^**HTP** cells, and *integrated concentration* (nM × min) for ^**+**^**HTP** cells. The metrics of cell-specificity thus become:

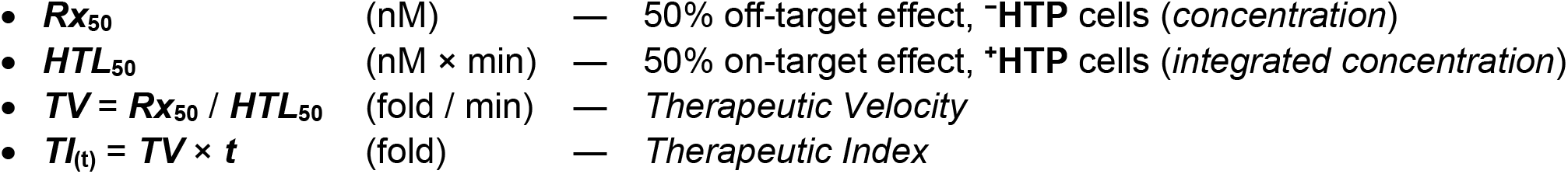

For example, given a reagent in which off-target effects occur at ***Rx***_**50**_ = 10,000 nM, and on-target covalent tethering requires ***HTL***_**50**_ = 5,000 nM × min, the ratio of these terms specifies the *Therapeutic Velocity*, ***TV*** = 2-fold / min, a constant that describes the rate at which cellular specificity is achieved. The textbook quantity, *Therapeutic Index* (***TI***), thus grows with time: at 15 min, ***TI***_**15m**_ = 30-fold; and at 60 min, ***TI***_**60m**_ = 120-fold.

A point that merits clarification is that ***TI*** can, in principle, become arbitrarily large with ***t***. This is because experiments that permit a longer, pre-allocated incubation time, would permit tethering at a proportionally lower concentration. The technical limit to trading time for concentration is degradation of **HTP** after it has captured an **Rx**^**DART**^; this limits linear regime to ∼10-20 hr (Shields et al., 2017). However, faster manipulations are often needed for other reasons. In particular, whole-cell electrophysiology protocols are typically short in duration, and behavioral causality is more interpretable if acute effects can be disambiguated from chronic changes.

### Development of the DART.2 Platform

To improve speed and cellular specificity, we studied the atomic structure of **HTP** in complex with the original **HTL.1** and noted few stabilizing contacts within the inner surface of the **HTP** tunnel (**Figure 1A**, top right). We pursued parallel efforts for chemical and protein engineering to improve shape/charge complementarity. Of the two strategies, chemistry succeeded, via a strategy analogous to a synthon-based approach (Sadybekov et al., 2022). In particular, we modeled the atomic position of minimal chemical moieties (cyan rings in **Figure 1A**, right) known to have some affinity for **HTP** at exterior (Los et al., 2008) and interior regions of the tunnel (Neklesa et al., 2013). We then chemically synthesized a small library of variants in which we interconnected the moieties with linkers of varying length and composition. We synthesized each candidate **HTL** as a fusion to a polyethylene glycol linker (PEG_12_) and biotin. We then tested capture efficiency in the cellular microenvironment of interest, leveraging ^**+**^**HTP** neurons in culture. We incubated cells with ligand for 15 min, washed, and quantified surface biotin (**Figure 1A**, left). The best variant, named **HTL.2**, yields ***HTL***_**50**_ = 150 nM × min (= 10 nM × 15 min), which in side-by-side assays, is 40-times faster than **HTL.1**, with ***HTL***_**50**_ = 6,000 nM × min (= 400 nM × 15 min).

**Figure 1.**
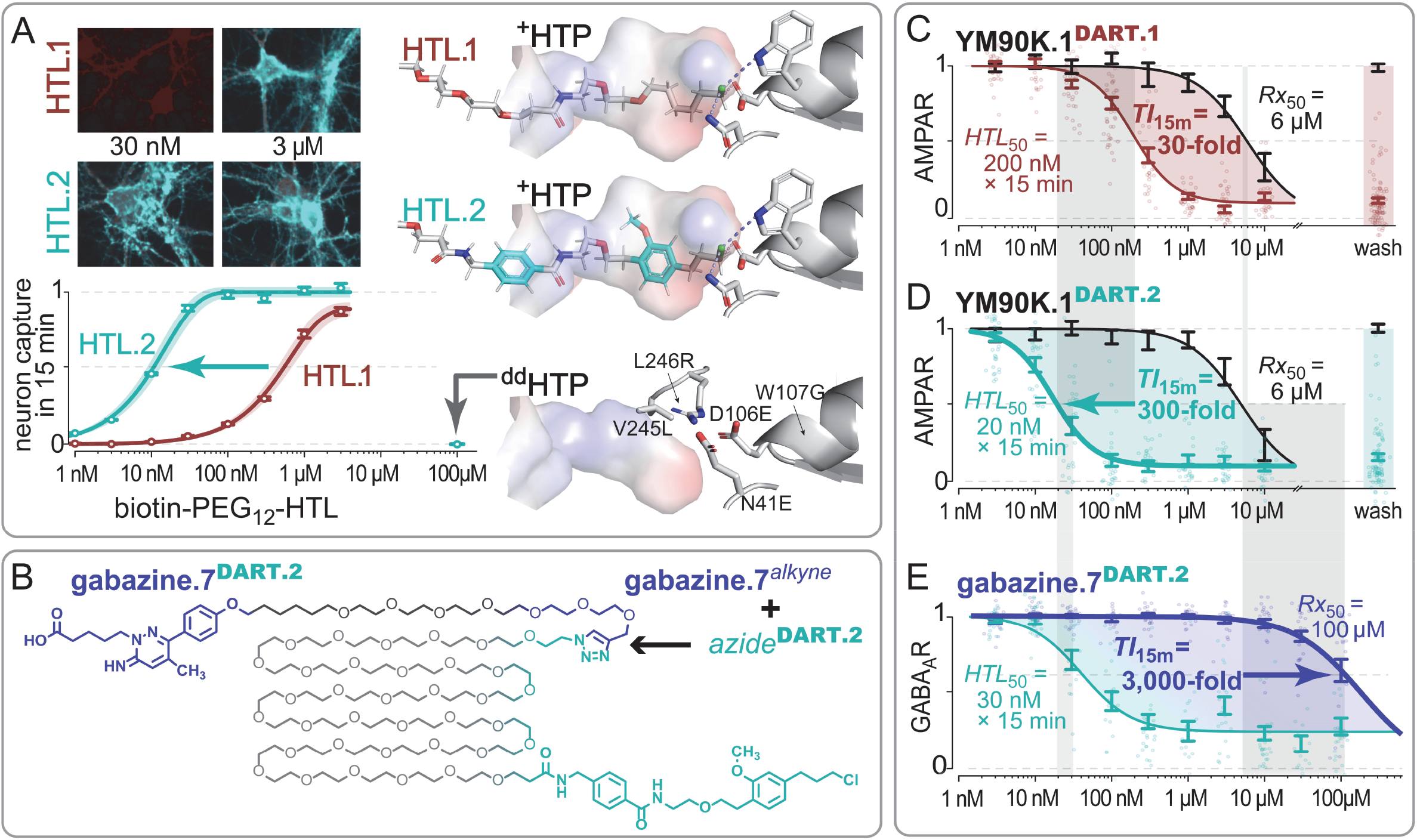
The DART.2 Platform Provides Thousandfold Cell Specificity. **(A) Development of HTL.2**. Top Left: ***HTL***_**50**_ measured on ^**+**^**HTP** neurons; biotin-PEG_12_-HTL applied (at a given dose) for 15 min. Streptavidin label (cyan) quantifies captured biotin, dTomato (dark red) estimates **HTP** expression. Bottom Left: capture dose-response, y-axis is the fraction of surface HTP bound to HTL; error bars, 95% CI from n >1000 cells, via regression of streptavidin vs dTomato (**methods**). Right: Structural model (PDB 4KAJ) of **HTP** in complex with **HTL.1** (top), **HTL.2** (middle), and _**dd**_**HTP** mutations (bottom). **(B)**Modular chemistry. ‘**Rx** fragment’ (**Rx**―short spacer―*alkyne*) is small. ‘**HTL** module’ (*azide*―long linker―**HTL**) is larger due to the long PEG_36_ linker. Final assembly is via *alkyne*-*azide* cycloaddition (e.g., **gabazine.7**^*alkyne*^ + *azide*^DART.2^ → **gabazine.7**^DART.2^). **(C-D)** AMPAR assay. We prepared ChR2-expressing ‘presynaptic’ neurons, co-cultured with ‘postsynaptic’ neurons expressing GCaMP6s and ^**+**^**HTP** or _**dd**_**HTP**. Assay isolates AMPAR-signaling from pre- to post-synaptic cells, with other receptors blocked (**methods**). For **YM90K.1**^DART.1^, ***HTL***_**50**_ = 200 nM × 15 min (on ^+^**HTP** neurons) and ***Rx***_**50**_ = 6,000 nM (on ^**dd**^**HTP** neurons), yielding ***TI***_**15m**_ = 30-fold. For **YM90K.1**^DART.2^, ***HTL***_**50**_ = 20 nM × 15 min (on ^+^**HTP** neurons) and ***Rx***_**50**_ = 6,000 nM (on ^**dd**^**HTP** neurons), yielding ***TI***_**15m**_ = 300-fold. **(E)** GABA_A_R assay. ChR2 and GCaMP6s are co-expressed in the same neuron. Excitatory synapses are blocked, leaving ChR2 as the only trigger of neural activity (**methods**). Activation of endogenous GABA_A_Rs (10 µM GABA throughout) largely overpowers ChR2 to blunt GCaMP activity (=1 on vertical axis indicating maximum GABA_A_R activity). We then perform a dose-response, wherein the antagonism of GABA_A_Rs allows ChR2-mediated GCaMP signals to reawaken. Full GABA_A_R antagonism (=0 on vertical axis) is calibrated with 30 µM traditional gabazine (**methods**). For **gabazine.7**^DART.2^, ***HTL***_**50**_ = 60 nM × 15 min (^+^**HTP** neurons) and ***Rx***_**50**_ = 180,000 nM (^**dd**^**HTP** neurons), yielding ***TI***_**15m**_ = 3,000-fold. Side-by-side assays of other gabazine variants appear in **Figure S1**.

To facilitate synergy among investigators, we standardized the design of ‘**Rx** fragments’ to allow independent development of ‘**HTL** modules’ that can later be co-assembled with one another via the universal *alkyne-azide* click-chemistry reaction (Rostovtsev et al., 2002). Given that reagents will naturally evolve, the chemical structure of each fragment and module is uniquely identified by a version number, delimited by a period for readability (**Figure 1B**). We began by synthesizing an ‘**Rx** fragment’ named **YM90K.1**^*alkyne*^, and two ‘**HTL** modules’ named *azide*^DART.1^ and *azide*^DART.2^. Following assembly, which crosslinks the *alkyne* and *azide*, we obtained **YM90K.1**^DART.1^ and **YM90K.1**^DART.2^, which differ only in their **HTL.1** vs **HTL.2** moieties (**methods**).

To improve rigor of control experiments, we developed an inactive **HTP** that cannot capture an **Rx**^**DART**^, thus enabling measurement of ambient drug effects (***Rx***_**50**_) while controlling for any effects of **HTP** expression. We first explored mutation of **HTP** at its D106 residue, to which **HTL** forms a covalent bond; however, these variants either disrupted surface trafficking in neurons (e.g., D106A) or retained residual capture (e.g., D106E). We thus explored distributed mutations of the catalytic triad, combining the subtle D106E mutation with W107G and N41E. We combined this with a second strategy, which blocks access to the catalytic site with bulky side-chains V245L and L246R (**Figure 1A**, bottom right). Together, these yielded a ‘double dead’ ^**dd**^**HTP**, which exhibited zero measurable capture of **HTL.2** while maintaining proper surface trafficking (**Figure 1A**, 100 µM).

### Antagonizing the AMPAR and GABA_A_R with Improved Cellular Specificity

We tested **YM90K.1**^**DART.1**^ vs **YM90K.1**^**DART.2**^ side-by-side in neuronal assays of synaptic AMPAR signaling (**Figure 1C-D**, see legend for assay details). Ambient, off-target effects (^**dd**^**HTP** cells, 50% AMPAR block) occurred at ***Rx***_**50**_ = 6,000 nM for both reagents, as expected given their identical **YM90K.1** moiety (**Figure 1C-D**, black data). In contrast, on-target effects (^**+**^**HTP** cells, 50% AMPAR block) differed considerably between the reagents, with **YM90K.1**^DART.1^ requiring ***HTL***_**50**_ = 200 nM × 15 min, corresponding to ***TI***_**15m**_ = 30-fold, and the upgraded variant, **YM90K.1**^DART.2^, achieving ***HTL***_**50**_ = 20 nM × 15 min, yielding ***TI***_**15m**_ = 300-fold (**Figure 1C-D**).

We next focused on the GABA_A_R. Antagonists of this receptor are highly epileptogenic, with even transient off-target effects risking seizure and permanent damage to the brain. Given the unforgiving nature of this concern, we anticipated that optimization of ***Rx***_**50**_ would be needed, in concert with ***HTL***_**50**_ optimization. We thus explored the structure-activity literature of gabazine, a selective GABA_A_R antagonist, and made several variants, including **gabazine.1**^**DART.2**^ (2-butyric acid, 5-H), **gabazine.5**^**DART.2**^ (2-pentanoic acid, 5-H), and **gabazine.7**^**DART.2**^ (2-pentanoic acid, 5-CH_3_) (**Figure 1B, Figure S1B-D**). We then developed an all-optical assay of endogenous GABA_A_Rs in primary neuronal culture (**Figure 1E**, see legend). The most potent **gabazine.1**^**DART.2**^ yields ***TI***_**15m**_ = 400-fold, intermediate **gabazine.5**^**DART.2**^ yields ***TI***_**15m**_ = 1,700-fold, and attenuated **gabazine.7**^**DART.2**^ achieves ***TI***_**15m**_ = 3,000-fold, which is our maximum to date (**Figure 1E, Figure S1B-D**).

### GABA_A_R Antagonism in Behaving Animals

We next explored GABA_A_R pharmacology in behaving mice. We began with the input layer of the cerebellar cortex, wherein glutamatergic mossy-fiber input is transformed onto a larger number of granule cells (Cayco-Gajic and Silver, 2019). Many features remain unknown regarding this transformation, which underlies the influential Marr-Albus theory of cerebellar learning (Albus, 1971; Marr, 1969). In particular, the extent to which the GABA microcircuit shapes granule cell activity *in vivo* has been a longstanding question (Eccles et al., 1966). Existing tools that silence GABA interneurons (e.g., Golgi cell, **Figure 2A**) can have unintended consequences, as GABA would be withheld from all cells in the microcircuit (Dino et al., 2000). In contrast, **DART** can alter how granule cells listen to GABA, without disrupting GABA release or its impact on neighboring cells. As such, the approach can isolate the impact of GABA on granule cells without imposing unwanted circuit perturbations.

**Figure 2.**
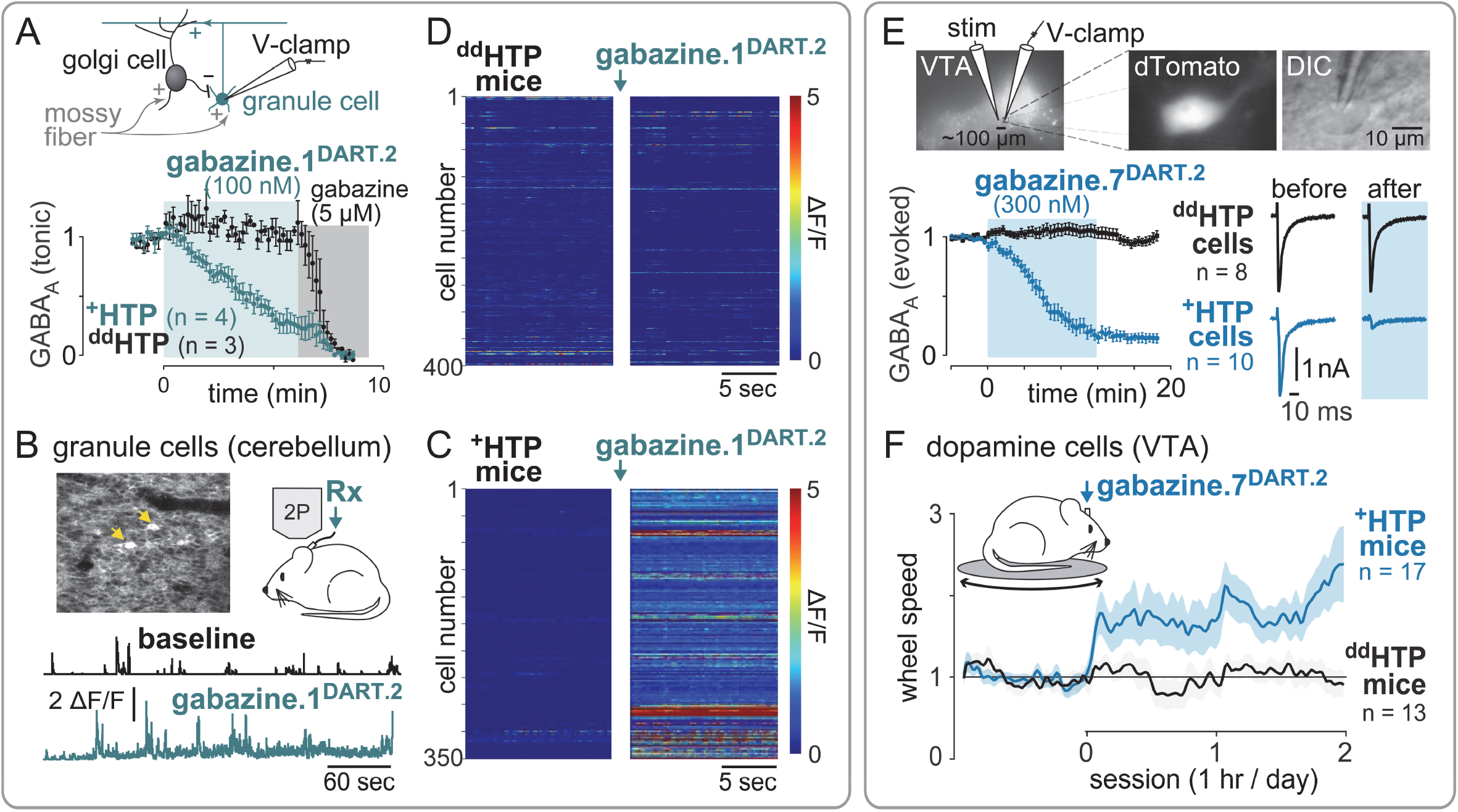
DART.2 enables safe delivery of epileptogenic drugs in behaving animals. **(A)** Validation of **gabazine.1**^**DART.2**^ in cerebellar slice. Top: voltage-clamp schematic. Bottom: tonic GABA_A_R in granule cells expressing ^**+**^**HTP** or ^**dd**^**HTP**. Application of **gabazine.1**^**DART.2**^ (100 nM) antagonizes the GABA_A_R in ^**+**^**HTP** but not ^**dd**^**HTP** cells. Subsequent application of traditional gabazine blocks all GABA_A_R current. Data are mean ±SEM, cells normalized to baseline. Dose-response in **Figure S2A**. **(B)** In vivo delivery of **gabazine.1**^**DART.2**^ to cerebellar granule cells. Top right: schematic of experimental approach, GCaMP6f granule-cell activity obtained in the absence of sensory stimuli. Top left: example field of view, GCaMP6f signal averaged across trials. Bottom: example calcium traces (ΔF/F) in an awake ^**+**^**HTP** mouse before (baseline) and after (1 µL of 1 µM **gabazine.1**^**DART.2**^) infusion. **(C)** In vivo, 1 µL of 1 µM **gabazine.1**^**DART.2**^ to ^**+**^**HTP** mice. Color indicates ΔF/F, each row is one cell before (left) and after (right) **gabazine.1**^**DART.2**^ infusion. Full dataset from 6 mice in **Figure S2C** (n = 1707 cells, p < 0.0001, paired t-test, before vs after infusion). **(D)** In vivo, 1 µL of 1 µM **gabazine.1**^**DART.2**^ to _**dd**_**HTP** mice. Format as in panel C. Full dataset from 4 mice appears in **Figure S2B** (n = 1084 cells, p = 0.76, paired t-test, before vs after infusion). **(E)** Validation of **gabazine.7**^**DART.2**^ in VTA slice. Top: evoked-IPSC configuration. Bottom: 300 nM **gabazine.7**^**DART.2**^ has no impact on _**dd**_**HTP** neurons, while blocking IPSCs on ^**+**^**HTP** cells in under 15 min. Data show mean ±SEM of cells normalized to baseline. Example traces to the right. Control experiments appear in **Figure S2D-H**, confirming the GABA_A_R specificity of the manipulation. **(F)** In vivo, 1.2 µL of 10 µM **gabazine.7**^**DART.2**^ infusion into the VTA increases locomotor speed in VTA_DA **+**_**HTP** mice (n=17) but not in _**dd**_**HTP** control mice (n=13). Data are mean ±SEM, normalized to baseline speed of each animal. The difference in ^**+**^**HTP** vs _**dd**_**HTP** post-infusion speed demonstrates efficacy of the tethered, but not the ambient compound (p < 0.01, unpaired t-test).

We optimized a matched pair of AAV reagents, **AAV-DIO-**^**+**^**HTP**_**GPI**_ and **AAV-DIO-**^**dd**^**HTP**_**GPI**_, to achieve robust expression in granule cells; with key improvements including use of a GPI anchor in place of the transmembrane anchor used in our earlier work (Shields et al., 2017) and selection of a capsid serotype that fortuitously improved expression in this cell type (**methods**). To limit expression to granule cells, we incorporated the DIO (double inverted open) switch, making AAV expression dependent on Cre recombinase, and used BACα6Cre-A mice, which express Cre only in cerebellar granule cells (Aller et al., 2003). We validated potency and cellular specificity of reagents in acute cerebellar slices (**methods**), leveraging whole-cell electrophysiology to measure tonic GABA_A_R current (Brickley et al., 1996; Rossi and Hamann, 1998). On ^**+**^**HTP** cells, GABA_A_R signaling was antagonized by **gabazine.1**^**DART.2**^ at ***HTL***_**50**_ ∼ 20 nM × 15 min. In contrast, off-target effects on ^**dd**^**HTP** cells required a higher concentration of ***Rx***_**50**_ ∼ 1,000 nM (**Figure 2A, Figure S2A)**

To allow optical recording of neural activity, we used **AAV-DIO-**^**+**^**HTP**_**GPI**_ and **AAV-DIO-**^**dd**^**HTP**_**GPI**_ in BACα6Cre-A mice crossed with Ai148 mice (Chen et al., 2013), which provide stable GCaMP6f expression in granule cells. We implanted a cannula and imaging window, enabling visualization of granule cell activity before and after drug delivery (**Figure 2B, methods**). Infusing 1 µL of 1 µM **gabazine.1**^**DART.2**^ in awake ^**+**^**HTP** mice caused a pronounced ∼9-fold increase in granule-cell activity (i.e., GCaMP6f ΔF/F, **Figure 2B-C, Figure S2C**). In contrast, the same infusion had no effect in ^**dd**^**HTP** mice, indicating no measurable impact of the ambient compound (**Figure 2D, Figure S2B**). Together, these data indicate that **gabazine.1**^**DART.2**^ is efficacious only when tethered onto ^**+**^**HTP** cells. With regard to cerebellar function, we revealed a pronounced role of GABA_A_R signaling on granule cells, supporting predictions of optimal signal-to-noise encoding (Duguid et al., 2012). These findings are further elaborated in a companion study wherein **gabazine.1**^**DART.2**^ is used to interrogate Marr-Albus theory in mice performing a cerebellar-dependent sensorimotor discrimination task (Fleming et al., 2022).

To test for general utility of **DART.2** tools, we examined a different brain region and cell type. In particular, we focused on VTA (ventral tegmental area) dopamine neurons (VTA_DA_ neurons). A notable feature of these cells is their intrinsic pacemaker activity, wherein their non-zero baseline activity allows a single neuron to encode valence in the positive or negative direction (Schultz et al., 1997). **DART.2** offers a unique opportunity in this context, as it can edit chemical neurotransmission onto VTA_DA_ neurons without directly perturbing their intrinsic rhythm―a conceptually needed but thus far absent manipulation in the neuroscience toolkit (Bolkan et al., 2022; Pan et al., 2021). In pilot experiments, we found the VTA to be more sensitive to epileptogenic side effects than the cerebellum; a challenge exacerbated by the larger brain volume of interest, which requires more ligand than the prior example. We thus focused on **gabazine.7**^**DART.2**^ to allow safe infusion at a higher dose. We expressed **AAV-DIO-**^**+**^**HTP**_**GPI**_ or **AAV-DIO-**^**dd**^**HTP**_**GPI**_ in DAT-Cre mice, limiting expression to VTA_DA_ neurons (**methods**). We validated tools in acute slice, which confirmed a pronounced block of evoked GABA_A_R current onto ^**+**^**HTP** VTA_DA_ neurons, with no measurable effect on ^**dd**^**HTP** cells (**Figure 2E**). We further tested receptor specificity of **gabazine.7**^**DART.2**^ when tethered onto ^**+**^**HTP** cells and found no impact on NMDA or AMPA-mediated transmission, nor on other measurable electrophysiological parameters including pacemaker firing rate or action potential waveform (**Figure S2D-G**). Altogether, these findings indicate a selective receptor-specific manipulation of the GABA_A_R by **gabazine.7**^**DART.2**^.

We next looked for locomotor effects of the manipulation, using the same viral parameters along with an implanted cannula. Infusion of 1.2 µL of 10 µM **gabazine.7**^**DART.2**^ had no impact on locomotion in ^**dd**^**HTP** animals, nor any other observable behavioral effects. In contrast, the same infusion produced a significant increase in running speed in ^**+**^**HTP** mice, indicative of cell-specific potency (**Figure 2F**). These findings implicate the GABA_A_R on VTA_DA_ neurons as a causal determinant of locomotion, mirroring the locomotor sensitization typically produced by psychostimulant drugs of abuse, which also engage the dopamine system (Grace, 1995). More generally, these data establish the safety and efficacy of a set of **gabazine**^**DART.2**^ reagents in behaving mice―a milestone in safe, cell-specific delivery of an epileptogenic pharmaceutical.

### Quantitative Cellular Target Engagement

A behavioral difference in manipulated vs control animals is the gold standard of technical success in behavioral neuroscience. However, in the absence of a behavioral effect, it can be unclear whether to infer biological insight or suspect technical failure. With neuropharmacology in particular, ground-truth calibration of the dose-response requires an *ex-vivo* brain slice, wherein drug concentration is precise. Yet, it remains unclear how to equivalently dose a given brain region *in vivo*―wherein drug concentration exhibits complex spatiotemporal profiles, and can be inconsistent across animals due to variable coordinates and fluid handling.

We sought to address the issue by developing **tracer**^**DART.2**^ reagents, which mimic an **Rx**^**DART.2**^ with regard to mass, electrostatics, and capture efficiency, yet with a dye in place of the drug. Mixing these reagents in a consistent one-to-ten ratio would allow each **tracer**^**DART.2**^ molecule to serve as a visible proxy for ten **Rx**^**DART.2**^ molecules nearby. Moreover, because capture is covalent, fluorescence could be quantified many hr later, making brain-wide quantitation of cellular capture possible. Thus, by the logic of ratiometric dosing, any two cells with similar dye capture would be assured to have accumulated comparable amounts of drug, irrespective of the ambient dose given earlier―whether it was infused into a live animal or onto a brain slice.

We developed **Alexa488.1**^**DART.2**^ (green) and **Alexa647.1**^**DART.2**^ (far red) reagents (**methods**), and tested ratiometric dosing in a live-mouse locomotor assay. We focused on the dorsal striatum, wherein delivery of the original **YM90K**^**DART.1**^ reagent to D1 neurons in one hemisphere is known to cause ipsiversive turning (Shields et al., 2017). In similar fashion, we injected **AAV-DIO-**^**+**^**HTP**_**GPI**_ or **AAV-DIO-**^**dd**^**HTP**_**GPI**_ in D1-Cre mice and implanted a cannula in the left dorsal striatum (**methods**). We performed open-field behavior to quantify turning via automated video tracking. This was repeated before and after infusion of a one-to-ten mixture of **Alexa488.1**^**DART.2**^ + **YM90K.1**^**DART.2**^ (3 µM dye + 30 µM drug, infusion: 1 µL in 10 min). We then perfused the mice for histology, performed measurement of dye capture integrated over the dorsal striatum (**methods**), and plotted the magnitude of open-field turning vs dye capture for each animal (**Figure 3A**).

**Figure 3.**
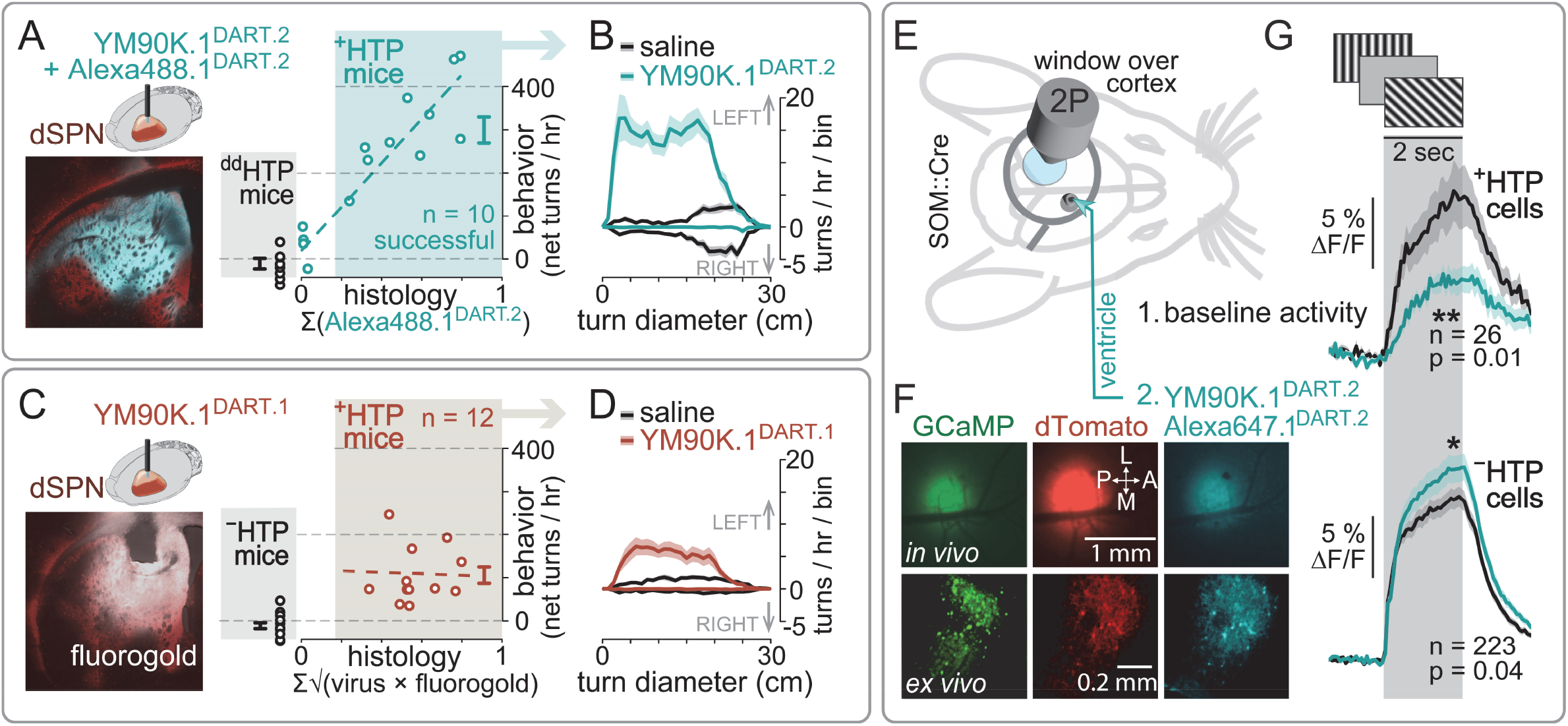
Quantitative Target Engagement and Whole-Brain Delivery. **(A)** Mice with ^**+**^**HTP** or _**dd**_**HTP** in D1 cells of left dorsal striatum. Open field (1 hr) before and after **Alexa488.1**^**DART.2**^ + **YM90K.1**^**DART.2**^ (3 µM dye + 30 µM drug, 1 µL in 10 min). Histology performed 24 hr later. Left, dTomato expression (red) and **Alexa488.1**^**DART.2**^ capture (cyan) from a ^**+**^**HTP** mouse. No dye capture detected in _**dd**_**HTP** mice. Right, open-field turning vs dye capture (**methods**). Each symbol is one mouse. _**dd**_**HTP** mice (black) exhibit little turning. ^**+**^**HTP** mice (cyan) turn in proportion to dye capture. Dye capture can be used as a behavior-independent inclusion criterion (shading), allowing exclusion of 4 ^**+**^**HTP** mice in this dataset (cyan data outside of the shading). **(B)** Fine-scale analysis for ^**+**^**HTP** mice that met the histology inclusion criteria (n = 10). Data is a histogram of turn diameter (1 cm bins), with left turns (upward histogram) tallied separately from right turns (downward histogram). Data are mean ±SEM over 10 mice. Black data (saline infusion) shows symmetric large-diameter turning, indicating low viral toxicity. Cyan data (**Alexa488.1**^**DART.2**^ + **YM90K.1**^**DART.2**^ infusion in the same 10 mice), eliminated right turns (downward histogram) and promoted narrow-diameter left turns (upward histogram). **(C)** Older technology lacked **tracer**^**DART**^. Here, we attempted to estimate technical success with a non-DART dye (fluorogold) and a numerical algorithm to estimate dye-virus overlap. This did not provide a predictive histology marker of target engagement (right, correlation is absent when considering only the ^**+**^**HTP** mice). Format as in panel A; adapted from (Shields et al., 2017). **(D)** Fine-scale analysis for ^**+**^**HTP** mice with older technology (n = 12). Format as in panel B; adapted from (Shields et al., 2017). **(E)** Mice expressing pan-neuronal GCaMP8s with SOM-specific ^**+**^**HTP**. Window over V1. DART delivered via the contralateral ventricle. **(F)** In-vivo and ex-vivo expression and dye capture (see **Figure S3A** for ambient pharmacokinetics). **(G)** Mice presented oriented visual gratings, 2P Ca^2+^ imaging obtained before (black traces) and 5-6 hr after (cyan traces) infusion of 0.3 nmole **Alexa647.1**^**DART.2**^ + 3 nmole **YM90K.1**^**DART.2**^ (2 µL volume) into the contralateral ventricle. Data show mean ± SEM of cells from 3 mice. Following the manipulation, responses in ^**+**^**HTP** SOM cells is reduced (p=0.01, n=26, paired t-test), and responses in ^**–**^**HTP**, putative pyramidal cells, were unchanged or increased (p=0.04, n=223, paired t-test). See **Figure S3B** for cell-by-cell analysis.

Three takeaways merit emphasis. First, we saw a turning bias in ^**+**^**HTP** mice, with no turning bias in ^**dd**^**HTP** animals, confirming that behavioral effects are mediated by tethered drug, not ambient drug. Second, the data exhibit a tight correlation between behavioral potency and histology; in particular, the four animals with the weakest behavioral impact corresponded to those with little dye capture, providing an unbiased parameter with which to exclude these animals due to technical failure (**Figure 3A-B**). A similar correlation was not seen with the original technology, despite our efforts to quantify viral expression and chemical spread (**Figure 3C**). Third, the ability to exclude technical failures in our new dataset enabled a more accurate measurement of the behavioral effect: now 3-fold larger than originally estimated with the older approach (**Figure 3B** vs **Figure 3D)**. These data support **tracer**^**DART**^ as a behavior-independent metric of cellular target engagement.

### Whole-Brain Delivery

We next asked whether **DART.2** could lessen the need for cannulation directly into the investigated brain region. In preliminary experiments, we tested dosing via the cisterna magna, which showed promise with regard to rapid (∼30 min) bio-distribution to the cortex (Iliff et al., 2012). However, we ultimately favored cannulation of the lateral ventricle, which we found to be more consistent with regard to stereotactic coordinates and mechanical stability. Given that the ventricle is upstream of the cisterna magna, we began by characterizing ambient pharmacokinetics in ^**–**^**HTP** mice by implanting a cannula in one ventricle and glass window over contralateral visual cortex (**Figure 3E**). Infusion of **Alexa647.1**^**DART.2**^ into the ventricle produced an ambient fluorescence signal through the window, which peaked at ∼2 hr, and cleared by 24 hr (**Figure S3A**). The ambient signal was sustained and relatively constant over the 1-8 hr timeframe, suggesting distribution of the initial bolus over time.

Next, we expressed **AAV-DIO-**^**+**^**HTP**_**GPI**_ in the visual cortex of SOM-Cre mice for cell-specific expression in somatostatin interneurons (SOM cells), along with pan-neuronal **AAV-GCaMP8s** to enable recording of neural activity in all cells. We used the same cannula and window configuration, and presented oriented visual gratings to mice before and after infusion of 0.3 nmole **Alexa647.1**^**DART.2**^ + 3 nmole **YM90K.1**^**DART.2**^ (2 µL volume) into the contralateral ventricle. Several observations were made. First, we observed **Alexa647.1**^**DART.2**^ capture in the virus-expressing region, brighter than the neighboring ambient dye (**Figure 3F**, top right). Regarding tethered pharmacokinetics, dye capture plateaued after 4 – 8 hr. Animals with dense viral expression took the longest, and appeared to label ^**+**^**HTP** cells on the outer perimeter of expression before those in the center, suggesting ligand-limited dynamics. In animals with sparse expression, capture rose uniformly throughout the virus-expression region, suggesting that ambient ligand was in excess. The latter case provides an opportunity to estimate the ambient cortical concentration. In particular, sparse-expressing mice exhibited full capture at ∼4 hr (half-maximal at ∼2 hr), and thus given ***HTL***_**50**_ ∼ 300 nM × min (**Figure 1D**), we can estimate that ambient levels in the cortex remained in the low nM range (300 nM × min / 120 min = ∼2.5 nM drug + dye). This represents an upper bound of this estimate, as these mice exhibited the fastest capture kinetics.

With regard to functional effects, ^**+**^**HTP** SOM cells exhibited an attenuated response to visual gratings following drug delivery (**Figure 3G**, top; and **Figure S3B**), consistent with antagonized sensitivity to glutamate excitation. There was no evidence of off-target, ambient **YM90K.1**^**DART.2**^ effects: ^**–**^**HTP** cells were not attenuated at any timepoint (**Figure S3A-B**). This confirms that ambient concentrations stayed below ***Rx***_**50**_ in the visual cortex, and suggests a similarly safe ambient drug profile in distant brain regions, which provide the visual cortex with long-range afferents (Harris et al., 2019; Yao et al., 2021). Third, a subset of ^**–**^**HTP** cells showed an amplified response to visual gratings following **YM90K.1**^**DART.2**^ (**Figure 3G**, bottom; and **Figure S3B**). Because SOM cells are a major source of inhibition in the cortical microcircuit, disinhibition of neighboring cells is an expected finding, in qualitative agreement with optogenetic perturbations of SOM-cells (Adesnik, 2017; Kato et al., 2017; Lee et al., 2012). In contrast to these prior studies, **YM90K.1**^**DART.2**^ alters the glutamatergic connectome onto SOM-cells without directly perturbing their voltage―a key distinction to be elaborated upon in a companion study. Collectively, these findings represent a milestone with regard to dosing from a distance via the CSF.

### Positive Allosteric Modulator Design

Thus far, we succeeded in developing competitive antagonist reagents, which act via occlusive binding. We next sought to develop allosteric modulators, which must not only bind, but also induce the desired conformational change. For example, the benzodiazepine site of the GABA_A_R binds many drugs, ranging from negative allosteric modulation (NAM) to positive allosteric modulation (PAM). Given the diversity of allosteric possibilities at this site, we anticipated that a tether could disrupt the conformational impact of a given benzodiazepine. We began the endeavor with diazepam, a full-strength PAM, and the most clinically significant of the class. An atomic structure of the GABA_A_R bound to diazepam suggests that its amide is well positioned to provide access to free solution (Masiulis et al., 2019). We thus synthesized a diazepam analog with a short alkyne attached to this amide, and used click-chemistry to assemble **diazepam.1**^**DART.2**^ (**methods**). We next adapted our all-optical neuronal assay of endogenous GABA_A_Rs to maximize sensitivity for GABA_A_R PAMs (**methods**). We find that **diazepam.1**^**DART.2**^ exhibits pronounced cellular specificity (**Figure S4A**), with ***Rx***_**50**_ = 38 µM on ^**dd**^**HTP** neurons and ***HTL***_**50**_ = 50 nM × 15 min on ^**+**^**HTP** neurons, altogether yielding a ***TI***_**15m**_ = 760-fold. Notably, the maximum allostery of **diazepam.1**^**DART.2**^ appears equivalent to traditional diazepam (**Figure S4A**). Altogether, we observe a reduction in drug binding affinity, without a change in the allosteric impact once bound―a pattern reminiscent of our experience with antagonists.

The ease with which diazepam was developed into **diazepam**^**DART.2**^ was fortuitous, albeit unexpected. Given that diazepam is a full-strength PAM, we wondered whether partial or neutral allosteric drugs would be as amenable to development into a DART. In particular, flumazenil binds the same site as diazepam yet has neither positive nor negative allosteric impact once bound, providing a clinical antidote to diazepam overdose and a useful experimental reagent for local occlusion of systemic benzodiazepines. We developed **flumazenil.1**^**DART.2**^ via conjugation to an analogous amide of the flumazenil core and click-chemistry assembly (**methods**). To our surprise, **flumazenil.1**^**DART.2**^ became a full-strength PAM, with little resemblance to traditional flumazenil in our assay (**Figure S4B**). Though tentative, this example reveals a putative design principle wherein drugs that impart a maximal conformational change may be more likely to retain their original function upon tether attachment than those with partial or neutral conformational impact.

To complete the bidirectional toolkit, we explored AMPAR PAMs. For many pharmaceuticals of this class, including cyclothiazide, two drug molecules bind in close proximity in an inter-clamshell interface (Sun et al., 2002). We chose to focus on newer variants that bind both sites simultaneously, reasoning that a single tether may be easier to accommodate. We focused on CMPDA (phenyl-1,4-bisalkylsulfonamide), owing to availability of an experimental atomic structure and its potent allosteric effects, which are stronger than cyclothiazide (Timm et al., 2011). We devised a synthetic scheme to resynthesize CMPDA while installing a short PEG spacer domain at a point of attachment, predicted to allow tether access from the allosteric binding site to free solution (**methods**). The resulting **CMPDA.1**^**alkyne**^ retained its ability to positively modulate the AMPAR; however, its function was lost upon conversion to a full-length **CMPDA.1**^**DART**^ (**Figure S4C**). The reagent did not occlude other AMPAR PAMs, suggesting that the failure mode was a lack of binding rather than a change in allosteric efficacy once bound. Upon further inspection of the atomic structure, we noted that the path from the binding pocket to free solution had an overall negative surface potential, which we speculated could be repulsive to the electronegative properties of the PEG spacer. We thus performed computational docking of candidate spacers, of which one electropositive amine-rich design was developed into **CMPDA.2**^**DART.2**^ (**methods**). Consistent with intuition and docking studies, this design gained the ability to bind and also produce positive allosteric modulation of the AMPAR (**Figure S4D**).

### Bidirectional Editing of Chemical Neurotransmission

We next asked whether bidirectional modulation of the AMPAR and GABA_A_R could be achieved in the same cell type. We focused on a preparation in which electrophysiology is the primary readout, enabling resolution of the amplitude and millisecond time-constant of synaptic transmission. We selected the retina as the tissue of interest, given potential opportunities afforded by the whole-mount preparation, wherein the intact circuit can be studied under conditions amenable to electrophysiology (Masland and Raviola, 2000). We focused on a subset of retinal ganglion cells (RGCs) and leveraged a combination of viral serotype and mouse strain to genetically instruct parvalbumin-positive RGCs to express ^**+**^**HTP**_**GPI**_ or ^**dd**^**HTP**_**GPI**_ (**Figure 4A, methods**). We later prepared whole-mount retina and measured evoked synaptic currents under conditions that isolate the AMPAR or GABA_A_R (**methods**). For each cell, we established a stable baseline, then delivered a given **Rx**^**DART.2**^ (300 nM) + **Alexa647.1**^**DART.2**^ (30 nM) for 15 min, followed by washout. At the end of each experiment we imaged the **Alexa647** channel and observed robust capture on the surface of ^**+**^**HTP** cells, and no detectible capture on ^**dd**^**HTP** or ^**-**^**HTP** cells (**Figure 4B**).

**Figure 4.**
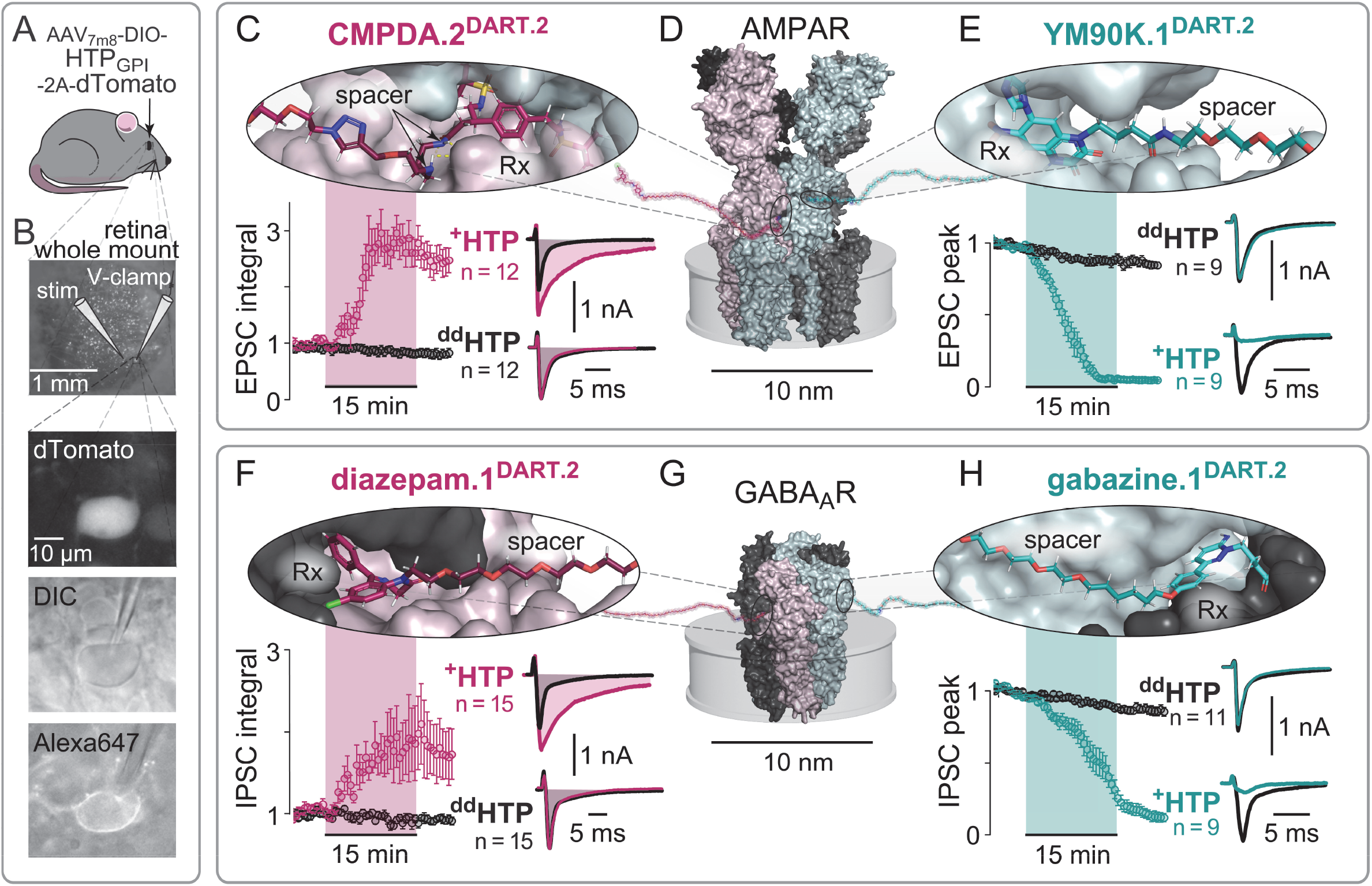
Bidirectional Editing of AMPAR and GABA_A_R Neurotransmission. **(A) Viral strategy**. Intravitreal AAV_7m8_-DIO-HTP_GPI_-2A-dTomao achieves retinal ganglion cell (RGC) expression in PV::cre mice. **(B) Experimental overview**. Top: whole retina; synaptic current is evoked (stim) and recorded (voltage-clamp of dTomato+ RGC), with ionic and pharmaceutical conditions chosen to isolate the AMPAR or GABA_A_R (**methods**). A 10 min baseline is followed by 15 min application of 300 nM **Rx**^**DART.2**^ + 30 nM **Alexa647.1**^**DART.2**^. After wash, tethered dye (bottom panel) is seen on ^**+**^**HTP** but not _**dd**_**HTP** cells. **(C) CMPDA.2**^**DART.2**^. Top: electropositive spacer (PDB 3RNN, design elaborated in **Figure S4D**). Bottom left: AMPAR evoked EPSC integral (total charge transfer) is boosted 3-fold on ^**+**^**HTP** but not _**dd**_**HTP** cells (mean ±SEM, cells normalized to baseline). Bottom right: EPSC waveform before (gray) and after (pink) **CMPDA.2**^**DART.2**^ (summary of EPSC peak and time constant in **Figure S4F**). **(D)** AMPAR structural model (composite of PDB 5WEO, 3RNN, and 1FTL). Cylinder depicts cell membrane. **CMPDA.2**^**DART.2**^ binds within an inter-clamshell interface (2 per receptor). **YM90K.1**^**DART.2**^ binds the orthosteric glutamate binding site in the clamshell (4 per receptor). **(E) YM90K.1**^**DART.2**^. Top: structural model (PDB 1FTL). Bottom left: AMPAR evoked EPSC peak is blocked on ^**+**^**HTP** cells but not on _**dd**_**HTP** cells (mean ±SEM, cells normalized to baseline). Bottom right: EPSC waveforms before (gray) and after (cyan) **YM90K.1**^**DART.2**^. **(F) diazepam.1**^**DART.2**^. Top: structural model (PDB 6HUP). Bottom left: GABA_A_R evoked IPSC integral (total charge transfer) is boosted 2-fold on ^**+**^**HTP** but not _**dd**_**HTP** cells (mean ±SEM, cells normalized to baseline). Bottom right: IPSC waveform before (gray) and after (pink) **diazepam.1**^**DART.2**^ (summary of IPSC peak and time constant in **Figure S4E**). **(G)** GABA_A_R structural model (composite of PDB 6HUP and 6HUK). Cylinder depicts cell membrane; **diazepam.1**^**DART.2**^ binds the α/δ interface (1 per receptor). **gabazine.1**^**DART.2**^ binds the orthosteric GABA-binding site in the α/β interface (2 per receptor). **(H) gabazine.1**^**DART.2**^. Top: structural model (PDB 6HUK). Bottom left: GABA_A_R evoked IPSC peak is blocked on ^**+**^**HTP** cells but not on _**dd**_**HTP** cells (mean ±SEM, cells normalized to baseline). Bottom right: IPSC waveforms before (gray) and after (cyan) **gabazine.1**^**DART.2**^.

Starting with PAM reagents, **CMPDA.2**^**DART.2**^ strongly potentiated AMPAR-mediated EPSC (excitatory post-synaptic current) on ^**+**^**HTP** RGCs, both by augmenting the peak amplitude and by prolonging the decay time (**Figure S4F**), altogether yielding a three-fold increase in the integrated EPSC charge transfer (**Figure 4C**, pink). No impact of **CMPDA.2**^**DART.2**^ was seen in analogous experiments in ^**dd**^**HTP** RGCs (**Figure 4C**, black). We next evaluated the impact of **diazepam.1**^**DART.2**^ on the GABA_A_R-mediated IPSC (inhibitory post-synaptic current). We observed that this reagent significantly potentiated IPSCs, with an analogous increase in peak amplitude and prolongation of decay time (**Figure S4E**). Again, the effects were only seen on ^**+**^**HTP** RGCs, with no effect on ^**dd**^**HTP** RGCs (**Figure 4F**). Finally, we confirmed that **gabazine.1**^**DART.2**^ blocked the GABA_A_R-mediated IPSC (**Figure 4H**), and that **YM90K.1**^**DART.2**^ blocked the AMPAR-mediated EPSC (**Figure 4E**). With all reagents, maximal effects took hold within 15 min for ^**+**^**HTP** RGCs, and no effects were seen on ^**dd**^**HTP** RGCs.

## DISCUSSION

Here, we describe an optimized technology for cell-specific pharmacology, offering the efficiency and technical checkpoints needed for safe delivery of diverse pharmaceuticals in a breadth of experimental contexts. In particular, we provide a toolkit for bidirectional modulation of the AMPAR and GABA_A_R, which transduce the majority of rapid synaptic neurotransmission between brain cells. These reagents set the stage for an emerging kind of causal neuroscience experiment, one that modulates how a cell listens to a given chemical dialect. In particular, cells can now be made less or more sensitive to glutamate inputs. They can alternately be made less or more responsive to incoming GABA signals. Whether these chemically discrete signals encode behaviorally distinct variables―and how these four manipulations (scaling neural sensitivity to transmitter input) will compare to the traditional two (adding vs subtracting neural output)―represents a frontier in causal neuroscience.

### Technology Comparison

The study of endogenous receptors (e.g., AMPAR and GABA_A_R) has historically favored genetic knockouts. However, the weeks needed for protein turnover can lead to compensatory false negatives, or over-compensation with paradoxical behavioral effects (Bailey et al., 2006). For faster knockouts, acute protein degradation is a promising avenue (Verma et al., 2020). This was achieved for gephryn, a scaffolding protein, which when degraded causes redistribution of GABA_A_Rs to extrasynaptic locations while permitting their continued response to GABA (Gross et al., 2016). We see these as complementary tools that address distinct questions. In particular, redistribution of GABA_A_Rs differs from their antagonism (e.g., gabazine). Moreover, elimination of proteins cannot foreseeably mimic allosteric modulation (e.g., diazepam), nor distinguish ionotropic from structural signaling roles (Li et al., 2016; Nabavi et al., 2013; Zachariassen et al., 2016).

Regarding cell-specific pharmacology, prodrugs can be enzymatically unmasked to yield active drug within the confines of the cytoplasm (Gruber et al., 2018; Tian et al., 2012; Yang et al., 2015), offering promise for intracellular pharmacology. For extracellular pharmacology, tethering is needed to counteract diffusion. This was first achieved via chemical attachment to a fortuitous cysteine residue on a native receptor (Lester et al., 1980), and later via deliberate placement of a cysteine into many other receptors (Banghart et al., 2004; Barber et al., 2016; Berlin et al., 2016; Broichhagen et al., 2015; Broichhagen and Levitz, 2022; Chambers and Kramer, 2008; Donthamsetti et al., 2017; Fortin et al., 2011; Janovjak et al., 2010; Kramer et al., 2005; Levitz et al., 2017; Levitz et al., 2013; Levitz et al., 2016; Morstein et al., 2022; Reiner et al., 2015; Sandoz et al., 2012; Tochitsky et al., 2012; Volgraf et al., 2006). Viral delivery of these tools has been used to add exogenous receptors to a defined cell type (Berry et al., 2017; Szobota et al., 2007; Wyart et al., 2009), while leaving alone endogenous receptors that lack the cysteine. Conversely, cysteine knock-in into the genome enables tethering to endogenous receptors (Lin et al., 2015), albeit this has typically sacrificed cell specificity.

The advance provided by **DART** is that drugs are tethered to cells, not receptors themselves (Shields et al., 2017). This provides cell specificity while recapitulating the modus operandi of traditional pharmacology, wherein the endogenous receptor of interest is neither virally overexpressed, nor manipulated in the genome via knock-in. An additional advantage is modularity of the reagents, wherein a universal **HTP** virus can select the cell type of interest, enabling later selection of any **Rx**^**DART**^ chemical, which specifies the receptor and pharmaceutical class of the manipulation. As with traditional pharmacology, the rapid onset of drug capture enables acute, causal relationships to be established. Thereafter, covalent attachment persists for days, enabling chronic therapeutic efficacy to be assessed. If desired, re-dosing can be performed to extend therapeutic duration, or to deliver an entirely new pharmaceutical class, all in the same individual animal.

### Community Guidelines for Extending DART

To facilitate community design, ‘**Rx** fragments’ are made separately from ‘**HTL** modules’ for later assembly via *alkyne-azide* click chemistry (Rostovtsev et al., 2002). Here, we outline design standards and define a naming and numbering convention to uniquely identify fragments, modules, and their co-assembled counterparts.

With regard to **Rx** fragment design, the ideal ***Rx***_**50**_ must balance potency and cellular specificity. In three of four pharmaceutical classes, our first design attempt yielded an acceptable ***Rx***_**50**_ in the µM range (**YM90K.1**^**DART.2**^ ***Rx***_**50**_ = 6 µM, **gabazine.1**^**DART.2**^ ***Rx***_**50**_ = 12 µM, **diazepam.1**^**DART.2**^ ***Rx***_**50**_ = 30 µM). We decided to attenuate one of these further given epileptogenic concerns (**gabazine.7**^**DART.2**^ ***Rx***_**50**_ = 180 µM). We experienced one failure with CMPDA, wherein the electronegative spacer in **CMPDA.1**^**DART**^ prevented AMPAR binding, and the electropositive spacer in **CMPDA.2**^**DART.2**^ restored binding in the µM range.

A second parameter, separate from affinity, is conformational impact of a drug once bound. We observed an example in which a neutral allosteric drug (**flumazenil.1**^**DART.2**^) changed its functional identity upon tether addition, while its full-strength PAM counterpart (**diazepam.1**^**DART.2**^) remained true to form. In many biological processes, pushing a parameter to its saturation can yield resilience to perturbation. In this case, allosteric strength is the parameter of interest, and we speculate that full-strength allosteric drugs may be more resilient to a tether than their partial-allosteric counterparts. Consistent with this, CMPDA, a full-strength PAM for the AMPAR, also retained its functional identity so long as the tether permitted binding to occur. Additional examples will be needed to deepen this intuition and assess applicability to other drug classes.

Third, we previously established the requirement for a long chemical linker between **Rx** and **HTL**. In particular, PEG_12_ was too short to reach the AMPAR, whereas PEG_24_ and PEG_36_ were both acceptable (Shields et al., 2017). Since then, PEG_36_ has sufficed for all receptors tested. Given its common utility, the long linker is grouped into the ‘**HTL** module’ (e.g., *azide*^DART.2^ contains *azide*-**PEG**_**36**_-**HTL.2**), enabling the majority of investigators to focus on developing small **Rx**^*alkyne*^ reagents with traditional drug-like properties (**Figure 1B**).

Finally, we advise that the name of an **Rx** fragment should uniquely identify its structure, with the convention of including the name of its commonly known precursor followed by a version number that defines conjugation and spacer details (e.g., **gabazine.7**^*alkyne*^). In the interest of readability, we settled on consistent use of a period to delimit the version number, as many precursor names contain numbers (e.g., **Alexa647.1**^*alkyne*^). After the final *alkyne-azide* assembly step, the resulting full-length reagent should retain both version numbers for clarity (e.g., **gabazine.7**^*alkyne*^ + *azide*^DART.2^ → **gabazine.7**^DART.2^). In sum, we envision that distributed teams of scientists will be able to grow the platform in a manner that is organized, collaborative, and scalable.

### Thousandfold Cell-Specificity Insights

The core advance with **DART.2** is improved cellular specificity, achieved via optimization of **HTL** and **Rx** moieties. Each had impact in the expected direction, albeit with two surprises. First, improving ***HTL***_**50**_ had no impact on ***Rx***_**50**_ (as expected). However, the 40-fold ***HTL***_***50***_ improvement seen with a biotin payload (**Figure 1A**), became a 10-fold ***HTL***_***50***_ benefit with YM90K (**Figure 1C-D**). We suspect this discrepancy can be attributed to an avidity phenomenon with **YM90K.1**^DART.1^ (Wilhelm et al., 2021). In particular, because **HTL.1** is inefficient, transient interactions mediated by YM90K can assist the HTL moiety, enabling capture of **YM90K.1**^DART.1^ to outperform **HTL.1** on its own. Avidity becomes negligible with **YM90K.1**^DART.2^, wherein **HTL.2** singlehandedly drives capture before YM90K can assist. Thus, avidity is unequal in the two cases. To compare **HTL.1** vs **HTL.2** fairly, a biotin payload is preferred owing to its consistent, lack of interaction with cells over a broad dosing range.

Next, with regard to **Rx** optimization, **gabazine.7**^**DART.2**^ requires 15-times as many molecules in the ambient (***Rx***_**50**_), yet surprisingly only 2-fold more when tethered (***HTL***_**50**_) in comparison to **gabazine.1**^**DART.2**^. This observation runs counter to avidity and could not be explained with binding saturation. We thus considered pharmaceutical ligand depletion, which is known to arise as volume is miniaturized (Carter et al., 2007). We began with a model wherein the length of an extended DART (∼16 nm) constrains the ‘nanodomain’ volume over which a GABA_A_R can interact with tethered drug (**Figure S1A**). Since this volume is tiny (∼9 × 10^−21^ L), there would be few drug molecules in a nanodomain, each contributing a substantial concentration (∼180 µM). In this parameter regime, one must account for each individual drug. In particular, only GABA_A_R-unbound drugs contribute to the mobile nanodomain concentration, as only these can collide to drive new binding events. Conversely, only GABA_A_R-bound drug can modulate the channel, but at the price of depleting the mobile concentration. Because depletion is a subtractive phenomenon (governed by the number of binding sites on the GABA_A_R), replenishing the depletion would require an additive (not multiplicative) right-shift in the dose-response. As such, ***HTL***_**50**_ would be impacted most for **gabazine.1**^**DART.2**^, narrowing the gap to **gabazine.7**^**DART.2**^ on the logarithmic dosing axis (**Figure S1A**).

Given that ligand depletion is accentuated as volume shrinks, we asked whether our dataset could provide an experimental constraint on the nanodomain volume. Indeed, we find that the 16-nm hemispherical radius, estimated from molecular structure, recapitulates the 2-fold change in ***HTL***_**50**_ data across **gabazine**^**DART**^ variants (**Figure S1B-D**). If the nanodomain volume were increased, the fold-change in ***HTL***_**50**_ across variants would broaden, approaching the 15-fold change seen in the ambient. The opposite is true of smaller nanodomains, which would restrict ***HTL***_**50**_ more tightly than seen in the data. Thus, the volume parameter is well constrained. The model also explains why tethered **gabazine.1**^**DART.2**^ achieves 97% block, whereas **gabazine.7**^**DART.2**^ achieves ∼72% block at saturation. Both findings are consistent with a stoichiometry of ∼4 **HTP** proteins per nanodomain (**Figure S1A**). This number sets the maximum concentration in the nanodomain, which we estimate can be as high as ∼720 µM (= 4 × 180 µM) if all 4 **Rx** molecules remain mobile. For high-affinity gabazine variants, up to 2 **Rx** would be depleted upon binding the GABA_A_R, leaving ∼360 µM (= 2 × 180 µM) mobile. These estimates provide a conceptual framework to better understand, design, and use **DART** reagents.

### Reagents for Rigor and Reproducibility in Behaving Animals

With regard to receptor specificity, high tethered **Rx** concentrations run the risk of producing unforeseen receptor interactions on ^**+**^**HTP** cells. Interestingly, the ligand depletion phenomenon (**Figure S1**) could mitigate this concern to some extent, as the desired receptor could outcompete off-target sites for a limited number of drugs. Nevertheless, given the unknowns of receptor density and drug binding in the tethered regime, we recommend experimental characterization in the cell type of interest. For example, we confirmed that **gabazine.7**^**DART.2**^ blocks evoked GABA_A_R current onto ^**+**^**HTP** VTA_DA_ neurons, and that the same amount of tethered **gabazine.7**^**DART.2**^ has no impact on NMDA or AMPA-mediated transmission, nor impact on other measurable electrophysiological parameters, including pacemaker firing rate or action potential waveform in these cells (**Figure S2D-G**). As an additional control, we developed a **blank.1**^**DART.2**^, with neither drug nor dye (**methods**), and used this to confirm that there is no pharmaceutical impact of the **Alexa647.1**^**DART.2**^ tracer or impact of chemical accumulation on the surface of a neuron (**Figure S2H**). While this example provides a template for reagent validation, the characterization of each specific reagent will require community engagement in diverse biological contexts.

In addition to providing cellular specificity, **DART.2** resolves two technical issues that have plagued neuropharmacology. First, the ^**dd**^**HTP** reagent makes it possible to rule out effects of the ambient drug, and in so doing eliminates the nagging concern that effects may be mediated by an unanticipated receptor in an untested cell type. This allows characterization to focus on ^**+**^**HTP** cells, as delineated in the example above. Second, the cell-by-cell **tracer**^**DART.2**^ signal can be used to calibrate dosing across preparations―whether drug was infused into a live animal or onto a brain slice. This ensures that slice characterizations on ^**+**^**HTP** cells are matched to those in a live animal. Together the reagents provide a new standard of rigor, wherein the ^**dd**^**HTP** reagent allows one to rule out over-dosing (false-positive) behavioral effects, while **tracer**^**DART.2**^ allows one to rule out under-dosing (false-negatives). As such, **DART.2** can support or disprove hypotheses without ambiguity.

### Future Directions

Preclinical and clinical pharmacology studies depend on reliable target engagement. To this end, the tracer approach described here can serve as a template for near-infrared dyes and MRI contrast agents, which enable real-time deep-tissue observation in living animals. With regard to the efficiency of drug capture, we achieved a milestone with whole-brain dosing from a distance via the CSF. A future goal is to achieve safe, cell-specific delivery over the whole brain and body without the need for intracranial infusion. Capture efficiency improvement continues to be a promising avenue. Branched ligands (Acosta-Ruiz et al., 2020) could improve stoichiometry by delivering multiple drugs per tether. The PEG_36_ linker may be amenable to added functionality to promote blood-brain-barrier crossing and membrane permeation to access intracellular targets. Additional features of interest include delivery of two or more drugs to distinct cellular populations, subcellular pharmacology, and rapid reversibility. In particular, photo-controlled versions of DART are possible (Donthamsetti et al., 2021; Donthamsetti et al., 2019), and likely of great interest for questions that require millisecond precision over small brain volumes. Conversely, light-independent reversibility would simplify use over large brain volumes, facilitating large-animal and clinical studies. Finally, extending the method to diverse clinical pharmaceuticals―and the conceptual questions now within reach―remain areas of untapped promise.

## STAR ★ METHODS

Detailed methods are provided in the online version of this paper and include the following:

- KEY RESOURCES TABLE
- RESOURCE AVAILABILITY
- METHOD DETAILS

## SUPPLEMENTAL INFORMATION

Supplemental information includes the following:

- SUPPLEMENTARY **FIGURES S1-S4**
- Detailed Author Contributions (**Table S1**)
- Chemical Synthesis (**Supplementary Document 1**)

## ACKNOWLEDGEMENTS

We would like to thank Victoria Z. Goldenshtein for piloting of binding assays in recombinant protein; James M. Roach for technical support in open field behavioral assays and data processing; S. Aryana Yousefzadeh for piloting of intraventricular cannulation strategies; Isaac A. Weaver for development of a web-based distribution platform and chemical depictions; Tom Haimowitz for synthesis and co-design of the first batch of *azide*^DART.1^; Sandeep Thana for support with synthesis of CMPDA.2^DART.2^; Javier Izquierdo-Ferrer for contributing to the methods and partial synthesis of gabazine^DART^ reagents; Ankit Choudhury and Jiyong Hong for insightful feedback on the manuscript. This work was supported by Duke University Startup Funds (to CH, GDF, LLG, and MRT), Duke University Holland Trice Scholars Award (to LLG), Duke University DIBS Awards (to MRT), and by NIH grants 1F3-1NS113742-01A1 (to EAF); R01-NS096289 and R01-NS112917 (to CH); R34-NS111645 and R01-EY031396 (to GDF); R01-EY031716 (to LLG); and RF1-MH117055, DP2-MH1194025, R01-NS107472, and R61-DA051530 (to MRT).

## AUTHOR CONTRIBUTIONS

See **Table S1** for detailed author contributions. **Conceptualization**―BCS, HY, SSXL, SCB, EAF, CMC, TMH, CH, GDF, LLG, and MRT. **Methodology**―BCS, HY, SSXL, SCB, EAF, CMC, EWK, PPV, MHL, LZ, MEM, MLS, TMH, MT, ABR, GES, CH, GDF, LLG, and MRT. **Software**―SCB and MRT. **Validation**―BCS, HY, SSXL, SCB, EAF, CMC, EWK, MHL, MEM, MLS, TMH, MT, and MRT. **Formal Analysis**―BCS, HY, SCB, EAF, CMC, PPV, LZ, GES, and MRT. **Investigation**―BCS, HY, SSXL, SCB, EAF, CMC, EWK, PPV, MHL, LZ, TMH, MT, and MRT. **Resources**―ABR, GES, CH, GDF, LLG, and MRT. **Data Curation**―BCS, HY, SCB, EAF, CMC, EWK, and MRT. **Writing, Original Draft**―BCS, HY, SSXL, PPV, LZ, and MRT. **Writing, Review and Editing**―BCS, HY, SSXL, SCB, EAF, CMC, PPV, GES, CH, GDF, LLG, and MRT. **Visualization**―BCS, HY, SSXL, SCB, EAF, CMC, and MRT. **Supervision**―BCS, CMC, MEM, ABR, GES, CH, GDF, LLG, and MRT. **Project Administration**―BCS, EWK, ABR, GES, CH, GDF, LLG, and MRT. **Funding Acquisition**―EAF, CH, GDF, LLG, and MRT.

## STAR **★** METHODS

### KEY RESOURCES TABLE

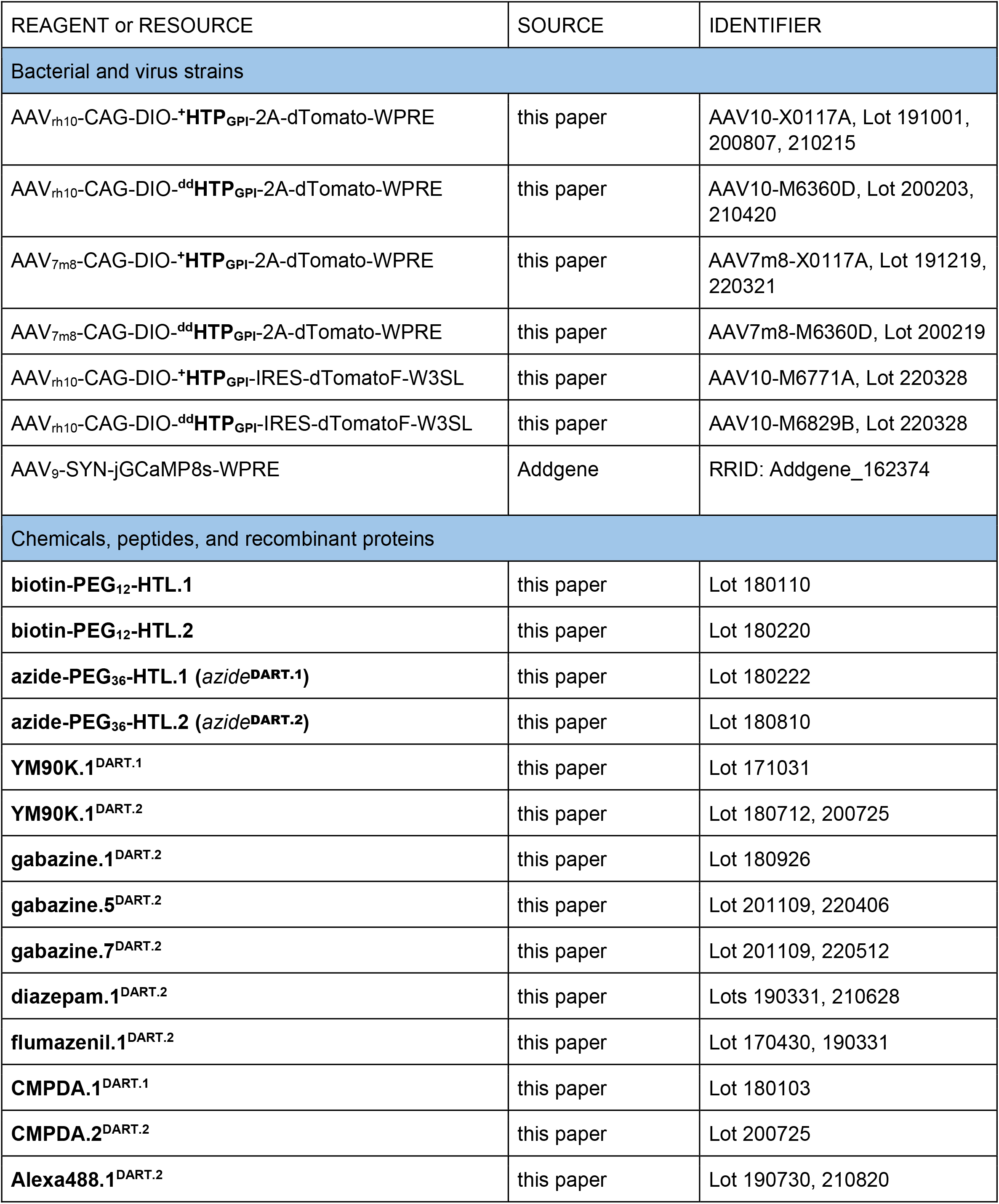
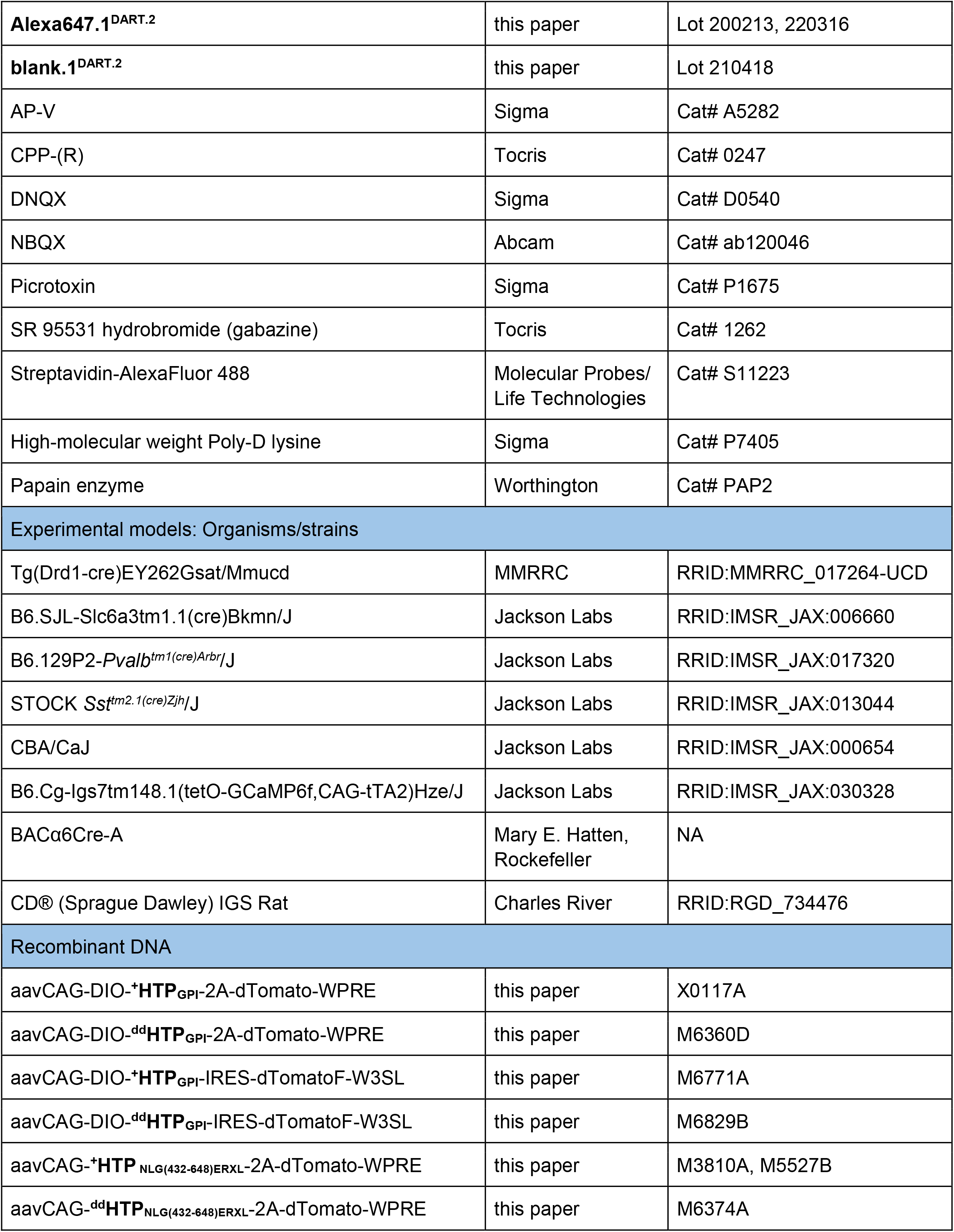
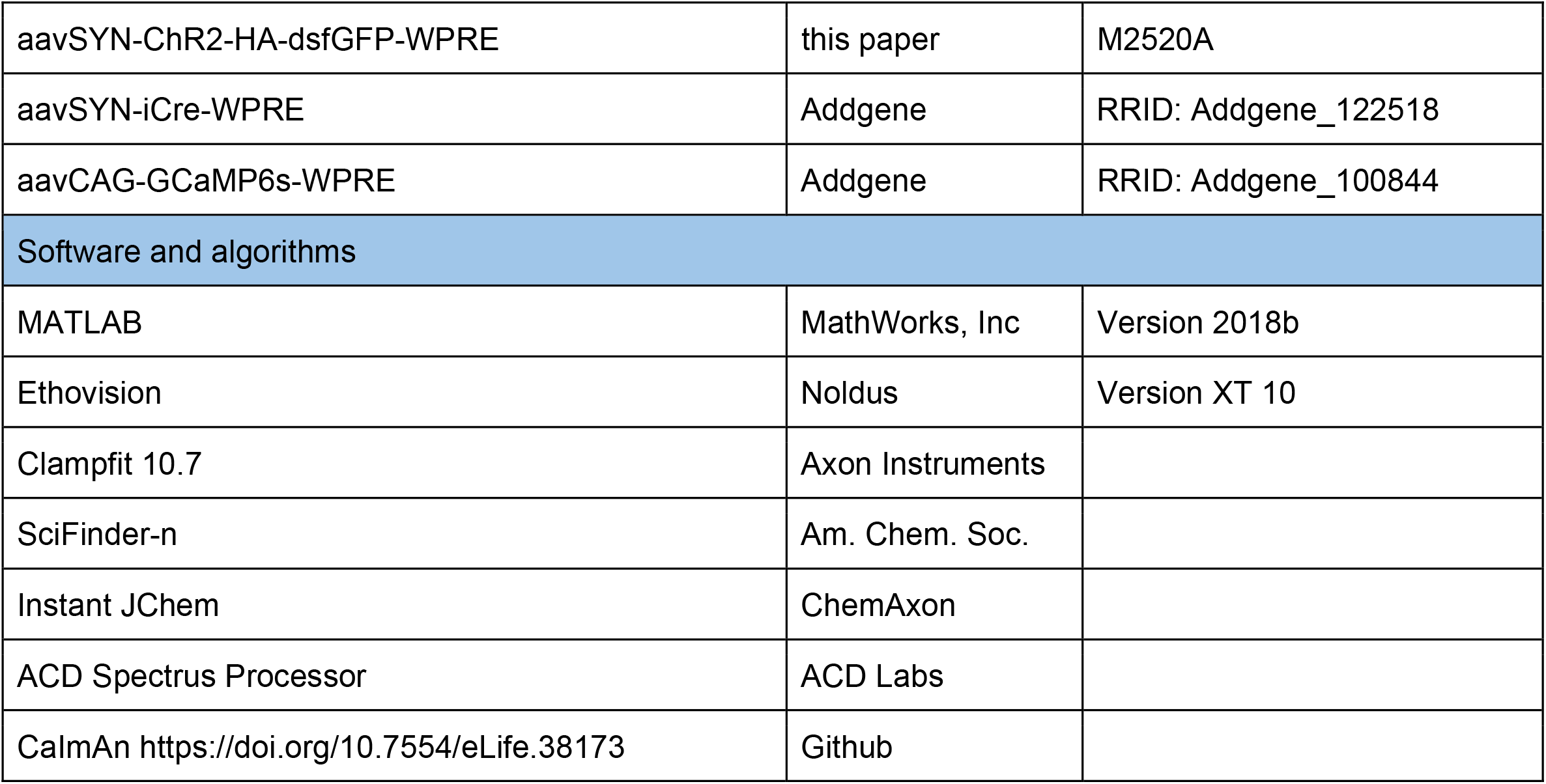

### CONTACT FOR REAGENT AND RESOURCE SHARING

Further information and requests for resources and reagents should be directed to and will be fulfilled by the Lead Contact Michael R. Tadross, MD, PhD.

### EXPERIMENTAL MODEL AND SUBJECT DETAILS

All experiments involving animals were approved by the Duke Institutional Animal Care and Use Committee, an AAALAC accredited program registered with both the USDA Public Health Service and the NIH Office of Animal Welfare Assurance, and conform to all relevant regulatory standards.

#### Rats

Timed-pregnant female Sprague-Dawley rats (Charles River) were individually housed in a standard temperature and humidity environment, under a normal 12-hr light/dark cycle and with food and water provided *ad libitum*.

#### Mice

Drd1A-Cre (GENSAT EY262, striatum D1 medium spiny neuron), DAT-*Ires*-cre (Jackson Labs 006660, VTA dopamine neuron), SST-Cre (Jackson Labs 013044, V1 SOM interneuron), PV-Cre (Jackson Labs 017320, retinal ganglion cell), BACα6Cre-A (Hatten Lab, Rockefeller, cerebellar granule cell), and Ai148D (Jackson Labs 030328, GCamp6f). Mice were group housed by age and gender (max 5 per cage) in a standard temperature and humidity environment, under a normal 12-hr light/dark cycle and with food and water provided *ad libitum*.

## METHOD DETAILS

### Genetic Construct Design

Genetic elements were concatenated via PCR, restriction digest, ligation, and sequence verification, as follows:

- aavCAG-DIO-^**+**^**HTP**_**GPI**_-2A-dTomato-WPRE. Coding: SS_nlg_-^+^HTP-(GGSGG)_8_-Thy1_GPI_-2A-dTomato. SS_nlg_, the signal peptide (residues 1–49) of mouse neuroligin-1; ^+^HTP, mammalian codon-optimized variant of the HaloTag Protein (Los *et al*., 2008); (GGSGG)8, linker with 8 repeats of gly-gly-ser-gly-gly; Thy1_GPI_ from Addgene_163696. Marker of Expression: 2A-dTomato, P2A ribosomal skip sequence and dTomato. Backbone: aavCAG-DIO-WPRE vector is from Addgene_100842.
- aavCAG-DIO-^**dd**^**HTP**_**GPI**_-2A-dTomato-WPRE. Coding: SS_nlg_-_dd_HTP-(GGSGG)_8_-Thy1_GPI_-2A-dTomato.
- _dd_HTP is the HTP with N41E, D106E, W107G, V245L, and L246R mutations. Other elements as above.
- aavCAG-DIO-^**+**^**HTP**_**GPI**_-IRES-dTomatoF-W3SL. Coding: SS_nlg_-^+^HTP-(GGSGG)_8_-Thy1_GPI_-IRES-dTomatoF.
- IRES is the internal ribosomal entry site. dTomatoF is dTomato followed by the Farnesylation sequence KLNPPDESGPGCMSCKCVLS. W3SL is from Addgene_61463. Other elements as above.
- aavCAG-DIO-^**dd**^**HTP**_**GPI**_-IRES-dTomatoF-W3SL. Coding: SS_nlg_-_dd_HTP-(GGSGG)_8_-Thy1_GPI_-IRES-dTomato. Elements all as above
- aavSYN-ChR2-HA-dsfGFP-WPRE. Coding: ChR2(H134R)-HA-dsfGFP. ChR2(H134R) is the optogenetic channelrhodopsin2; dsfGFP is superfolder GFP with T65G and Y66G mutations to eliminate fluorescence. Backbone: aavSYN-WPRE vector is from Addgene_100843.
- aavCAG-^**+**^**HTP**_**NLG(432-648)ERXL**_-2A-dTomato-WPRE. Coding: SS_nlg_-HA-^+^HTP-NLG(432-648)-ERXL.
- HA, the hemagglutinin epitope tag; NLG(432-648), the esterase-truncated 71-residue extracellular domain, the 19-residue predicted transmembrane domain, and the 127-residue C terminus of mouse neuroligin-1 (Kim et al., 2012); ERXL, the peptide sequence KSRITSEGEYIPLDQIDINVGGSGFCYENEV, a fusion of the trafficking and ER export signals from Kir2.1 (Gradinaru et al., 2010). Other elements as above.
- aavCAG-^**dd**^**HTP**_**NLG(432-648)ERXL**_-2A-dTomato-WPRE. Coding: SS_nlg_-HA-_dd_HTP-NLG(432-648)-ERXL.Elements all as above

### Chemical Synthesis

See Supplementary Document 1.

### Hippocampal Cultured Neurons

Mixed glial and neuronal cultures were prepared from the hippocampus of postnatal day 0 to 1 Sprague-Dawley rat littermates as previously described (Shields et al., 2017). Pups were decapitated and brains quickly excised into ice-cold neural dissection solution (NDS), consisting of Hepes-buffered HBSS, pH 7.4 (Sigma, H3375; Gibco, 24020-117). Hippocampal tissue was dissected and transferred to a separate petri dish containing ice-cold NDS, quartered, and washed several times with fresh ice-cold NDS. The hippocampal pieces were collected and incubated in papain enzyme (Worthington, PAP2; 105U per prep in NDS) in a 37 °C water bath for 25-35 min, inverted to mix twice during incubation. The papain solution was decanted, and the tissue was washed 3 times in plating medium (PM), consisting of MEM (Gibco, 51200-038) with 10% (v:v) heat-inactivated FBS (HyClone, SH30071.03), and 27.8 mM glucose, 2.4 mM NaHCO_3_, 0.1 mg/mL transferrin, 0.025 mg/mL insulin, and 1% (v:v; 2 mM) L-glutamine plus 1% (v:v) Penicillin/ Streptomycin. The tissue was further dissociated via mechanical trituration through a 10-mL serological pipette, followed by two fire-polished glass pipettes, and the final cell suspension was filtered through a 0.22 µm cellulose acetate membrane.

Cells were nucleofected (LONZA, V4SP-3096) with a selection of high-quality plasmid constructs (0.8 -1 µg dna per cuvette). After a 10 min recovery period, cells were plated individually or mixed with a different pool of cells (for co-cultures). Cells were cultured on coverslips (Deckglaser) pre-treated with high molecular weight poly-D-lysine (Sigma, P7405; 0.05 mg/mL) in 24-well plates and maintained in NbActiv4 (BrainBits, NB4) at 37 °C, 5% CO2. Media was changed by half at DIV3 or DIV4 and then weekly thereafter; experiments were performed between DIV16 to DIV18.

For any given neuronal assay, coverslips were first rinsed briefly in a resting Tyrode solution containing (in mM): 4 KCl, 2 CaCl_2_, 2 MgCl_2_, 150 NaCl, 10 HEPES, 10 glucose, pH 7.4, before transfer to a glass-bottom, 24-well imaging plate (Cellvis, P24-1.5H-N). Live cell imaging was performed on an Olympus IX83 inverted fluorescent microscope (UPlanSApo 10X objective, NA 0.40) with Spectra X LED illumination and filter cubes as follows:

|  | <u>LED (nm)</u> | <u>Excitation (nm)</u> | <u>Dichroic (nm)</u> | <u>Emission (nm)</u> |
| --- | --- | --- | --- | --- |
| • **GFP** | 470/24 | 472/30 | 495 | 520/35 |
| • **TRITC** | 550/15 | 543/22 | 562 | 593/40 |
| • **Cy5** | 640/30 | 628/40 | 660 | 692/40 |

### Biotin-HTL Assay

Neurons were nucleofected with aavCAG-^+^HTP _**NLG(432-648)ERXL**_-2A-dTomato-WPRE, plated and cultured for 2 weeks. A dilution series was made for the **biotin-PEG**_**12**_**-HTL.1** and **biotin-PEG**_**12**_**-HTL.2** compounds in Tyrode solution containing 1% BSA, representing 12 final concentrations (in µM) between 10 and 0.001. After a brief rinse in Tyrode +BSA to acclimate the live neurons to room temperature. Each coverslip was incubated in one of the twelve dilutions for 15 min, washed twice in Tyrode +BSA, then incubated in a 1:500 dilution of Streptavidin-AlexaFluor488 (2 mg/mL; Life Technologies, S11223) for 15 min. The neurons were rinsed four times then transferred to a glass-bottom imaging plate containing Tyrode +BSA. Each coverslip was imaged in its entirety by stitching together several 10X images (TRITC at 10ms; GFP at 100ms) using the Olympus CellSens software stitching algorithm. A dose-response curve was created using a custom MATLAB algorithm, involving two steps: segmentation, and estimation of the fraction of HTP bound to HTL, as follows.

Regarding segmentation, we used an automated method given the number of cells per coverslip (thousands). We used the red dTomato channel (genetic expression) for segmentation, reserving the green streptavidin channel (surface HTL) as the readout. The dTomato intensity was first normalized to a 0 to 1 scale, with gamma = 0.4 to enable detection of dim cells. We performed adaptive background subtraction (90 pixel-radius), applied a fixed threshold = 0.15 to obtain a binary mask, applied an open radius of 9 pixels, and detected regions with the ‘regionprops’ command from the MATLAB Image Processing Toolkit. This typically yielded >1000 regions per coverslip. The quality of segmentation was confirmed by visual inspection; ∼80% of regions corresponded to individual neuronal cell bodies, with the remaining being cell clusters or processes. For each region, we calculated the mean dTomato (red, R) and streptavidin (green, G) intensities.

With regard to estimating the fraction of HTP bound to HTL, data from coverslips that had been incubated in high-dose biotin-PEG_12_-HTL.2 (≥ 300 nM × 15 min) appeared to be fully saturated, and were used to estimate the relationship between red (R) and green (G), which fit well to G = ***G***_**max**_ × R / (R + ***R***_**half**_). We interpret this equation to reflect surface trafficking of HTP, which saturates as genetic expression rises. Because all coverslips used the identical genetic construct, we took the estimates of ***G***_**max**_ and ***R***_**half**_ from the saturated coverslips as a constant (held fixed for all coverslips). We then fit data from each coverslip to the equation G = ***HTP***_**FracBound**_ × ***G***_**max**_ × R / (R + ***R***_**half**_). We used the ‘fit’ command from the MATLAB Curve Fitting Toolbox, which provides a nonlinear regression estimate of the only free parameter, ***HTP***_**FracBound**_. This estimate includes a 95% confidence interval, which we plot as the error bars in **Figure 1A**.

### All-optical Cultured-Neuron AMPAR Assay

Two pools of dissociated neurons were nucleofected separately to express either:

1. aavSYN-ChR2-HA-dsfGFP-WPRE (**ChR2**)
2. aavCAG-^+^HTP _**NLG(432-648)ERXL**_-2A-dTomato-WPRE + aavCAG-GCaMP6s-WPRE (**HTP/GC**)

The ChR2 transfected cells were then mixed equally with the HTP/GC cells, plated, and cultured for 16-18 days. The assay was performed in Tyrode (described above) supplemented with 10 µM CPP (NMDAR antagonist) and 10 µM gabazine (GABA_A_R antagonist). This recipe isolates the AMPAR, and is thus named Tyrode^AMPAR^. The assay is synaptic and AMPAR-specific because we ensure that ChR2 and GCaMP6s are expressed in separate cells, such that light must first stimulate ‘presynaptic’ ChR2 neurons to release glutamate, which then activates ‘postsynaptic’ HTP/GC neurons only via the AMPAR (isolated via Tyrode^AMPAR^). Thus, light-evoked HTP/GC activity is a proxy for postsynaptic AMPAR function.

We performed assays 12 coverslips at a time. Plates were removed from the incubator, coverslips briefly rinsed in Tyrode^AMPAR^ and transferred to a glass-bottom imaging plate with 500 µL of Tyrode^AMPAR^ per well. A parallel drug plate with 500 µL Tyrode^AMPAR^ per well was used to add reagents via manual pipetting. Between dosing rounds, each well of the drug plate was mixed with its corresponding well in the imaging plate (add, gently triturate once, remove). Thus, the imaging plate maintains a set volume, and we have independent control over drug additions in each well. Overall, we perform 6 dosing rounds. We begin with two controls (no Rx^DART^, with mixing) to assess assay stability, followed by three drug rounds (**YM90K**^**DART**^, with mixing), and a final wash (mix 6 times with fresh Tyrode^AMPAR^). In all the entire assay lasts ∼2 hr (20 min per dose × 6 dosing rounds). We settled on three **YM90K**^**DART**^ concentrations per coverslip to balance throughput and cell health. To obtain a 9-point dose-response, we distribute doses 1:9 over coverslips A, B, C (with A = 1, 4, 7; B = 2, 5, 8; C = 3, 6, 9).

Each 20 min dosing round involves ∼4 min of pipetting, 4 min waiting, and 12 min of optogenetic imaging. The optogenetic protocol visits each well once every 2 min for 12 min (i.e., 6 reps), and thus there are 72 stage movements (=12 wells × 6 reps). After each stage movement, we obtain a dTomato image (TRITC channel) followed by an optogenetic pulse train (GFP channel, 16 pulses at 3 Hz, 50ms per pulse). We typically see a rise in the GCaMP signal over the 16-pulse (5 sec) train, with dynamics that are largely reproduced on each of the 6 reps. As described below (data analysis), response variability over reps provides a quality-control metric, and we use the average of the last 4 reps to correspond to a ∼15 min **Rx**^**DART**^ incubation (since pipetting midpoint).

Analysis is performed in custom MATLAB code in four steps. First we perform image alignment using the dTomato channel to account for stage jitter and coverslip drift. Second, cell bodies are segmented using a static image comprised of dTomato and ΔF/F from the first control dosing round. Thus, segmentation is performed without knowledge of the GCaMP6s changes over subsequent doses. We use a combination of automated and manual segmentation, and we proofread each cell to ensure exclusion of pixels wherein cells overlap. Third, we calculate the GCaMP6s waveform for each cell by obtaining the raw trace (mean over segmented pixels of each frame), baseline subtracting (to define 0), and normalizing (to define 1). We account for a small amount of photobleaching by allowing the baseline to vary with time (a linear fit to the first timepoint of each pulse train). We also account for assay rundown so that the three YM90K^DART^ doses on a coverslip can be compared fairly. The correction factors were mild: 0.92, 0.84, and 0.82 (1.0 would reflect no rundown), and were estimated from mock assays with no Rx^DART^, as done previously (Shields et al., 2017). Fourth, we apply inclusion criteria to restrict analysis to cells with GCaMP expression within 1e3 - 1e4 AFU and with < 50% change in activity in the two control doses. Cells with dTomato > 1e4.5 AFU are designated ^+^HTP and used to determine ***HTL***_**50**_ curves **Figure 1C-D**. Cells with dTomato < 1e2.5 AFU are designated ^‒^HTP. A subset of experiments were performed with aavCAG-_dd_HTP_**NLG(432-648)ERXL**_-2A-dTomato-WPRE (_dd_HTP); we saw no statistical difference between _dd_HTP and ^‒^HTP data, and the data were combined to produce the ***Rx***_**50**_ curves **Figure 1C-D**.

With regard to the statistical unit, neurons on a coverslip were highly correlated, such that the majority of the variance was at the level of coverslips rather than cells. This likely reflects the synaptic nature of the assay, wherein cells are highly interconnected. As such, we define a statistical unit as the coverslip, and we weighted the ‘n’ of each cell so that each coverslip could contribute at most n = 1 to the ***Rx***_**50**_ curve, and n=1 to the ***HTL***_**50**_ dataset. The data thus represent mean ±SEM over coverslips. All data were obtained with experimental groups tested side-by-side, such that comparison groups were equally represented in each batch of cells.

### All-optical Cultured-Neuron GABA_A_R Assay

Dissociated neurons were nucleofected with aavCAG-^+^**HTP**_**NLG(432-648)ERXL**_-2A-dTomato-WPRE + aavSYN-**ChR2**-HA-dsfGFP-WPRE + aavCAG-**GCaMP6s**-WPRE. Thus, all three elements (**HTP/ChR/GC**) were in the same cell. Neurons were plated on coverslips and cultured until the assay was performed at 16-18 DIV. We used Tyrode (described above) supplemented with 10 µM CPP (NMDAR antagonist) and 10 µM NBQX (AMPAR antagonist) and 10 µM GABA (low dose GABA_A_R agonist). This combination isolates the GABA_A_R, and further activates endogenous GABA_A_Rs (10 µM GABA) and is thus named Tyrode^GABAR+10^. The underlying principle of the assay is that ChR2 is the only source of excitation for a cell (all excitatory synapses blocked) yet endogenous GABA_A_Rs largely overpower ChR2. We thus take blunted light-triggered activity as an indication of active endogenous GABA_A_Rs, and the reinstatement of light-triggered activity as an indication of GABA_A_R antagonism. The assay is otherwise similar to the AMPAR assay.

With regard to dosing, we performed two control rounds (no Rx^DART^, with mixing) to establish stability, and three drug rounds (**gabazine**^**DART**^, with mixing). However, in lieu of a wash, we used a high dose of regular gabazine (30 µM) as the 6^th^ dosing round, to serve as a calibration for the maximal neural activity (and thus the lowest GABA_A_R function). Imaging is identical to the AMPAR assay, with a dTomato image (TRITC channel) followed by optogenetic stimulation/recording (GFP channel, 6 pulses at 3 Hz, 50ms per pulse), with all 12 stage positions visited 6 times during a single dosing round. Analysis is performed in custom MATLAB code, following the same procedure for the AMPAR assay. The key difference is in regard to the statistical unit, as we found that neurons on a coverslip were not significantly correlated, such that the majority of the variance was at the level of neurons rather than coverslips. This likely reflects the cell-autonomous (non-synaptic) nature of the assay, wherein all excitatory synapses are blocked. As such, we define a statistical unit as the neuron, and plot mean ±SEM over neurons. As before, experimental groups were tested side-by-side, such that comparison groups were equally represented in each batch of cells.

To assay GABA_A_R PAMs, we used a modified Tyrode^GABAR+3^, with a lowered 3 µM GABA level such that ChR2 is not overpowered to start, but becomes overpowered upon positive allostery of the GABA_A_R. Regarding the dosing scheme, we performed only one control round (no Rx^DART^, with mixing), three drug rounds (**diazepam**^**DART**^, with mixing). We next applied regular diazepam (1 µM) to establish a positive control for a full-strength PAM, followed by the 6^th^ and final dose in which we applied gabazine (30 µM) to block all GABA_A_Rs. Imaging alignment, proof-reading and analysis were performed in a similar fashion to the GABAR assay using MATLAB. As with the gabazine assay, a statistical unit is a neuron, and we plot mean ±SEM over neurons. Experimental groups were tested side-by-side, such that comparison groups were equally represented in each batch of cells. In particular, given a lower overall signal to noise in the GABA_A_R PAM assay, we ran side-by-side groups wherein **Rx**^**DART**^ was replaced with a negative control (no drug) or positive control (regular diazepam) to account for assay stability over time (grey data in **Figures S4A**).

### AAV Functional Titer Assay

Dissociated neurons were either plated without modification, for use with non-DIO flexed viruses, or were nucleofected with an aavSyn-iCre construct (AddGene), for use with DIO flexed viruses, and cultured as described above. At DIV 3 or DIV4 along with the first media change, neurons were infected with a serial dilution of virus made in pre-acclimated NbActiv4, then maintained for 2 weeks. Coverslips were rinsed briefly in Tyrode solution containing 1% BSA (Sigma, A7904), incubated with either 1 µM **Alexa647.1**^**DART.2**^ for 15 min at room temperature (diluted in Tyrode +BSA) or with 10 nM **Alexa647.1**^**DART.2**^ for 60 min at 37 °C (diluted in conditioned media+BSA), then rinsed two to three times in Tyrode +BSA. Coverslips were transferred to a glass-bottom imaging plate containing Tyrode +BSA. Each coverslip was imaged in its entirety by combining several 10X images (TRITC at 10ms; Cy5 at 300ms) together using the Olympus CellSens software stitching algorithm. Where possible, a serotype-specific virus with a previously determined functional titer based on the fluorescent dTomato marker was used as a reference for new lots or viruses with the same serotype. If not available, other markers including the ^**+**^**HTP** capture of the **Alx647.1-DART.2** were used to estimate functional titers. The median pixel intensity at a fixed scale was recorded for a given fluorescent channel, and the values for the coverslips used in the serial dilution of a virus were plotted using Excel, against the reference virus, as a function of dose to determine the functional titer.

### Recombinant adeno-associated viral (rAAV) vectors

All viral vectors were produced by the Duke Viral Vector Core. HEK293 cells were transfected with a triple transient transfection protocol of the adenovirus helper plasmid, the AAV helper plasmid and the inverted terminal repeat (ITR) transgene cassette plasmid. The adenovirus helper cassettes encodes the adenovirus proteins (E1A, E1B, E4 and E3A) and the adenovirus virus-associated RNAs required for helper functions. The AAV helper plasmid encodes the wild-type AAV genome lacking ITRs. The rAAV was harvested from the nuclei of transfected cells 48-72 hr after transfection and purified from the cell homogenate using a double round cesium chloride gradient protocol. Genome titers were estimated using real time polymerase chain reaction.

### Acute Brain Slice Preparation

Mice (8-10 weeks) were injected with either AAV_rh10_-CAG-DIO-^**+**^**HTP**_**GPI**_-2A-dTomato-WPRE or AAV_rh10_-CAG-DIO-^**dd**^**HTP**_**GPI**_-2A-dTomato-WPRE to ventral tegmental area (DAT-*Ires*-cre; 6 males, 3 females:titer 2e12, 100nL per site at -3.2mm AP, +-0.5mm ML, -5.0/-4.5mm DV). After 3-5 weeks for expression, mice were deeply anesthetized with isoflurane and euthanized by decapitation. Coronal brain slices (300 µm) containing striatum or VTA were prepared by standard methods using a Vibratome (Leica, VT 1200S), in ice-cold high sucrose cutting solution containing (in mM) 220 sucrose, 3 KCl, 1.25 NaH_2_PO4, 25 NaHCO_3_, 12 MgSO_4_.7H2O, 10 glucose, and 0.2 CaCl2 bubbled with 95% O2 and 5% CO2. The slices were placed into modified artificial cerebrospinal fluid (aCSF) containing (in mM): 120 NaCl, 3.3 KCl, 1.23 NaH_2_PO_4_, 1 MgSO_4_, 2 CaCl_2_, 25 NaHCO_3_, and 10 glucose at pH 7.3, previously saturated with 95% O_2_ and 5% CO_2_. For recovery, the slices were incubated at 33°C for 40-60 min in the bubbled ACSF solution and then allowed to cool to room temperature (22–24°C) until the recordings were initiated.

### Retinal Whole-Mount Preparation

PV-Cre mice (22 males, 38 females, 12-24 weeks) were intravitreally injected with either AAV_7m8_-CAG-DIO-^**+**^**HTP**_**GPI**_-2A-dTomato-WPRE or AAV_7m8_-CAG-DIO-^**dd**^**HTP**_**GPI**_-2A-dTomato-WPRE (4.4E+11 VG/mL, 1.0 µL per eye). After 8-12 weeks, the mice were sacrificed by cervical dislocation. Both eyes were removed and dissected under a stereo-microscope. Retinas were isolated from the eyecups and placed in oxygenated (95% O_2_/5% CO_2_) cold-Ames medium (Sigma-Aldrich), supplemented with 21 mM NaHCO_3_. After the vitreous body was removed, a retina was cut into 3 pieces, leaving each retina piece attached to a filter paper with 2 mm hole by retina ganglion-cell-side-down. The pieces of retina were stored in bubbled Ames solution at room temperature, then transferred to recording chamber with retina ganglion-cell-side-up for recording.

### Acute Brain Slice and Retina Electrophysiology

The brain slices or retinas were perfused with bubbled aCSF at 29-30°C with a 2 ml/min flow rate. Recordings were made by whole cell patch recording techniques using a Multiclamp 700B amplifier (Molecular Devices, Axon Instruments Inc., Union City, CA). The signals were filtered at 10 kHz and acquired using a Digitate 1440A and pClamp 10.7 (Molecular Devices). The recording pipettes (4 MΩ–6 MΩ) were filled with internal solutions. To measure GABA_A_R inhibitory postsynaptic current (IPSC), the pipettes were filled with a cesium chloride-based internal solution (in mM) 135 CsCl, 2 MgCl_2_, 0.5 EGTA, 10 HEPES, 4 Mg-ATP, 0.5 Na-GTP, 10 Na_2_-phosphocreatine, and QX314 (lidocaine N-ethyl bromide), pH adjusted to 7.3 with CsOH (290 mOsm). For AMPAR excitatory postsynaptic current (EPSC), the pipette solution contained (in mM) 130 CsMeS, 1 MgCl_2_, 0.5 EGTA, 10 HEPES, 4 Mg-ATP, 0.5 Na-GTP, 10 Na_2_-phosphocreatine, and 4 QX314, pH adjusted to 7.3 with CsOH (290 mOsm). Evoked IPSC or EPSC signals were elicited by electrical stimuli of 0.3 ms duration and 150-300 μA (60-70% maximum responses), with a repetition interval of 15 sec. The stimulating electrode was placed 60–100 μm from the recorded neuron. GABA_A_R mediated IPSCs were isolated in the presence of DNQX (20 mM, AMPAR antagonist) and AP-V (50 mM, NMDA antagonist) in the bath solution. For AMPAR-mediated EPSCs, picrotoxin (50 mM, GABA_A_R antagonist) and AP-V (50 mM) were added to the bath solution. For current-clamp recording, the pipette solution contained (in mM) 130 K-gluconate, 5 KCl, 2 MgCl_2_, 0.2 EGTA, 10 HEPES, 4 Mg-ATP, 0.5 Na-GTP, and 10 phosphocreatine, pH adjusted to 7.3 with KOH (290 mosM). The liquid junction potential was estimated to be 15.9 mV for the normal aCSF solution and was not corrected. Our inclusion criteria required that cells maintain stable access and holding currents for at least 5 min. In particular, series resistance is monitored using 5–10 mV hyperpolarizing steps interleaved with our stimuli, and cells are discarded if series resistance changed more than ∼15% during the experiment. The stored data signals were processed using Clampfit 10.7 (Axon Instruments). All averaged data are presented as mean ± SEM and *n* represents the number of cells tested per condition. Statistical significance was determined using Student’s *t* or one-way or two-way ANOVA tests. IPSC or EPSC decay time constants were obtained by fitting a single-exponential function, *I*(*t*) = *I* exp^(−t/τ)^ + *I*_*SS*_, where *I*(*t*) is the amplitude of the current at time *t* and *I*_*SS*_ is the steady-state current, *I* is instantaneous current subtracted from *Iss*, and t is the time constant of decay.

### Cerebellum Experiments

BACα6Cre-A x Ai148D-Cre mice (6 males, 4 females, P50–60) were given a 3-mm diameter craniotomy over Crus I at approximately 3.0 mm lateral and 4.3 mm posterior to lambda. Crus I was injected (WPI UMP3) with 150 nL of either AAV_7m8_-CAG-DIO-^**+**^**HTP**_**GPI**_-2A-dTomato-WPRE (1 × 10^12^; Duke Viral Vector Core) or AAV_7M8_-CAG-DIO-^**dd**^**HTP**_**GPI**_-2A-dTomato-WPRE (1 × 10^12^; Duke Viral Vector Core) at a rate of 30 nl/min and a depth of 350 µm at 2–3 sites. Glass windows consisting of two 3-mm coverslips bonded to a 5-mm coverslip (Warner Instruments No. 1) with index matched adhesive (Norland No. 1) were installed in the craniotomy using Metabond. Imaging mice receiving saline and drug infusions received a plastic cannula (Plastics One; C315GS/PK length 0.5 mm) positioned immediately rostral to the imaging window and attached with Metabond. All mice were individually housed after cannula placement and given 8 weeks to allow viral expression, including 1–2 weeks of habituation to head restraint.

Imaging was performed with a resonant scanning microscope (Neurolabware) equipped with a 16× water immersion objective (Nikon CF175 LWD 16xW 0.8NA). The cerebellum was scanned with a TI:Sapphire laser tuned to 920 nm (SpectraPhysics, Mai Tai eHP DeepSee) using a resonant galvanometer (8 kHz, Cambridge Technology) at a frame rate of 30 Hz and a field of view of 278 µm × 117 µm (796 × 264 pixels). Data was collected through a green filter (510 ± 42 nm band filter (SEMrock)) onto GaAsP photomultipliers (H10770B-40, Hamamatsu). Throughout imaging, a polymer (MakingCosmetics, 0.4% Carbomer 940) was used TO stabilize the immersion solution. Images were processed using the open source Python toolbox for large scale calcium imaging data analysis CaImAn and custom written MATLAB code, as described in our companion manuscript (Fleming et al., 2022).

### VTA_DA_ Locomotion Experiments

Adult DAT-IRES-cre mice (16 males, 14 females; 10-20 weeks old) were anesthetized and stereotaxically injected with 400 nL of either AAV_rh10_-CAG-DIO-^**+**^**HTP**_**GPI**_-2A-dTomato-WPRE or AAV_rh10_-CAG-DIO-^**dd**^**HTP**_**GPI**_-2A-dTomato-WPRE (titer 2e12, 100nL per site, two tracks with two depths per track: -3.2mm AP, +-0.5mm ML, -5.0/-4.5mm DV) and immediately implanted with a bilateral metal cannula above the VTA (P1Tech; C235G-1.0) lowered to -3.75mm. Mice were fitted with a plastic head bar adhered to the skull with UV-glue and dental cement, enabling head fixation to facilitate drug infusions in awake animals. Assays were performed at least 3 weeks after viral injection to allow for recombinant protein expression. Mice were singly housed post-surgery, in a 12-hr reverse light cycle, with food and water provided ad libitum. Mice were acclimated to head-fixation for 3 consecutive days prior to behavioral experiments.

Mice were head-fixed on a round plastic treadmill (Delvie’s Plastics, 8” plexiglass disk covered with silicone rubber) attached to a rotary encoder to collect rotation data (U.S. Digital H5-100-NE-S). Rotary encoder data was collected by a National Instruments card (NI USB-6351 X Series DAQ) and a custom MATLAB script. Mice were water restricted to 80-85% body weight and delivered occasional sucrose rewards (Lee Company solenoid LHDA1233315H HDI-PTD-Saline-12V-30PSI) to induce motivation to run. Behavior sessions consisted of 1 hr per day on consecutive days. After 10 baseline running sessions to fully acclimate to the treadmill, mice had reagents infused into the brain via the cranial cannula as previously described (Shields et al., 2017). DART reagents were freshly thawed and diluted in sterile ACSF to 10 µM **gabazine.7**^**DART.2**^ + 1 µM **Alexa647.1**^**DART.2**^, with 0.6nL per hemisphere infused at a rate of 0.1uL/min. Mice were then recorded on the treadmill at 2 hr and 24 hr post-infusion. Wheel speed was analyzed by a custom MATLAB script, and was normalized per mouse to their average wheel speed during the final training session.

### Open-Field Turning Assay

Adult (10-20 week old) Drd1-Cre mice (6 males, 8 females) were anesthetized and stereotaxically injected with 960 nL of AAV (titer 2e12, 80 nL per site, four tracks with three depths per track: 0.55mm AP, -1.7mm ML, -2.8/-2.3/-1.8mm DV; 1.35mm AP, -1.3mm MP, -3.0/-2.5/-2.0mm DV; 1.45mm AP, -2.1mm ML, -3.1/-2.6/-2.1mm DV; 0.65mm AP, -2.5mm ML, -3.1/-2.6/2.1mm DV); we found that distributed injections of lower volumes of virus resulted in consistent and robust expression without toxicity. Mice were immediately implanted with a plastic (peek) cannula (P1Tech, C315G/PK) at -1.9mm ML and 1.0mm AP lowered to -2.5mm; peek was favored for these experiments to eliminate the hypothetical possibility that nitroaromatic compounds like YM90K could be reduced by iron-containing surfaces. Mice were fitted with a plastic head bar adhered to the skull with UV-glue and dental cement, enabling head fixation to facilitate drug infusions in awake animals.

Reagents were infused into the brain via the cranial cannula as previously described (Shields et al., 2017). Following infusion, mice were placed into open-field chambers (27cm x 27cm) in the dark, and behavior was recorded using infrared video. The positions of the nose, tail and center of mass of each mouse were tracked using Noldus Ethovision XT 10, and analyzed offline in custom MATLAB scripts. Changes in orientation of the vector pointing from tail to nose were analyzed to identify vector rotations of at least 360° with no more than 25% cumulative rotation in the opposite direction (e.g., a 400° rotation to the right could contain no more than 100° cumulative hesitations to the left). Rotations were normalized (e.g., 400° / 360° = 1.11 turns). False rotations attributed to nose/tail assignment flips were eliminated by excluding frames in which the nose-tail distance decreased to less than half of the median value. Left and right rotations were separately tallied, and net turns (e.g., Figure 3A) is total left minus right turns per hr. For analysis of rotations binned according to turn diameter (e.g., Figure 3B), a sliding-window analysis was performed over each 360° portion of a turn; diameter was defined as the maximum Euclidean distance between all center of mass positions. Separate histograms were tallied for left and right turns.

### Dorsal Striatum Histology

Mice were deeply anesthetized with isoflurane and fixed by transcardial perfusion of 15mL PBS followed by 50mL ice-cold 4% paraformaldehyde (PFA) in 0.1M PB, pH 7.4. Brains were excised from the skull, further post-fixed in 50mL of 4% PFA at 4 C overnight then washed three times with PBS. Brains were embedded in 5% agarose and sliced along the coronal axis at 50 µM (Leica, VT1200S). Sections were mounted onto glass slides (VWR 48311-703) and coverslipped with Vectashield mounting medium (Vector Labs, H-1400 or H-1800). Fluorescent images (DAPI, GFP, TRITC, Cy5) were collected at 10X magnification with either an Olympus IX83 inverted microscope or an Olympus VS200 slide scanner. Images were analyzed for pixel intensity using MATLAB. For each coronal section, the striatum was manually segmented in both hemispheres. Background fluorescence (median of right hemisphere) was subtracted. Dye capture in the left hemisphere was calculated via a pixel-wise summation over 25 coronal sections (cannula center ±12 sections; 50 µm per section).

### Visual Cortex Surgeries

SOM::Cre mice (JAX#013044, 2 males and 1 female, for two-photon experiments) of a 50% CBA/CaJ background, or CBA mice (JAX # :000654, 2 males, for ambient pharmacokinetic experiments) aged > P45 were administered Dexamethasone (3.2 mg/kg, s.c.) <2 hr before surgery. Mice were anesthetized with isoflurane (1.25–2% in 100% O2), ketamine (200 mg/kg, i.p.), xylazine (30 mg/kg, i.p.), then an incision was made in the scalp and the dorsal side of the skull was scraped clean of remaining tissue. A guide cannula (F11552, P1 Technologies) with a complementary dummy cannula (F11372, P1 Technologies) was directed to the right lateral ventricle using the following coordinates from bregma: 1.10mm lateral, 0.20mm posterior, 2.30 from the skull surface. The cannula was secured to the skull with Metabond (Parkell).

Within the same surgery, a custom titanium headpost was cemented to the skull with Metabond and a 4.5 mm craniotomy was made over the left visual cortex centered on the following coordinates from lambda: 3.10 mm lateral, 1.64 mm anterior. The craniotomy was fit with a custom-made glass window composed of a 6 mm coverslip bonded to two 4.5 mm coverslips (Warner, no. 1) with refractive index-matched adhesive (Norland, no. 71), and the window was fixed in place with Metabond. Buprenorphine (0.05 mg/kg) and cefazolin (50 mg/kg) were delivered s.c. for 48 hr following surgery. Following at least 7d recovery from the implantation surgery, mice were gradually habituated to head restraint. Mice to be used for two-photon experiments then underwent retinotopic mapping using widefield autofluorescence imaging (Jin and Glickfeld, 2020) to locate the primary visual cortex within the cranial window, which served as the target for viral injections.

The mice used for two-photon imaging underwent an additional surgery for viral injection. Dexamethasone (3.2 mg/kg, s.c.) was administered at least 2 hr before surgery. After anesthesia with isoflurane (1.25–2% in 100% O2), the cranial window was removed. AAV_rh10_-CAG-DIO-^**+**^**HTP**_**GPI**_-IRES-dTomatoF-W3SL (titer 9 × 10^13^ GC/mL) mixed with AAV_9_-SYN-jGCaMP8s-WPRE (titer 2.8 × 10^13^ GC/mL) in a 1:1 ratio was injected via a glass micropipette mounted on a Hamilton syringe, driving expression of HTP and dTomato in somatostatin+ interneurons while driving GCaMP expression in all neurons. Two hundred nanoliters of virus were injected at 170-230 µM below the pia (30 nl/min); the pipette was left in the brain for an additional 3 min to allow the virus to infuse into the tissue. Following injection, a new coverslip was sealed in place with Metabond. We then waiting a minimum of two weeks for viral expression to mature before performing two-photon experiments.

### Lateral Ventricle Ambient Pharmacokinetics

For YM90K.1^DART.2^ and Alexa647.1^DART.2^ delivery, mice were headfixed on a running wheel and the dummy cannula removed. An internal cannula (F11373, P1 Technologies) connected to a Hamilton syringe on an infusion pump was inserted into the guide cannula and secured in place. Compounds were delivered at 100nL/min, followed by at 10-20 min waiting period before the internal cannula was removed. The dummy cannula was then reinserted and secured.

We visualized Alexa647.1^DART.2^ through the cranial window using widefield microscopy. An excitation wavelength of 624±40 nm was delivered through the cranial window and emitted light was filtered at 692±40m. Images were collected using a CCD camera (Rolera EMC-2, QImaging) through a 5 air-immersion objective [0.14 numerical aperture (NA), Mitutoyo], using Micromanager acquisition software (NIH), and were analyzed in imageJ (NIH) to qualitatively (for two-photon experiments) or quantitatively (for ambient pharmacokinetics experiments) measure changed in fluorescence.

### Visual Cortex 2P Imaging and Analysis

Images were collected using a two-photon microscope controlled by Scanbox software (Neurolabware). A Mai Tai eHP DeepSee laser (Newport) was directed into a modulator (Conoptics) and raster scanned on the visual cortex using resonant galvanometers (8 kHz; Cambridge Technology) through a 16× (0.8 NA, Nikon) water-immersion lens. Emitted photons were directed through a green filter (510 ± 42 nm band filter; Semrock) or a red filter (607 ± 70 nm band filter; Semrock) onto GaAsP photomultipliers (H10770B-40, Hamamatsu). Frames were collected at 15Hz. At the start of each experiment, an excitation wavelength of 1040 nm was used to visualize dTomato fluorescence, allowing identification of ^+^HTP cells. All subsequent imaging employed an excitation wavelength of 920 nm. Mice were headfixed on running wheel, and locomotion was monitored with a digital encoder (US Digital, H5-32-NE-S) at 10 Hz. Full-field drifting gratings in eight directions (from 0 to 315 degrees in intervals of 45 degrees) at three contrasts (25%, 50%, and 100%; data shown is for 50% contrast) were presented to the right visual field; each trial constituted a 2s stimulus presentation and a 4s interstimulus interval. Data were collected at 180 – 250 µM below the cortical surface.

Custom code written in MATLAB (MathWorks) was used to analyze two-photon data. To adjust for x-y motion, we registered all frames from each imaging session to a stable reference image selected out of several 500-frame-average images, using Fourier domain subpixel 2D rigid body registration. We manually segmented cell bodies, first using the dTomato fluorescence to segment and identify ^+^HTP cells, then segmenting all other visible cells from images of the average dF/F signal during presentation of each stimulus, a time-averaged image of the full stack, and a local correlation map. These later two segmentation strategies allowed detection of cells that were active only weakly visually responsive. All segmented cells that were not identified based on dTomato fluorescence were labelled as -HTP and assumed to be putative pyramidal cells.

We derived fluorescence time courses by averaging all pixels in a cell mask. To exclude signal from the neuropil, we first selected shell around each neuron (excluding neighboring neurons), then estimated the neuropil scaling factor by maximizing the skew of the resulting subtraction, and finally removed the identified component from each cell’s time course. Time courses were then split into 6s epochs corresponding to stimulus trials, and visually-evoked responses were measured as the average dF/F in the 2s stimulus period (where F was designated at the mean fluorescence in the 1s period preceding the stimulus).

For each mouse we performed a baseline imaging session the day before YM90K^DART^ ICV delivery, then re-identified the same imaging field of view in a second imaging session after delivery. We matched cells across imaging sessions using a custom MATLAB script; briefly, after registration the image stack from the baseline session was rotated to fully align with the image stack from the session following drug delivery, then for each cell found in the post-ICV session we examined a small field of view in the corresponding region of the stack from the baseline session to determine whether the cell was visible in the baseline session. Among cells that we could identify in both imaging session, we included for analysis any cell that was visually responsive (demonstrated a statistically significant elevation in dF/F during the stimulus period for at least one stimulus condition as defined by a Bonferroni corrected paired t-test) in at least one of the sessions. We then found the preferred direction of visual grating for each cell on each day, and analyses were performed on the subset of trials at that grating direction for each cell. We further restricted the present analyses to stationary periods (wheel speed < 2cm/s).

### QUANTIFICATION AND STATISTICAL ANALYSIS

Reported values are mean ± SEM. Statistical tests were performed via Student’s t-test (paired or unpaired), or nonlinear regression (MATLAB curve fitting toolbox). The statistical test, sample size and *p* value are indicated in each figure legend. No statistical methods were used to predetermine sample size. All experiments involving a quantitative comparison of an experimental parameter were performed in a manner to counterbalance uncontrolled sources of variability. In cultured neurons, data obtained to compare two or more reagents (e.g., YM90K.1^DART.1^ vs YM90K.1^DART.2^) was obtained with experimental groups tested side-by-side, such that all comparison groups were equally represented in each batch of cells. For slice, retina, and mouse behavior, positive vs negative groups (e.g., ^+^HTP vs _dd_HTP mice) were interleaved and run side-by-side, using cage-mates when possible, and balancing groups to minimize confounding differences (e.g., animal sex, experimenter, time of year), with the goal of isolating the experimental variable of interest.

## DATA AND SOFTWARE AVAILABILITY

All data and software are available upon request to the corresponding author.

## SUPPLEMENTARY FIGURES

**Figure S1.**
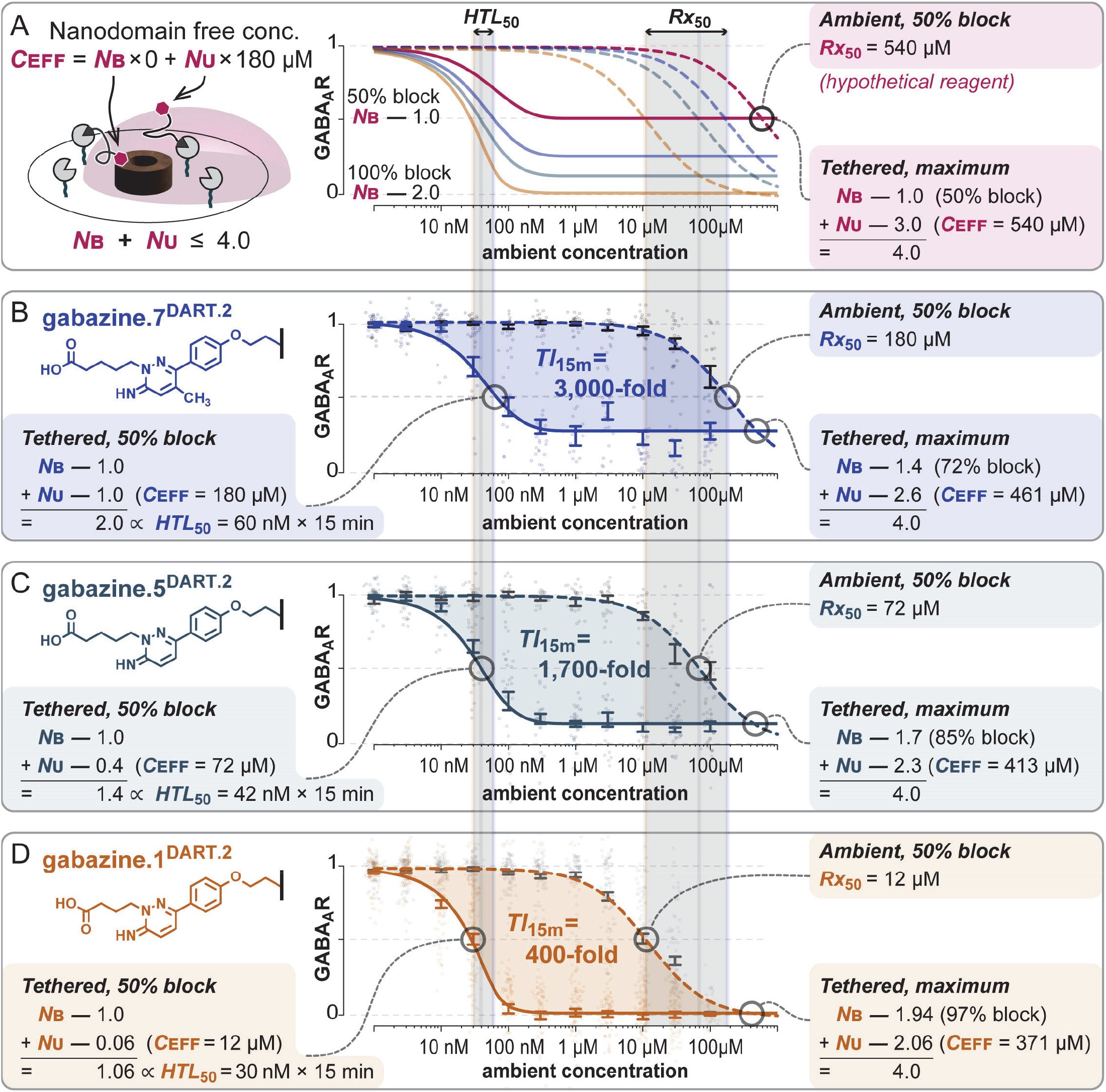
Supporting Materials for Figure 1. **(A) gabazine**^**DART**^ **model**. In a small nanodomain, the number of **Rx** molecules is few. Thus, the number that are GABA_A_R-bound (***N***_**B**_) becomes significant, leading to ligand depletion, as only the GABA_A_R-unbound (***N***_**U**_) subset contributes to the mobile nanodomain concentration (***C***_**EFF**_). The relevant equations and parameters are described in the supplementary text. Three points merit emphasis. First, we estimate a volume so small that each unbound **Rx** contributes an effective 180 µM concentration. Second, the channel is half-blocked when ***N***_**B**_ = 1 and fully blocked when ***N***_**B**_ = 2 (there are two binding sites per channel). Third, we estimate a total of ∼4 HTP proteins per nanodomain, which sets an upper limit on the nanodomain concentration. We illustrate the model with a hypothetical low-affinity reagent, with ***Rx***_**50**_ = 540 µM. Ligand depletion would be subtle in this case, with only 1 of 4 tethered drugs depleted at the tethered maximum. Given 3 remaining (GABA_A_R-unbound) drugs, ***C***_**EFF**_ = 3 × 180 µM = 540 µM, which matches the **Rx**_**50**_ for this hypothetical drug. As such, the GABA_A_R would be only half-blocked at the tethered maximum. **(B-D) gabazine**^**DART**^ **dataset**. Chemical structure of each **Rx** is shown on the top left of each panel. Raw data, with means ±SEM shown in the center. Model fits shown in dashed (**Equation 1**) and solid (**Equation 2**) curves, with numerical values as indicated. Starting on the right hand side, we note that **gabazine.7**^**DART.2**^ achieves 72% tethered maximum block, and the most potent **gabazine.1**^**DART.2**^ achieves 97% tethered block. In parallel to the increase in max block, we also observe accentuated ligand depletion (reduction in maximum ***C***_**EFF**_) for **gabazine.1**^**DART.2**^. If we now examine the left-hand side of the fits (near ***HTL***_**50**_), we again observe moderate ligand depletion for **gabazine.7**^**DART.2**^ wherein half of the tethered drug is depleted near its ***HTL***_**50**_ (***N***_**B**_ = 1.0 and ***N***_**U**_ = 1.0). As we move to **gabazine.1**^**DART.2**^, ligand depletion becomes severe, with nearly all drug depleted near its ***HTL***_**50**_ (***N***_**B**_ = 1.0 and ***N***_**U**_ = 0.06). Thus, the impact of ligand depletion is unequal; ***HTL***_**50**_ is impacted most for **gabazine.1**^**DART.2**^, narrowing the gap to **gabazine.7**^**DART.2**^.

### Derivation of gabazine^DART^ model

Experimental **gabazine**^**DART**^ data in **Figure S1** (summarized to the right), contains a surprise in that large changes in ***Rx***_***50***_ correspond to smallnchanges in ***HTL***_***50***_.

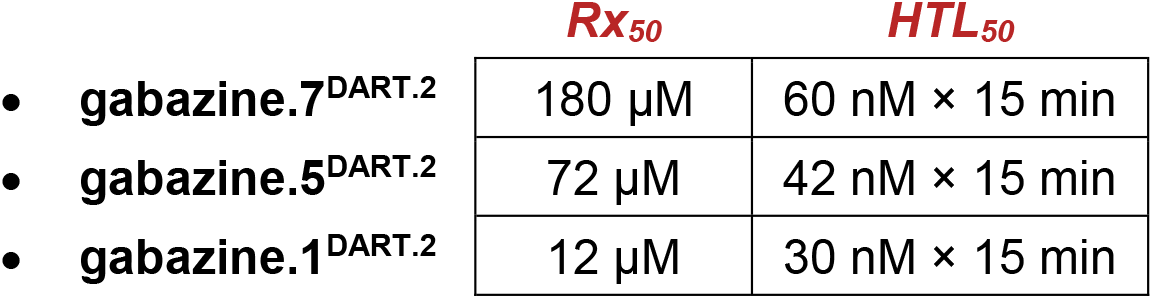

The paradox can be stated explicitly in **Equations 1-2**, which indicate that both *ambient* and *tethered* configurations are subject to similar Langmuir binding rules, with the key difference being that the tethered concentration (***C***_**EFF**_) would be a boosted version of the ambient concentration (***C***_**ambient**_). If this boost were linear (***C***_**EFF**_ = β × ***C***_**ambient**_) then we would expect ***Rx***_**50**_ and ***HTL***_**50**_ to also be linearly proportional to one another (***Rx***_**50**_ = β × ***HTL***_**50**_). Moreover, if we consider nonlinearities wherein ***C***_**EFF**_ saturates with increasing ***C***_**ambient**_, we find that this could only broaden the spread of ***HTL***_***50***_, exacerbating the paradox.

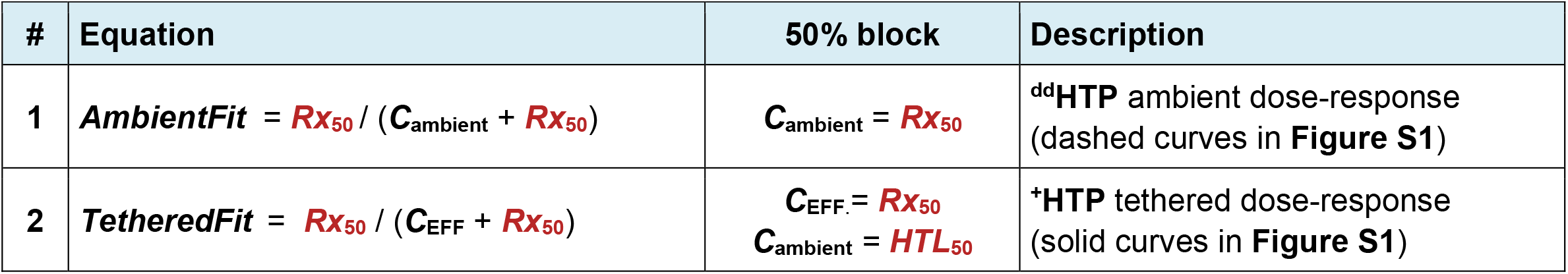

We next considered pharmaceutical ligand depletion, which is known to arise in miniaturized volumes (Carter *et al*., 2007). We started with a physical ∼16 nm length scale over which a GABA_A_R can interact with a fully extended tethered drug (**Figure S1A**, left cartoon). Within these dimensions, there would be few **Rx** molecules in a nanodomain, each contributing a substantial concentration. As such, one must account for each individual molecule. In particular, only the GABA_A_R-unbound (***N***_**U**_) subset of **Rx** molecules contribute to ***C***_**EFF**_, as only these can collide to drive new binding events. Conversely, only the GABA_A_R-bound (**N**_**B**_) subset can modulate the channel. However, binding comes with a price, as even one GABA_A_R-bound drug (***N***_**B**_ = 1) would substantially lower the mobile concentration (***C***_**EFF**_) in a nanodomain.

Equilibrium of this model obeys **Equations 3-5**, which describe how a given total number of nanodomain drugs (***N***_**T**_) is equilibrated into GABA_A_R-bound (***N***_**B**_) and unbound (***N***_**U**_) subsets. **Equation 6** describes the capture kinetics, whereby ***C***_**ambient**_ determines ***N***_**T**_. Thus, the only parameter allowed to vary for each **gabazine**^**DART**^ variant is ***Rx***_**50**_ (in red), while the physical parameters (in blue) are common to all.

| # | Equation | Description |
| --- | --- | --- |
| 3 | **C<sub>EFF</sub>** = **N<sub>B</sub>** × 0 + **N<sub>U</sub>** × **C<sub>x</sub>** | Effective mobile concentration in the nanodomain |
| 4 | **N<sub>U</sub>** = **N<sub>T</sub>** – **N<sub>B</sub>** | Number of nanodomain **Rx** unbound to GABA<sub>A</sub>R |
| 5 | **N<sub>B</sub>** = **N<sub>B,max</sub>** × **C<sub>EFF</sub>** / (**C<sub>EFF</sub>** + **Rx<sub>50</sub>**) | Number of nanodomain **Rx** bound to GABA<sub>A</sub>R. |
| 6 | **N<sub>T</sub>** = **N<sub>T,max</sub>** × (1 – e<sup>–C<sub>ambient</sub> × 15 min × **k<sub>T</sub>**</sup>) | Number of nanodomain **Rx** total |

The data for all three reagents could be jointly fit with the common parameter set shown below. There is little wiggle room in these parameters, and thus the values are tightly constrained by data. Additional insight is provided in the legend of **Figure S1**, as well as in the **Discussion** of the main text.

| Name | Value | Description |
| --- | --- | --- |
| **C<sub>x</sub>** | 180 μM | **Conc. per Rx**. Related to nanodomain radius: **r<sub>nano</sub>** = 92.6 [nm × μM<sup>1/3</sup>] / **C<sub>x</sub>**<sup>1/3</sup> |
| **N<sub>B,max</sub>** | 2 | Maximum # of **Rx** bound to GABA<sub>A</sub>R (from GABA<sub>A</sub>R structure/function). |
| **N<sub>T,max</sub>** | 4 | Maximum # of **Rx** total in the nanodomain (= number of HTP in nanodomain). |
| **k<sub>T</sub>** | 6.6 × 10<sup>5</sup> M<sup>–1</sup>min<sup>–1</sup> | Tethering rate constant (rate of **gabazine<sup>DART</sup>** capture by HTP) |

**Figure S2.**
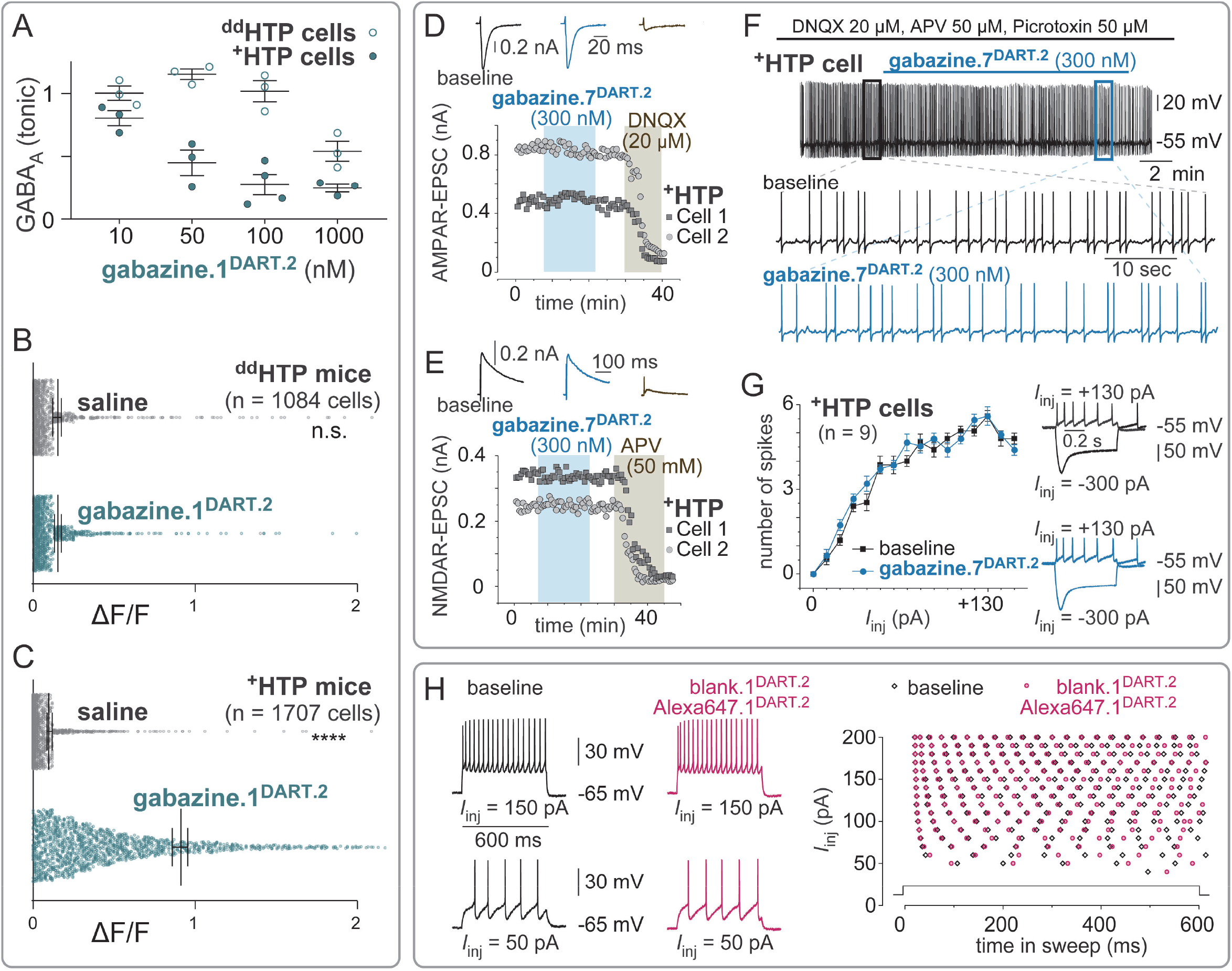
Supporting Data for Figure 2 **Cell-specific manipulations of** ^**+**^**HTP cerebellar neurons with gabazine.1**^**DART.2**^. **(A)** Voltage clamp of ^**+**^**HTP** vs ^**dd**^**HTP** granule cells, measuring tonic GABA_A_ receptor currents. The 100 nM dose, along with details of the experiment, are shown in **Figure 2A**. Here, we show summary data for the **gabazine.1**^**DART.2**^ dose-response. On-target effects on ^**+**^**HTP** cells yielded GABA_A_R antagonism at ***HTL***_**50**_ ∼ 20 nM × 15 min. Off-target effects on ^**dd**^**HTP** cells required ***Rx***_**50**_ ∼ 1,000 nM. **(B)** Supporting data for **Figure 2D**. Granule cells in _**dd**_**HTP** mice showed no significant change in GCaMP signals after **gabazine.1**^**DART.2**^. n = 1084 cells from 4 mice, p = 0.76, paired t-test, before vs after infusion. **(C)** Supporting data for **Figure 2C**. Granule cells in ^**+**^**HTP** mice exhibited a significant increase in GCaMP signals after **gabazine.1**^**DART.2**^ n = 1707 cells from 6 mice, p < 0.0001, paired t-test, before vs after infusion. **Receptor Specificity Assays:** **(D)** AMPA receptor mediated evoked excitatory postsynaptic potentials (evoked EPSC) of ^+^HTP VTA neurons (black trace) were not affected by tethered gabazine.7^DART.2^ (blue trace) but were blocked by the AMPA antagonist DNQX (tan trace). **(E)** NMDA receptor mediated EPSC of ^+^HTP VTA neurons (black trace) were not affected by tethered gabazine.7^DART.2^ (blue trace) but were blocked by the NMDA antagonist AP5 (tan trace). **(F)** Current clamp experiments shows that gabazine.7^DART.2^ does not affect action potential firing patterns when GABA_A_ receptors were blocked by picrotoxin, AMPA receptors blocked by DNQX, and NMDA receptors blocked by AP5, suggesting that all other receptors are not affected by gabazine.7^DART.2^. **(G)** Quantification of the number of action potentials fired over a 500 ms period, as a function of injected current. No change was observed before vs after gabazine.7^DART.2^ was tethered. Representative traces shown on the right. **Validation that 300 nM blank.1**^**DART.2**^ **+ 30 nM Alexa647.1**^**DART.2**^ **is pharmacologically inert**. **(H)** Spike timing of ^**+**^**HTP** VTA neuron in response of different magnitudes of injected current. Left: representative traces. Right: spike raster for all injected currents.

**Figure S3.**
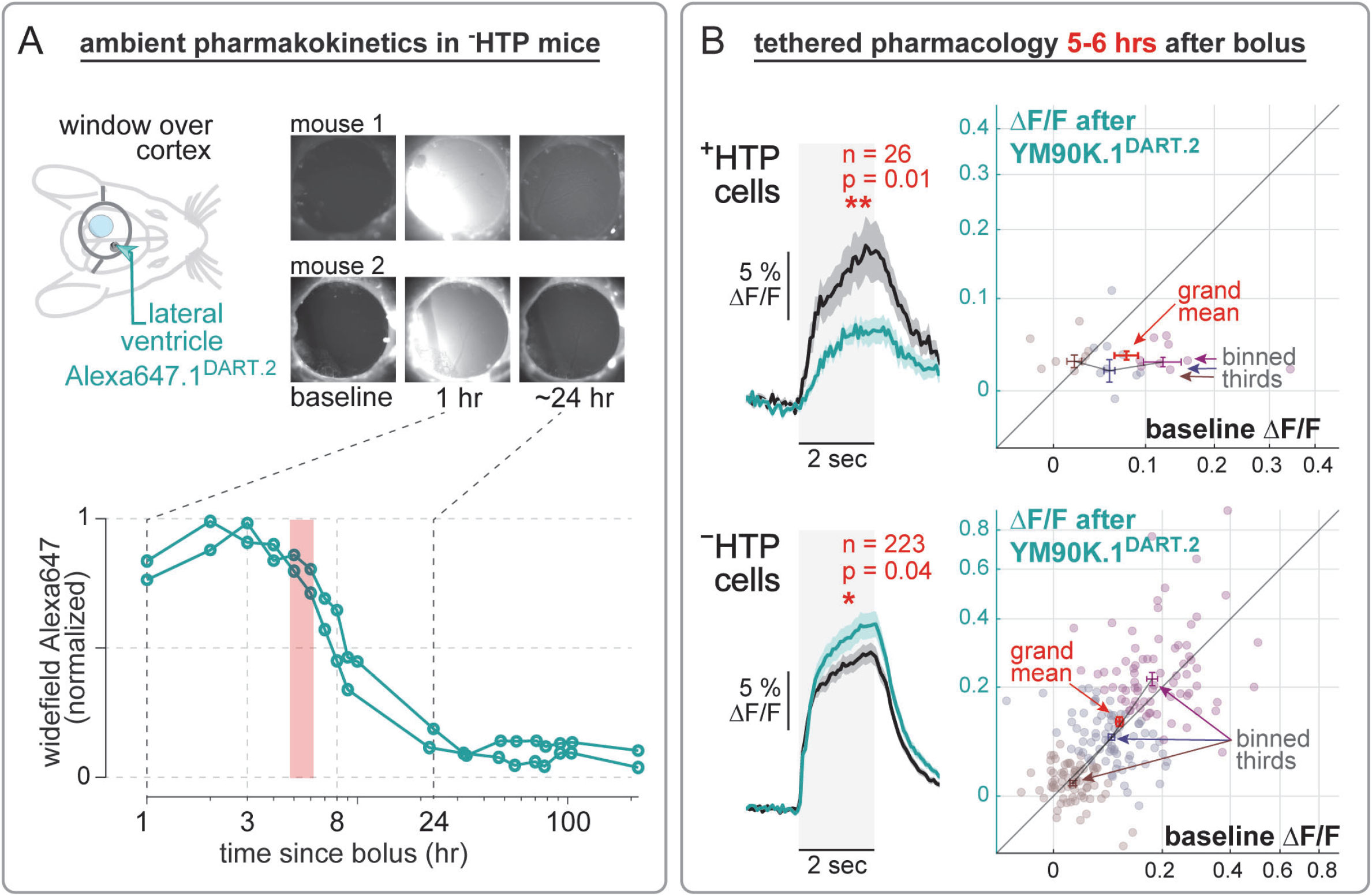
Supporting Data for Figure 3 **(A)** Two wild-type mice were infused with **Alexa647.1**^**DART.2**^ (2 μL over 20 min). We performed widefield imaging of the far-red channel starting one hr after infusion, every hr for the initial 10 hr, then approximately twice a day for the following several days. Upper panel: example widefield images from each mouse. Lower panel: Fluorescence values were determined by taking the mean pixel intensity of an ROI at the center of the field of view. Each value is normalized to the maximum and minimum of the time course. We note that the widefield signal was remarkably constant over the first ∼6 hr, consistent with dilution of the bolus over time. We thus selected the 5-6 hr timepoint for the next experiments (red shading). **(B)** Supporting data for **YM90K.1**^**DART.2**^ experiments in SOM::Cre mice. Left: traces (from **Figure 3G**) represent the mean ±SEM over all cells from three mice during a 2 sec stimulus period. We use data corresponding to the preferred direction of each cell on trials when mice are stationary. Black is before, cyan is 5-6 hr after bolus delivery of **YM90K.1**^**DART.2**^ into the contralateral ventricle. Right: cell-by-cell scatterplot (each symbol is one cell). We plot ∆F/F before vs after the **YM90K.1**^**DART.2**^ manipulation. Red symbols show the grand mean ±SEM. We additionally bin the data into thirds, along an angle defined by the data variance in horizontal vs vertical axes. The binned mean ±SEM suggest that effects of **YM90K.1**^**DART.2**^ are most pronounced for cells with initially high activity. In particular, for ^**+**^**HTP** SOM cells (top) the cells with the highest baseline activity are the most dramatically attenuated by their accumulation of **YM90K.1**^**DART.2**^. Likewise, for ^**–**^**HTP** putative pyramidal cells (bottom) the cells with the highest baseline activity are the most dramatically disinhibited, presumably due to attenuation of neighboring SOM cells by **YM90K.1**^**DART.2**^.

**Figure S4.**
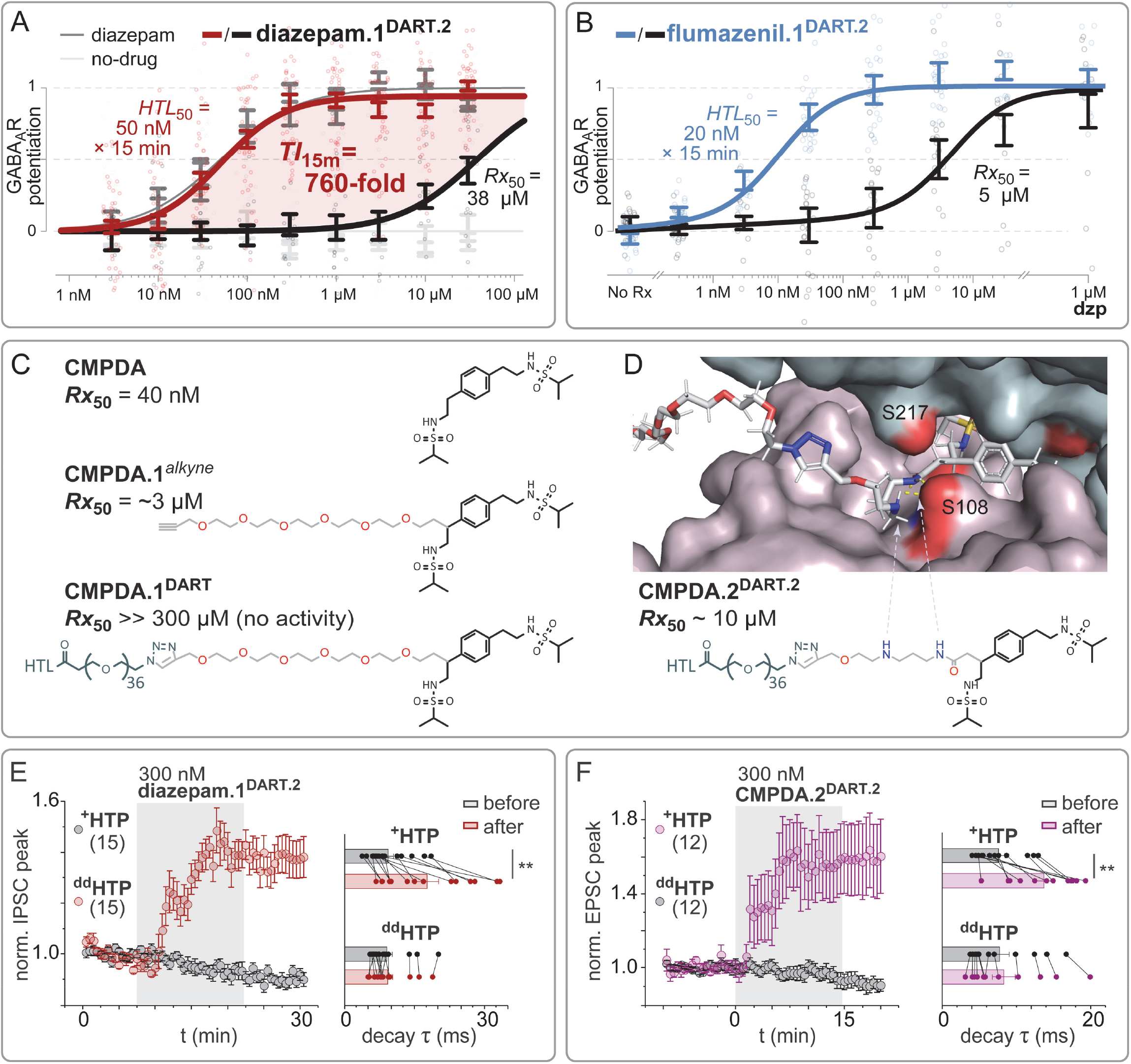
Supporting Data for Figure 4 **(A)** GABA_A_R assay adapted for PAMs. ChR2 and GCaMP6s are co-expressed in the same neuron. Excitatory synapses are blocked, leaving ChR2 as the only trigger of neural activity. We begin the assay by gently activating endogenous GABA_A_Rs (3 µM GABA present throughout), which only partially attenuates ChR2. We then add a putative GABA_A_R PAM at a given dose, which if successful will potentiate the impact of GABA to further overpower the ChR2-triggered GCaMP (**methods**). We performed side-by-side assays with no drug (light gray) and traditional diazepam (dark grey), wherein data from ^+^**HTP** and _**dd**_**HTP** neurons were indistinguishable. We thus normalized all data to 0 (no drug) and 1 (diazepam saturation). Given this calibration, **diazepam.1**^DART.2^ functioned as a full-strength PAM on par with diazepam. ***HTL***_**50**_ = 50 nM × 15 min (^+^**HTP** neurons) and ***Rx***_**50**_ = 38,000 nM (^**dd**^**HTP** neurons), yielding ***TI***_**15m**_ = 760-fold. **(B)** Full-strength positive allostery produced by **flumazenil.1**^**DART.2**^. Format as in panel A. Final dose is 1 µM diazepam. **(C)** Structure and ***Rx***_**50**_ of **CMPDA, CMPDA.1**^***alkyne***^, and **CMPDA.1**^**DART**^ (**methods**). Note electronegative (red atoms) in spacer. **(D)** Top: computational design of spacer compatible with electrostatics of the AMPAR (PDB 3RNN). Electropositive (blue atoms) of spacer predicted to hydrogen bond to electronegative S108 of the receptor. Bottom: Structure and ***Rx***_**50**_ of **CMPDA.2**^**DART.2**^ (**methods**). **(E)** Supporting data for **Figure 4F**. Retinal ganglion cells in voltage-clamp, GABA_A_R-mediated evoked IPSC configuration (**methods**). Left: normalized IPSC peak is augmented ∼1.4-fold by **diazepam.1**^**DART.2**^. Data are mean ±SEM of cells, normalized to their baseline. Right: IPSC time constant is prolonged ∼1.9-fold by **diazepam.1**^**DART.2**^. Values are exponential fits of the IPSC decay of each cell. **(F)** Supporting data for **Figure 4C**. Retinal ganglion cells in voltage-clamp, AMPAR-mediated evoked EPSC configuration (**methods**). Left: normalized EPSC peak is augmented ∼1.6-fold by **CMPDA.2**^**DART.2**^. Data are mean ±SEM of cells, normalized to their baseline. Right: EPSC time constant is prolonged ∼1.8-fold by **CMPDA.2**^**DART.2**^. Values are exponential fits of the EPSC decay of each cell.

**Supplementary Table S1.**
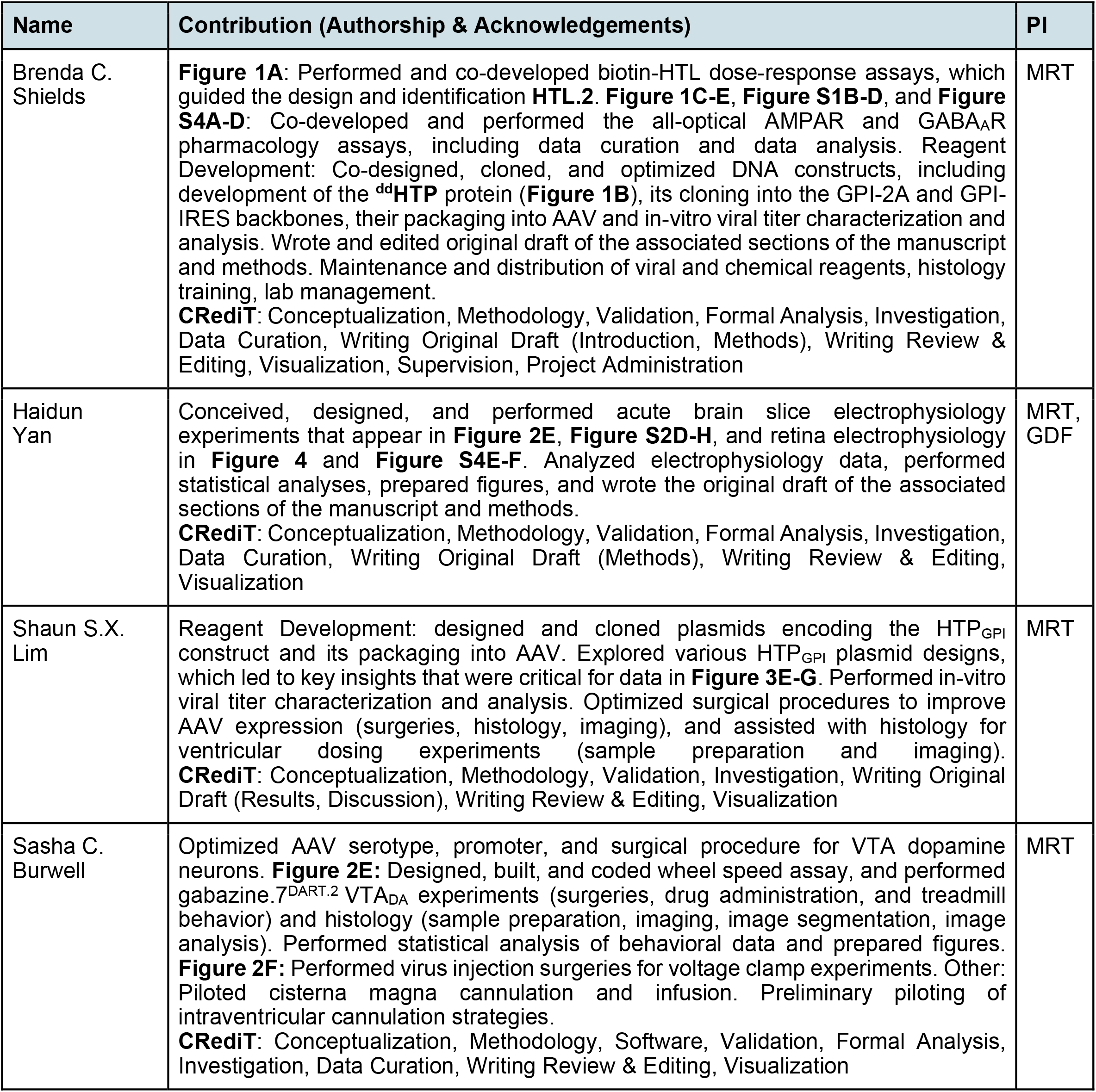
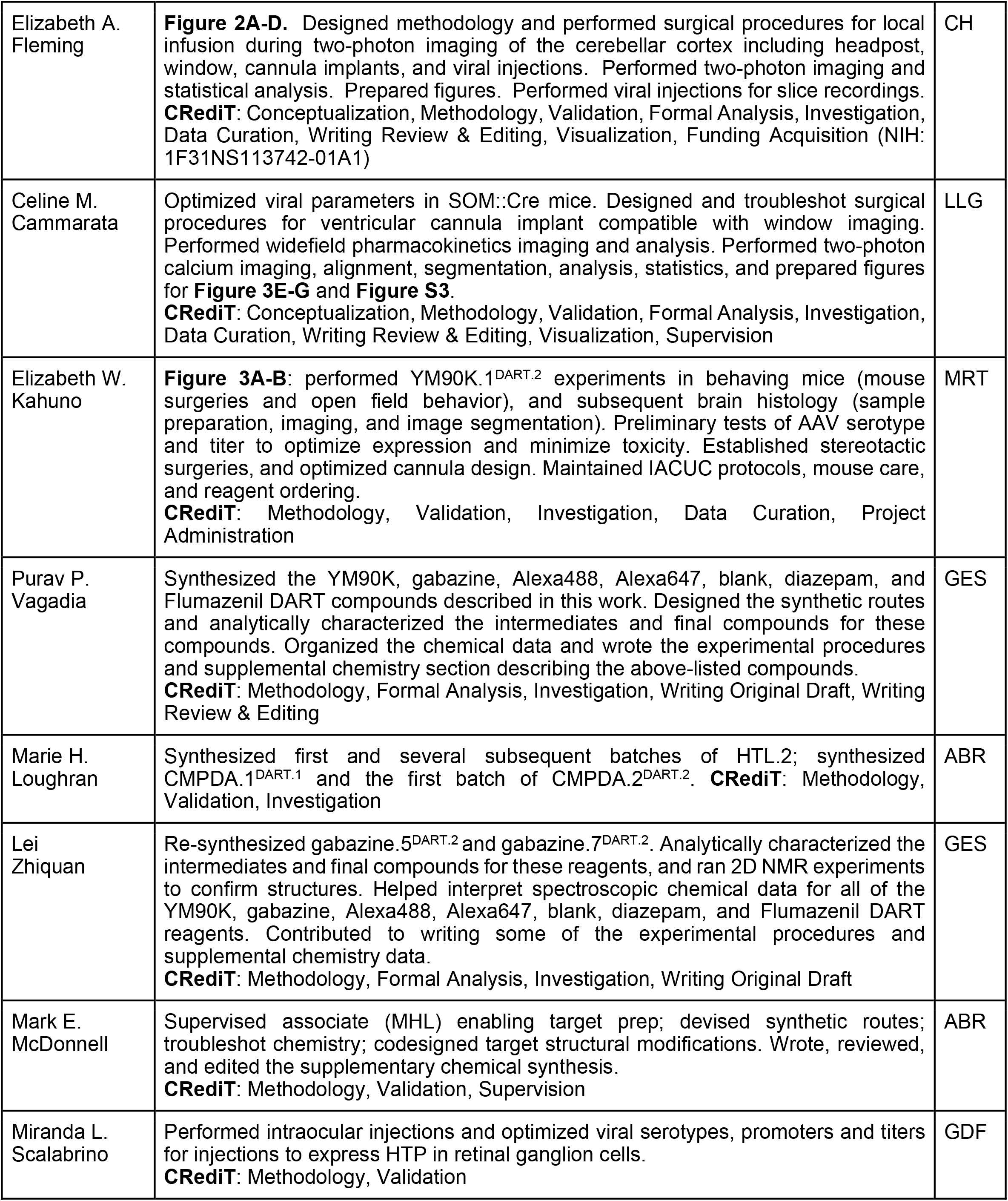
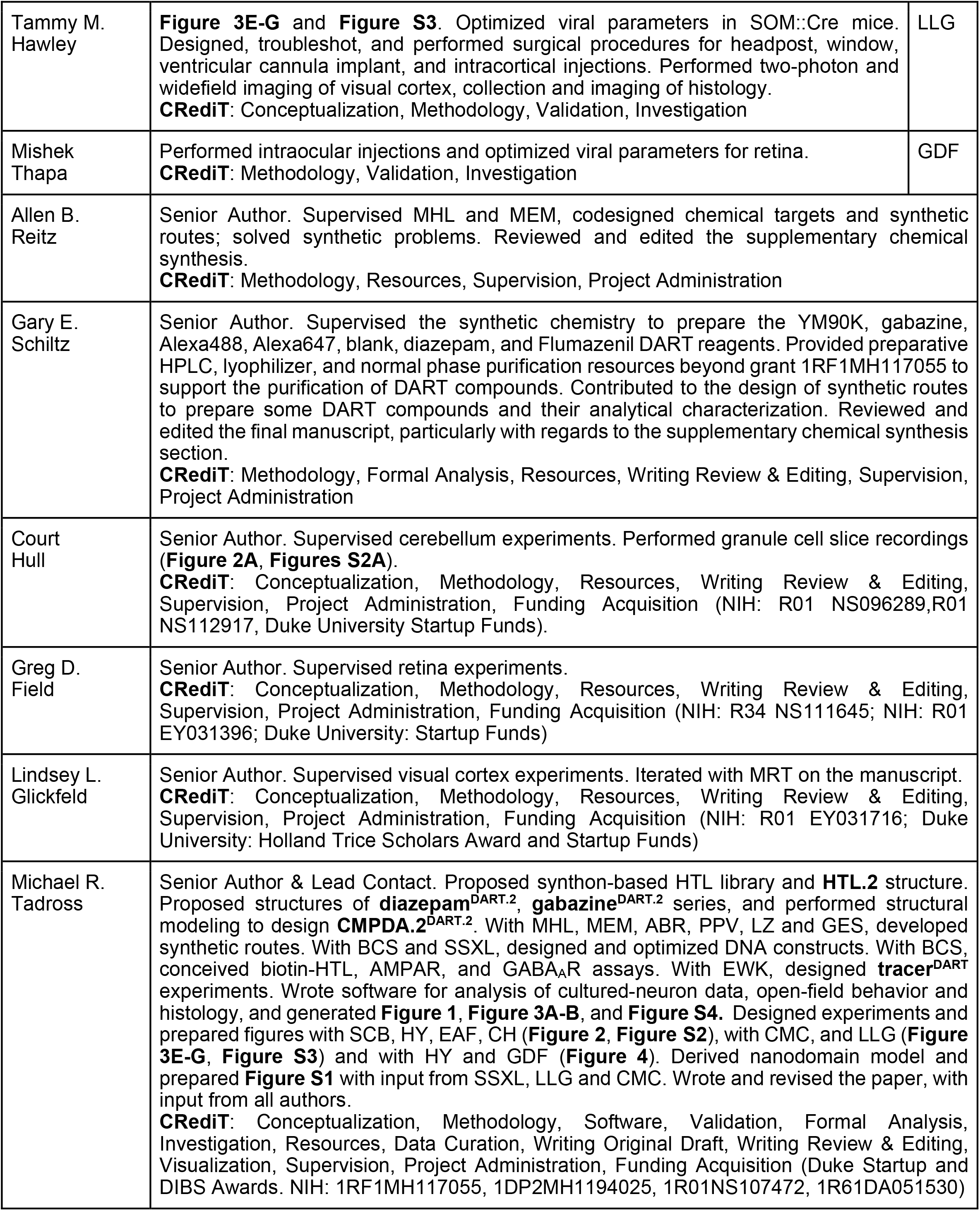
Detailed Author Contributions. **Attribution Process:** Each Principle Investigator (**PI**, right column) worked with their corresponding team members to outline individual contributions across a standard set of CRediT domains in the link below. Contributions were entered and edited on a live google-sheet, allowing all individuals to discuss and reach consensus. Authors were advised of a mediation process, whereby disagreements could be voiced, discussed, and resolved with the relevant PIs and the Lead PI. Decisions regarding the delineation between authorship and acknowledgement were decided by the supervising PI of each individual, and authorship order was decided by the lead PI with input and discussions with all authors. https://www.cell.com/pb/assets/raw/shared/guidelines/CRediT-taxonomy.pdf

