## Supplementary Document 1 for "Thousandfold Cell-Specific Pharmacology of Neurotransmission"

### Supplementary Document 1—Chemical Synthesis

- R1 — biotin-PEG$_{12}$-HTL.1
- R2 — biotin-PEG$_{12}$-HTL.2
- R3 — azide-PEG$_{36}$-HTL.1 (*azide*$^{\mathbf{DART.1}}$)
- R4 — azide-PEG$_{36}$-HTL.2 (*azide*$^{\mathbf{DART.2}}$)
- R5 — YM90K.1$^{DART.1}$
- R6 — YM90K.1$^{DART.2}$
- R7 — gabazine.1$^{DART.2}$
- R8 — gabazine.5$^{DART.2}$
- R9 — gabazine.7$^{DART.2}$
- R10 — Alexa488.1$^{DART.2}$
- R11 — Alexa647.1$^{DART.2}$
- R12 — blank.1$^{DART.2}$
- R13 — diazepam.1$^{DART.2}$
- R14 — flumazenil.1$^{DART.2}$
- R15 — CMPDA.1$^{DART.1}$
- R16 — CMPDA.2$^{DART.2}$

### General Chemical Synthesis Information

All chemical reagents were obtained from commercial suppliers and used without further purification unless otherwise stated. Anhydrous solvents were purchased from Sigma-Aldrich, and dried over 3 Å molecular sieves when necessary. Normal-phase flash column chromatography was performed using Biotage KP-Sil 50 µm silica gel columns and ACS grade solvents on a Biotage Isolera flash purification system or Teledyne Combiflash using Silicycle silica gel columns. Reverse phase preparative HPLC was performed with the following conditions: Phenomenex Gemini-NX C18, 110 Å, 150 x 21.2 mm; 5 µm. Eluting with a gradient of acetonitrile:water specified for each DART with 0.1 % formic acid over 25 min, then 1 min at 100 % acetonitrile. For Alexa647.1$^{DART.2}$ (R11), 0.1 % TFA rather than 0.1 % formic acid was used and the flow rate was 20 mL/min. For Alexa488.1$^{DART.2}$ (R10), the following prep HPLC conditions were used: Phenomenex Kinetex C18 100 Å, 50 x 30 mm; 5 µm, and a gradient of 15 to 80 % acetonitrile in water (0.1 % FA) at 50 mL/min for 12 min. Analytical thin layer chromatography (TLC) was performed on EM Reagent 0.25 mm silica gel 60 F254 plates and visualized by UV light. Proton (1 H), and carbon (13C) NMR spectra were recorded on a 500 MHz Bruker Avance III with direct cryoprobe spectrometer. Chemical shifts were reported in ppm (δ) and were referenced using residual nondeuterated solvent as an internal standard ($CDCl_3$ at 7.24 ppm for 1 H-NMR and 77.0 for 13C-NMR. $CD_3OD$ at 3.33 ppm for 1 H-NMR and 47.6 for 13C-NMR. DMSO-$d_6$ at 2.52 ppm for 1 H-NMR and 39.9 ppm for 13C-NMR). Proton coupling constants are expressed in hertz (Hz). The following abbreviations are used to denote spin multiplicity for proton NMR: s = singlet, d = doublet, t = triplet, q = quartet, m = multiplet, br.s = broad singlet, dd = doublet of doublets, dt = doublet of triplets, quin = quintet, tt = triplet of triplets. Low-resolution liquid chromatography/mass spectrometry (LCMS) was performed on a Waters Acquity-H UPLC/MS system with a 2.1 mm × 50 mm, 1.7 µm, reversed phase BEH C18 column and LCMS grade solvents. A gradient elution from 95% water +0.1% formic acid/5% acetonitrile +0.1% formic acid to 95% acetonitrile +0.1% formic acid/5% water +0.1% formic acid over 2 min plus a further minute continuing this mixture at a flow rate of 0.85 mL/min was used as the eluent (for Alexa647.1$^{DART.2}$ (R11), 0.1 % TFA rather than 0.1 % FA was used). Total ion current traces were obtained for electrospray positive and negative ionization (ESI+/ESI-). High-resolution mass spectra were obtained using an Agilent 6210 LC-TOF spectrometer in the positive ion mode using electrospray ionization with an Agilent G1312A HPLC pump and an Agilent G1367B autoinjector at the Integrated Molecular Structure Education and Research Center (IMSERC), Northwestern University. All microwave-assisted reactions were carried out in a Biotage® initiator. All IUPAC compound names were generated using Chemdraw Version 19.1.1.21 or Instant J Chem.

### Synthesis of R1—biotin-PEG$_{12}$-HTL.1

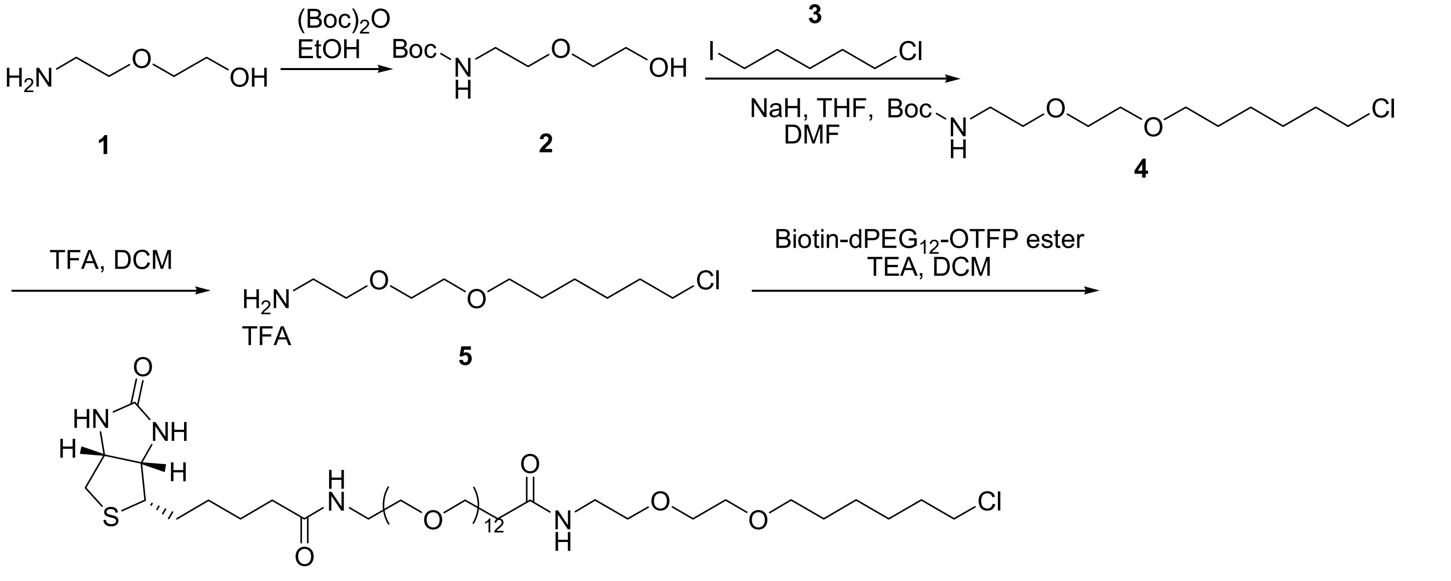

**biotin-PEG$_{12}$-HTL.1 (R1)**

Step 1: *tert*-butyl (2-(2-hydroxyethoxy)ethyl)carbamate, **2**

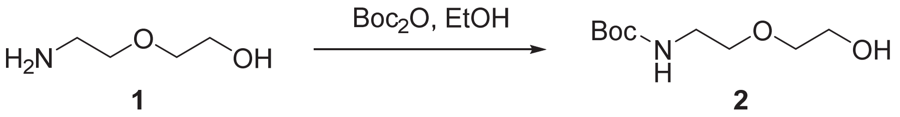

To a solution of 2-(2-aminoethoxy)ethan-1-ol (3.2 g, 30.4 mmol) in EtOH (10 mL) was added di-*tert*-butyl dicarbonate (6.64 g, 30.4 mmol ) and reaction mixture was maintained at room temperature for 24 h. After the complete consumption of starting material (LC-MS), the volatiles were removed *in vacuo*, and the residue was concentrated. Purification by flash chromatography (24 g silica cartridge, 0-10% MeOH/$CH_2Cl_2$) afforded the title compound in 5.8 g, 92% yield. (ES-LCMS) *m*/*z* 228 $(M+Na)^+$.

Step 2: *tert*-butyl (2-(2-((6-chlorohexyl)oxy)ethoxy)ethyl)carbamate, **4**

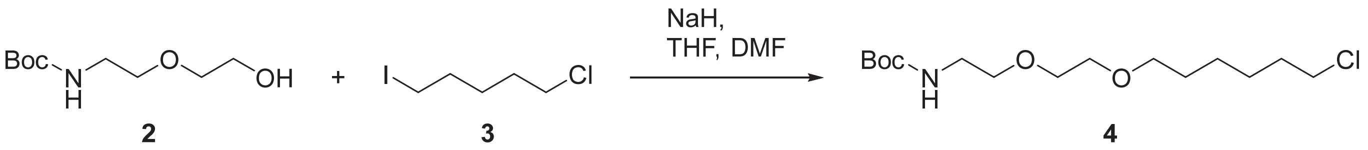

In a flame-dried 250 mL round-bottom flak under nitrogen was placed sodium hydride (60% dispersion in mineral oil, 0.76 g, 19.0 mmol) followed by dry THF (30 mL). The mixture was cooled in an ice bath to 0 °C and a solution of *tert*-butyl (2-(2-hydroxyethoxy)ethyl)carbamate (3 g, 14.6 mmol) in dry THF (5 mL) was slowly added. After the reaction was stirred for one hour at 0 °C, a solution of 1-chloro-5-iodopentane (5.41 g, 21.9 mmol) in dry THF (5 mL) was slowly added. The reaction was kept at 0 °C overnight then poured into water and the product extracted with ethyl acetate (3x). The combined organic extracts were filtered to remove dark brown precipitates then washed with brine, dried over $Na_2SO_4$, filtered and concentrated *in vacuo*. The crude material was purified by flash chromatography (40 g silica column, 0-100% EtOAc/hexane gradient) to obtain the product as a light yellow oil (3.1 g, 65% yield). (ES-LCMS) *m*/*z* 343 $(M+Na)^+$.

Step 3: 2-(2-((6-chlorohexyl)oxy)ethoxy)ethan-1-amine TFA salt, **5**

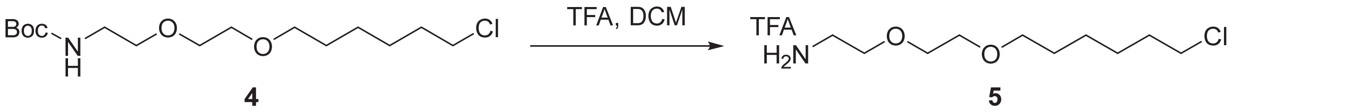

To a solution of *tert*-butyl (2-(2-((6-chlorohexyl)oxy)ethoxy)ethyl)carbamate (3 g, 9.28 mmol) in dichloromethane (20.0 mL) added trifluoroacetic acid (5 mL) and maintained at room temperature for 3.0 h. The reaction was concentrated under vacuum and triturated with two 5 ml portions of ether and ether decanted away and residual ether of was concentrated to yield a dark yellow product 3.11 g, 104% yield. $^{1}H$ NMR (CHLOROFORM-d ,300MHz): δ = 11.01 - 11.63 (m, 1 H), 3.71 - 3.80 (m, *J*=10.0 Hz, 2 H), 3.65 - 3.71 (m, 2 H), 3.63 (br. s., 2 H), 3.43 - 3.57 (m, 4 H), 3.16 - 3.31 (m, 2 H), 1.76 (s, 2 H), 1.59 (s, 2 H), 1.25 - 1.50 ppm (m, 4 H); (ES-LCMS) *m*/*z* 224 (M+Na)$^{+}$.

Step 4: *N*-(2-(2-((6-chlorohexyl)oxy)ethoxy)ethyl)-1-(5-((3a*S*,4*S*,6a*R*)-2-oxohexahydro-1*H*-thieno[3,4-*d*]imidazol-4-yl)pentanamido)-3,6,9,12,15,18,21,24,27,30,33,36-dodecaoxanonatriacontan-39-amide, **R1**

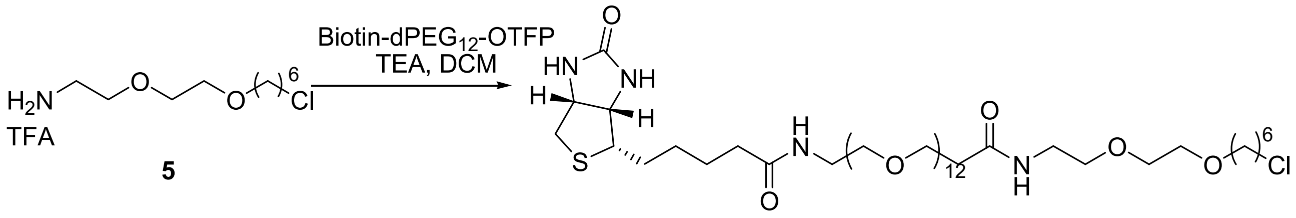


**biotin-PEG$_{12}$-HTL.1 (R1)**

DIPEA (125 µL, 0.717 mmol) was added to a solution of 2-(2-((6-chlorohexyl)oxy)ethoxy)ethan-1-amine TFA salt (**5**) (20 mg, 0.089 mmol) and biotin-dPEG12-OTFP ester (71 mg, 0.071 mmol, Quanta Biodesign) in DCM (1 mL). After two hours the solvent was removed *in vacuo* and the residue was purified by Gilson prep HPLC (Sunfire Prep C18 OBD, 10 µm, 30 x 150 mm column, 15-75% ACN/$H_2O$ (both containing 0.1% TFA) gradient). The product fractions were combined and concentrated to remove the acetonitrile before freeze drying overnight to obtain a colorless semi-solid product (20 mg, 21 % yield). ES-LCMS: m/z = 1049.5(M+H)$^{+}$; $^{1}H$ NMR (METHANOL-d$_4$ ,300MHz): δ = 4.45 - 4.53 (m, 1 H), 4.26 - 4.35 (m, 1 H), 3.45 - 3.76 (m, 58 H), 3.33 - 3.41 (m, 5 H), 3.21 (dd, *J*=8.8, 4.7 Hz, 1 H), 2.87 - 2.98 (m, 1 H), 2.66 - 2.76 (m, 1 H), 2.45 (s, 2 H), 2.22 (s, 2 H), 1.32 - 1.84 ppm (m, 14 H); HRMS (ESI+ ): m/z calcd for $C_{47}H_{90}ClN_4O_{17}S$: 1049.5711 [M+H]$^{+}$; found: 1049.5704, [M+H]$^{+}$

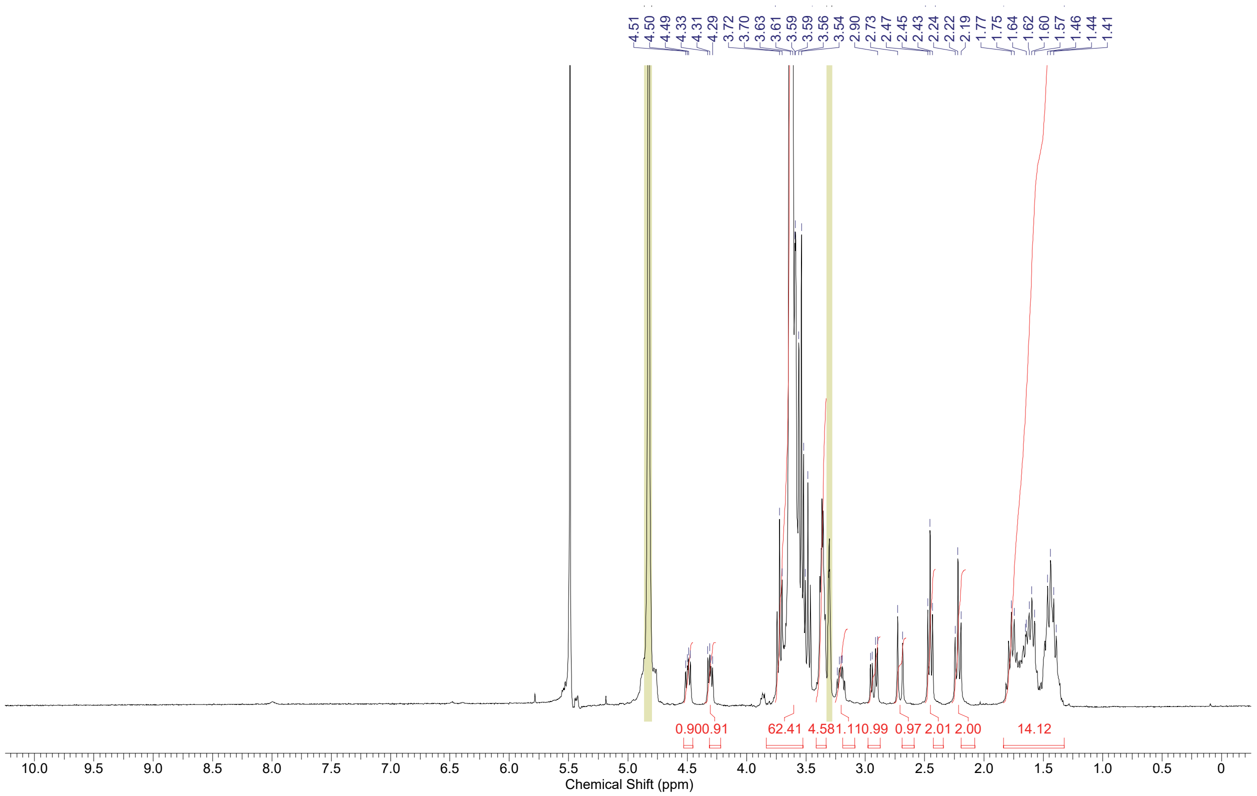


**$^{1}H$ NMR of biotin-PEG$_{12}$-HTL.1 (R1)**

### Synthesis of R2—biotin-PEG$_{12}$-HTL.2

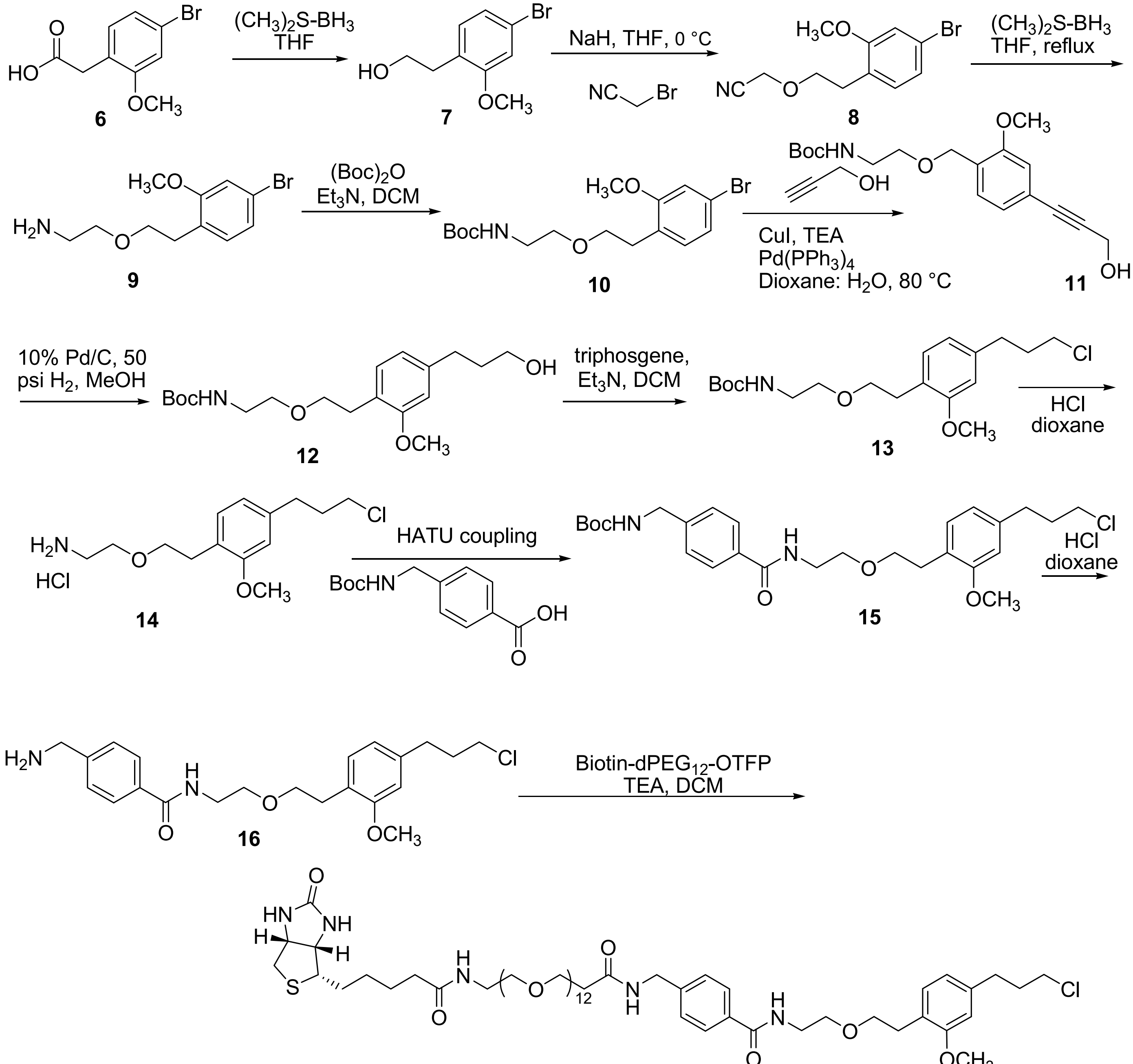

**biotin-PEG$_{12}$-HTL.2 (R2)**

Step 1: 2-(4-bromo-2-methoxyphenyl)ethan-1-ol, **7**

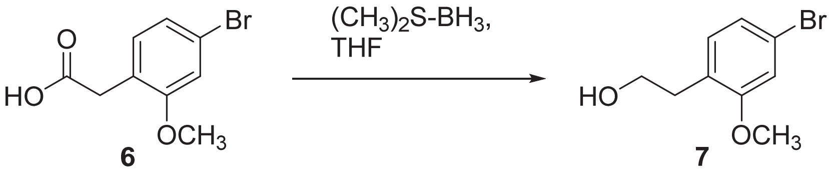

To a solution of 2-(4-bromo-2-methoxyphenyl)acetic acid (10.0 g, 41.0 mmol, CombiBlocks) in THF (100 mL) at 0 °C was added dropwise borane dimethylsulfide (21.1 mL, 246.0 mmol), keeping the temperature below 5 °C during the addition. The reaction was allowed to stir overnight as the ice bath warmed to ambient temperature. The reaction was poured into dry methanol (100 mL), stirred for 5 minutes and then the solvent was removed *in vacuo*. The crude product was obtained as a yellowish brown oil (13.45 g, 98% yield). $^1$H NMR (METHANOL-d$_4$, 300MHz): δ = 6.94 - 7.09 (m, *J*=8.8 Hz, 3 H), 3.78 (s, 3 H), 3.67 (t, *J*=7.0 Hz, 2 H), 2.77 ppm (t, *J*=7.0 Hz, 2 H); ES-LCMS: m/z 232 (M+H)$^+$

Step 2: [2-(4-Bromo-2-methoxy-phenyl)-ethoxy]-acetonitrile, **8**

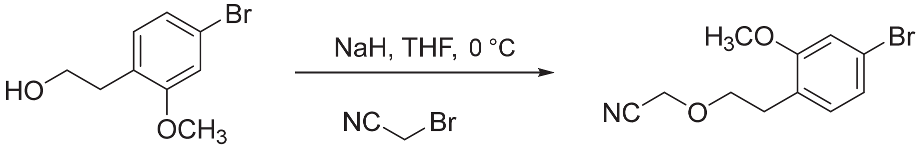


**7** **8**

In a flame-dried 100 mL round-bottom flak under nitrogen was placed sodium hydride (60% dispersion in mineral oil, 9.35 g, 70.1 mmol) followed by dry THF (90 mL). The mixture was cooled in an ice bath to 0 °C and a solution of 2-(4-bromo-2-methoxy-phenyl)-ethanol (13.45 g, 58.4 mmol) in dry THF (50 mL) was slowly added. After the reaction was stirred for one hour at 0 °C, a solution of bromoacetonitrile (4.89 mL, 70.1 mmol) in dry THF (50 mL) was slowly added. The reaction was kept at 0 °C overnight then poured into water and the product extracted with ethyl acetate (3x). The combined organic extracts were filtered to remove dark brown precipitates then washed with brine, dried over $Na_2SO_4$, filtered and concentrated *in vacuo*. The crude material was purified by flash chromatography (40 g silica column, 0-2% MeOH/DCM gradient) to obtain the product as a colorless oil (8.57 g, 54% yield). $^1$H NMR (CHLOROFORM-d ,300MHz): δ = 7.03 (br. s., 3 H), 4.23 (br. s., 2 H), 3.67 - 3.91 (m, 5 H), 2.89 ppm (d, *J*=5.9 Hz, 2 H); ES-LCMS: m/z 271.5 $(M+H)^+$

Step 3: 2-[2-(4-Bromo-2-methoxy-phenyl)-ethoxy]-ethylamine, **9**

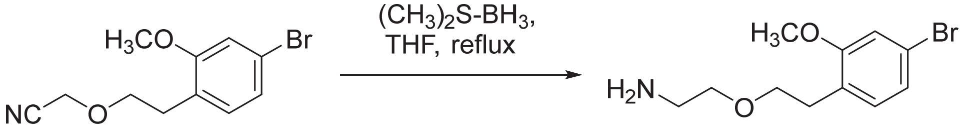


**8** **9**

A 2M solution of borane dimethyl sulfide complex in THF (47.8 mL, 95.5 mmol) was added to a solution of [2-(4-bromo-2-methoxy-phenyl)-ethoxy]-acetonitrile (8.57 g, 31.8 mmol) in THF (50 mL) and the mixture was heated at 70 °C for 3 hours. The reaction as cooled and then quenched with methanol (50 mL). The reaction was concentrated *in vacuo* and then concentrated twice more from methanol. The crude oil was place on under high vacuum overnight to afford compound **9** (5.2 g, 60% yield). ES-LCMS: m/z 274.5, 276.4 (M, $M+2H)^+$

Step 4: *tert*-butyl (2-(4-bromo-2-methoxyphenethoxy)ethyl)carbamate, **10**

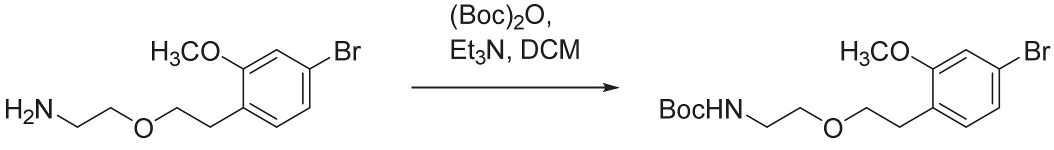


**9** **10**

To a solution of 2-(4-bromo-2-methoxyphenethoxy)ethan-1-amine (8.6 g, 31.3 mmol) in DCM (50 mL) was added di-*tert*-butyl dicarbonate (8.9 g, 40.8 mmol ) and the reaction mixture was maintained at 45 °C for 24 h. After the complete consumption of starting material (LC-MS), the volatiles were removed *in vacuo*, and the residue was dissolved in dichloromethane (30 mL). The solution was washed successively with 1% HCl (60 mL), brine (2 x 30 mL), and deionized water (30 mL), then dried over $MgSO_4$, filtered, and concentrated. Purification by flash chromatography (40 g silica cartridge, 0-100% EtOAc/Hexanes) afforded the title compound in 9 0g, 77% yield. $^1$H NMR (CHLOROFORM- δ ,300MHz): d = 6.92 - 7.12 (m, 3 H), 4.70 - 4.93 (m, 1 H), 3.73 - 3.93 (m, 3 H), 3.54 - 3.66 (m, 2 H), 3.43 - 3.53 (m, 2 H), 3.21 - 3.33 (m, 2 H), 2.76 - 2.90 (m, 2 H), 1.44 ppm (br. s., 9 H); (ESI) *m/z* 397, 399 $(M+Na, M+Na+2)^+$.

Step 5: *tert*-butyl (2-((4-(3-hydroxyprop-1-yn-1-yl)-2-methoxybenzyl)oxy)ethyl)carbamate, **11**

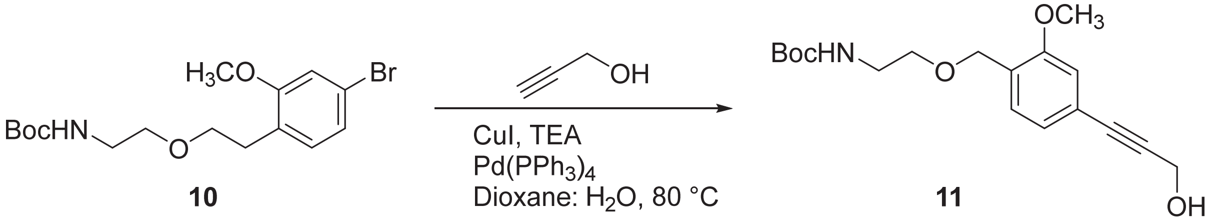


To mixture of compounds *tert*-butyl (2-(4-bromo-2-methoxyphenethoxy)ethyl)carbamate (7.0 g, 19.0 mmol), $K_2CO_3$ (5.25 g, 38.0 mmol), copper(I) iodide (0.36 g, 1.90 mmol), $Pd(PPh)_4$ (1.06 g, 0.95 mmol), and dioxane/$H_2O$ (100 mL/10 mL) were added to a round bottom flask. The reaction mixture was placed under vacuum and then vented with nitrogen (3 times). The prop-2-yl-1-ol (13.3 g, 190.0 mmol) was added and the reaction heated to 75°C overnight. Reaction monitored by LC-MS until complete. The solvent was removed under reduced pressure and crude residue purified by flash chromatography (40 g silica cartridge, 0-70% EtOAc/Hexanes) to afford the title compound in 2.16 g 33% yield. $^1H$ NMR (CHLOROFORM-d ,300MHz): δ = 7.03 - 7.11 (m, 1 H), 6.93 - 7.01 (m, 1 H), 6.86 - 6.92 (m, 1 H), 4.77 - 5.01 (m, 1 H), 4.46 (br. s., 2 H), 4.24 - 4.33 (m, 1 H), 3.78 (s, 3 H), 3.59 (s, 2 H), 3.46 (br. s., 2 H), 3.20 - 3.31 (m, 2 H), 2.85 (s, 2 H), 1.43 ppm (s, 9 H); (ESI) *m/z* 372 $(M+Na)^+$.

Step 6: *tert*-butyl (2-(4-(3-hydroxypropyl)-2-methoxyphenethoxy)ethyl)carbamate, **12**

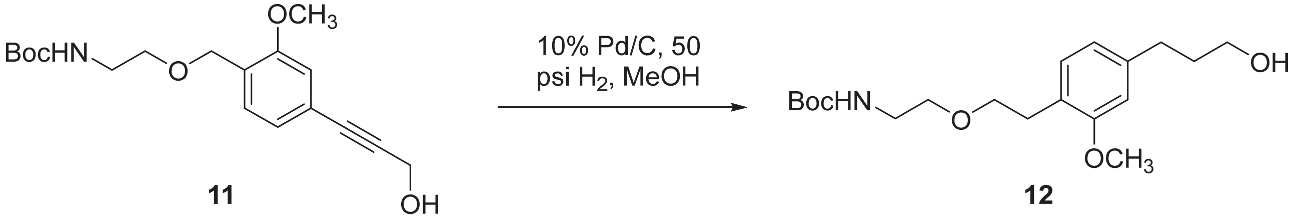


*tert*-Butyl (2-((4-(3-hydroxyprop-1-yn-1-yl)-2-methoxybenzyl)oxy)ethyl)carbamate (1.8 g, 5.15 mmol) in EtOH (20 mL) was hydrogenated using 5% Pt/C (~100 mg) in a small Parr vessel (250 mL) with hydrogen (55 psi) overnight. The reaction was filtered through celite while under nitrogen and the filtrate was then concentrated *in vacuo*. The crude product was purified by flash chromatography (24 g silica, 0-100% EtOAc/hexane gradient) to obtain product **12** (1.11 g, 61% yield). $^1H$ NMR (CHLOROFORM-d ,300MHz): δ = 6.99 - 7.15 (m, 1 H), 6.69 (m, 2 H), 4.91 (br. s., 1 H), 3.80 (s, 3 H), 3.56 - 3.72 (m, *J*=6.2, 6.2 Hz, 4 H), 3.49 (t, *J*=5.0 Hz, 2 H), 3.27 (d, *J*=5.3 Hz, 2 H), 2.85 (t, *J*=7.3 Hz, 2 H), 2.61 - 2.74 (m, *J*=7.6, 7.6 Hz, 1 H), 1.81 - 1.95 (m, 2 H), 1.44 ppm (s, 9 H); ES-LCMS: m/z 377 $(M+Na)^+$

Step 7: *tert*-butyl (2-(4-(3-chloropropyl)-2-methoxyphenethoxy)ethyl)carbamate, **13**

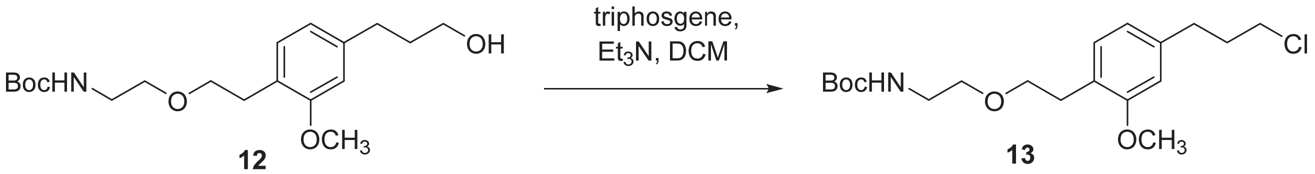


To *tert*-butyl (2-(4-(3-hydroxypropyl)-2-methoxyphenethoxy)ethyl)carbamate (1.11 g, 3.14 mol) and triethylamine (1.09 mL, 7.85 mmol) in DCM (20 mL) was added triphosgene (0.46 g, 1.57 mmol) and the reaction stirred at room temperature for 1h. Aqueous sodium bicarbonate was added and the water layer extracted twice with dichloromethane. The organic extracts were combined, dried ($Na_2SO_4$), and concentrated under reduced pressure to provide crude product. Purification by flash chromatography (24 g silica cartridge, 0-10% MeOH/$CH_2Cl_2$) afforded the title compound (1.05 g, 89% yield). $^1H$ NMR (CHLOROFORM-d ,300MHz): δ = 7.07 (d, *J*=7.6 Hz, 1 H), 6.65 - 6.77 (m, 2 H), 4.85 (br. s., 1 H), 3.82 (s, 3 H), 3.44 - 3.66 (m, 6 H), 3.28 (q, *J*=4.9 Hz, 2 H), 2.86 (t, *J*=7.0 Hz, 2 H), 2.75 (t, *J*=7.3 Hz, 2 H), 2.01 - 2.14 (m, 2 H), 1.45 ppm (s, 9 H); (ES-LCMS) *m/z* 395 $(M+Na)^+$.

Step 8: 2-(4-(3-chloropropyl)-2-methoxyphenethoxy)ethan-1-amine, **14**

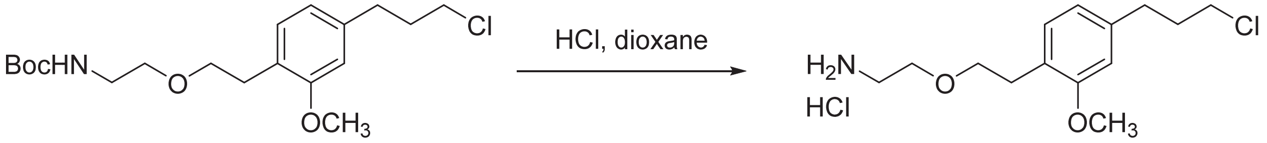


**13** **14**

To a solution of *tert*-butyl (2-(4-(3-chloropropyl)-2-methoxyphenethoxy)ethyl)carbamate (203 mg, 0.55 mmol) in dichloromethane (2.0 mL) was added 4N HCl in dioxane (0.5 mL) and maintained at room temperature for 3.0h. The reaction was concentrated under vacuum and triturated with two portions of ether (5 ml ) and the ether decanted off to yield product (160.2 mg, 99% yield). $^1$H NMR (CHLOROFORM-d ,300MHz): δ = 8.29 (br. s., 2 H), 7.07 (d, *J*=7.6 Hz, 1 H), 6.62 - 6.78 (m, 2 H), 3.80 (s, 3 H), 3.62 - 3.76 (m, *J*=5.0, 5.0 Hz, 4 H), 3.52 (t, *J*=6.4 Hz, 2 H), 3.20 (br. s., 2 H), 2.88 (t, *J*=7.0 Hz, 2 H), 2.73 (t, *J*=7.3 Hz, 2 H), 1.92 - 2.20 ppm (m, 2 H); (ES-LCMS) *m*/*z* 272 $(M+H)^+$.

Step 9: *tert*-butyl (4-((2-(4-(3-chloropropyl)-2-methoxyphenethoxy)ethyl)carbamoyl)benzyl)carbamate, **15**

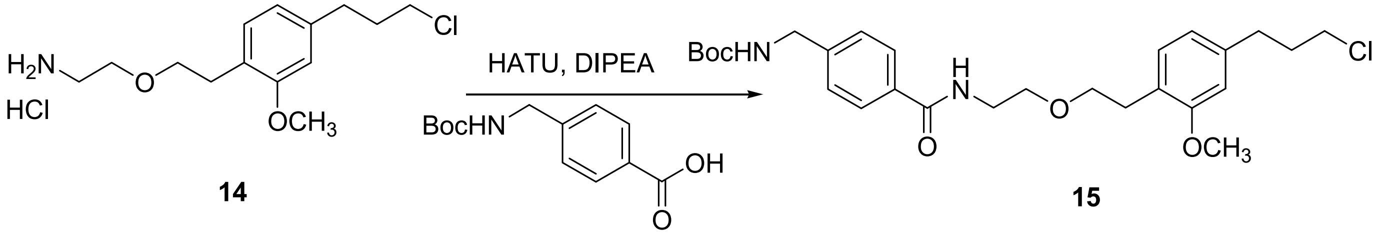


**14** **15**

2-(4-(3-Chloropropyl)-2-methoxyphenethoxy)ethan-1-amine (276 mg, 1.01 mmol), 4-Boc-aminomethyl)benzoic acid (305.9 mg, 1.21 mmol), HATU (964.5 mg, 2.53 mmol), and DIPEA (708 µL, 4.05 mmol) were combined in DMF (5 mL). The mixture was stirred for 4 hours then poured into water and the product extracted with ethyl acetate (3x). The combined organic extracts were washed with brine then dried over $Na_2SO_4$, filtered and concentrated *in vacuo*. The crude material was purified by flash chromatography (24 g silica, 15-70% EtOAc/hexane gradient) to obtain the product (339.8 mg, 66% yield). $^1$H NMR (CHLOROFORM-d, 300MHz): δ = 7.68 (d, *J*=7.6 Hz, 2 H), 7.34 (d, *J*=8.2 Hz, 2 H), 7.08 (d, *J*=7.6 Hz, 1 H), 6.67 (br. s., 2 H), 6.44 (br. s., 1 H), 4.86 - 5.00 (br. s, 1 H), 4.22 - 4.47 (m, 2 H), 3.80 (s, 3 H), 3.47 - 3.71 (m, 8 H), 2.61 - 2.98 (m, *J*=6.7, 6.7 Hz, 4 H), 2.05 (t, *J*=6.7 Hz, 2 H), 1.47 ppm (s, 9 H); (ES-LCMS) *m*/*z* 506 $(M+H)^+$.

Step 10: 4-(aminomethyl)-N-(2-(4-(3-chloropropyl)-2-methoxyphenethoxy)ethyl)benzamide, **16**

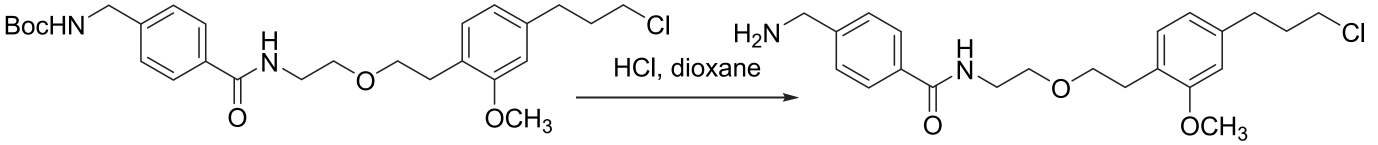


**15** **16**

To a solution of *tert*-butyl (2-(4-(3-chloropropyl)-2-methoxyphenethoxy)ethyl)carbamate (203 mg, 0.55 mmol) in dichloromethane (2.0 mL) was added 4N HCl (0.5 mL) in dioxane and the reaction maintained at room temperature for 3.0 h. The reaction was then concentrated under vacuum and triturated with two portions of ether (5 ml) that were subsequently decanted off to yield product (160.2 mg, 99% yield). $^1$H NMR (CHLOROFORM-d ,300MHz): δ = 8.41 (br. s., 2 H), 7.52 (br. s., 2 H), 7.31 (br. s., 2 H), 6.96 - 7.04 (br. s., 1 H), 6.59 - 6.70 (m, 2 H), 3.99 (s., 2 H), 3.70 - 3.78 (s, 3 H), 3.39 - 3.67 (m, 8 H), 2.83 (m, 2 H), 2.57 - 2.74 (m, 2 H), 1.90 - 2.09 ppm (m, *J*=5.3 Hz, 2 H); (ES-LCMS) *m*/*z* 406 $(M+H)^+$.

Step 11: Synthesis of reagent Biotine-PEG12-HTL.2 (**R2**)

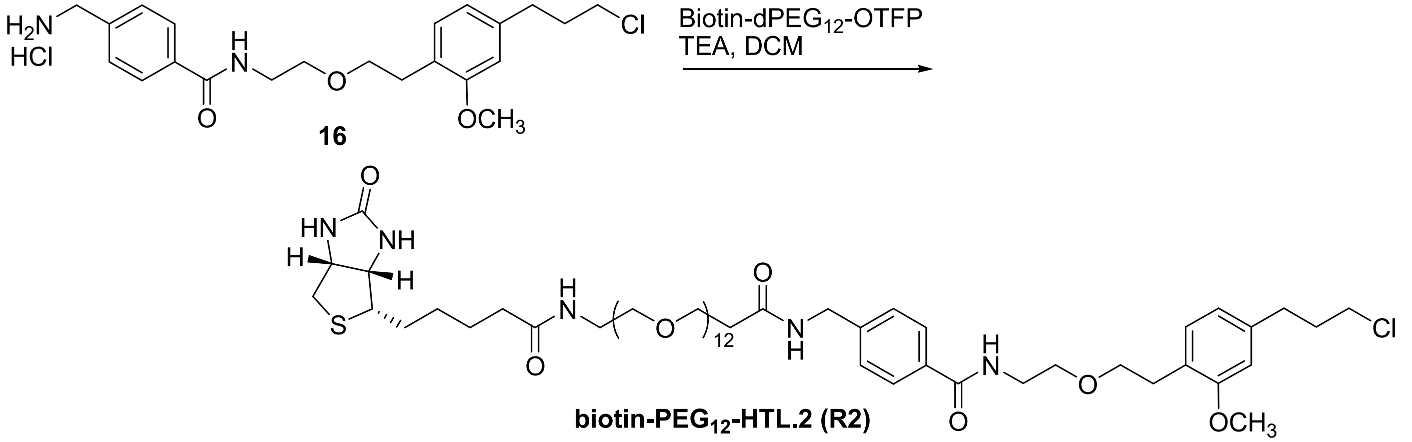


**biotin-PEG12-HTL.2 (R2)**

To a solution of solution of 2-(2-((6-chlorohexyl)oxy)ethoxy)ethan-1-amine HCl salt (**16**) (20 mg, 0.089 mmol) and biotin-dPEG12-OTFP (71 mg, 0.071 mmol, Quanta Biodesign) in DCM (1 mL) was added DIPEA (69 µL, 0.395 mmol) and the reaction stirred at room temperature overnight. The solvent was then removed *in vacuo* and the residue was purified by Gilson prep HPLC (Sunfire Prep C18 OBD, 10 µm, 30 x 150 mm column, 15-75% ACN/$H_2O$ gradient (both solvents containing 0.1% TFA). The product fractions were combined and concentrated to remove acetonitrile before freeze drying overnight to obtain a colorless semi-solid product (20 mg, 21 % yield). $^1$H NMR (METHANOL-d$_4$, 300MHz): δ = 7.77 (d, *J*=8.2 Hz, 2 H), 7.40 (d, *J*=8.2 Hz, 2 H), 7.05 (d, *J*=7.6 Hz, 1 H), 6.75 (s, 1 H), 6.64 (d, *J*=7.6 Hz, 1 H), 4.42 - 4.51 (m, 3 H), 4.29 (dd, *J*=7.9, 4.4 Hz, 1 H), 3.73 - 3.81 (m, 4 H), 3.56 - 3.68 (m, 48 H), 3.47 - 3.56 (m, 6 H), 3.36 (d, *J*=5.3 Hz, 2 H), 3.15 - 3.24 (m, *J*=8.5, 5.0 Hz, 1 H), 2.79 - 2.96 (m, *J*=12.9, 4.7 Hz, 3 H), 2.66 - 2.75 (m, *J*=2.3 Hz, 3 H), 2.52 (t, *J*=6.2 Hz, 2 H), 2.21 (t, *J*=7.3 Hz, 2 H), 1.96 - 2.08 (m, 2 H), 1.53 - 1.80 (m, *J*=2.3 Hz, 4 H), 1.44 ppm (q, *J*=7.0 Hz, 2 H); (ES-LCMS) *m*/*z* 1231.63 (M+H)$^+$; HRMS (ESI): m/z calcd for $C_{59}H_{97}ClN_5O_{18}S$: 1230.6236 [M+H]$^+$; found: 1230.6232, [M+H]$^+$

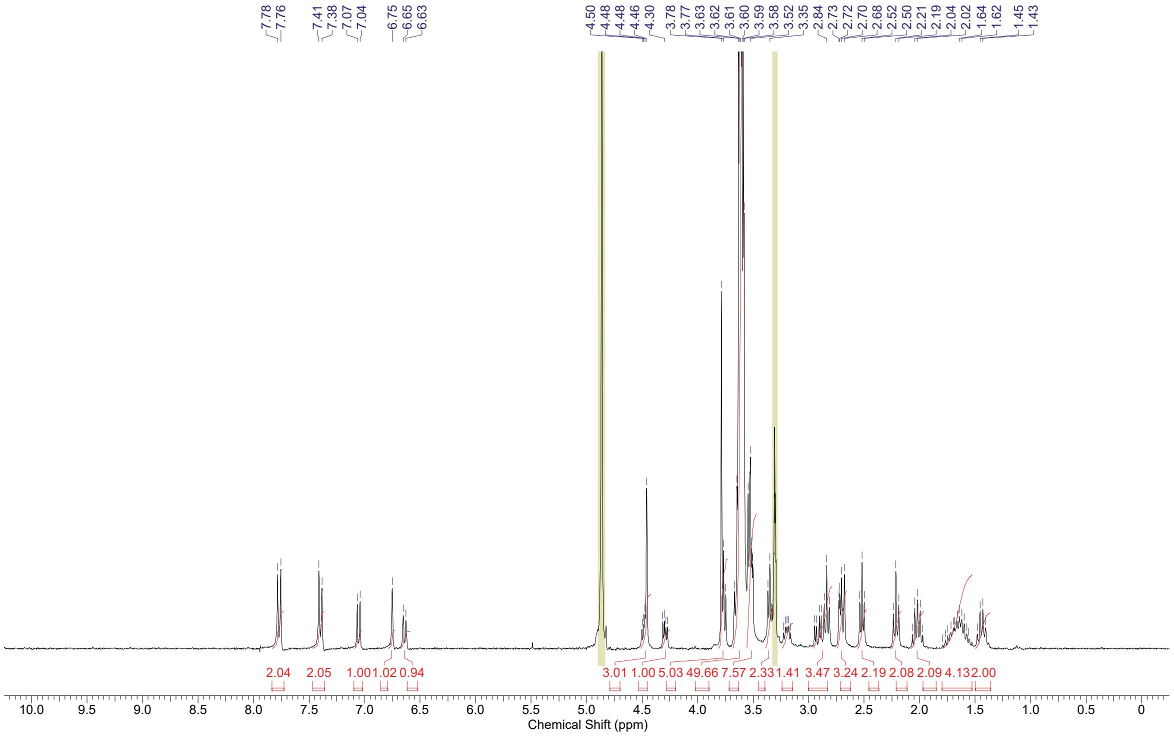


**$^1$H NMR of Biotin-PEG$_{12}$-HTL.2 (R2)**

### Synthesis of R3—azide-PEG$_{36}$-HTL.1 (*azide*$^{DART.1}$)

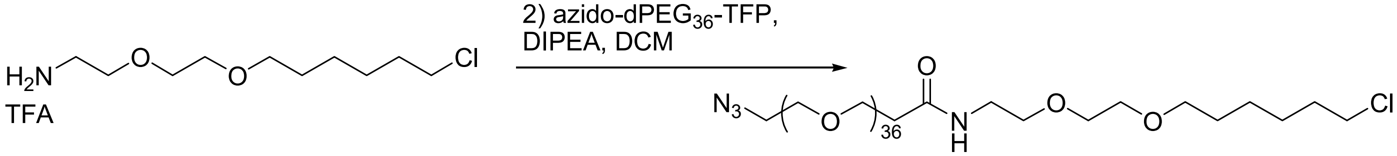


**5**

**azido-PEG$_{36}$-HTL.1 (R3)**

DIPEA (125 µL, 0.717 mmol) was added to a solution of 2-(2-((6-chlorohexyl)oxy)ethoxy)ethan-1-amine TFA salt (**5**) (20.5 mg, 0.109 mmol) and biotin-dPEG$_{36}$-OTFP (202 mg, 0.109 mmol, Quanta Biodesign) in DCM (2 mL). After two hours the solvent was removed *in vacuo* and the residue was purified by Gilson prep HPLC (Sunfire Prep C18 OBD, 10 µm, 30 x 150 mm column, 15-80% MeCN/$H_2O$ gradient (both solvents 0.1% TFA). The product fractions were combined and concentrated to remove the acetonitrile before freeze drying overnight to obtain a colorless semi-solid for the product (54 mg, 23 % yield). ES-LCMS: m/z = 1929.3$(M+Na)^+$; $^1$H NMR (METHANOL-$d_4$ ,300MHz): δ = 3.45 - 3.88 (m, 158 H), 3.33 - 3.42 (m, 6 H), 2.39 - 2.50 (m, 2 H), 1.71 - 1.84 (m, 2 H), 1.54 - 1.66 (m, 2 H), 1.32 - 1.53 ppm (m, 4 H); HRMS (ESI): m/z calcd for $C_{85}H_{169}ClN_4NaO_{39}$: 1928.0937 $[M+Na]^+$; found: 1928.0944, $[M+Na]^+$

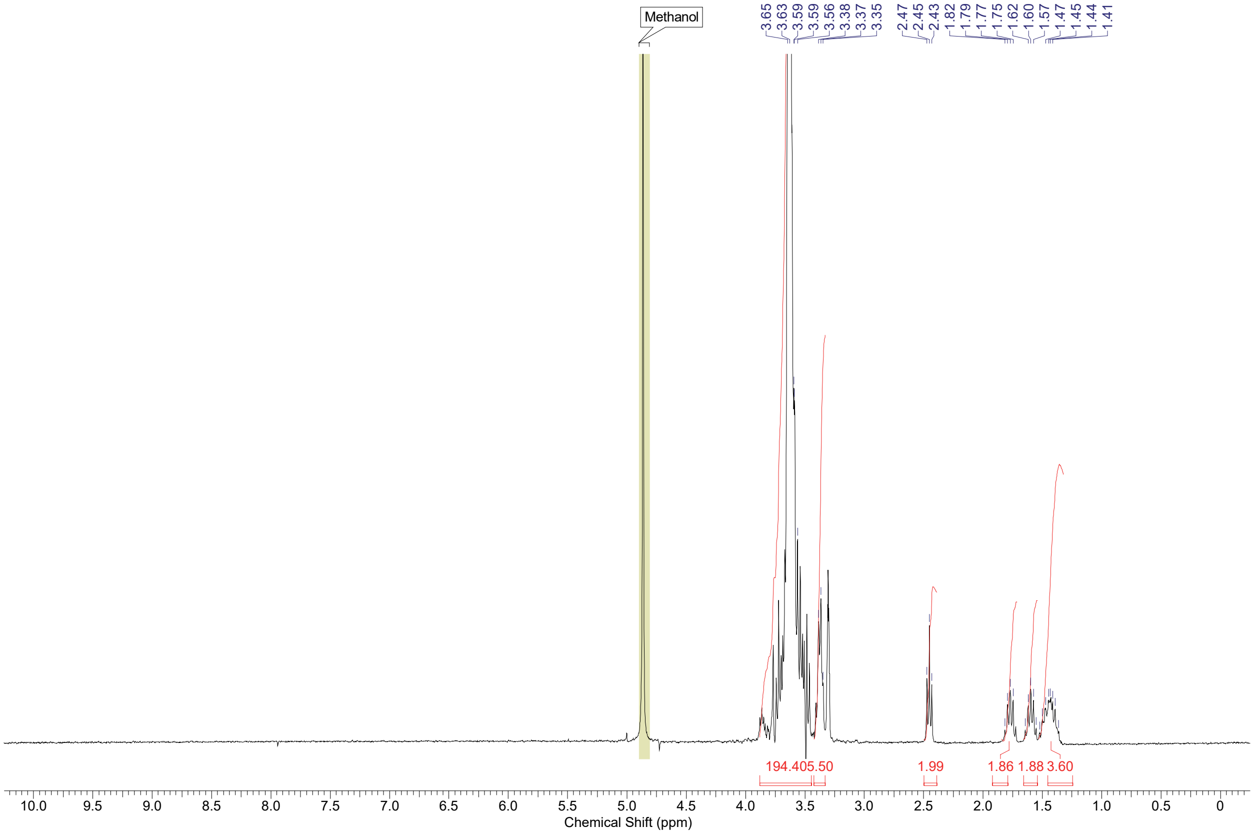


**H NMR of azide-PEG$_{36}$-HTL.1 (azide$^{DART.1}$, R3)**

### Synthesis of R4—azide-PEG36-HTL.2 (*azide*$^{DART.2}$)

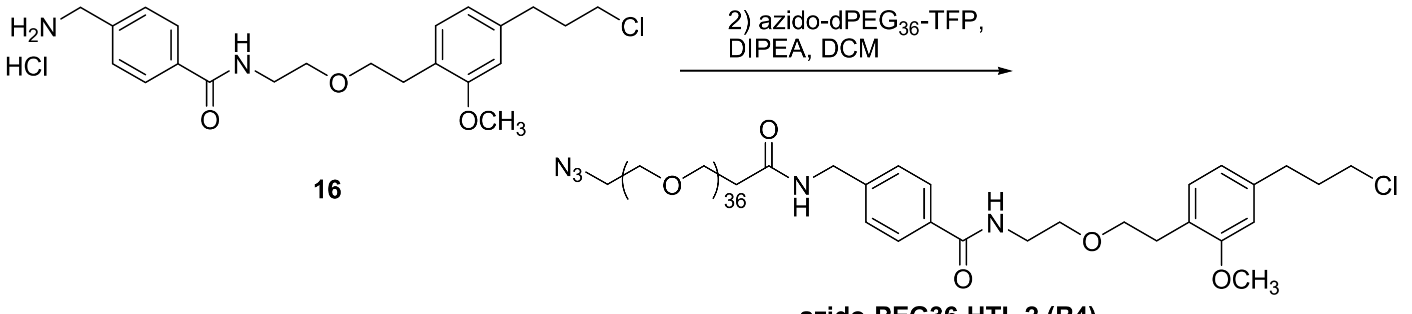


DIPEA (125 µL, 0.717 mmol) was added to a solution of 2-(2-((6-chlorohexyl)oxy)ethoxy)ethan-1-amine TFA salt (**5**) (20.5 mg, 0.109 mmol) and biotin-dPEG$_{36}$-OTFP ester (202 mg, 0.109 mmol, Quanta Biodesign) in DCM (2 mL). After two hours the solvent was removed *in vacuo* and the residue was purified by Gilson prep HPLC (Sunfire Prep C18 OBD, 10 µm, 30 x 150 mm column, 15-80% ACN/$H_2O$ gradient (both 0.1% TFA). The product fractions were combined and concentrated to remove the acetonitrile before freeze drying overnight to obtain a colorless semi-solid for the product (54 mg, 24%). $^1$H NMR (METHANOL-d$_4$ ,300MHz): δ = 7.77 (d, *J*=8.2 Hz, 2 H), 7.40 (d, *J*=8.2 Hz, 2 H), 7.04 (s, 1 H), 6.75 (s, 1 H), 6.59 - 6.69 (m, 1 H), 4.43 - 4.51 (m, 2 H), 3.78 - 3.80 (m, 3 H), 3.53 - 3.71 (m, 150 H), 3.34 - 3.41 (m, 4 H), 2.80 - 2.88 (m, 2 H), 2.66 - 2.75 (m, 2 H), 2.47 - 2.55 (m, 2 H), 2.02 ppm (s, 2 H); (ES-LCMS) *m/z* 1053.74 ((M+Na)/2)$^+$; HRMS (ESI): m/z calcd for $C_{97}H_{177}ClN_5O_{40}$: 2087.1645 [M+H]$^+$; found: 2087.1652, [M+H]$^+$

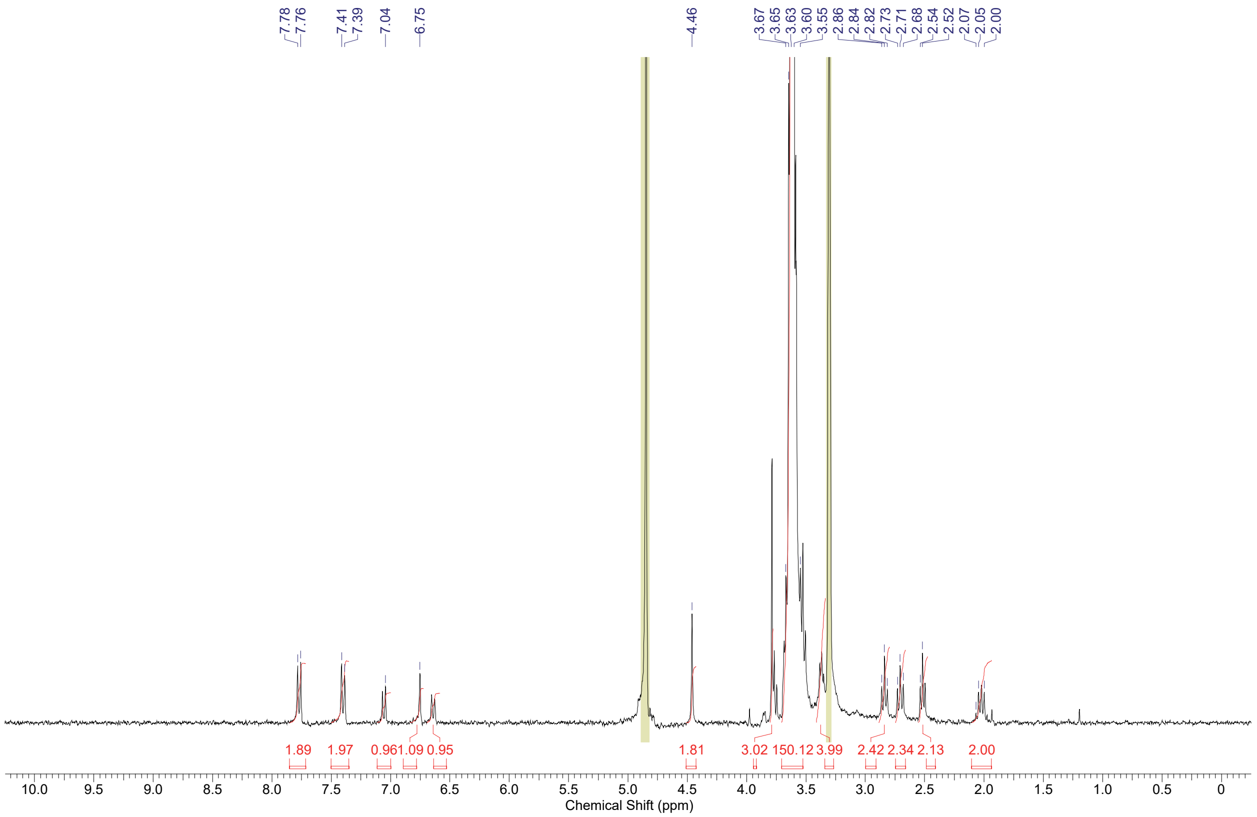


**$^1$H NMR of azide-PEG$_{36}$-HTL.2 (azide$^{DART.2}$, R4)**

**Synthesis of R5–YM90K.1DART.1**

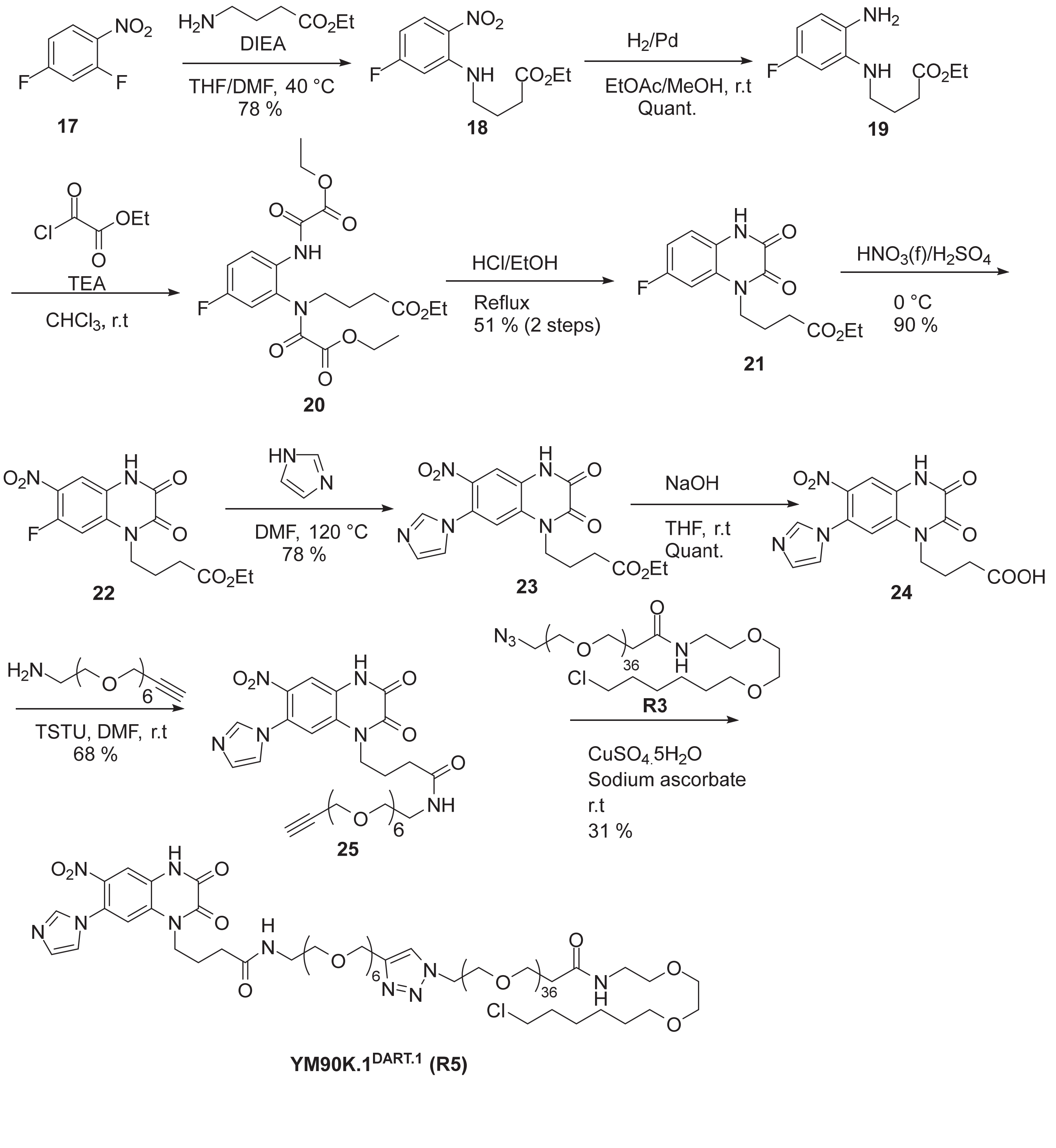

Step 1: Synthesis of ethyl 4-((5-fluoro-2-nitrophenyl)amino)butanoate, **18** (reported in reference 1)

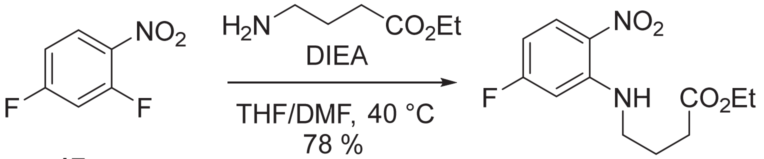


To a vial was added 2,4-difluoro-1-nitrobenzene, **17** (1.993 g, 1.374 mL, 12.53 mmol) and THF (10 mL)/DMF (5 mL) after which ethyl-4-aminobutanoate.HCl (2.000 g, 11.93 mmol) and triethylamine (1.449 g, 2.0 mL, 14.32 mmol) were added. The reaction was stirred at 40°C overnight after which the mixture was concentrated, taken up in DCM (~200 mL), washed with water (2 x 20 mL), brine (20 mL), filtered through an isolute phase separator, and concentrated to a yellow oil which was purified by silica gel chromatography eluting with 0 to 50 % ethyl acetate in hexanes to give ethyl-4-((5-fluoro-2-nitrophenyl)amino)butanoate, **18** (2.5 g, 78 %) as a yellow/orange oil. $^1$H NMR (500 MHz, Chloroform-*d*) δ 8.19 (dd, *J* = 9.5, 6.1 Hz, 2H), 6.52 (dd, *J* = 11.5, 2.6 Hz, 1H), 6.35 (ddd, *J* = 9.7, 7.2, 2.5 Hz, 1H), 4.14 (q, *J* = 7.2 Hz, 2H), 3.32 (q, *J* = 6.6 Hz, 2H), 2.44 (t, *J* = 7.0 Hz, 2H), 2.02 (p, *J* = 7.1 Hz, 2H), 1.30 – 1.20 (m, 3H).
(ES-LCMS) *m*/*z* 271.2 $(M+H)^+$.

Step 2: Synthesis of ethyl 4-((2-amino-5-fluorophenyl)amino)butanoate, **19** (reported in reference 1)

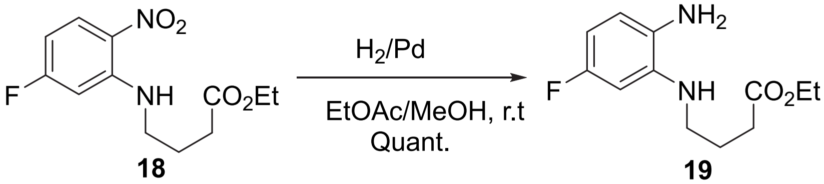


To a flask containing palladium on carbon (0.30 g, 0.28 mmol) was added (under a stream of nitrogen) a solution of ethyl 4-((5-fluoro-2-nitrophenyl)amino)butanoate, **18** (2.5 g, 9.3 mmol) in methanol (30 mL)/ethyl acetate (30 mL). The reaction was stirred under an atmosphere of hydrogen overnight after which it was filtered through celite and the filtrate concentrated to give ethyl 4-((2-amino-5-fluorophenyl)amino)butanoate, **19** (2.3 g, quant.) as a purple oil.$^1$H NMR (500 MHz, Chloroform-*d*) δ 6.59 (dd, *J* = 8.3, 5.6 Hz, 1H), 6.36 – 6.23 (m, 2H), 4.12 (q, *J* = 7.1 Hz, 2H), 3.12 (t, *J* = 6.9 Hz, 2H), 3.07 (s, 2H), 2.44 (t, *J* = 7.1 Hz, 2H), 1.98 (p, *J* = 7.0 Hz, 2H), 1.24 (t, *J* = 7.1 Hz, 4H).
(ES-LCMS) *m*/*z* 241.3 $(M+H)^+$.

Steps 3 / 4: Synthesis of ethyl 4-(7-fluoro-2,3-dioxo-3,4-dihydroquinoxalin-1(2H)-yl)butanoate, **21** (reported in reference 1)

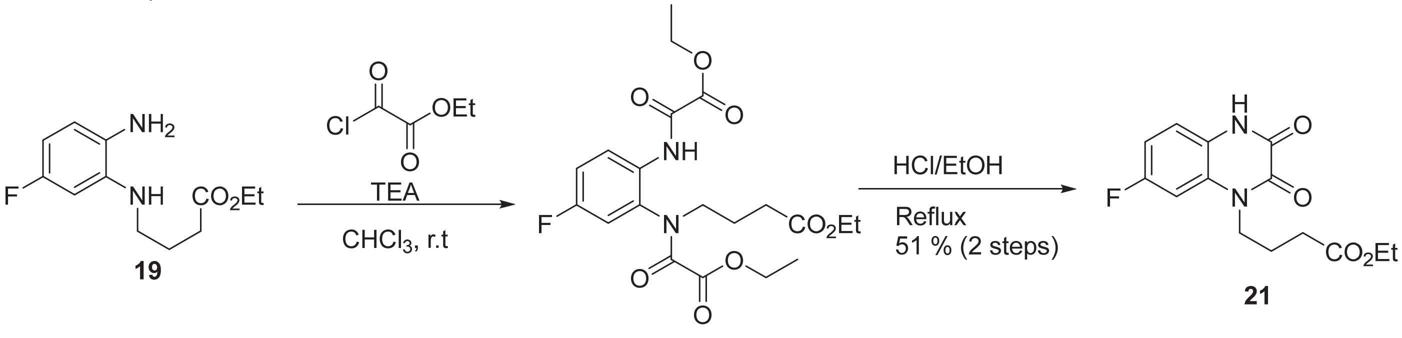


A solution of ethyl 4-((2-amino-5-fluorophenyl)amino)butanoate, **19** (2.3 g, 9.6 mmol) and triethylamine (1.9 g, 2.7 mL, 19 mmol) in chloroform (40 mL) was cooled in an ice/water bath and placed under nitrogen. Ethyl 2-chloro-2-oxoacetate (2.6 g, 2.1 mL, 19 mmol) in chloroform (5 mL) was added dropwise. The ice bath was removed upon dropwise addition and the reaction was allowed to stir at room temperature for 1.5 h after LC/MS indicated the desired intermediate, **20.** The reaction was diluted with chloroform (15 mL) and transferred into a separatory funnel. The organic layer was washed with water (15 mL), saturated aqueous

$NaHCO_3$ solution (2 x 15 mL), saturated $NH_4Cl$ solution (2 x 15 mL), brine (15 mL), and filtered through an isolute phase separator. The filtrate was concentrated to a black/purple oil which was taken up in ethanol (30 mL). Concentrated HCl (0.5 mL) was added and the reaction was heated to 100°C for 1 h after which LC/MS indicated starting material. Additional concentrated HCl (0.5 mL) was added and heating was continued for another 1.5 after the reaction was allowed to cool to room temperature and solids precipitated out of solution. The mixture was filtered and the collected solids washed with ethanol to give ethyl 4-(7-fluoro-2,3-dioxo-3,4-dihydroquinoxalin-1(2H)-yl)butanoate, **21** (1.45 g, 51 %) as a tan solid. $^1$H NMR (500 MHz, DMSO-$d_6$) δ 12.04 (s, 1H), 7.43 (dd, $J$ = 11.1, 2.6 Hz, 1H), 7.19 (dd, $J$ = 8.8, 5.5 Hz, 1H), 7.06 (td, $J$ = 8.5, 2.5 Hz, 1H), 4.17 – 3.98 (m, 4H), 2.47 (t, $J$ = 7.2 Hz, 2H), 1.87 (p, $J$ = 7.3 Hz, 2H), 1.20 (t, $J$ = 7.1 Hz, 3H).
(ES-LCMS) *m/z* 295.2 (M+H)$^+$.

Step 5: Synthesis of ethyl 4-(7-fluoro-6-nitro-2,3-dioxo-3,4-dihydroquinoxalin-1(2H)-yl)butanoate, **22** (reported in reference 1)

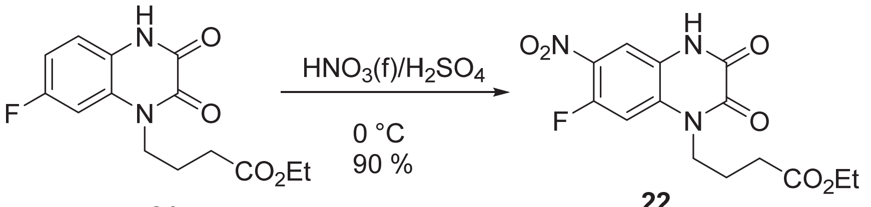


**21** **22**

A round bottom flask containing ethyl 4-(7-fluoro-2,3-dioxo-3,4-dihydroquinoxalin-1(2H)-yl)butanoate, **21** (1.45 g, 4.93 mmol) was immersed in an ice/water bath and concentrated sulfuric acid (15 mL) was added. The mixture was stirred and nitric acid (0.34 g, 0.24 mL, 4.93 mmol) (red, fuming) was added dropwise. The resulting reaction mixture was stirred for 30 min after which it was poured into a mixture of ice and water, and yellow solids precipitated out. The suspension was thoroughly stirred then filtered to give ethyl 4-(7-fluoro-6-nitro-2,3-dioxo-3,4-dihydroquinoxalin-1(2H)-yl)butanoate, **22** (1.51 g, 90.3 %) as a light yellow solid. $^1$H NMR (500 MHz, DMSO-$d_6$) δ 12.25 (s, 1H), 7.92 (d, $J$ = 7.1 Hz, 1H), 7.73 (d, $J$ = 13.4 Hz, 1H), 4.19 – 3.99 (m, 4H), 2.49 (t, $J$ = 7.2 Hz, 2H), 1.87 (p, $J$ = 7.3 Hz, 2H), 1.20 (t, $J$ = 7.1 Hz, 3H). (ES-LCMS) *m/z* 340.2 (M+H)$^+$.

Step 6: Synthesis of ethyl 4-(7-(1H-imidazol-1-yl)-6-nitro-2,3-dioxo-3,4-dihydroquinoxalin-1(2H)-yl)butanoate, **23** (reported in reference 1)

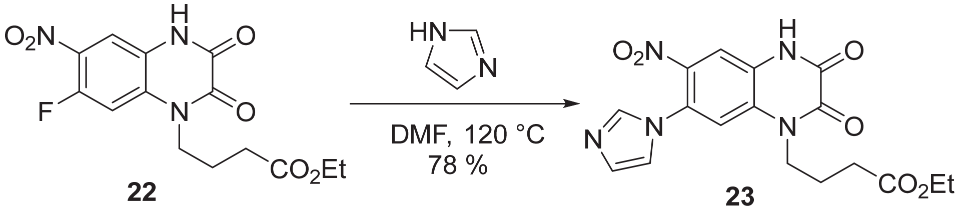


**22** **23**

To a vial containing ethyl 4-(7-fluoro-6-nitro-2,3-dioxo-3,4-dihydroquinoxalin-1(2H)-yl)butanoate, **22** (1.5 g, 4.4 mmol) was added DMF (10 mL) after which imidazole (0.63 g, 9.3 mmol) was added and the reaction was heated overnight at 80°C after which LC/MS still indicated starting material. Additional imidazole (100 mg, 0.7 mmol) was added and the temperature was increased to 100°C for 2 h after which LC/MS still starting material. Additional imidazole (100 mg, 0.7 mmol) was added and heating was continued at 100°C for 2 h then 120°C for 2 h after which the reaction was finally complete. The reaction mixture was poured into a mixture of ice/water and stirred vigorously then filtered. The collected solid was isolated to give ethyl 4-(7-(1H-imidazol-1-yl)-6-nitro-2,3-dioxo-3,4-dihydroquinoxalin-1(2H)-yl)butanoate**, 23** (1.34 g, 78 %) as an orange solid. $^1$H NMR (500 MHz, DMSO-$d_6$) δ 12.57 (s, 1H), 9.45 (s, 1H), 8.17 (s, 1H), 8.05 (d, $J$ = 1.7 Hz, 1H), 8.00 (s, 1H), 7.89 (d, $J$ = 1.7 Hz, 1H), 4.13 (t, $J$ = 7.5 Hz, 2H), 2.40 (t, $J$ = 7.3 Hz, 2H), 1.87 (p, $J$ = 7.4 Hz, 2H). (ES-LCMS) *m/z* 388.3 (M+H)$^+$.

Step 7: 4-(7-(1H-imidazol-1-yl)-6-nitro-2,3-dioxo-3,4-dihydroquinoxalin-1(2H)-yl)butanoic acid, **24** (reported in reference 1)

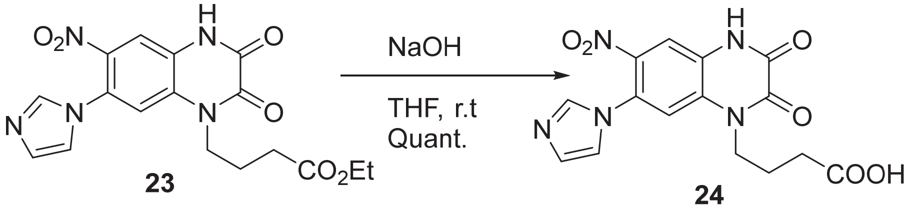


To a vial containing ethyl 4-(7-(1H-imidazol-1-yl)-6-nitro-2,3-dioxo-3,4-dihydroquinoxalin-1(2H)-yl)butanoate, **23** (1 g, 3 mmol) was added NaOH (1 M in water, 8 mL, 4 mmol) and THF (2 mL) and the reaction was stirred at room temperature for 4 h. THF was removed under a stream of nitrogen and to the resulting solution was added 2 M HCl (~ 6 mL) [pH ~ 2] and solids precipitated. The mixture was vigorously stirred then filtered to give 4-(7-(1H-imidazol-1-yl)-6-nitro-2,3-dioxo-3,4-dihydroquinoxalin-1(2H)-yl)butanoic acid, **24** (1.12 g, quant.) as a yellow solid. $^1$H NMR (500 MHz, DMSO-$d_6$) δ 12.57 (s, 1H), 9.45 (s, 1H), 8.17 (s, 1H), 8.05 (d, *J* = 1.7 Hz, 1H), 8.00 (s, 1H), 7.89 (d, *J* = 1.7 Hz, 1H), 4.13 (t, *J* = 7.5 Hz, 2H), 2.40 (t, *J* = 7.3 Hz, 2H), 1.87 (p, *J* = 7.4 Hz, 2H). (ES-LCMS) *m*/*z* 360.3 $(M+H)^+$.

Step 8: Synthesis of 4-(7-(1H-imidazol-1-yl)-6-nitro-2,3-dioxo-3,4-dihydroquinoxalin-1(2H)-yl)-N-(3,6,9,12,15,18-hexaoxahenicos-20-yn-1-yl)butanamide, **25**

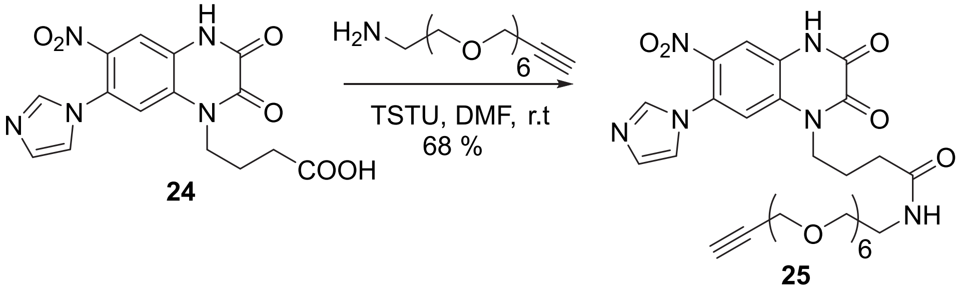


To 4-(7-(1H-imidazol-1-yl)-6-nitro-2,3-dioxo-3,4-dihydroquinoxalin-1(2H)-yl)butanoic acid, **24** (100 mg, 278 µmol) was added DMF (1.5 mL), DIPEA (71.9 mg, 97 µL. 557 µmol) and 1-(bis(dimethylamino)(tetrafluoro-l5-boraneyl)methoxy)pyrrolidine-2,5-dione (TSTU) (83.8 mg, 278 µmol). The reaction was stirred for 15 min after 3,6,9,12,15,18-hexaoxahenicos-20-yn-1-amine (88.9 mg, 278 µmol) was added and the reaction was stirred at room temperature for 1 h after which it was directly purified by reverse phase prep HPLC eluting from 5 to 50 % acetonitrile in water (0.1 % FA and conditions as described in the general chemical synthesis information) to give 4-(7-(1H-imidazol-1-yl)-6-nitro-2,3-dioxo-3,4-dihydroquinoxalin-1(2H)-yl)-N-(3,6,9,12,15,18-hexaoxahenicos-20-yn-1-yl)butanamide, **25** (125 mg, 68 %) as a yellow oil. $^1$H NMR (500 MHz, $CDCl_3$) δ 8.08 (s, 1H), 7.78 (s, 1H), 7.71 (s, 1H), 7.19 (s, 1H), 7.15 (s, 1H), 6.66 (t, *J* = 5.5 Hz, 1H), 4.27 (t, *J* = 7.5 Hz, 2H), 4.13 (d, *J* = 2.4 Hz, 2H), 3.69 – 3.55 (m, 20H), 3.51 (t, *J* = 5.1 Hz, 2H), 3.38 (q, *J* = 5.2 Hz, 2H), 2.49 – 2.26 (m, 3H), 2.11 (q, *J* = 7.0 Hz, 2H). (ES-LCMS) *m*/*z* 661.4 $(M+H)^+$.

Step 9: Synthesis of **YM90K.1$^{DART.1}$ (R5)**

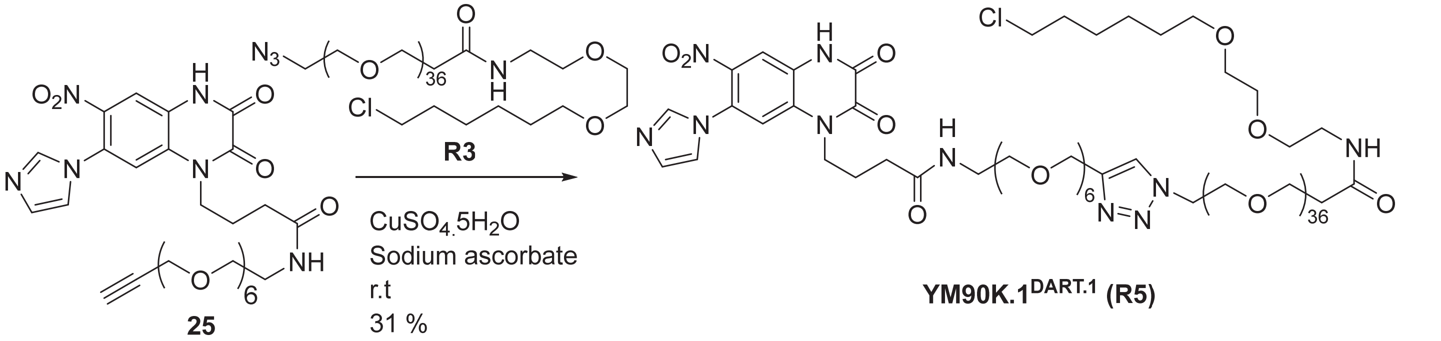

To 4-(7-(1H-imidazol-1-yl)-6-nitro-2,3-dioxo-3,4-dihydroquinoxalin-1(2H)-yl)-N-(3,6,9,12,15,18-hexaoxahenicos-20-yn-1-yl)butanamide, **25** (5 mg, 7.6 µmol) was added EtOH (0.4 mL)/t-BuOH (0.2 mL)/Water (0.2 mL) after which azido-PEG36-HTL1.0, **R3** (14 mg, 7.6 µmol) was added. Finally, Cu(II) sulfate pentahydrate (0.1 mg, 0.38 µmol) and sodium ascorbate (0.14 mg, 0.76 µmol) were added and the reaction mixture was stirred overnight at room temperature. The mixture was directly purified by reverse phase prep HPLC eluting from 5 to 70 % acetonitrile in water (0.1 % FA, 220 nm collection wavelength and other parameters as described in the general chemical synthesis section) and lyophilized to give 1-(4-(24-(7-(1H-imidazol-1-yl)-6-nitro-2,3-dioxo-3,4-dihydroquinoxalin-1(2H)-yl)-21-oxo-2,5,8,11,14,17-hexaoxa-20-azatetracosyl)-1H-1,2,3-triazol-1-yl)-N-(2-(2-((6-chlorohexyl)oxy)ethoxy)ethyl)-3,6,9,12,15,18,21,24,27,30,33,36,39,42,45,48,51,54,57,60,63,66,69,72,75,78,81,84,87,90,93,96,99,102,105,108-hexatriacontaoxaundecahectan-111-amide, **YM90K.1$^{DART.1}$, R5** (6 mg, 31 %).

$^1$H NMR (500 MHz, $CDCl_3$) δ 8.11 (s, 1H), 7.79 – 7.62 (m, 2H), 7.20 (s, 1H), 7.14 (s, 1H), 6.64 (s, 1H), 6.59 (s, 1H), 4.62 (s, 2H), 4.50 (q, *J* = 5.2 Hz, 2H), 4.27 (t, *J* = 7.4 Hz, 1H), 3.84 (t, *J* = 5.1 Hz, 2H), 3.77 – 3.34 (m, 178H), 2.45 (t, *J* = 6.1 Hz, 2H), 2.32 (t, *J* = 6.5 Hz, 1H), 2.11 – 2.00 (m, 2H), 1.75 (p, *J* = 6.9 Hz, 2H), 1.58 (p, *J* = 6.8 Hz, 2H), 1.43 (q, *J* = 8.3 Hz, 2H), 1.38 – 1.29 (m, 2H). HRMS (ESI+): *m/z* calcd for $C_{115}H_{211}ClN_{10}O_{50}$: 1284.1994 $[M+2H]^{2+}$; found: 1284.1989, $[M+2H]^{2+}$

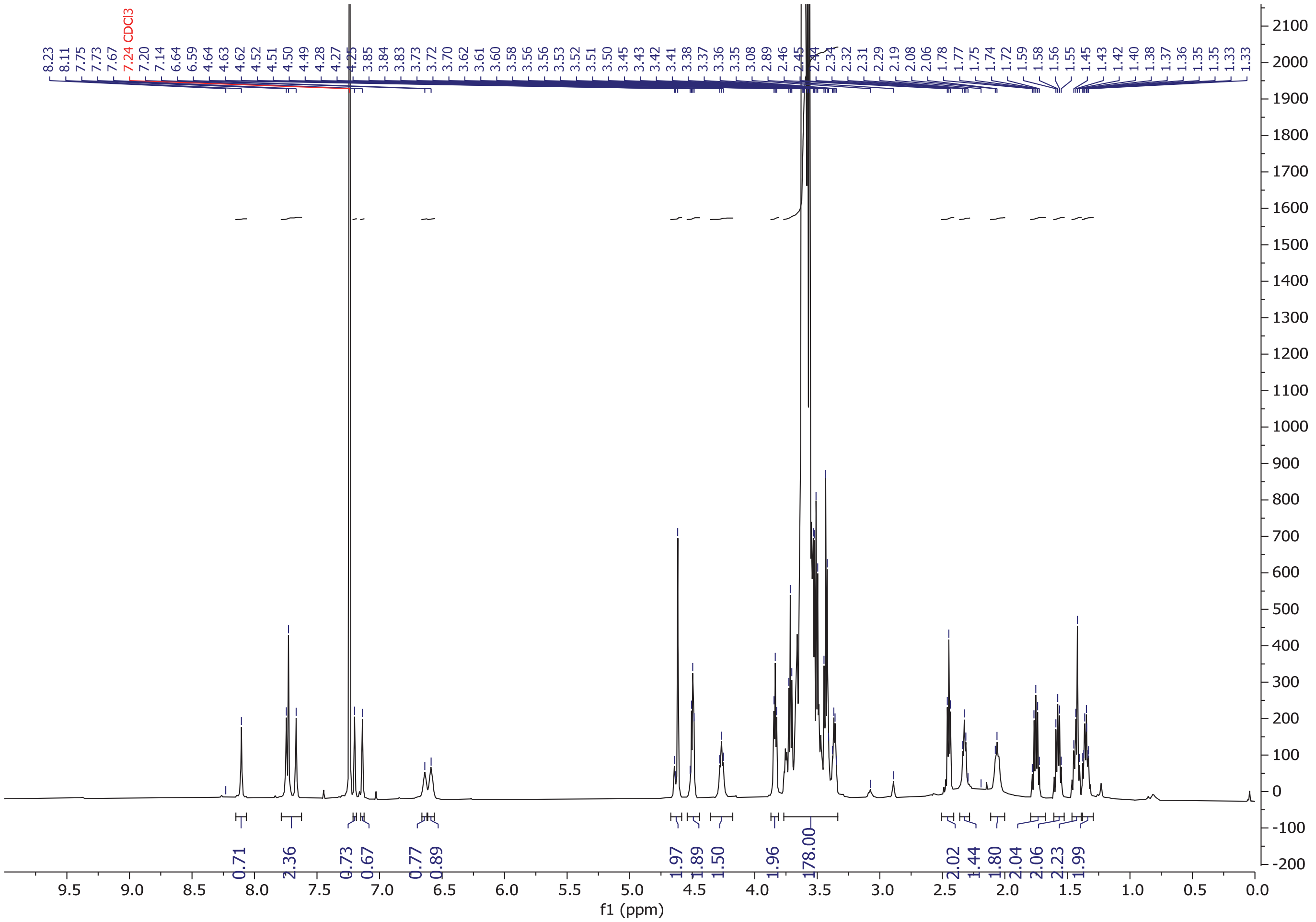

**$^1$H NMR of YM90K.1$^{DART.1}$ (R5) in $CDCl_3$**

#### Synthesis of R6–YM90K.1$^{DART.2}$

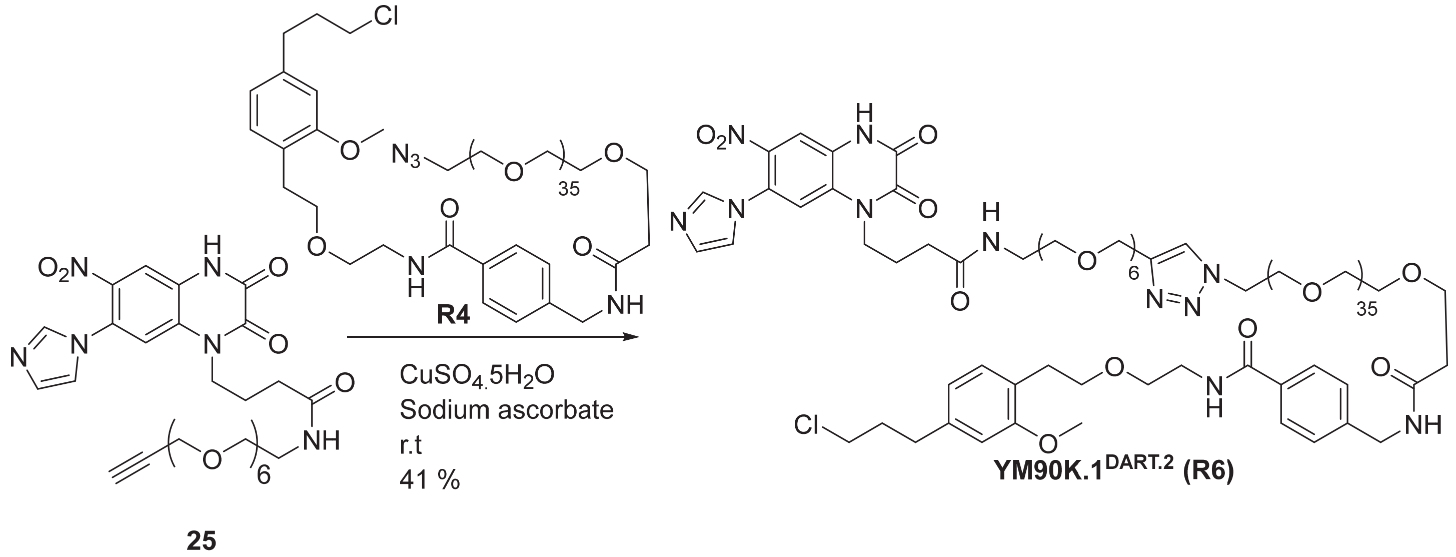

To 4-(7-(1H-imidazol-1-yl)-6-nitro-2,3-dioxo-3,4-dihydroquinoxalin-1(2H)-yl)-N-(3,6,9,12,15,18-hexaoxahenicos-20-yn-1-yl)butanamide, **25** (10 mg, 15 µmol) was added EtOH (0.4 mL)/t-BuOH (0.2 mL)/Water (0.2 mL) after which azido-PEG36-HTL2.0, **R4** (32 mg, 15 µmol) was added. Finally, Cu(II) sulfate pentahydrate (0.19 mg, 0.76 µmol) and sodium ascorbate (0.27 mg, 1.5 µmol) were added and the reaction mixture was stirred overnight at room temperature. The mixture was directly purified by reverse phase prep HPLC eluting from 5 to 70 % acetonitrile in water (0.1 % FA, 220 nm collection wavelength and other parameters as described in the general chemical synthesis section) and lyophilized to give 4-(113-(4-(24-(7-(1H-imidazol-1-yl)-6-nitro-2,3-dioxo-3,4-dihydroquinoxalin-1(2H)-yl)-21-oxo-2,5,8,11,14,17-hexaoxa-20-azatetracosyl)-1H-1,2,3-triazol-1-yl)-3-oxo-6,9,12,15,18,21,24,27,30,33,36,39,42,45,48,51,54,57,60,63,66,69,72,75,78,81,84,87,90,93,96,99,102,105,108,111-hexatriacontaoxa-2-azatridecahectyl)-N-(2-(4-(3-chloropropyl)-2-methoxyphenethoxy)ethyl)benzamide, **YM90K.1$^{DART.2}$, R6** (17 mg, 41 %). $^1$H NMR (500 MHz, $CDCl_3$) δ 8.15-8.12 (m, 2H), 7.90 (br.s, 1H), 7.73 (s, 2H), 7.66 (d, *J* = 8.1 Hz, 2H), 7.32 (d, *J* = 8.0 Hz, 2H), 7.25 – 7.12 (m, 2H), 7.04 (d, *J* = 7.8 Hz, 1H), 6.79 (d, *J* = 7.0 Hz, 1H), 6.65 (d, *J* = 6.3 Hz, 2H), 6.54 (d, *J* = 5.4 Hz, 1H), 4.61 (s, 2H), 4.51 – 4.42 (m, 4H), 4.24 (t, *J*=7.5Hz, 2H), 3.83 (t, *J* = 5.0 Hz, 2H), 3.77 (s, 3H), 3.75 (t, *J* = 5.7 Hz, 3H), 3.68 – 3.47 (m, 163H), 3.36 (q, *J* = 5.2 Hz, 2H), 2.85 (t, *J* = 7.0 Hz, 2H), 2.70 (t, *J* = 7.4 Hz, 2H), 2.53 (t, *J* = 5.6 Hz, 2H), 2.33 (d, *J* = 6.4 Hz, 2H), 2.06 – 2.00 (m, 4H).
HRMS (ESI+): *m/z* calcd for $C_{127}H_{218}ClN_{11}O_{51}$: 1374.7257 $[M+2H]^{2+}$; found: 1374.7217, $[M+2H]^{2+}$

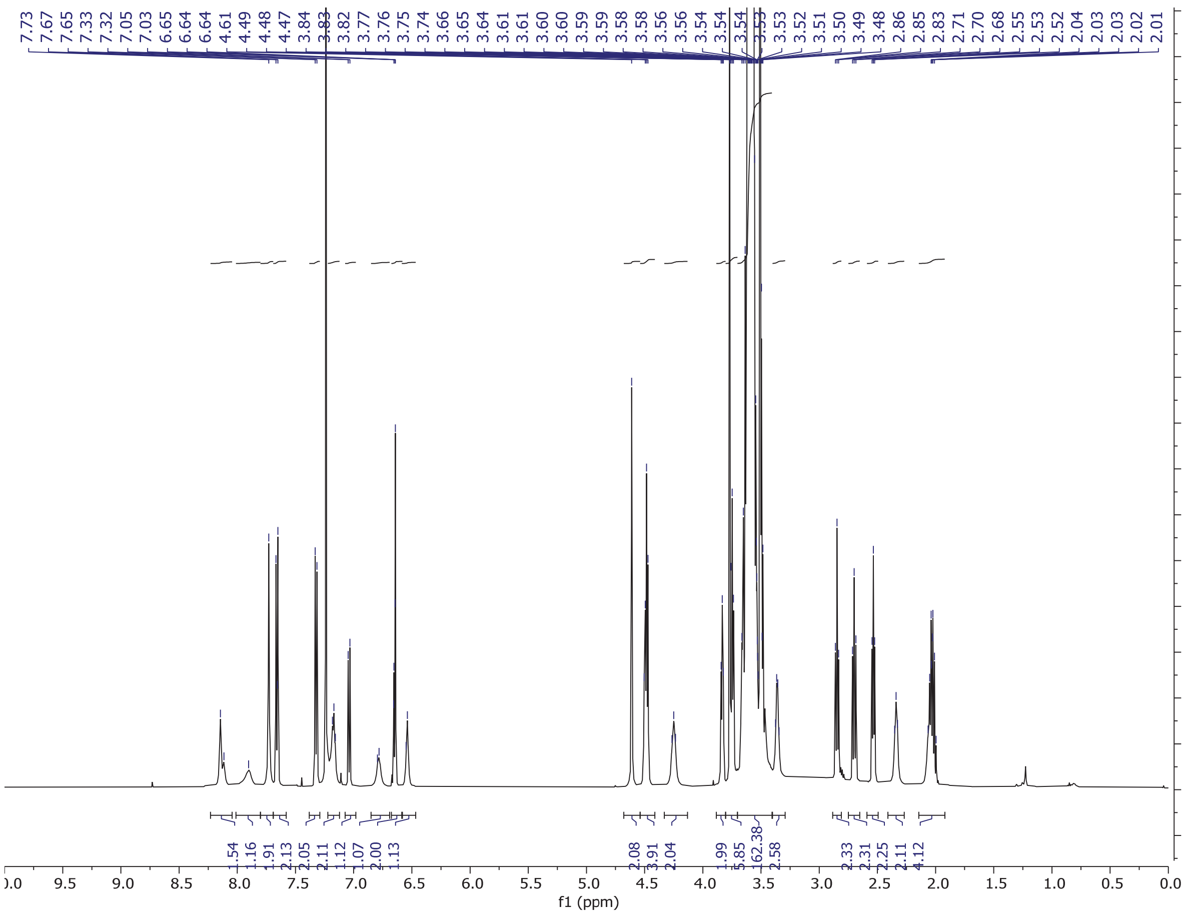

**$^1$H NMR of YM90K.1$^{DART.2}$ (R6) in $CDCl_3$**

### Synthesis of R7–gabazine.1[DART.2]

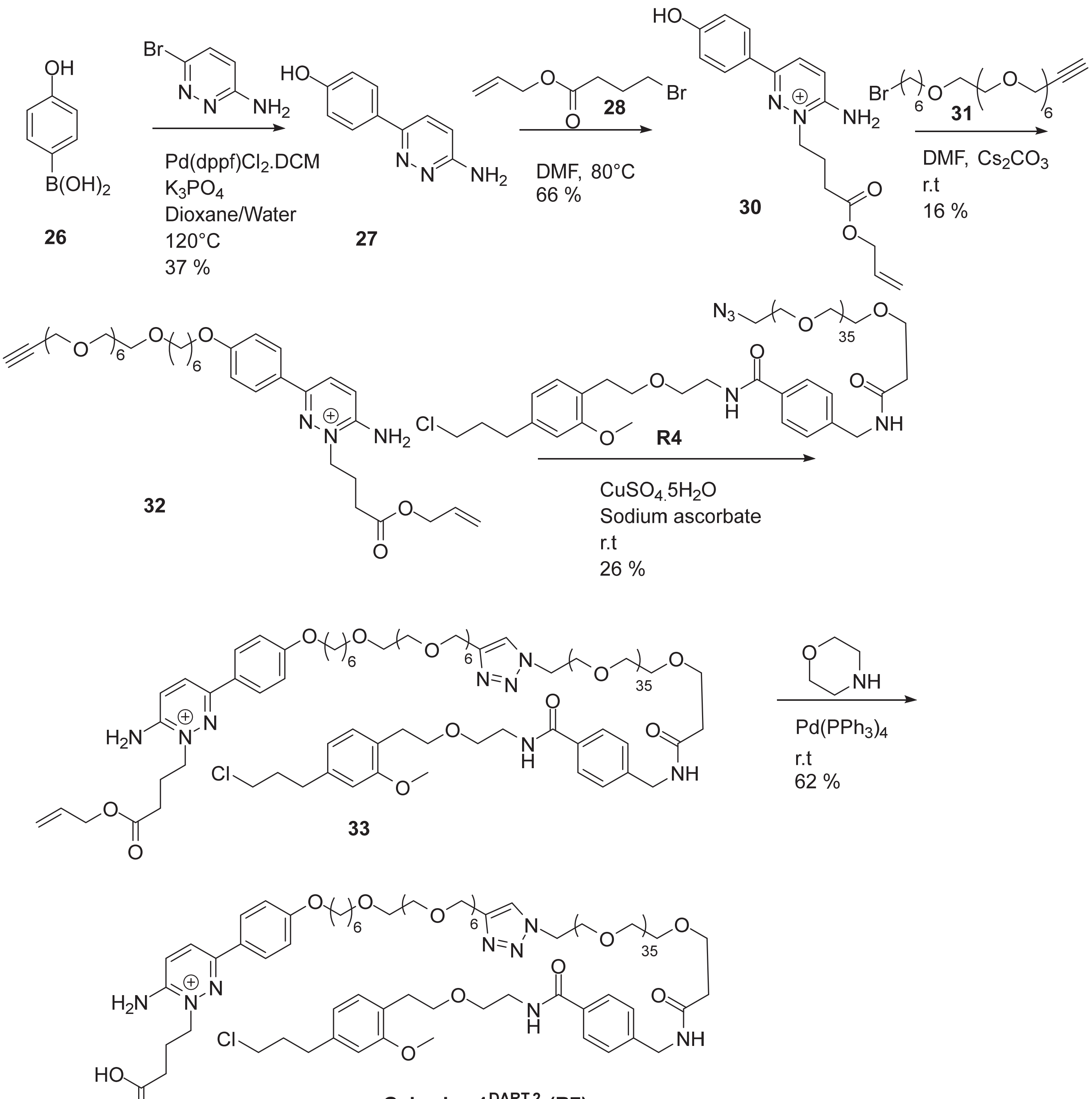


**Gabazine.1[DART.2] (R7)**

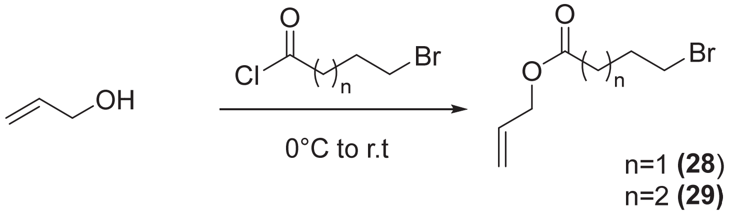


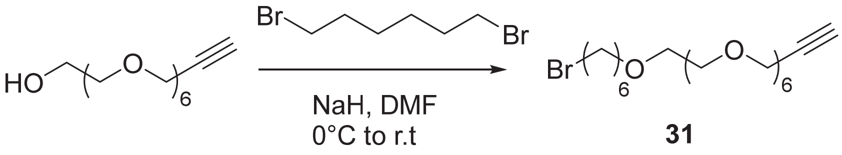


**Synthesis of Intermediates towards Gabazine-DARTs**

<u>Synthesis of allyl 4-bromobutanoate, **28** (Reported in reference 2)</u>

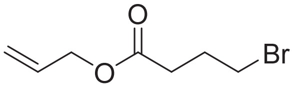

**28**

To cooled (ice/water bath) mixture of allyl alcohol (1.9 g, 2.2 mL, 6 equiv., 32 mmol) under nitrogen was added 4-bromobutanoyl chloride (1.0 g, 0.62 mL, 1 equiv., 5.0 mmol) dropwise. The reaction was allowed to stir at 0 °C for 2 h followed by room temperature for 3 h after TLC (1:1 EtOAc:Hex, iodine/silica stain) indicated conversion. The reaction mixture was directly concentrated to give allyl 4-bromobutanoate, **28** (1.035 g, 4.998 mmol, 93 %) as a brown oil. $^1$H NMR (500 MHz, $CDCl_3$) δ 5.90 (ddt, *J* = 16.7, 11.1, 5.6 Hz, 1H), 5.27 (dd, *J* = 38.1, 13.8 Hz, 2H), 4.59 (d, *J* = 5.8 Hz, 2H), 3.46 (t, *J* = 6.3 Hz, 2H), 2.54 (t, *J* = 7.1 Hz, 2H), 2.18 (p, *J* = 6.7 Hz, 2H).
(ES-LCMS) *m*/*z* 209.2 $(M+2H)^+$.

<u>Synthesis of allyl 5-bromopentanoate, **29** (Reported in reference 2)</u>

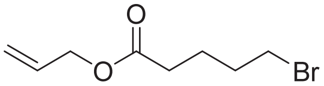

**29**

To a cooled (ice/water bath) mixture of allyl alcohol (1.7 g, 2.0 mL, 6 equiv., 30 mmol) under nitrogen was added 5-bromopentanoyl chloride (1.0 g, 0.62 mL, 1 equiv., 5.0 mmol) dropwise. The reaction was allowed to stir at 0 °C for 2 h followed by room temperature for 3 h after TLC (1:1 EtOAc:Hex, iodine/silica stain) indicated conversion. The reaction mixture was directly concentrated to give allyl 5-bromopentanoate, **29** (0.77 g, 3.48 mmol, 69 %) as a brown oil. $^1$H NMR (500 MHz, $CDCl_3$) δ 5.94-5.86 (m, 1H), 5.30 (d, *J* = 17.1 Hz, 1H), 5.22 (d, *J* = 10.4 Hz, 1H), 4.57 (d, *J* = 5.6 Hz, 2H), 3.40 (t, *J* = 6.5 Hz, 2H), 2.37 (t, *J* = 7.2 Hz, 2H), 1.89 (q, *J* = 7.0 Hz, 2H), 1.79 (q, *J* = 7.3 Hz, 2H).
(ES-LCMS) *m*/*z* 223.1 $(M+2H)^+$.

<u>Synthesis of 28-bromo-4,7,10,13,16,19,22-heptaoxaoctacos-1-yne, **31**</u>

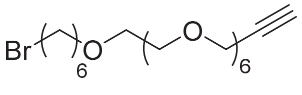

**31**

To 3,6,9,12,15,18-hexaoxahenicos-20-yn-1-ol (1.00 g, 1 equiv., 3.12 mmol) in DMF (12 mL) in an ice/water bath and under nitrogen was added NaH (137 mg, 60% Wt, 1.1 equiv., 3.43 mmol) and the mixture was stirred

for 20 min after which 1,6-dibromohexane (1.22 g, 768 µL, 1.6 equiv., 4.99 mmol) was added quickly. After 2 h of stirring as the ice bath expired, the reaction was quenched with saturated aqueous ammonium chloride solution and diluted with ethyl acetate and water. The aqueous layer was extracted once with ethyl acetate. Combined organics were washed with water, 10 % LiCl (aqueous) solution, brine, filtered through an isolute phase separator, and concentrated to a residue which was purified by silica gel chromatography eluting with 0 to 15 % MeOH in DCM to give 28-bromo-4,7,10,13,16,19,22-heptaoxaoctacos-1-yne, **31** (865 mg, 1.79 mmol, 57.3 %) as a light yellow oil. $^{1}$H NMR (500 MHz, $CDCl_3$) δ 4.18 (d, *J* = 2.4 Hz, 2H), 3.69 – 3.59 (m, 22H), 3.55 (dd, *J* = 5.9, 3.8 Hz, 2H), 3.43 (t, *J* = 6.6 Hz, 2H), 3.38 (t, *J* = 6.8 Hz, 2H), 2.41 (t, *J* = 2.4 Hz, 1H), 1.86-1.80 (m, 2H), 1.60-1.54 (m, 2H), 1.46 – 1.39 (m, 2H), 1.37 – 1.30 (m, 2H).
(ES-LCMS) *m*/*z* 485.3 $(M+2H)^{+}$.

Step 1: Synthesis of 4-(6-aminopyridazin-3-yl)phenol, **27**

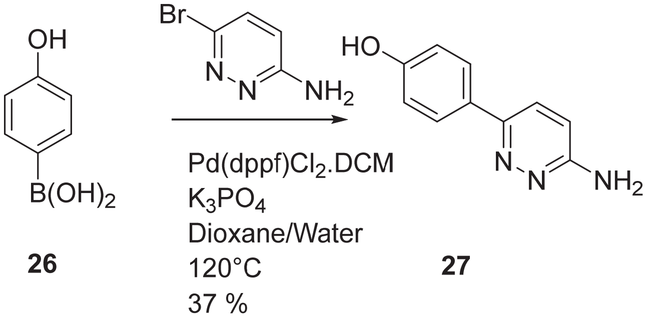


To (4-hydroxyphenyl)boronic acid, **26** (2 g, 14.5 mmol) was added $K_3PO_4$ (6.16 g, 29 mmol), Pd(dppf)Cl2·DCM (0.71 g, 0.87 mmol), 6-bromopyridazin-3-amine (2.47 g, 14.2 mmol) and dioxane (44 mL)/water (28 mL), and the reaction was heated in the microwave at 120°C for 10 min (note: separated into 4 portions which were each heated separately at 120°C for 10 min). The combined reaction mixture was diluted with ethyl acetate (50 mL), filtered through a pad of silica, and the contents transferred to a separatory funnel. The organic layer was washed with water (20 mL), brine (20 mL), and concentrated to a black solid which was triturated from dichloromethane. This solid was further purified by silica gel chromatography eluting with 0 to 35 % methanol in dichloromethane (product eluted in a wide band) to give 4-(6-aminopyridazin-3-yl)phenol, **27** (1 g, 5.34 mmol, 36.8 % yield) as a dark, brown solid. $^{1}$H NMR (500 MHz, $CD_3OD$) δ 7.73 (dd, *J* = 9.0, 2.5 Hz, 3H), 7.00 (d, *J* = 9.3 Hz, 1H), 6.92 – 6.84 (m, 2H).(ES-LCMS) *m*/*z* 188.1 $(M+H)^{+}$.

Step 2: Synthesis of allyl 4-(3-(4-hydroxyphenyl)-6-iminopyridazin-1(6H)-yl)butanoate, **30** (see references 3 and 4)

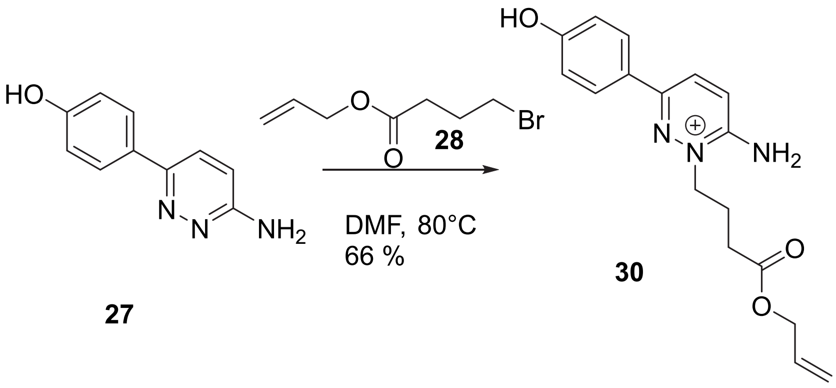


To 4-(6-aminopyridazin-3-yl)phenol, **27** (1 g, 5.34 mmol) was added allyl 4-bromobutanoate, **28** (1.14 g, 5.51 mmol) and DMF (26.7 mL) and the reaction was heated at 80°C for 7 h after which it was concentrated to remove DMF. The resulting thick, black oil was then triturated from acetonitrile give allyl 4-(3-(4-hydroxyphenyl)-6-iminopyridazin-1(6H)-yl)butanoate, **30** (1.1 g, 3.51 mmol, 65.7 % yield) as a brown solid. Since there are three nitrogens and a phenolic oxygen which may all react with the alkyl bromide, the assignment and confirmation of this product was described in the synthesis of **Gabazine.7** **DART.2** **(R9).** $^{1}$H NMR (500 MHz, Methanol-$d_4$) δ 8.23 (d, *J* = 9.5 Hz, 1H), 7.84 (d, *J* = 8.6 Hz, 2H), 7.55 (d, *J* = 9.5 Hz, 1H), 6.91 (dd,

$J$ = 9.1, 2.7 Hz, 2H), 5.86 (ddt, $J$ = 16.5, 11.0, 5.7 Hz, 1H), 5.33 – 5.13 (m, 2H), 4.50 (d, $J$ = 5.7 Hz, 2H), 4.43 (t, $J$ = 6.9 Hz, 2H), 2.63 (t, $J$ = 6.8 Hz, 2H), 2.27 (p, $J$ = 6.8 Hz, 2H). $^{13}$C NMR (125 MHz, $CD_3OD$) δ 172.65, 161.38, 152.44, 150.54, 132.00, 130.83, 127.91, 125.25, 123.22, 117.22, 115.99, 65.08, 55.19, 29.94, 21.12. (ES-LCMS) m/z 314.3 $(M+H)^+$.

Step 3: Synthesis of allyl 4-(3-(4-(4,7,10,13,16,19,22-heptaoxaoctacos-1-yn-28-yloxy)phenyl)-6-iminopyridazin-1(6H)-yl)butanoate, **32**

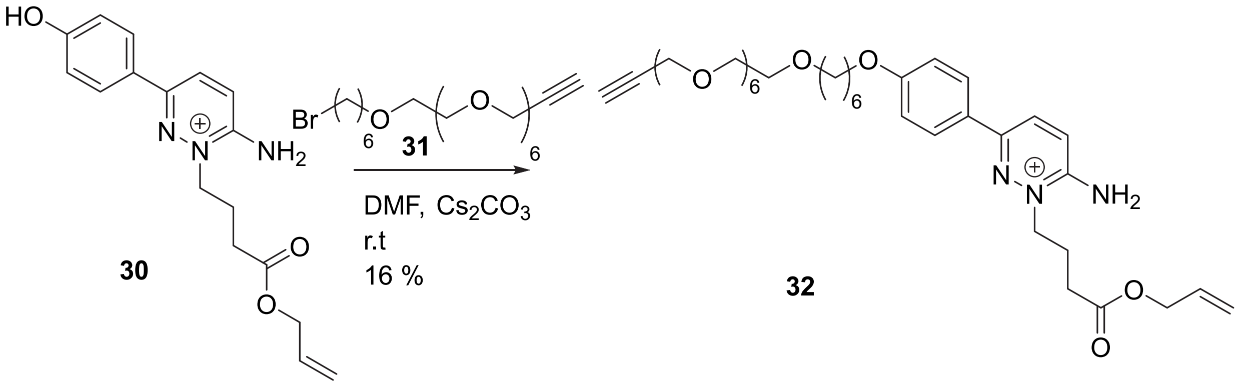


To allyl 4-(3-(4-hydroxyphenyl)-6-iminopyridazin-1(6H)-yl)butanoate, **30** (1.1 g, 3.51 mmol) was added anhydrous DMF (35.1 mL) after which 28-bromo-4,7,10,13,16,19,22-heptaoxaoctacos-1-yne, **31** (2 g, 4.13 mmol) was added. The solution was cooled to 0°C and anhydrous cesium carbonate (3.43 g, 10.53 mmol) was added and the reaction was stirred for 3 h as the ice bath expired after which water (20 mL) and ethyl acetate (60 mL) were added. The organic layer was washed with water (10 mL), brine (2 x 10 mL), dried (sodium sulfate), filtered and concentrated to give crude material as a yellow oil which was purified by reverse phase prep HPLC in several injections eluting with 5 to 50 % acetonitrile in water (0.1 % formic acid conditions and other conditions as described in the general chemical synthesis section) to give allyl 4-(3-(4-(4,7,10,13,16,19,22-heptaoxaoctacos-1-yn-28-yloxy)phenyl)-6-iminopyridazin-1(6H)-yl)butanoate, **32** (412 mg, 0.576 mmol, 16.39 % yield) as a red/yellow oil. $^1$H NMR (500 MHz, $CD_3OD$) δ 8.39 – 8.24 (m, 3H), 8.03 – 7.85 (m, 2H), 7.61 (d, $J$ = 9.6 Hz, 1H), 7.18 – 6.99 (m, 2H), 5.86 (ddt, $J$ = 17.3, 10.5, 5.7 Hz, 1H), 5.34 – 5.11 (m, 2H), 4.57 – 4.38 (m, 4H), 4.25 – 4.02 (m, 4H), 3.75 – 3.44 (m, 24H), 2.87 (t, $J$ = 2.4 Hz, 1H), 2.64 (t, $J$ = 6.7 Hz, 2H), 2.29 (p, $J$ = 6.8 Hz, 2H), 1.92 – 1.78 (m, 2H), 1.71 – 1.42 (m, 6H).
(ES-LCMS) *m/z* 716.5 $(M+H)^+$.

Step 4: Synthesis of allyl 4-(3-(4-((1-(1-(1-(4-((2-(4-(3-chloropropyl)-2-methoxyphenethoxy)ethyl)carbamoyl)phenyl)-3-oxo-6,9,12,15,18,21,24,27,30,33,36,39,42,45,48,51,54,57,60,63,66,69,72,75,78,81,84,87,90,93,96,99,102,105,108,111-hexatriacontaoxa-2-azatridecahectan-113-yl)-1H-1,2,3-triazol-4-yl)-2,5,8,11,14,17,20-heptaoxahexacosan-26-yl)oxy)phenyl)-6-iminopyridazin-1(6H)-yl)butanoate, **33**

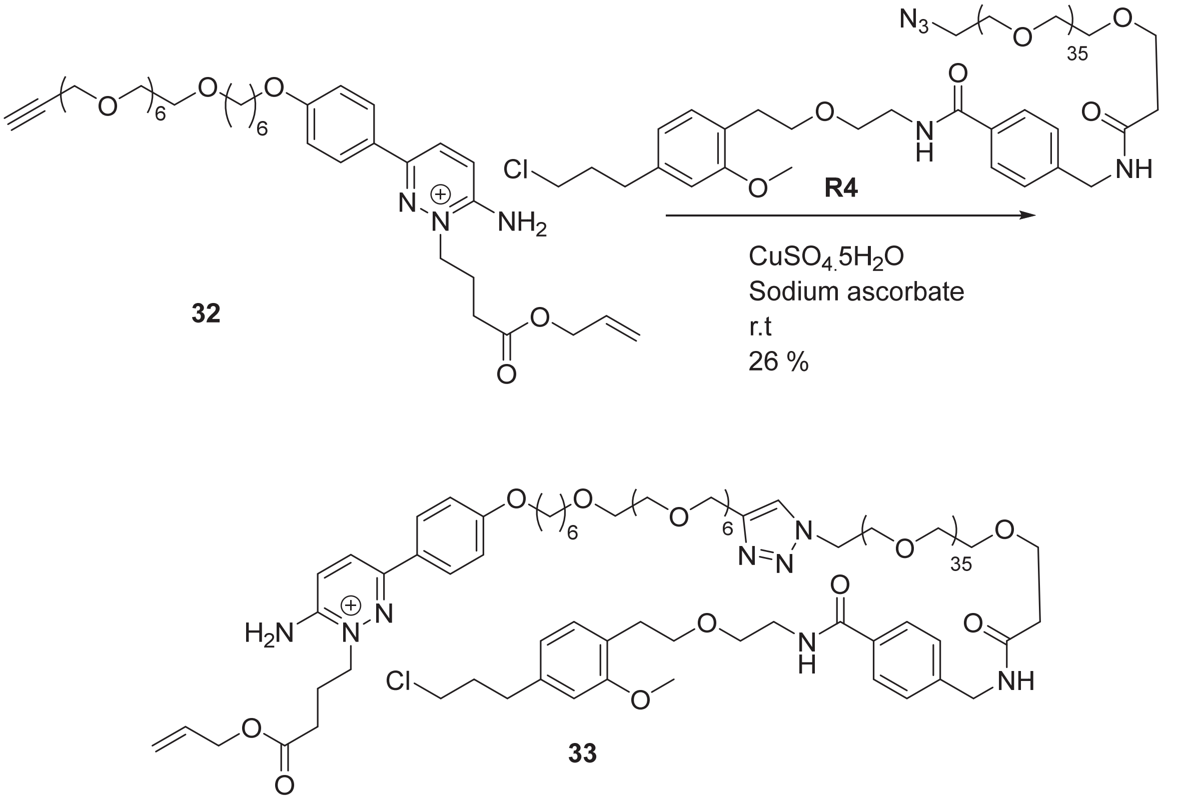


To a vial was added allyl 4-(3-(4-((4,7,10,13,16,19,22-heptaoxaoctacos-1-yn-28-yl)oxy)phenyl)-6-iminopyridazin-1(6H)-yl)butanoate, **32** (40 mg, 56 µmol) and EtOH/t-BuOH/water (2:1:1, 1.6 mL) after which azido-PEG36-HTL2.0, **R4** (0.12 g, 56 µmol) was added. Finally, Cu(II) sulfate pentahydrate (0.7 mg, 2.8 µmol) and sodium ascorbate (0.98 mg, 5.6 µmol) were added and the reaction mixture was stirred at room temperature for 3 h after which it was purified by reverse phase prep HPLC eluting with 50 to 80 % acetonitrile in water (0.1 % formic acid, 220 nm wavelength for collection, and other conditions reported in the general chemical synthesis section) to give allyl 4-(3-(4-((1-(1-(1-(4-((2-(4-(3-chloropropyl)-2-methoxyphenethoxy)ethyl)carbamoyl)phenyl)-3-oxo-6,9,12,15,18,21,24,27,30,33,36,39,42,45,48,51,54,57,60,63,66,69,72,75,78,81,84,87,90,93,96,99,102,105,108,111-hexatriacontaoxa-2-azatridecahectan-113-yl)-1H-1,2,3-triazol-4-yl)-2,5,8,11,14,17,20-heptaoxahexacosan-26-yl)oxy)phenyl)-6-iminopyridazin-1(6H)-yl)butanoate, **33** (45 mg, 26 %) as a yellow oil. $^1$H NMR (500 MHz, $CDCl_3$) δ 8.44 (s, 2H), 8.17 (d, *J* = 9.5 Hz, 1H), 7.83 (d, *J* = 9.5 Hz, 1H), 7.76 – 7.65 (m, 5H), 7.33 (d, *J* = 7.9 Hz, 2H), 7.05 (d, *J* = 7.8 Hz, 1H), 7.02 – 6.94 (m, 2H), 6.66 (d, *J* = 6.5 Hz, 2H), 6.53 (s, 1H), 5.91 – 5.77 (m, 1H), 5.42 – 5.11 (m, 2H), 4.65 (s, 2H), 4.58 (d, *J* = 5.9 Hz, 2H), 4.51 (t, *J* = 5.2 Hz, 2H), 4.48 (d, *J* = 6.0 Hz, 2H), 4.42 (t, *J* = 7.8 Hz, 2H), 3.99 (d, *J* = 6.4 Hz, 1H), 3.85 (t, *J* = 5.1 Hz, 2H), 3.78 (s, 3H), 3.75 (t, *J* = 5.6 Hz, 2H), 3.69 – 3.42 (m, 172H), 2.86 (t, *J* = 7.0 Hz, 2H), 2.71 (t, *J* = 7.4 Hz, 2H), 2.54 (dt, *J* = 15.9, 6.0 Hz, 4H), 2.23 (p, *J* = 6.8 Hz, 2H), 2.03 (dq, *J* = 8.3, 6.5 Hz, 3H), 1.79 (q, *J* = 6.8 Hz, 2H), 1.60 (p, *J* = 6.9 Hz, 2H), 1.53 – 1.30 (m, 3H). (ES-LCMS) *m*/*z* 571.2 $[(M+5H)]^{5+}$.

Step 5: Synthesis of **Gabazine.1$^{DART.2}$ (R7)**

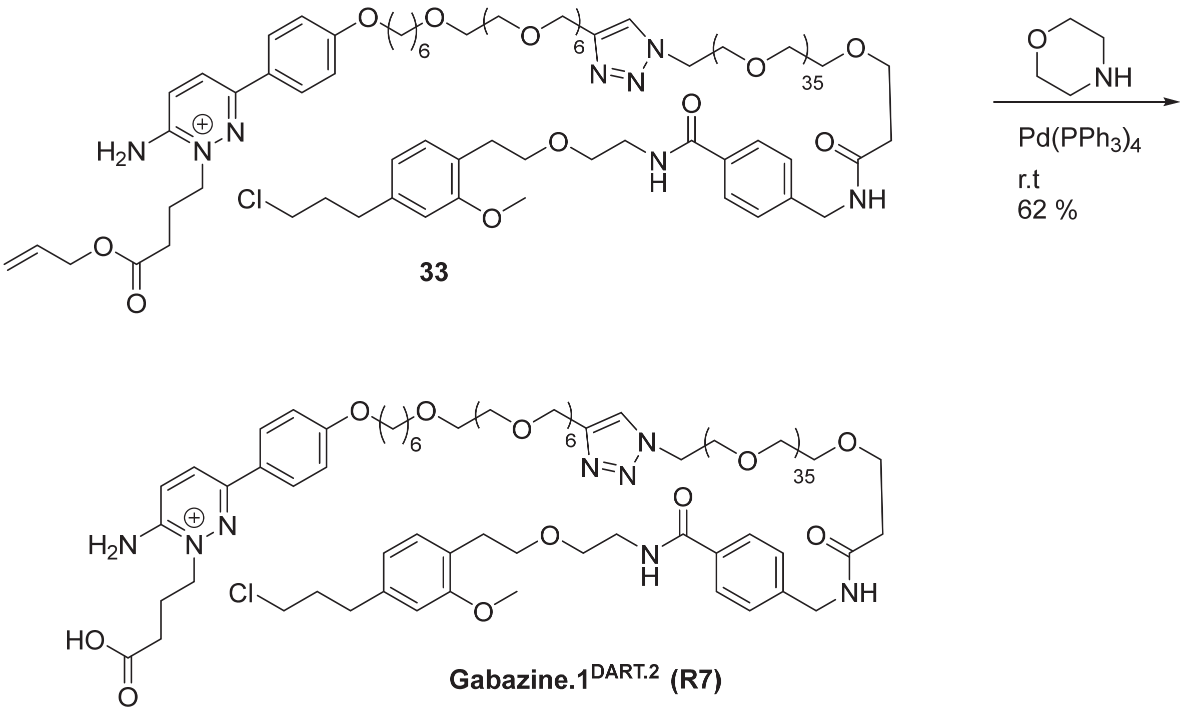


**33**

**Gabazine.1$^{DART.2}$ (R7)**

To a solution of allyl 4-(3-(4-((1-(1-(1-(4-((2-(4-(3-chloropropyl)-2-methoxyphenethoxy)ethyl)carbamoyl)phenyl)-3-oxo-6,9,12,15,18,21,24,27,30,33,36,39,42,45,48,51,54,57,60,63,66,69,72,75,78,81,84,87,90,93,96,99,102,105,108,111-hexatriacontaoxa-2-azatridecahectan-113-yl)-1H-1,2,3-triazol-4-yl)-2,5,8,11,14,17,20-heptaoxahexacosan-26-yl)oxy)phenyl)-6-iminopyridazin-1(6H)-yl)butanoate, **33** (52 mg, 19 µmol) in THF (0.8 mL) was added morpholine (8.1 mg, 8.1µL, 93 µmol) and $Pd(PPh_3)_4$ (3 mg, 11.1 µmol). The reaction was stirred under nitrogen for 1 h after which it was purified by reverse phase prep HPLC eluting with 5 to 80 % acetonitrile in water (0.1 % formic acid, 220 nm wavelength for collection, and conditions reported in the general chemical synthesis section) to give 4-(3-(4-((1-(1-(1-(4-((2-(4-(3-chloropropyl)-2-methoxyphenethoxy)ethyl)carbamoyl)phenyl)-3-oxo-6,9,12,15,18,21,24,27,30,33,36,39,42,45,48,51,54,57,60,63,66,69,72,75,78,81,84,87,90,93,96,99,102,105,108,111-hexatriacontaoxa-2-azatridecahectan-113-yl)-1H-1,2,3-triazol-4-yl)-2,5,8,11,14,17,20-heptaoxahexacosan-26-yl)oxy)phenyl)-6-iminopyridazin-1(6H)-yl)butanoic acid **Gabazine.1$^{DART.2}$, R7** (32 mg, 62 %). $^1$H NMR (500 MHz, $CDCl_3$) δ 10.97 (br.s, 1H), 8.55 (d, *J* = 9.7 Hz, 1H), 7.84 (d, *J* = 9.6 Hz, 1H), 7.77 – 7.72 (m, 2H), 7.71 (s, 1H), 7.69 – 7.65 (m, 2H), 7.33 (d, *J* = 8.0 Hz, 2H), 7.09 (t, *J* = 5.9 Hz, 1H), 7.05 (d, *J* = 7.9 Hz, 1H), 6.99 – 6.95 (m, 2H), 6.66-6.62 (m, 2H), 6.51 (s, 1H), 4.65 (s, 2H), 4.51(t, *J*=5.5 Hz, 2H), 4.48(d, *J* = 6.0Hz , 2H), 4.40 (t, *J*=7.5Hz, 2H), 3.99 (t, *J* = 6.5 Hz, 2H), 3.85 (t, *J* = 5.2 Hz, 2H), 3.77 (s, 3H), 3.75 (t, *J* = 5.6 Hz, 2H), 3.70 – 3.37 (m, 175H), 3.52-3.44 (m, 9H), 2.85 (t, *J* = 7.0 Hz, 2H), 2.71 (t, *J* = 7.4 Hz, 2H), 2.52 (t, *J* = 5.6 Hz, 2H), 2.50 – 2.41 (m, 2H), 2.12-2.07(m, 2H), 2.06-2.00 (m, 2H), 1.80 (p, *J* = 6.8 Hz, 2H), 1.65 – 1.54 (m, 2H), 1.51-1.37 (m, 4H), HRMS (ESI$^+$): *m/z* calcd for $C_{132}H_{229}ClN_8Na_2O_{50}$: 1404.7582 $[M+2Na]^{2+}$; found: 1404.7709 $[M+2Na]^{2+}$

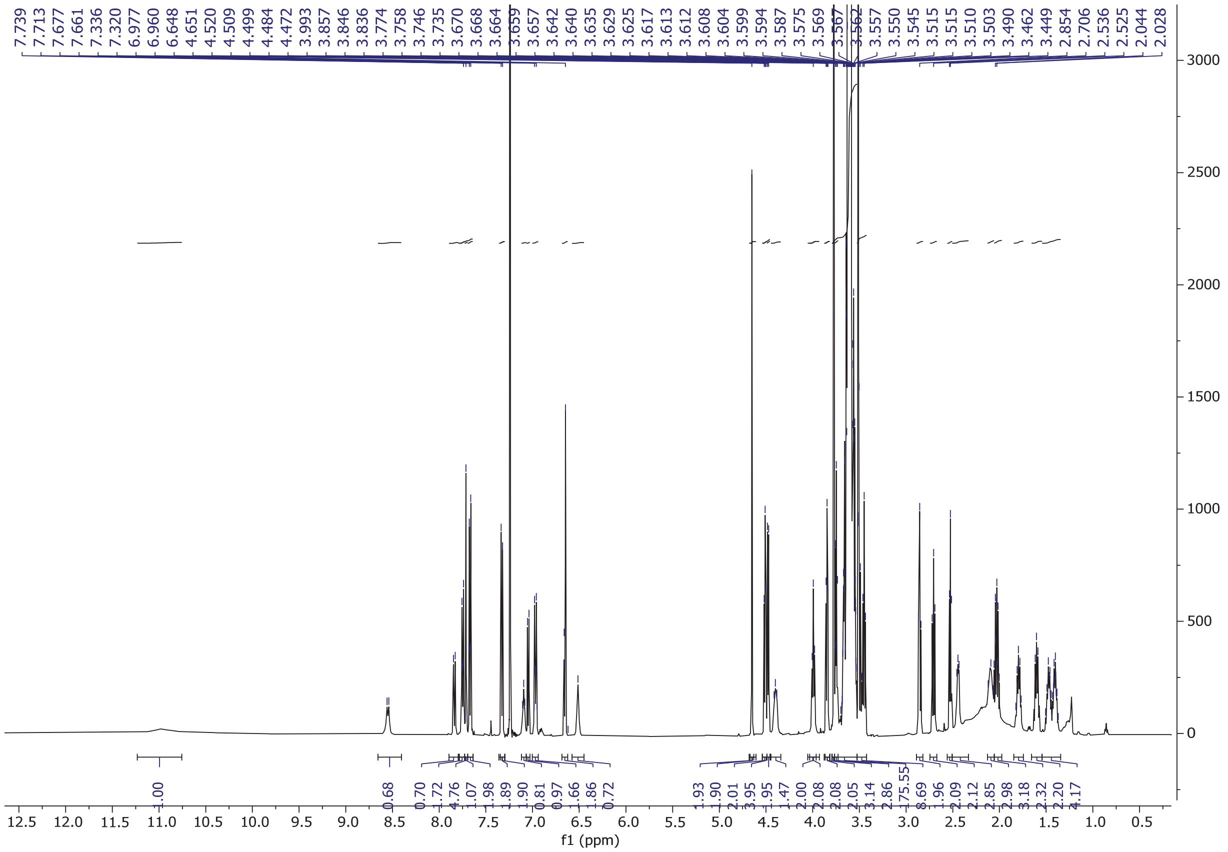

**$^{1}H$ NMR of Gabazine.1$^{DART.2}$ (R7) in $CDCl_3$**

### Synthesis of R8–gabazine.5[DART.2]

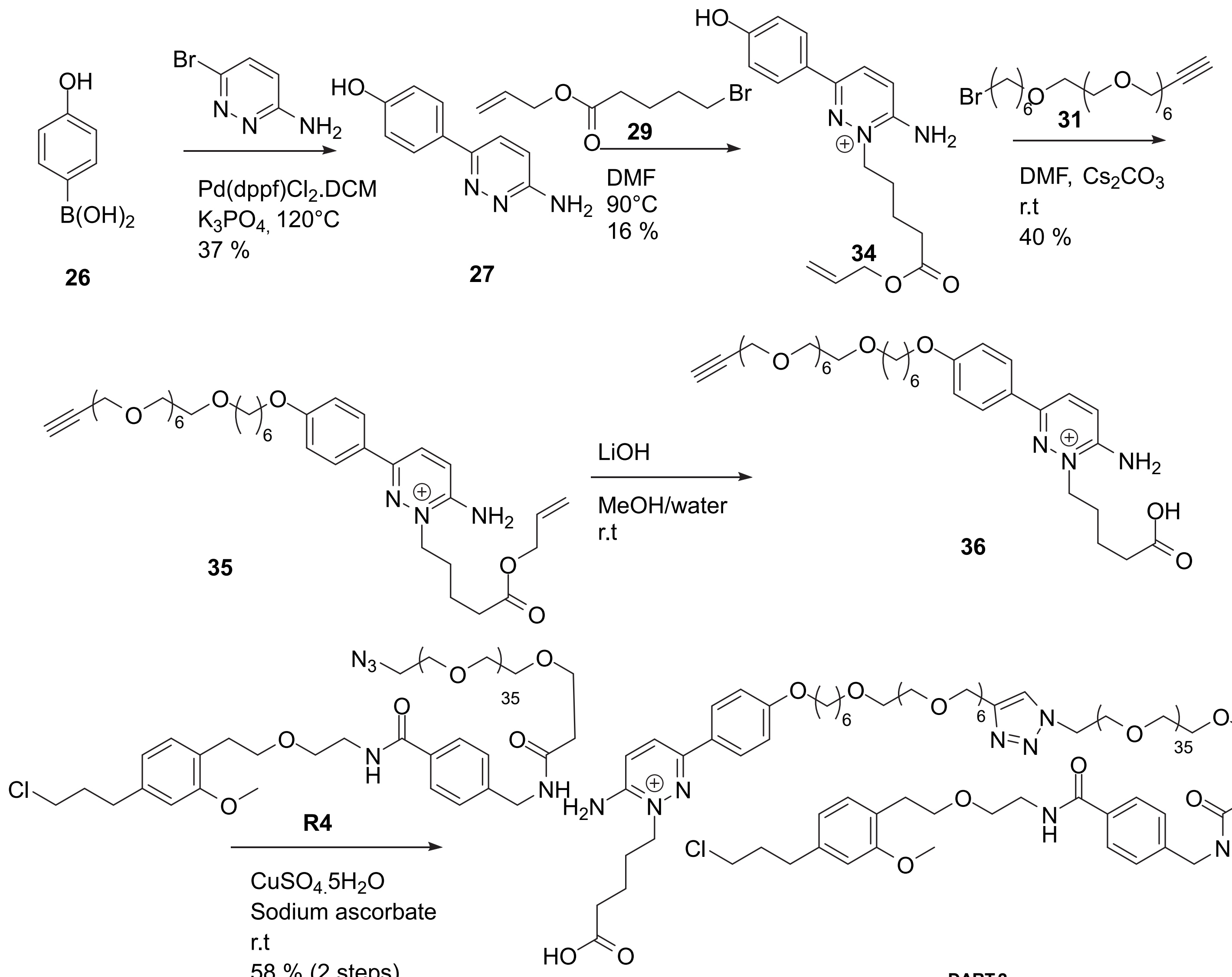


**Gabazine.5[DART.2] (R8)**

Step 1: Synthesis of 4-(6-aminopyridazin-3-yl)phenol, **27** (details in gabazine.1[DART.2])

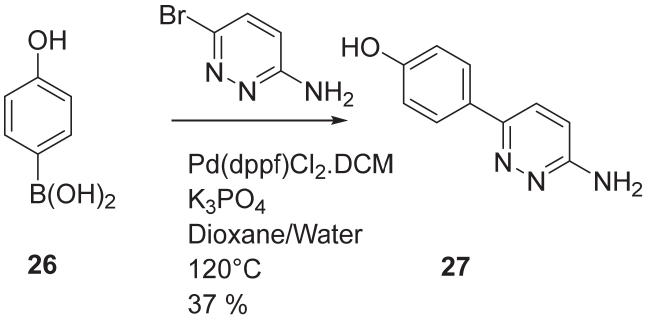

Step 2: Synthesis of allyl 5-(3-(4-hydroxyphenyl)-6-iminopyridazin-1(6H)-yl)pentanoate, **34** (see references 3 and 4)

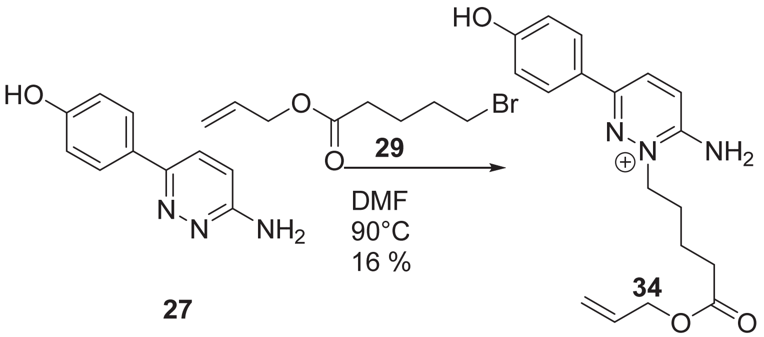


To 4-(6-aminopyridazin-3-yl)phenol, **27** (250 mg, 1.34 mmol) was added allyl allyl 5-bromopentanoate, **29** (295 mg, 1.34 mmol) and DMF (2 mL) and the reaction was heated at 90°C for 2 h. The mixture was directly purified by prep HPLC eluting with 10 to 90 % ACN in water (0.1 % FA, 220 nm wavelength for collection) in three 750 uL injections. The main peak(s) were isolated, and fractions were concentrated to give the desired product, **34** (71 mg, 16 %). Before this main peak, one byproduct with the same mass was isolated. Since three nitrogens in the product as well as the phenolic oxygen all may react with the alkyl bromide, the assignment and confirmation of product was described in the **Gabazine.7 DART.2 (R9)** section. $^1$H NMR (500 MHz, $CD_3OD$) δ 8.47 (br.s, 1H), 8.27 (d, *J* = 9.3 Hz, 1H), 7.87 (d, *J* = 8.4 Hz, 2H), 7.58 (d, *J* = 9.4 Hz, 1H), 6.94 (d, *J* = 8.4 Hz, 2H), 5.97-5.85 (m, 1H), 5.31 (dd, *J* = 17.3, 1.9 Hz, 1H), 5.21 (d, *J* = 10.5 Hz, 1H), 4.59 (d, *J* = 5.6 Hz, 2H), 4.40 (t, *J* = 7.2 Hz, 2H), 2.51 (t, *J* = 7.3 Hz, 2H), 2.05-1.99 (m, 2H), 1.83-1.77 (m, 2H).
$^{13}$C NMR (126 MHz, $CD_3OD$) δ 173.05, 160.45, 152.26, 150.88, 132.27, 131.05, 128.00, 125.02, 123.82, 116.95, 115.69, 64.77, 55.54, 32.75, 25.86, 21.21. (ES-LCMS) *m/z* 328.3 $(M+H)^+$.

Step 3: Synthesis of allyl 5-(3-(4-((4,7,10,13,16,19,22-heptaoxaoctacos-1-yn-28-yl)oxy)phenyl)-6-iminopyridazin-1(6H)-yl)pentanoate, **35**

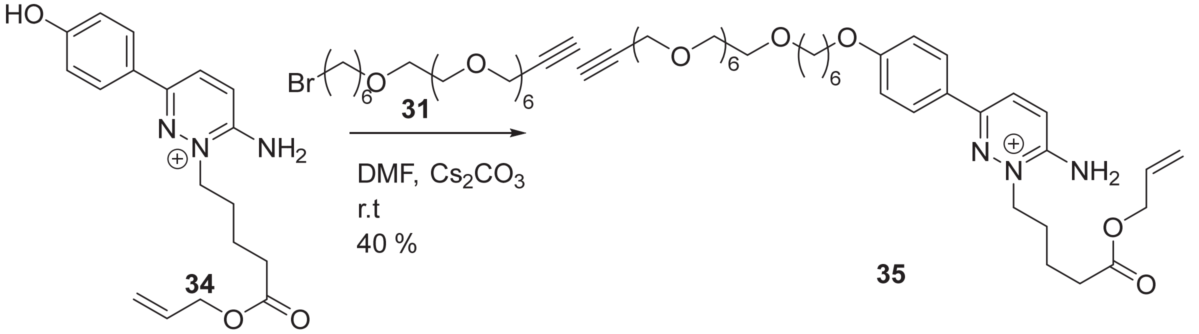


To allyl 5-(3-(4-hydroxyphenyl)-6-iminopyridazin-1(6H)-yl)pentanoate, **34** (40 mg, 0.12 mmol) was added anhydrous DMF (1 mL) after which 28-bromo-4,7,10,13,16,19,22-heptaoxaoctacos-1-yne, **31** (89 mg, 0.18 mmol) and anhydrous cesium carbonate (80 mg, 0.24 mmol) were added and the reaction was stirred for 3h. DMF was removed under a stream of nitrogen, and the crude was purified by reverse phase prep HPLC eluting with 5 to 50 % acetonitrile in water (0.1 % formic acid conditions and other conditions as described in the general chemical synthesis section) to give allyl 5-(3-(4-((4,7,10,13,16,19,22-heptaoxaoctacos-1-yn-28-yl)oxy)phenyl)-6-iminopyridazin-1(6H)-yl)pentanoate, **35** (36 mg, 49 µmol, 40 % yield) as a red/yellow oil. $^1$H NMR (500 MHz, $CD_3OD$) δ 8.53 (s, 1H), 8.30 (d, *J* = 9.6 Hz, 1H), 8.02 – 7.90 (m, 2H), 7.60 (d, *J* = 9.5 Hz, 1H), 7.16 – 7.03 (m, 2H), 5.98 – 5.90 (m, 1H), 5.31 (dq, *J* = 17.2, 1.6 Hz, 1H), 5.21 (dq, *J* = 10.5, 1.4 Hz, 1H), 4.59 (dt, *J* = 5.7, 1.5 Hz, 2H), 4.41 (t, *J* = 7.2 Hz, 2H), 4.20 (d, *J* = 2.4 Hz, 2H), 4.09 (t, *J* = 6.4 Hz, 2H), 3.69 – 3.63 (m, 21H), 3.61 – 3.59 (m, 2H), 3.52 (t, *J* = 6.5 Hz, 2H), 2.86 (t, *J* = 2.4 Hz, 1H), 2.51 (t, *J* = 7.3 Hz, 2H), 2.06 – 2.00 (m, 2H), 1.87-1.77 (m, 4H), 1.67-1.61 (m, 2H), 1.58-1.52 (m, 2H), 1.51 – 1.45 (m, 2H). (ES-LCMS) *m/z* 730.6 $(M+H)^+$.

Steps 4/5: Synthesis of **Gabazine.5$^{DART.2}$ (R8)**

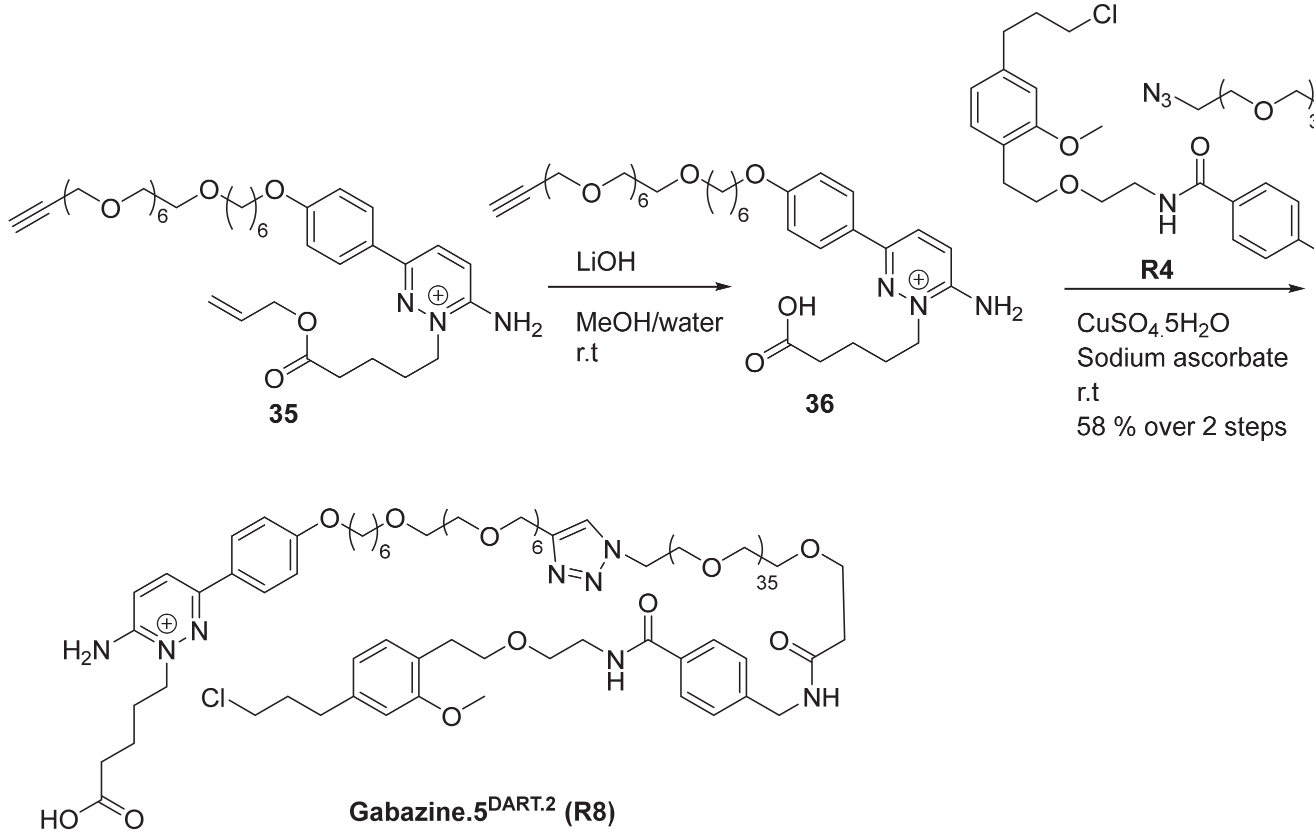


**Gabazine.5$^{DART.2}$ (R8)**

To allyl 5-(3-(4-((4,7,10,13,16,19,22-heptaoxaoctacos-1-yn-28-yl)oxy)phenyl)-6-iminopyridazin-1(6H)-yl)pentanoate, **35** (15 mg, 20 µmol) in MeOH/H2O(1/1, 2 ml) was added lithium hydroxide hydrate (5.1 mg, 120 µmol) at r.t, and the reaction was stirred for 2h. LCMS indicated formation of **36** and the pH was adjusted to 8 by addition of 1N HCl (110 ul). The mixture was concentrated under a stream of nitrogen. To the crude mixture was added EtOH/t-BuOH/water (2:1:1, 1 mL) after which azido-PEG36-HTL2.0, **R4** (42 mg, 20 µmol) was added. Finally, Cu(II) sulfate pentahydrate (0.45 mg, 1.8 µmol) and sodium ascorbate (0.32 mg, 1.8 µmol) were added and the reaction mixture was stirred at room temperature overnight after which it was purified by reverse phase prep HPLC eluting with 20 to 90 % acetonitrile in water (0.1 % formic acid, 200 nm wavelength for collection, and other conditions reported in the general chemical synthesis section) to give 5-(3-(4-((1-(1-(1-(4-((2-(4-(3-chloropropyl)-2-methoxyphenethoxy)ethyl)carbamoyl)phenyl)-3-oxo 6,9,12,15,18,21,24,27,30,33,36,39,42,45,48,51,54,57,60,63,66,69,72,75,78,81,84,87,90,93,96,99,102,105,108 ,111-hexatriacontaoxa-2-azatridecahectan-113-yl)-1H-1,2,3-triazol-4-yl)-2,5,8,11,14,17,20-heptaoxahexacosan-26-yl)oxy)phenyl)-6-iminopyridazin-1(6H)-yl)pentanoic acid, **Gabazine.5$^{DART.2}$, R8** (32.0 mg, 2 steps 58%) $^1$H NMR (500 MHz, $D_2O$, water suppression used- the peaks close to the water peak between 4.4 and 4.6 ppm were partially suppressed) δ 7.89 (s, 1H), 7.87 (d, J = 9.6 Hz, 1H), 7.62 (d, J = 8.4 Hz, 2H), 7.47 (d, J = 8.0 Hz, 2H), 7.33 (d, J = 9.4 Hz, 1H), 7.17 (d, J = 8.1 Hz, 2H), 6.79 (d, J = 8.5 Hz, 2H), 6.75 (d, J = 7.4 Hz, 1H), 6.46 (s, 1H), 6.31 (d, J = 7.6 Hz, 1H), 4.47 (s, 2H), 4.42 (t, J = 5.0 Hz, 2H), 4.23 (s, 2H), 4.07 (t, J = 7.4 Hz, 2H), 3.76 (t, J = 5.1 Hz, 4H), 3.70 – 3.34 (m, 170H), 3.29-3.23 (m, 5H), 2.55 (t, J = 6.6 Hz, 2H), 2.40 (t, J = 6.0 Hz, 2H), 2.34 (t, J = 7.5 Hz, 2H), 2.12 (t, J = 7.3 Hz, 2H), 1.72-1.65 (m, 4H), 1.53-1.43 (m, 4H), 1.38-1.32 (m, 2H), 1.22-1.10 (m, 4H). HRMS (ESI$^+$): *m/z* calcd for $C_{133}H_{232}ClN_8NaO_{50}$: 1400.2733 $[M+H+Na]^{2+}$; found: 1400.2734 $[M+H+Na]^{2+}$

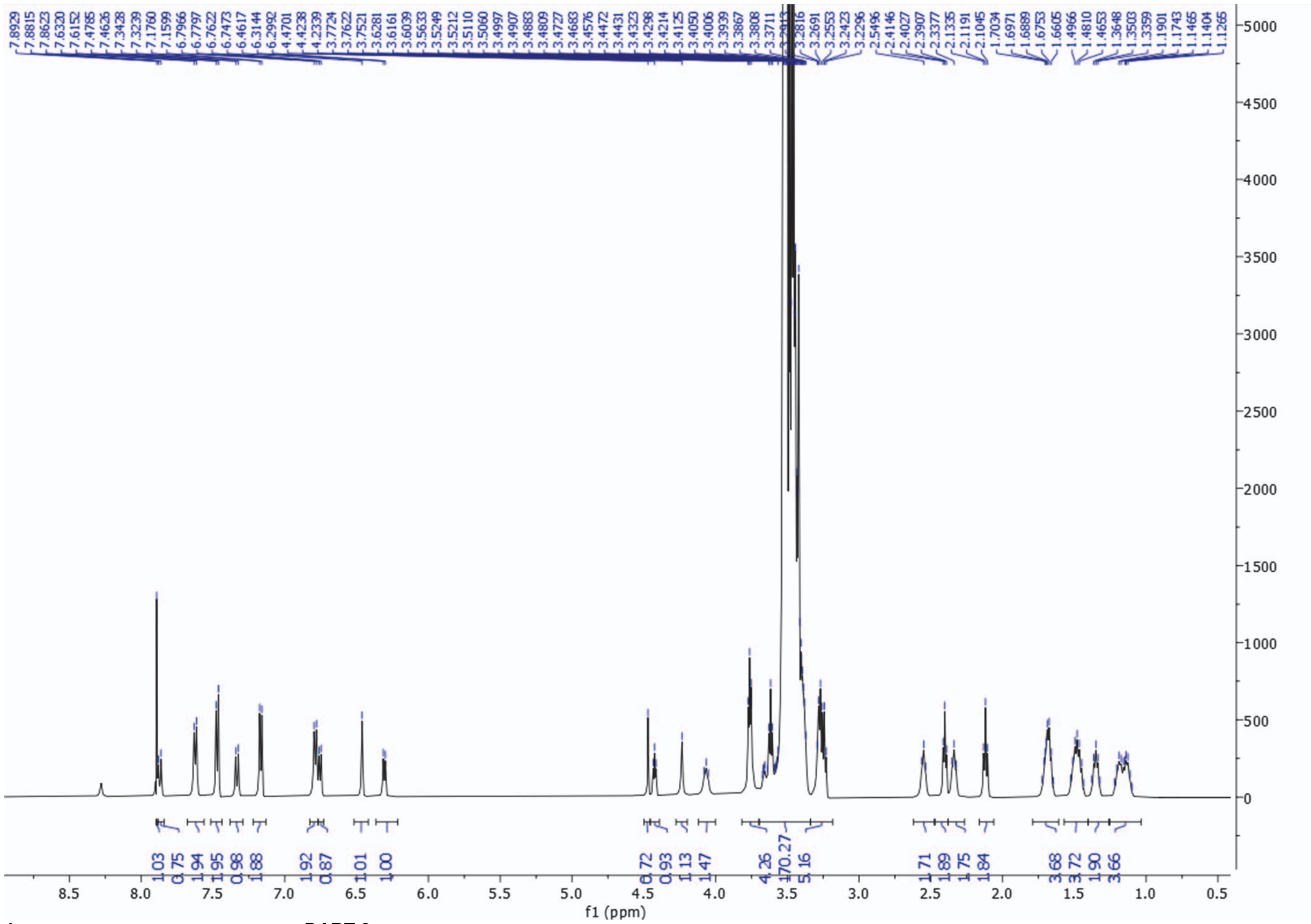

**$^{1}$H NMR of Gabazine.5$^{DART.2}$ (R8) in $D_2O$ (water suppression)**

### Synthesis of R9–gabazine.7$^{DART.2}$

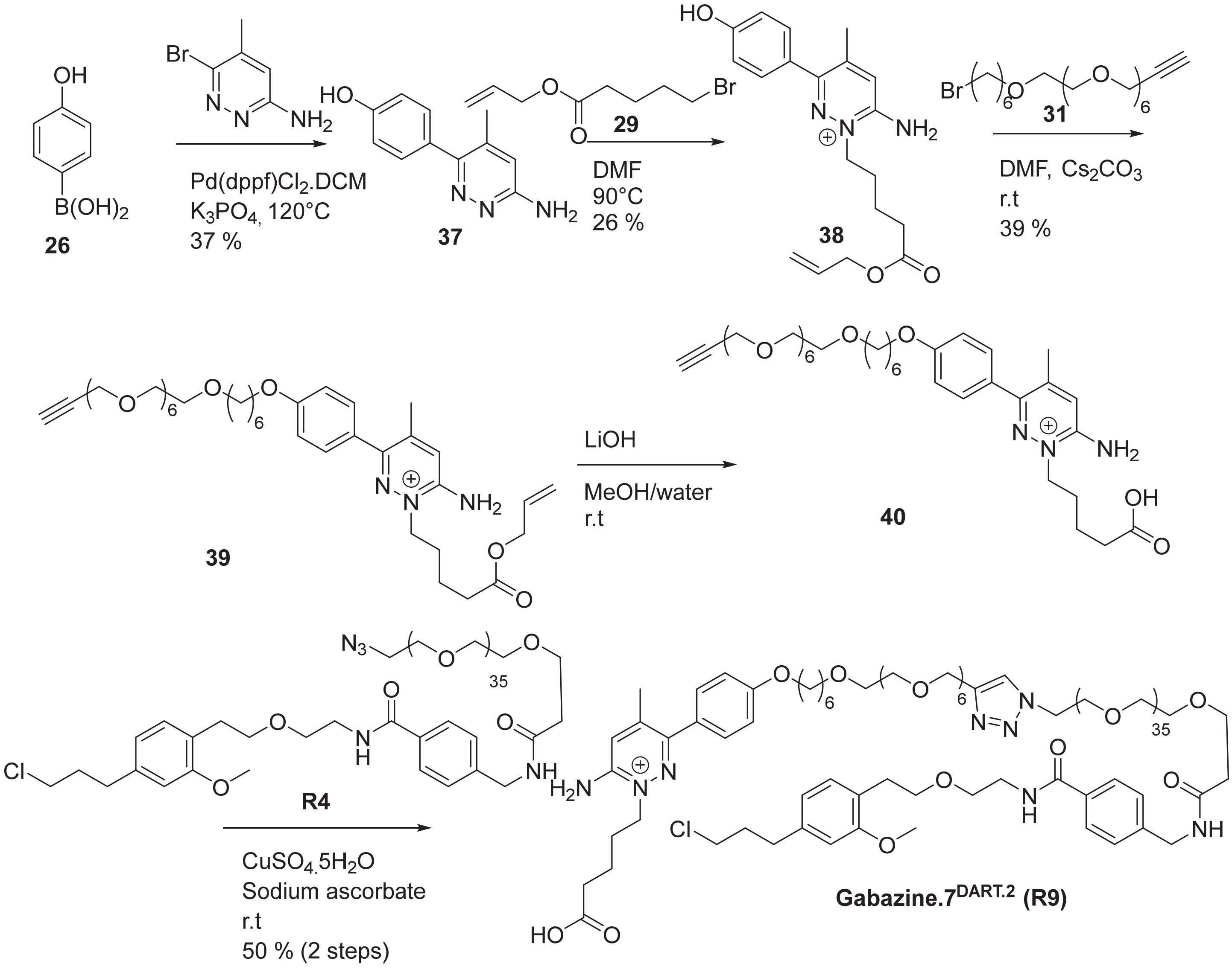

Step 1: Synthesis of 4-(6-amino-4-methylpyridazin-3-yl)phenol, **37**

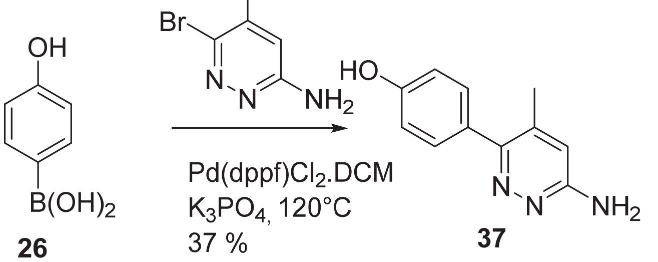

To (4-hydroxyphenyl)boronic acid, **26** (1 g, 7.25 mmol) was added $K_3PO_4$ (3.08 g, 14.5 mmol), Pd(dppf)Cl2·DCM (0.355 g, 0.435 mmol), 6-bromo-5-methylpyridazin-3-amine (0.937 g, 6.53mmol) and dioxane (7 mL)/water (4 mL). The suspension was stirred and degassed for 5 min after which it was heated at 120 degrees for 17 min, then concentrated. The material was then taken up in water and a slight amount of DCM, stirred thoroughly, then filtered. The mixture was filtered to give 4-(6-amino-4-methylpyridazin-3-yl)phenol, **37** (1.132 g, 5.62 mmol, 77 % yield) as a dark, brown solid, containing some aryl halide. Used directly without further purification. $^1$H NMR (500 MHz, $CD_3OD$) δ 7.30 (d, *J* = 8.6 Hz, 2H), 6.89 (d, *J* = 8.6 Hz, 2H), 6.86 (d, *J* = 1.1 Hz, 1H), 2.23 (d, *J* = 1.0 Hz, 3H). (ES-LCMS) *m/z* 202.2 (M+H)$^+$

Step 2: Synthesis of allyl 5-(3-(4-hydroxyphenyl)-6-imino-4-methylpyridazin-1(6H)-yl)pentanoate, **38** (see references 3 and 4)

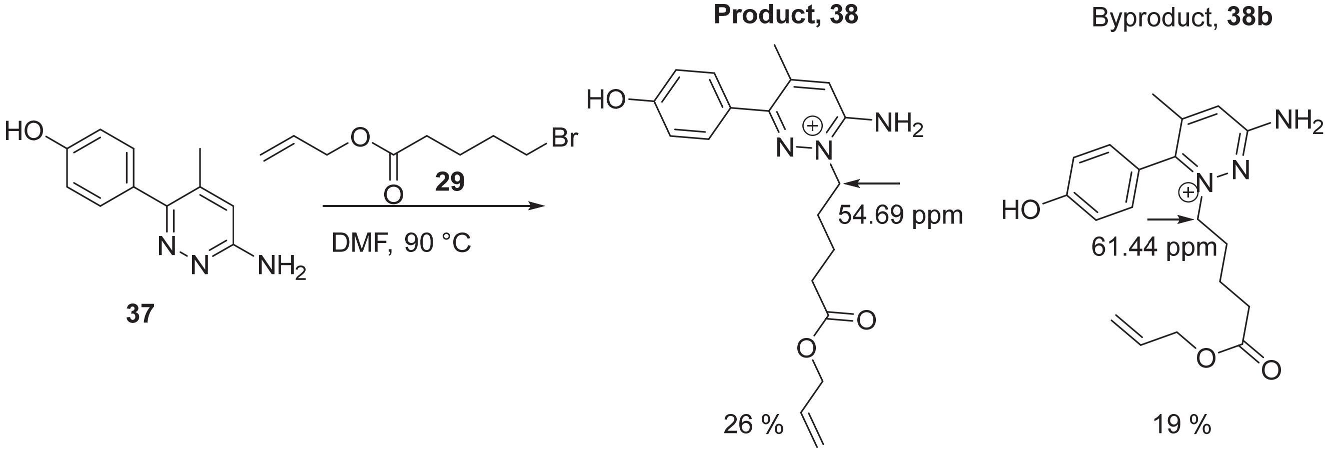


LCMS retention time: 1.13 min 1.08 min

To 4-(6-amino-4-methylpyridazin-3-yl)phenol, **37** (250 mg, 1.24 mmol) was added allyl allyl 5-bromopentanoate, **29** (275 mg, 1.24 mmol) and DMF (2 mL) and the reaction was heated at 90°C for 2 h. The mixture was directly purified by prep HPLC eluting with 10 to 90 % ACN in water (0.1 % FA, 220 nm wavelength for collection, and the rest of the conditions as described in the general chemistry synthesis section) in three 750 uL injections. Relevant fractions were combined and concentrated. The undesired byproduct eluted first and the desired product eluted later. *Since three nitrogens and the phenolic oxygen may all react with the alkyl bromide, the structures were fully assigned and confirmed by H/C NMR, HSQC, and HMBC in DMSO-d6. The most distinguishing peak for the product and byproduct is the $CH_2$ that is directly connected to the nitrogen which reacted. The $^{13}C$ NMR of the $CH_2$ immediately adjacent to the nucleophilic nitrogen has a shift of 54.69 ppm in the desired product vs 61.44 ppm in the undesired byproduct. The $^{13}C$ NMR shift of this $CH_2$ in the desired compound is the same as the shift of a $CH_2$ adjacent to the same nucleophilic nitrogen in a similar compound reported in reference 4 in which the structure was confirmed by X-ray crystallography. The regioselectivity of the alkylation of pyridazine has been documented in reference 3.* The reactivities of the structurally related analogs, Gabazine 1, Gabazine 5, and Gabazine 7, are very similar: in all cases, the undesired byproduct eluted first on RP HPLC. Additionally, the shifts of the $CH_2$ directly connected to the same nitrogen are very similar for all 3 analogs (see figure below). It worthwhile to mention that the products and byproducts exist as tautomers under acidic and basic conditions. Furthermore, the undesired byproduct does not react in the following alkylation step (the reaction with the alkyl bromide in DMF with cesium carbonate) while the desired product does react, further confirming our structural assignments. Desired product **38** yield: 108 mg (26 %). Byproduct **38b** yield: 80 mg (19 %). The NMR shifts of each compound in both $CD_3OD$ and DMSO-$d_6$ are reported below.

**Desired Product:** $^{1}H$ NMR (500 MHz, **$CD_3OD$**) δ 8.35 (br.s, 1H), 7.43 – 7.39 (m, 3H), 6.93 (d, $J$ = 8.6 Hz, 2H), 5.97-5.89 (m, 1H), 5.30 (dq, $J$ = 17.2, 1.7 Hz, 1H), 5.21 (dt, $J$ = 10.5, 1.5 Hz, 1H), 4.58 (dt, $J$ = 5.6, 1.5 Hz, 2H), 4.36 (t, $J$ = 7.2 Hz, 2H), 2.48 (t, $J$ = 7.3 Hz, 2H), 2.42 (s, 3H), 2.01 – 1.95 (m, 2H), 1.79-1.73 (m, 2H). $^{13}C$ NMR (126 MHz, **$CD_3OD$**) δ 173.06, 159.15, 154.36, 152.42, 145.60, 132.27, 130.21, 124.56, 123.28, 116.95, 115.04, 64.77, 55.03, 32.73, 25.92, 21.18, 18.99. $^{1}H$ NMR (500 MHz, **DMSO-$d_6$**) δ 8.48 (s, 1H), 7.50 (s, 1H), 7.39 (d, $J$ = 8.3 Hz, 2H), 6.91 (d, $J$ = 8.4 Hz, 2H), 5.95-5.87 (m, 1H), 5.29 (dt, $J$ = 17.2, 1.7 Hz, 1H), 5.21 (dd, $J$ = 10.5, 1.6 Hz, 1H), 4.55 (dd, $J$ = 4.4, 2.8 Hz, 2H), 4.29 (t, $J$ = 7.2 Hz, 2H), 2.43 (t, $J$ = 7.4 Hz, 2H), 2.32 (s, 3H), 1.85-1.79 (m, 2H), 1.67-1.60 (m, 2H). $^{13}C$ NMR (126 MHz, **DMSO-$d_6$**) δ 172.67, 159.57, 152.94, 152.48, 144.20, 133.10, 130.75, 124.41, 124.34, 118.06, 115.70, 64.67, 54.69, 33.18, 26.49, 21.52, 20.13. (ES-LCMS) $m/z$ 342.3 $(M+H)^+$

**Byproduct:** $^{1}H$ NMR (500 MHz, **$CD_3OD$**) δ 8.32 (br.s, 1H), 7.41 (s, 1H), 7.36 – 7.25 (m, 2H), 7.12 – 6.98 (m, 2H), 5.98-5.90 (m, 1H), 5.31 (dt, $J$ = 17.3, 1.7 Hz, 1H), 5.23 (dd, $J$ = 10.4, 1.6 Hz, 1H), 4.57 (dd, $J$ = 5.7, 1.6 Hz, 2H), 4.27 (t, $J$ = 7.4 Hz, 2H), 2.31 (t, $J$ = 7.2 Hz, 2H), 2.13 (s, 3H), 1.99-1.92 (m, 2H), 1.59-1.53 (m, 2H).

$^{13}$C NMR (126 MHz, **$CD_3OD$**) δ 172.80, 160.60, 160.04, 151.83, 146.60, 132.27, 129.52, 122.20, 119.60, 116.97, 116.25, 64.77, 61.52, 32.48, 28.54, 21.11, 18.70. $^{1}$H NMR (500 MHz, **DMSO-$d_6$**) δ 8.53 (br.s, 1H), 7.85 (s, 2H), 7.49 (s, 1H), 7.38 (d, *J* = 8.4 Hz, 2H), 7.06 (d, *J* = 8.3 Hz, 2H), 5.99-5.92 (m, 1H), 5.34 (dq, *J* = 17.2, 1.7 Hz, 1H), 5.27 (dt, *J* = 10.4, 1.5 Hz, 1H), 4.58 (dt, *J* = 5.5, 1.5 Hz, 2H), 4.19 (t, *J* = 7.4 Hz, 2H), 2.31 (t, *J* = 7.3 Hz, 2H), 2.08 (s, 3H), 1.90 – 1.84 (m, 2H), 1.52-1.46 (m, 2H). $^{13}$C NMR (126 MHz, **DMSO-$d_6$**) δ 172.41, 160.41, 160.35, 151.86, 146.48, 133.06, 130.10, 122.68, 119.39, 118.17, 116.68, 64.73, 61.44, 32.91, 28.64, 21.46, 19.97. (ES-LCMS) *m/z* 342.3 (M+H)$^+$

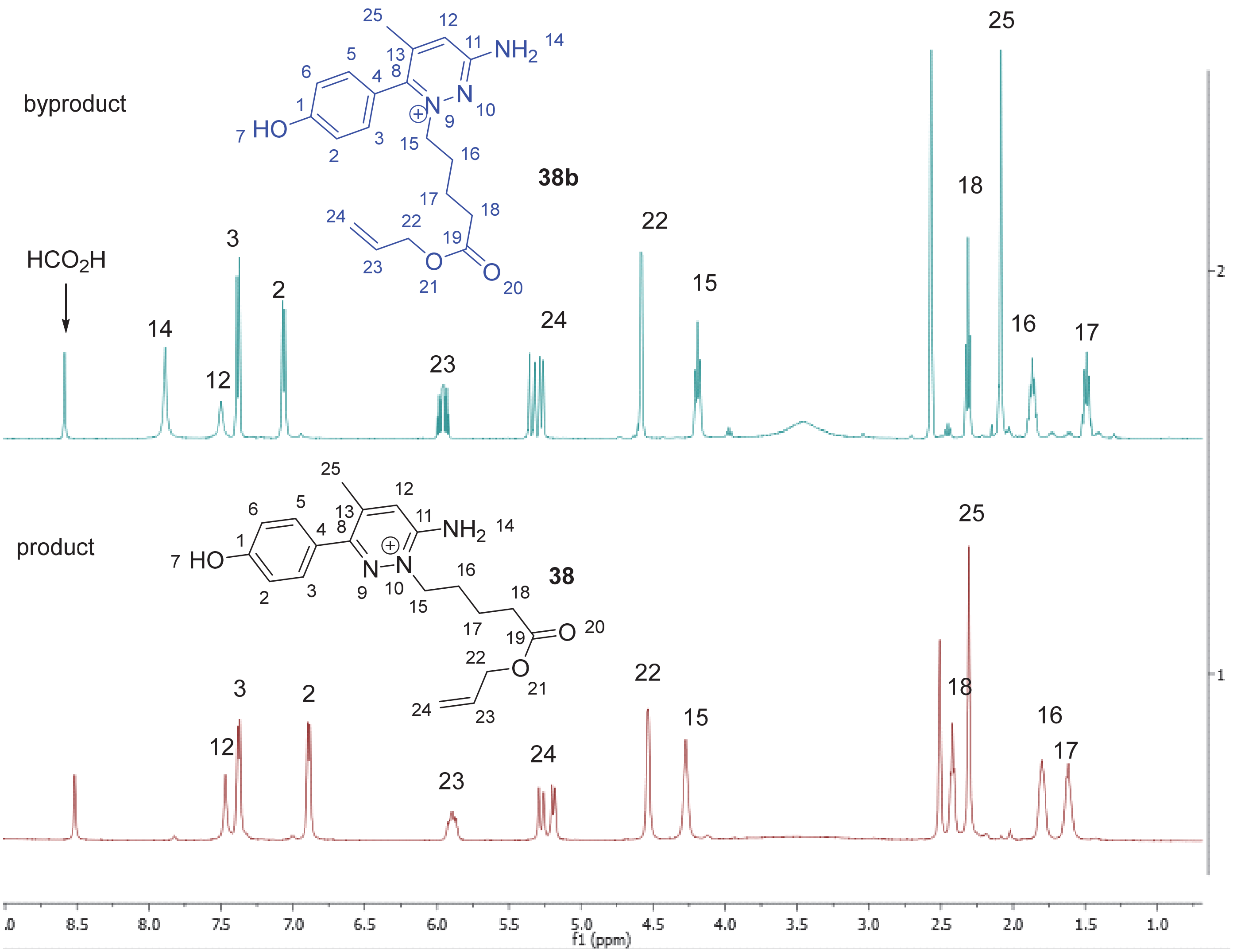


**Comparison of $^{1}$H NMR for product (38) and byproduct (38b) in DMSO-$d_6$**

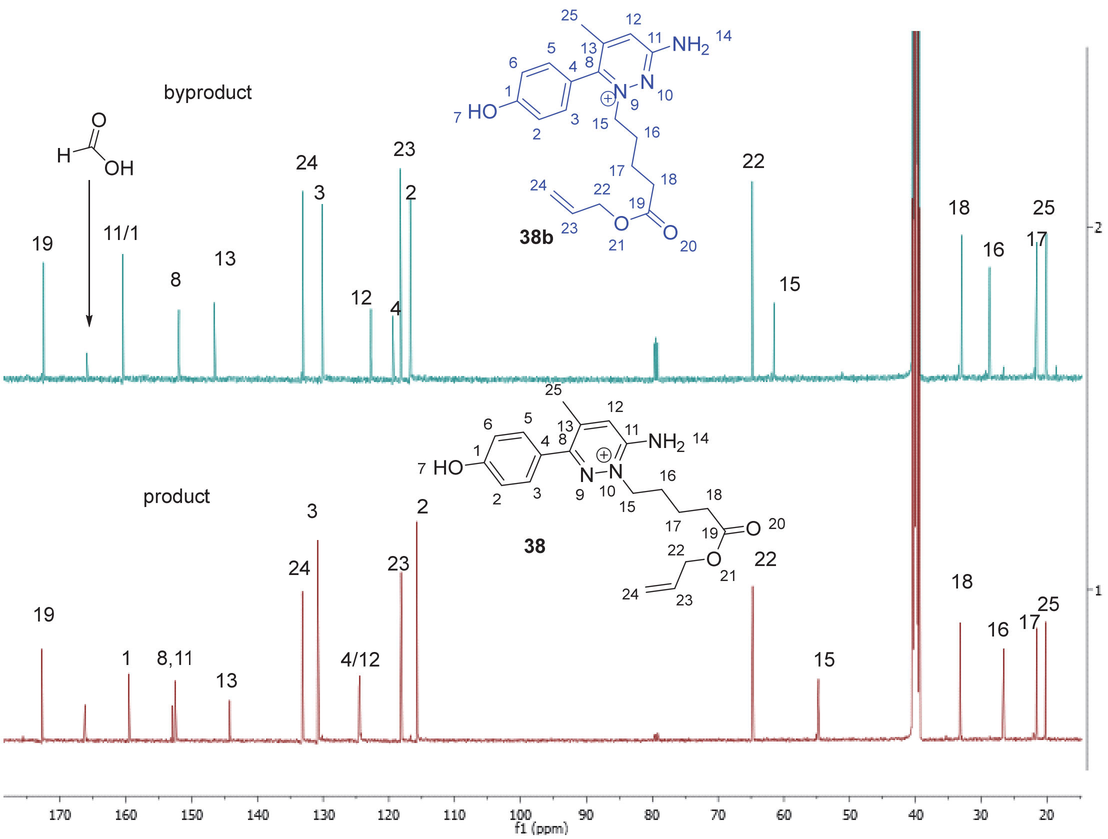


**Comparison of $^{13}C$ NMR for product (38) and byproduct (38b) in DMSO-d$_6$**

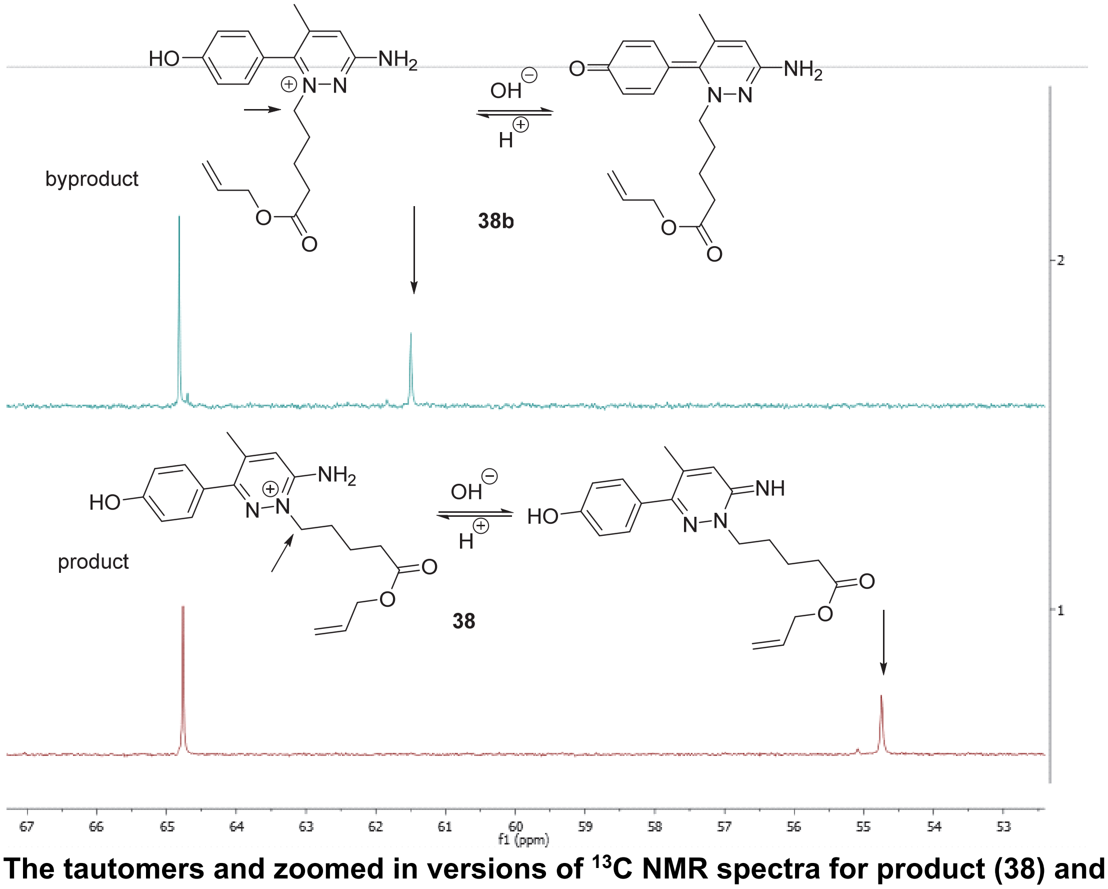


**The tautomers and zoomed in versions of $^{13}C$ NMR spectra for product (38) and byproduct (38b) in DMSO-d$_6$**

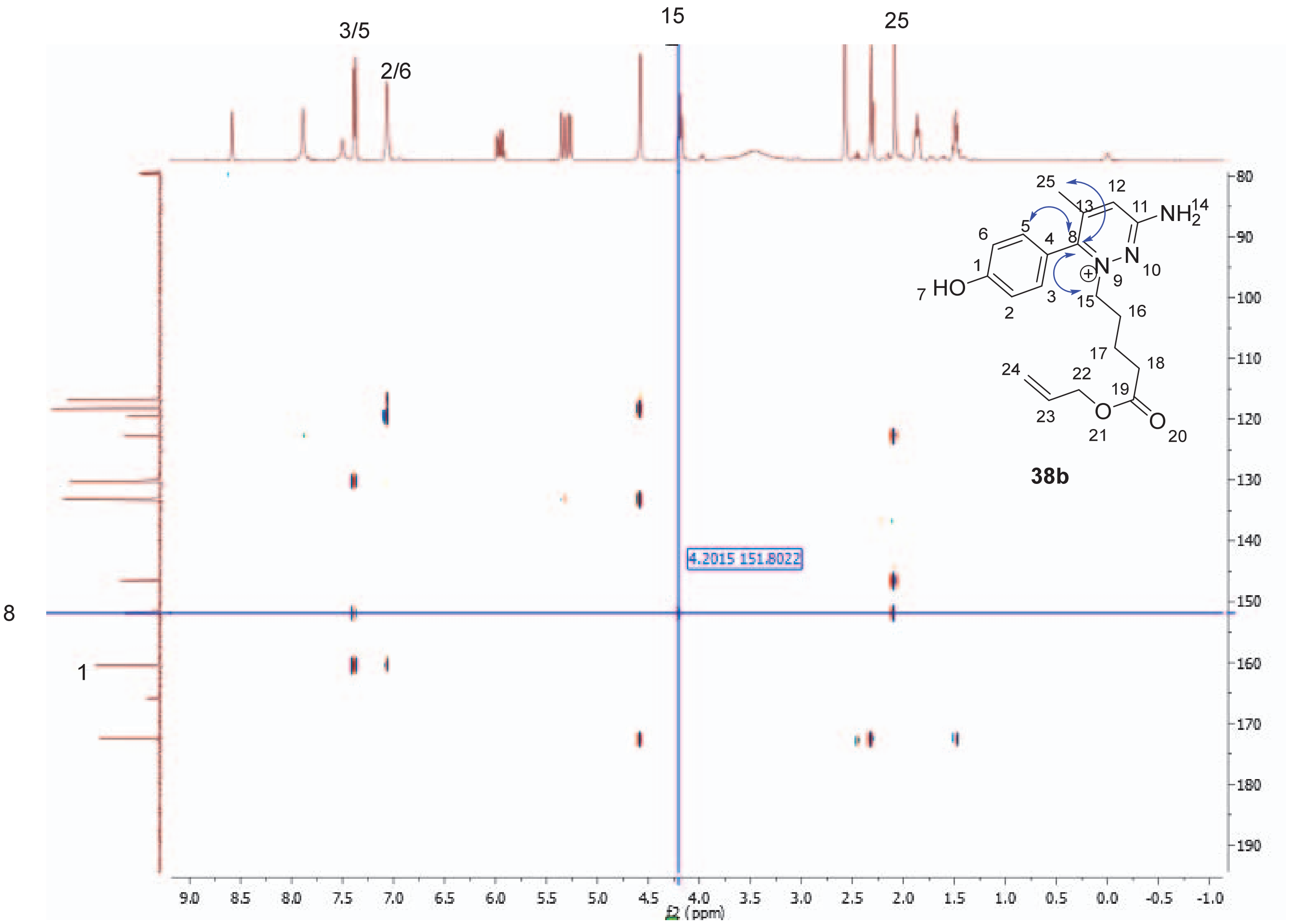


**The correlation of H5 and C8 & H15 and C8 in HMBC(DMSO-$d_6$) confirms the structure of the byproduct (38b)**

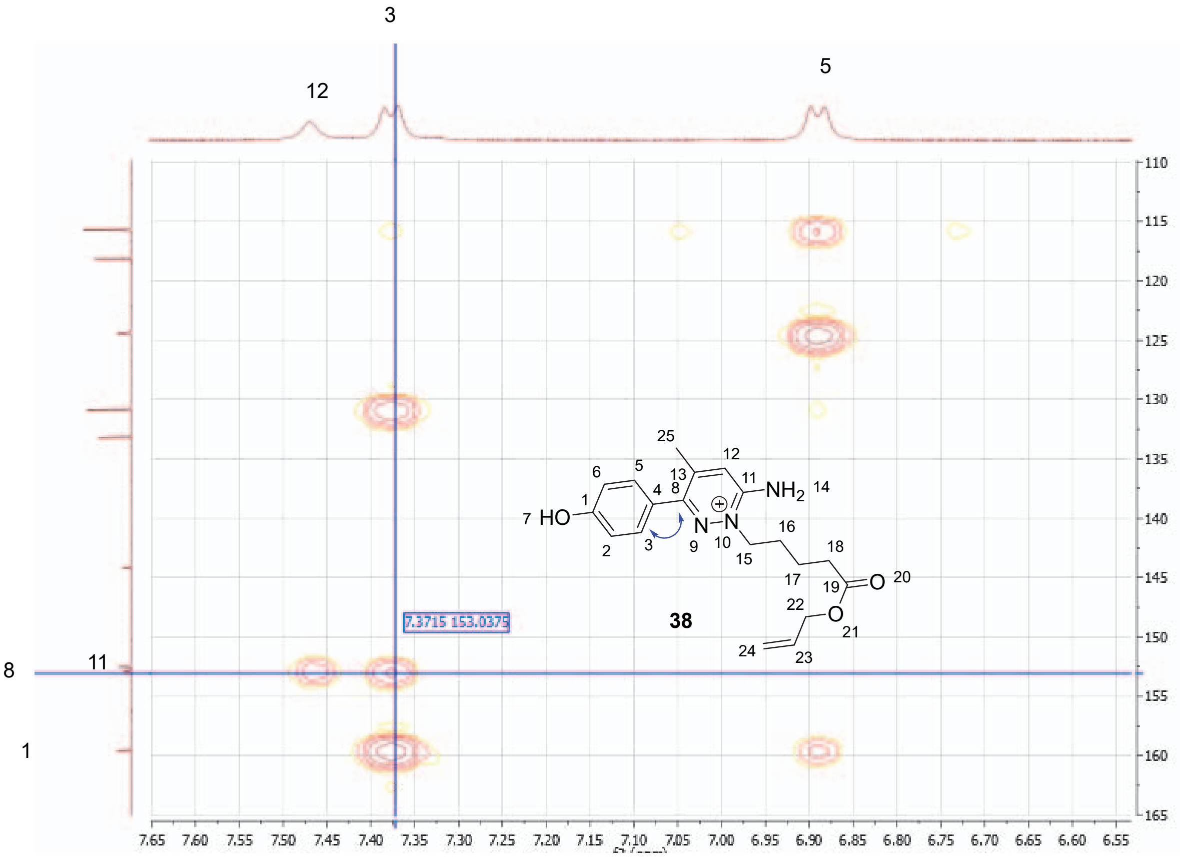


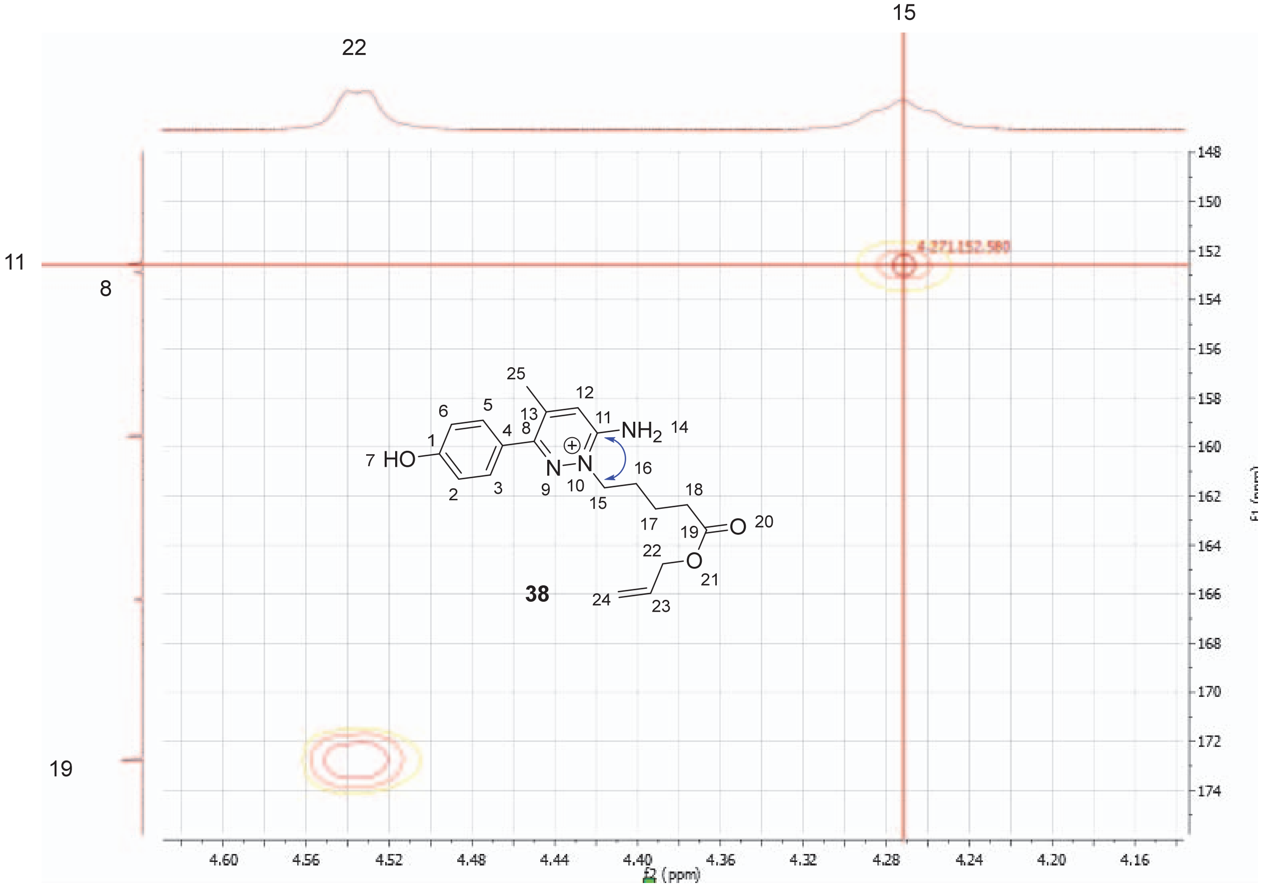


**The correlation of H3 and C8 (top) & H15 and C11 (bottom) in HMBC(DMSO-$d_6$) confirmed the structure of product, 38**

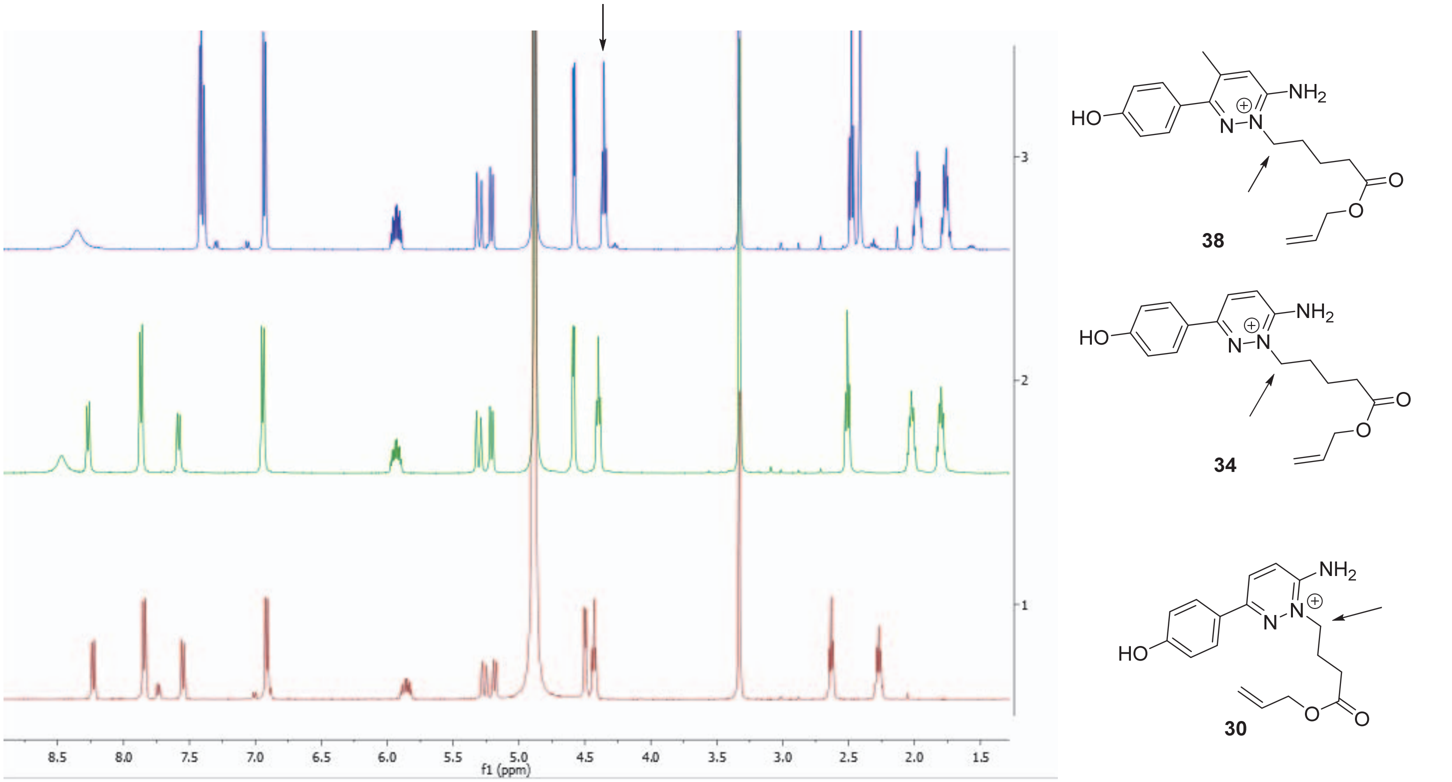


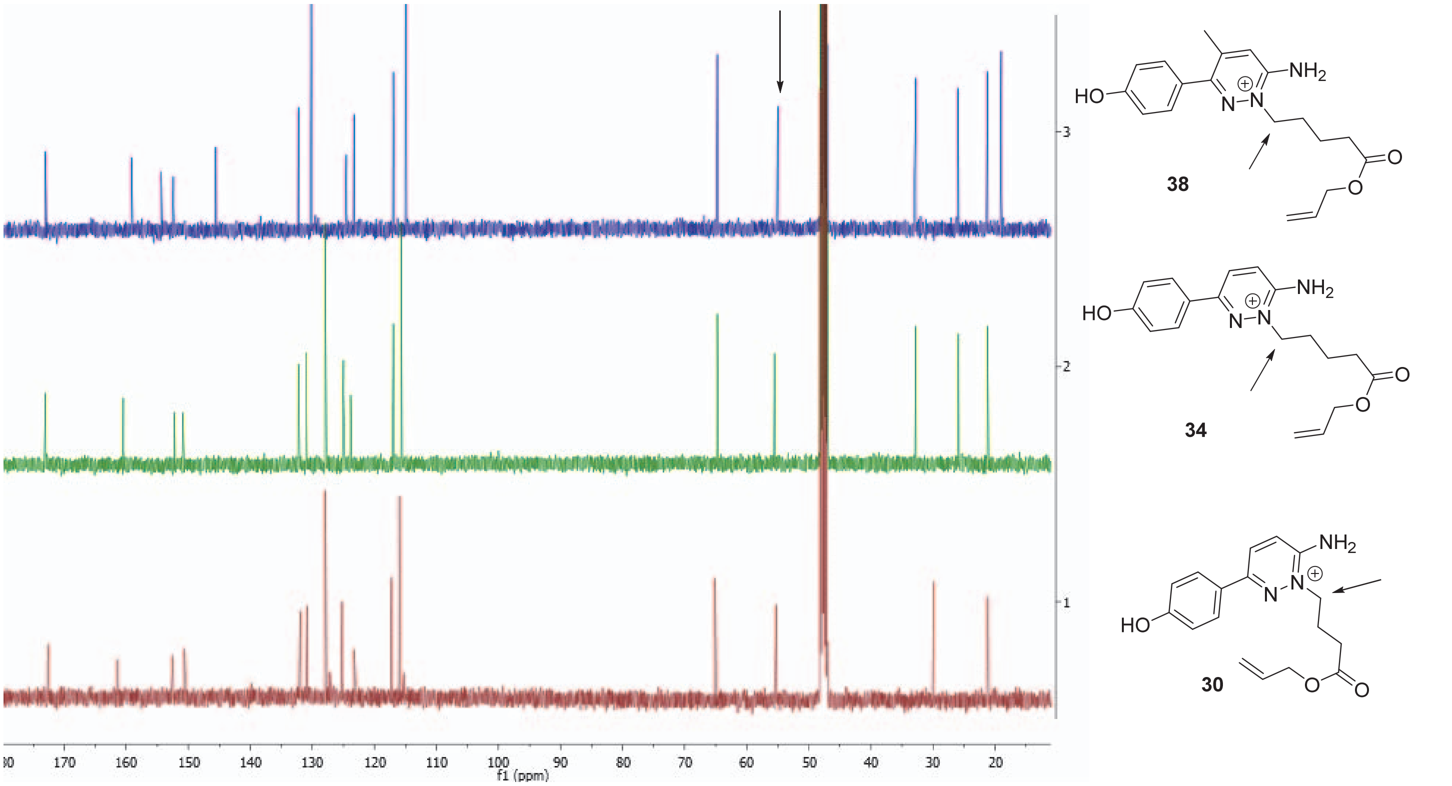


**Comparison of $^{1}$HNMR (top), $^{13}$C NMR (bottom) in $CD_3OD$ of desired intermediates for Gabazine.1 (30), Gabazine.5 (34), and Gabazine.7 (38)**

Step 3: allyl 5-(3-(4-((4,7,10,13,16,19,22-heptaoxaoctacos-1-yn-28-yl)oxy)phenyl)-6-imino-4-methylpyridazin-1(6H)-yl)pentanoate, **39**

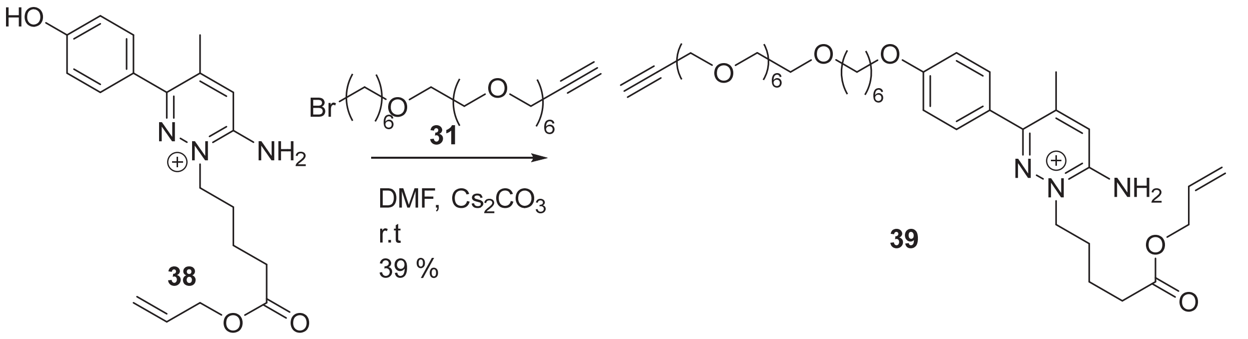


To allyl 5-(3-(4-hydroxyphenyl)-6-imino-4-methylpyridazin-1(6H)-yl)pentanoate, **38** (40 mg, 0.12 mmol) was added anhydrous DMF (1 mL) after which 28-bromo-4,7,10,13,16,19,22-heptaoxaoctacos-1-yne, **31** (85 mg, 0.18 mmol) and anhydrous cesium carbonate (76 mg, 0.23 mmol) were added and the reaction was stirred for 3 h. DMF was removed under a stream of nitrogen, and the crude was purified by reverse phase prep HPLC eluting with 5 to 50 % acetonitrile in water (0.1 % formic acid conditions and other conditions as described in the general chemical synthesis section) to give allyl 5-(3-(4-((4,7,10,13,16,19,22-heptaoxaoctacos-1-yn-28-yl)oxy)phenyl)-6-imino-4-methylpyridazin-1(6H)-yl)pentanoate, **39** (34 mg, 46 μmol, 39 % yield) as a red/yellow oil. $^1$H NMR (500 MHz, $CD_3OD$) δ 8.49 (s, 1H), 7.57 – 7.45 (m, 2H), 7.39 (d, *J* = 1.3 Hz, 1H), 7.09 – 7.06 (m, 2H), 5.97-5.89 (m, 1H), 5.30 (dq, *J* = 17.2, 1.6 Hz, 1H), 5.21 (dq, *J* = 10.4, 1.4 Hz, 1H), 4.59 (dt, *J* = 5.7, 1.5 Hz, 2H), 4.36 (t, *J* = 7.2 Hz, 2H), 4.20 (d, *J* = 2.4 Hz, 2H), 4.08 (t, *J* = 6.4 Hz, 2H), 3.70 – 3.64 (m, 21H), 3.61 – 3.59 (m, 2H), 3.52 (t, *J* = 6.5 Hz, 2H), 2.86 (t, *J* = 2.4 Hz, 1H), 2.48 (t, *J* = 7.3 Hz, 2H), 2.42 (s, 3H), 2.01 – 1.95 (m, 2H), 1.88 – 1.82 (m, 2H), 1.79-1.73 (m, 2H), 1.67-1.62 (m, 2H), 1.59-1.53 (m, 2H), 1.51-1.45 (m, 2H). ES-LCMS) *m/z* 744.6 $(M+H)^+$

Steps 4/5: Synthesis of **Gabazine.7$^{DART.2}$ (R9)**

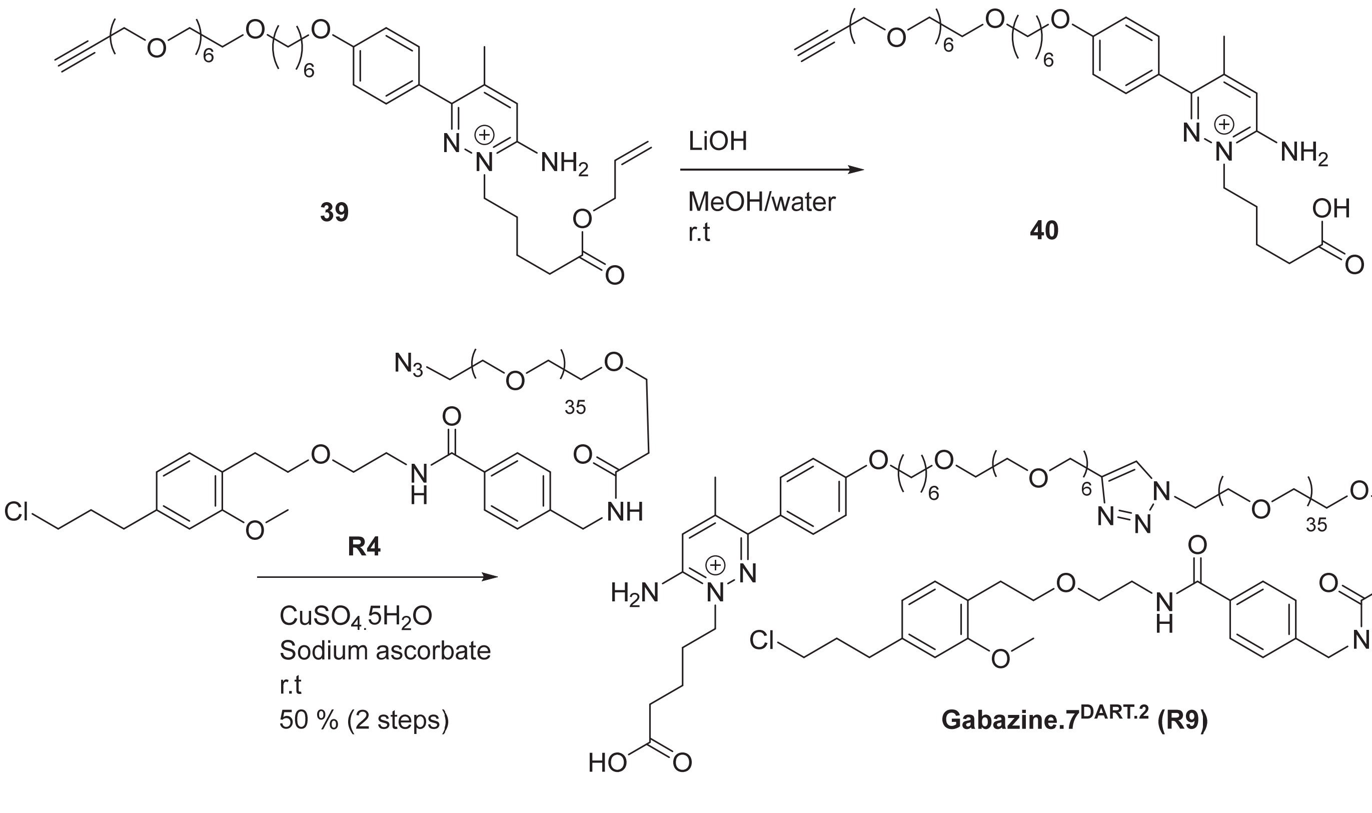

To allyl 5-(3-(4-((4,7,10,13,16,19,22-heptaoxaoctacos-1-yn-28-yl)oxy)phenyl)-6-imino-4-methylpyridazin-1(6H)-yl)pentanoate, **39** (15 mg, 20.0 μmol) in MeOH/water(1:1, 2 ml) was added lithium hydroxide hydrate (5.1 mg, 120 μmol) at room temperature and the reaction was stirred for 2h after which the pH was adjusted to 8 by addition of 1N HCl (110 μL) and solvent was removed under a stream of nitrogen. The crude material **40** was taken up in EtOH/t-BuOH/water (2:1:1, 1 mL) after which azido-PEG36-HTL2.0, **R4** (38 mg, 18 μmol) was added. Finally, Cu(II) sulfate pentahydrate (0.45 mg, 1.8 μmol) and sodium ascorbate (0.32 mg, 1.8 μmol) were added and the reaction mixture was stirred at room temperature overnight after which it was purified by reverse phase prep HPLC eluting with 20 to 90 % acetonitrile in water (0.1 % formic acid, 200 nm wavelength for collection, and other conditions reported in the general chemical synthesis section) to give 5-(3-(4-((1-(1-(1-(4-((2-(4-(3-chloropropyl)-2-methoxyphenethoxy)ethyl)carbamoyl)phenyl)-3-oxo-6,9,12,15,18,21,24,27,30,33,36,39,42,45,48,51,54,57,60,63,66,69,72,75,78,81,84,87,90,93,96,99,102,105,108,111-hexatriacontaoxa-2-azatridecahectan-113-yl)-1H-1,2,3-triazol-4-yl)-2,5,8,11,14,17,20-heptaoxahexacosan-26-yl)oxy)phenyl)-6-imino-4-methylpyridazin-1(6H)-yl)pentanoic acid, **Gabazine.7$^{DART.2}$, R9** (25.0 mg, 50% over 2 steps) $^{1}$H NMR (500 MHz, $D_2O$, water suppression used- the peaks close to the water peak between 4.4 and 4.6 ppm were partially suppressed) δ 7.91 (s, 1H), 7.49 (d, *J* = 7.9 Hz, 2H), 7.30 (d, *J* = 8.2 Hz, 2H), 7.22 (d, *J* = 8.0 Hz, 2H), 7.19 (s, 1H), 6.89 (d, *J* = 8.3 Hz, 2H), 6.83 (d, *J* = 7.6 Hz, 1H), 6.56 (s, 1H), 6.36 (d, *J* = 7.5 Hz, 1H), 4.49 (s, 2H), 4.44 (t, *J* = 5.2 Hz, 2H), 4.28 (s, 2H), 4.09 (t, *J* = 7.4 Hz, 2H), 3.87 (t, *J* = 6.5 Hz, 2H), 3.78 (t, *J* = 5.1 Hz, 3H), 3.68-3.62 (m, 5H), 3.53 – 3.43 (m, 183H), 3.38-3.28 (m, 6H), 2.60 (t, *J* = 6.6 Hz, 2H), 2.43 – 2.37 (m, 4H), 2.12 (s, 3H), 2.07 (t, *J* = 7.5 Hz, 2H), 1.75-1.66 (m, 4H), 1.61-1.56 (m 2H), 1.49-1.38 (dp, *J* = 28.1, 7.4 Hz, 4H), 1.30-1.24 (m, 2H), 1.23-1.17 (m, 2H). HRMS (ESI$^+$): *m/z* calcd for $C_{134}H_{234}ClN_8NaO_{50}$: 1407.2812 [M+H+Na]$^{2+}$; found: 1407.2808 [M+H+Na]$^{2+}$

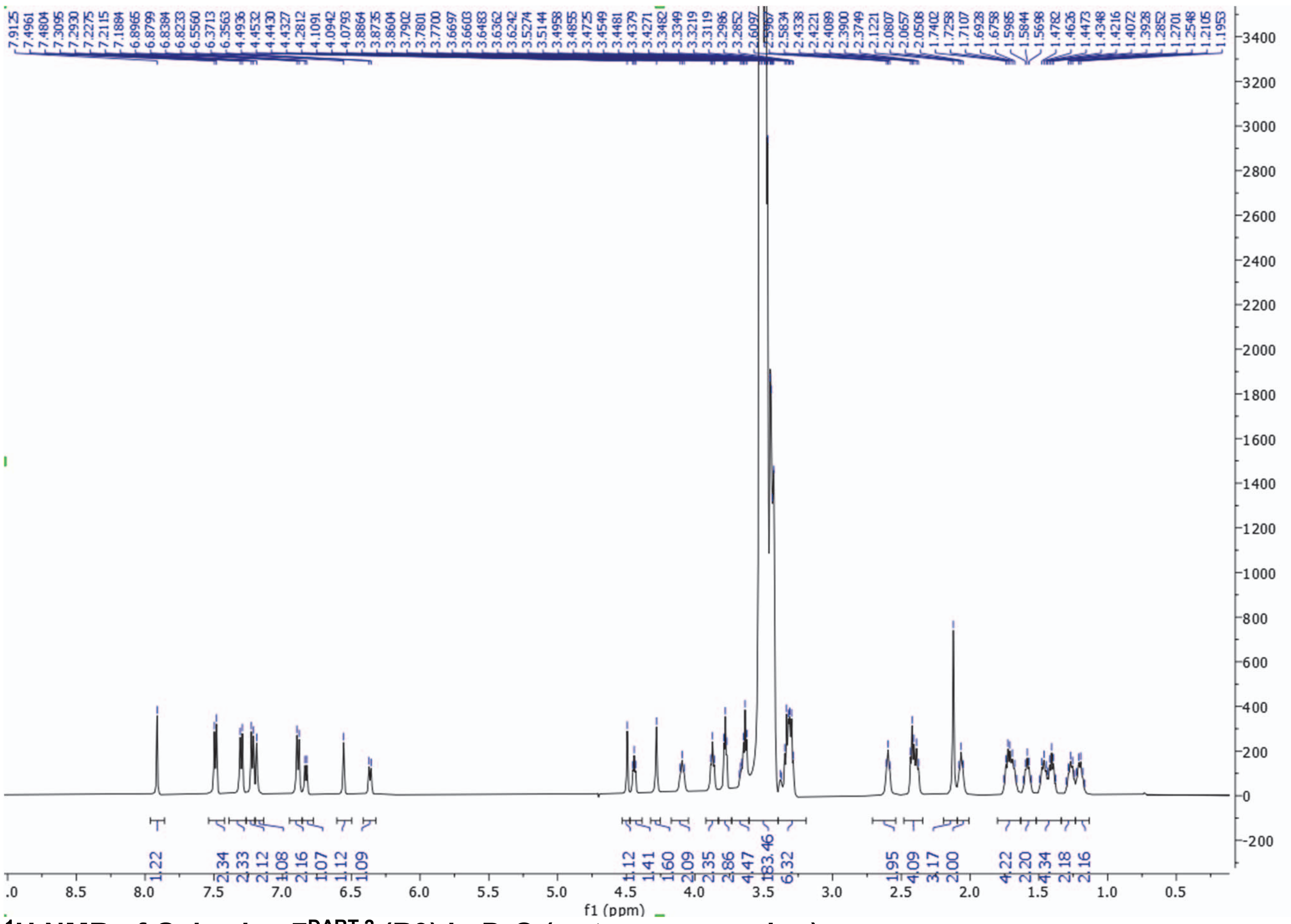

**$^{1}$H NMR of Gabazine.7$^{DART.2}$ (R9) in $D_2O$ (water suppression)**

**Synthesis of R10—Alexa488.1DART.2**

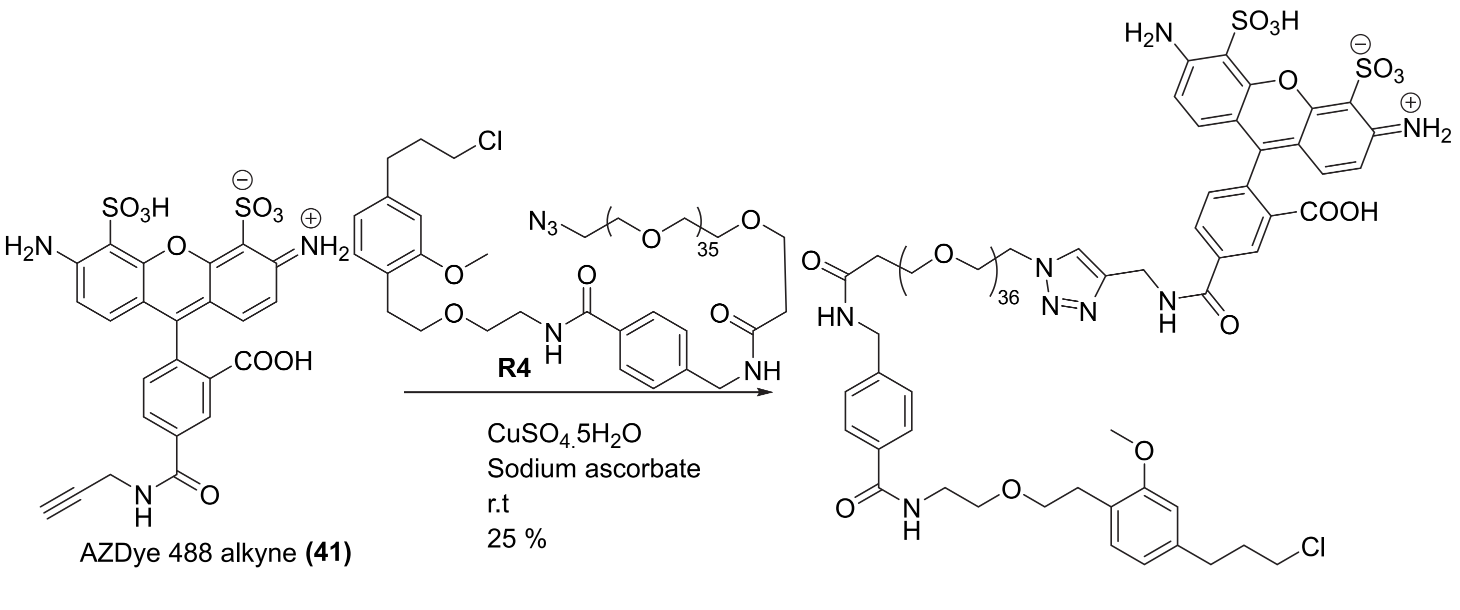


Note: the attachment point of the alkyne in AF488 alkyne, **41** (Click Chemistry Tools; Catalog # 1277-25, MW 773.91 g/mol reported, 571.53 g/mol protonated) or the linker length is not directly disclosed by the vendor, however, based on LCMS as well as $^{1}H$ NMR shown below, we are reasonably confident of the structure as drawn (presumably exists as a di-triethylamine salt).

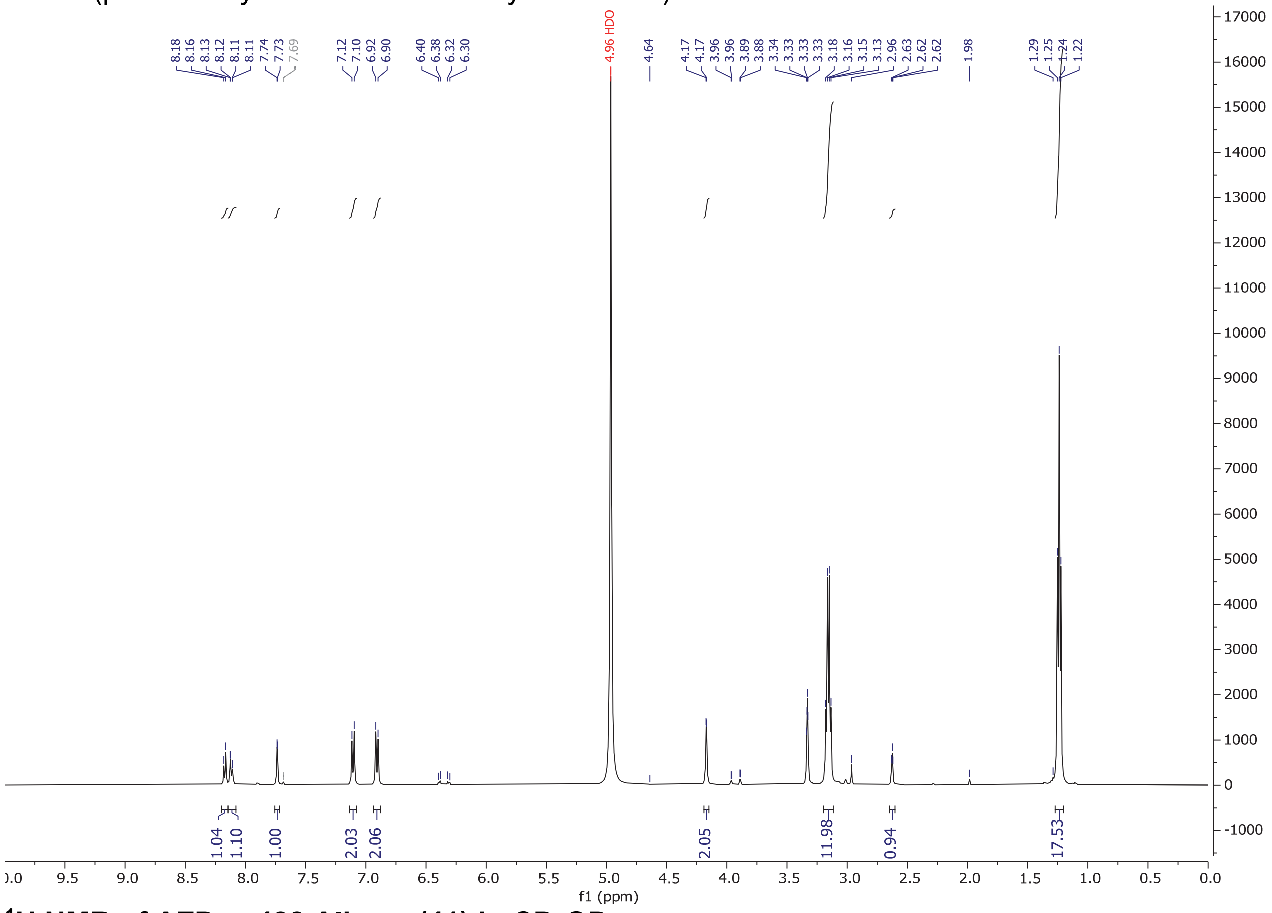


**$^{1}H$ NMR of AZDye 488 Alkyne (41) in $CD_3OD$**

To azido-PEG36-HTL2.0, **R4** (13 mg, 6.5 μmol) was added AF488 alkyne, **41** (5 mg, 6.5 μmol) followed by EtOH/t-BuOH/water (2:1:1, 0.8 mL). Then, copper (II) sulfate pentahydrate (0.1 mg, 0.325 μmol) and sodium ascorbate (0.1 mg, 0.65 μmol) were added and the reaction was stirred overnight after which it was concentrated under a stream of nitrogen, taken up in acetonitrile/DMSO and purified by 15 to 80 % acetonitrile in water (0.1 % FA) with a Phenomenex 50 x 30 mm column, a 12 min gradient, 50 mL/min flow rate, and collection wavelength of 230 nm. Relevant fractions were concentrated to give 6-amino-9-(2-carboxy-4-(((1-(1-(4-((2-(4-(3-chloropropyl)-2-methoxyphenethoxy)ethyl)carbamoyl)phenyl)-3-oxo-6,9,12,15,18,21,24,27,30,33,36,39,42,45,48,51,54,57,60,63,66,69,72,75,78,81,84,87,90,93,96,99,102,105,108,111-hexatriacontaoxa-2-azatridecahectan-113-yl)-1H-1,2,3-triazol-4-yl)methyl)carbamoyl)phenyl)-3-iminio-5-sulfo-3H-xanthene-4-sulfonate, **Alexa488.1$^{DART.2}$ (R10)** (5 mg, 29 %). $^{1}$H NMR (500 MHz, $CD_3OD$) δ 8.47 – 8.23 (m, 4H), 8.07 (s, 1H), 7.86 (s, 1H), 7.79 (d, *J* = 7.8 Hz, 2H), 7.42 (d, *J* = 8.0 Hz, 2H), 7.15 – 6.86 (m, 4H), 6.78 (s, 1H), 6.67 (d, *J* = 7.5 Hz, 1H), 4.62 (d, *J* = 61.7 Hz, 3H), 4.48 (d, *J* = 5.4 Hz, 2H), 3.80 (d, *J* = 11.4 Hz, 7H), 3.71 – 3.49 (m, 149H), 2.86 (t, *J* = 7.0 Hz, 2H), 2.73 (t, *J* = 7.4 Hz, 2H), 2.54 (t, *J* = 6.0 Hz, 2H), 2.05 (p, *J* = 6.9 Hz, 2H). (ES-LCMS) *m/z* 665.9 $(M+4H)^{4+}$; Compound decomposed on HRMS.

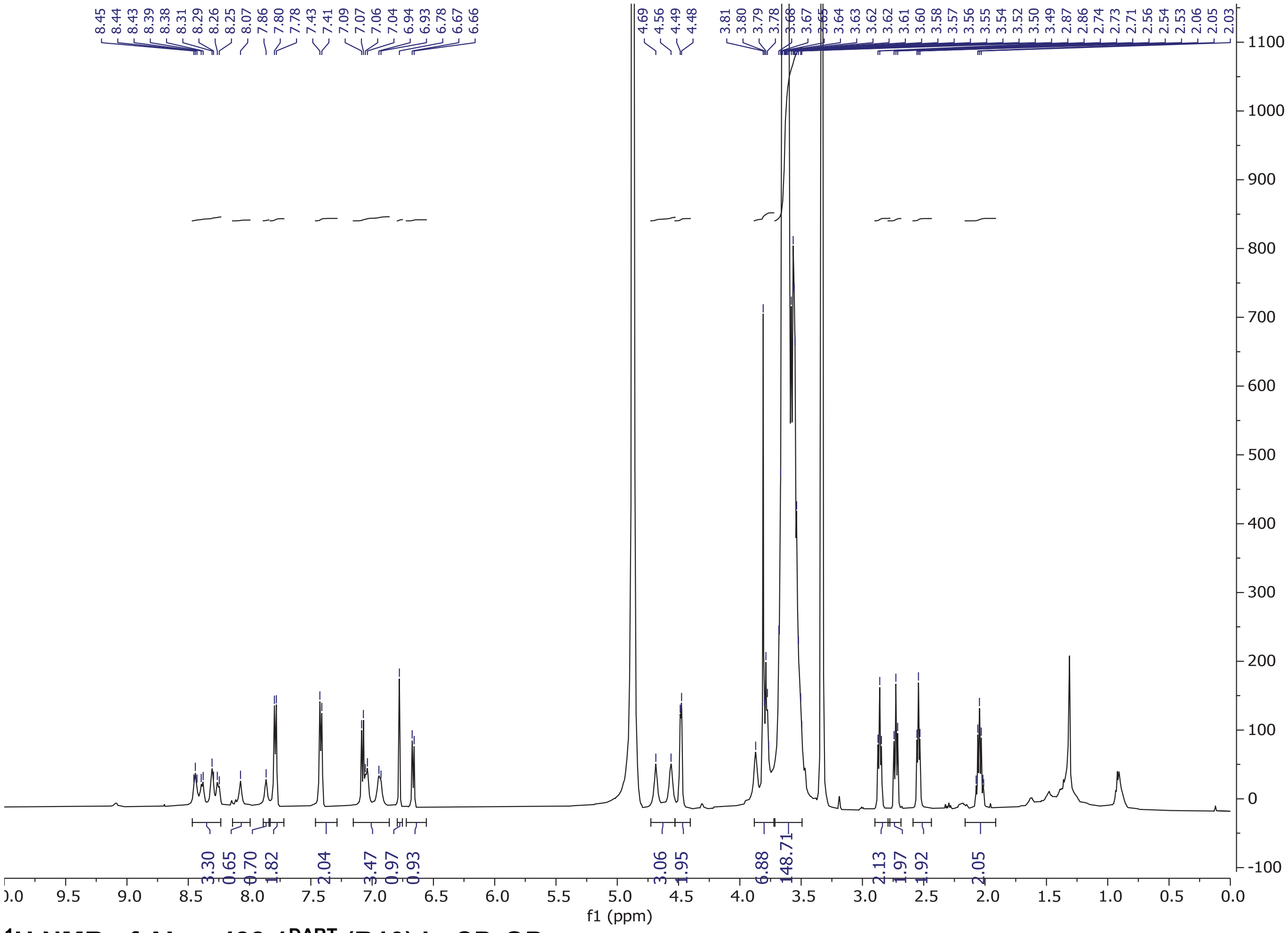

**$^{1}$H NMR of Alexa488.1$^{DART}$ (R10) in $CD_3OD$**

### Synthesis of R11–Alexa647.1$^{DART.2}$

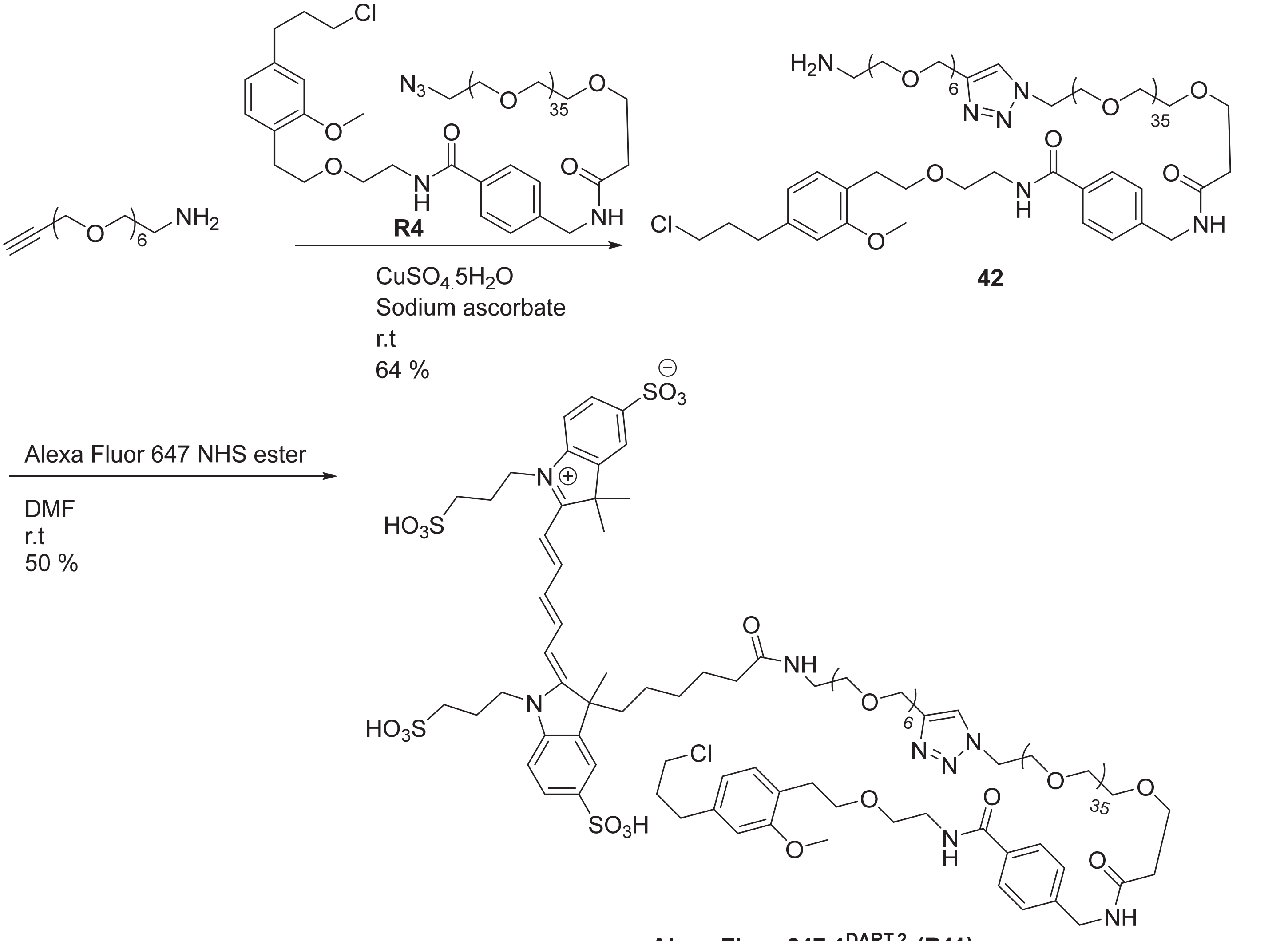


**Alexa Fluor 647.1$^{DART.2}$ (R11)**

<u>Step 1: Synthesis of 1-(4-(19-amino-2,5,8,11,14,17-hexaoxanonadecyl)-1H-1,2,3-triazol-1-yl)-N-(4-((2-(4-(3-chloropropyl)-2-methoxyphenethoxy)ethyl)carbamoyl)benzyl)-3,6,9,12,15,18,21,24,27,30,33,36,39,42,45,48,51,54,57,60,63,66,69,72,75,78,81,84,87,90,93,96,99,102,105,108-hexatriacontaoxaundecahectan-111-amide, **42**</u>

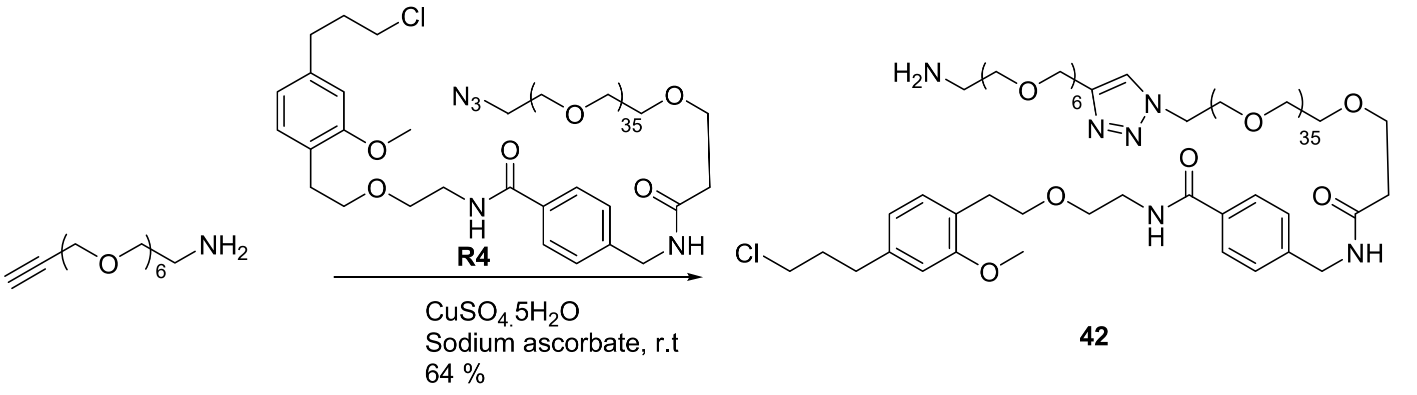


To 3,6,9,12,15,18-hexaoxahenicos-20-yn-1-amine (7.6 mg, 24 µmol) was added azido-PEG36-HTL2.0, **R4** (50 mg, 24 µmol) after which EtOH (0.4 mL)/t-BuOH(0.2 mL)/H2O(0.2 mL) was added. Then, copper (II) sulfate pentahydrate (0.3 mg, 1.2 µmol) and sodium ascorbate (0.42 mg, 2.4 µmol) were added and the reaction mixture was stirred at room temperature for 2h after which it was directly purified by reverse phase HPLC eluting with 20 to 90 % acetonitrile in water (0.1 % formic acid, 200 nm wavelength for collection and conditions as described in the general chemical synthesis section) to give 1-(4-(19-amino-2,5,8,11,14,17-hexaoxanonadecyl)-1H-1,2,3-triazol-1-yl)-N-(4-((2-(4-(3-chloropropyl)-2-methoxyphenethoxy)ethyl)carbamoyl)benzyl)-3,6,9,12,15,18,21,24,27,30,33,36,39,42,45,48,51,54,57,60,63,66,69,72,75,78,81,84,87,90,93,96,99,102,105,108-hexatriacontaoxaundecahectan-111-amide, **42** (37 mg, 64 %) as a light brown oil. $^1$H NMR (500 MHz, $D_2O$, water suppression used- the peaks close to the water peak between 4.4 and 4.6 ppm were partially suppressed) δ 7.94 (s, 1H), 7.57 – 7.39 (m, 2H), 7.37 – 7.18 (m, 2H), 6.94 (d, *J* = 7.6 Hz, 1H), 6.65 (d, *J* = 1.6 Hz, 1H), 6.43 (dd, *J* = 7.6, 1.6 Hz, 1H), 4.53 (s, 2H), 4.51 – 4.42 (m, 2H), 4.35 (d, *J* = 4.7 Hz, 2H), 3.82 (dd, *J* = 5.6, 4.5 Hz, 2H), 3.72 – 3.28 (m, 174H), 3.05 (t, *J* = 5.0 Hz, 2H), 2.68 (t, *J* = 6.2 Hz, 2H), 2.45 (td, *J* = 7.6, 6.7, 3.3 Hz, 4H), 1.79 (dq, *J* = 8.5, 6.6 Hz, 2H). (ES-LCMS) *m/z* 602.9 $(M+4H)^{4+}$.

Step 2: Synthesis of **Alexa Fluor-647.1$^{DART.2}$ (R11)**

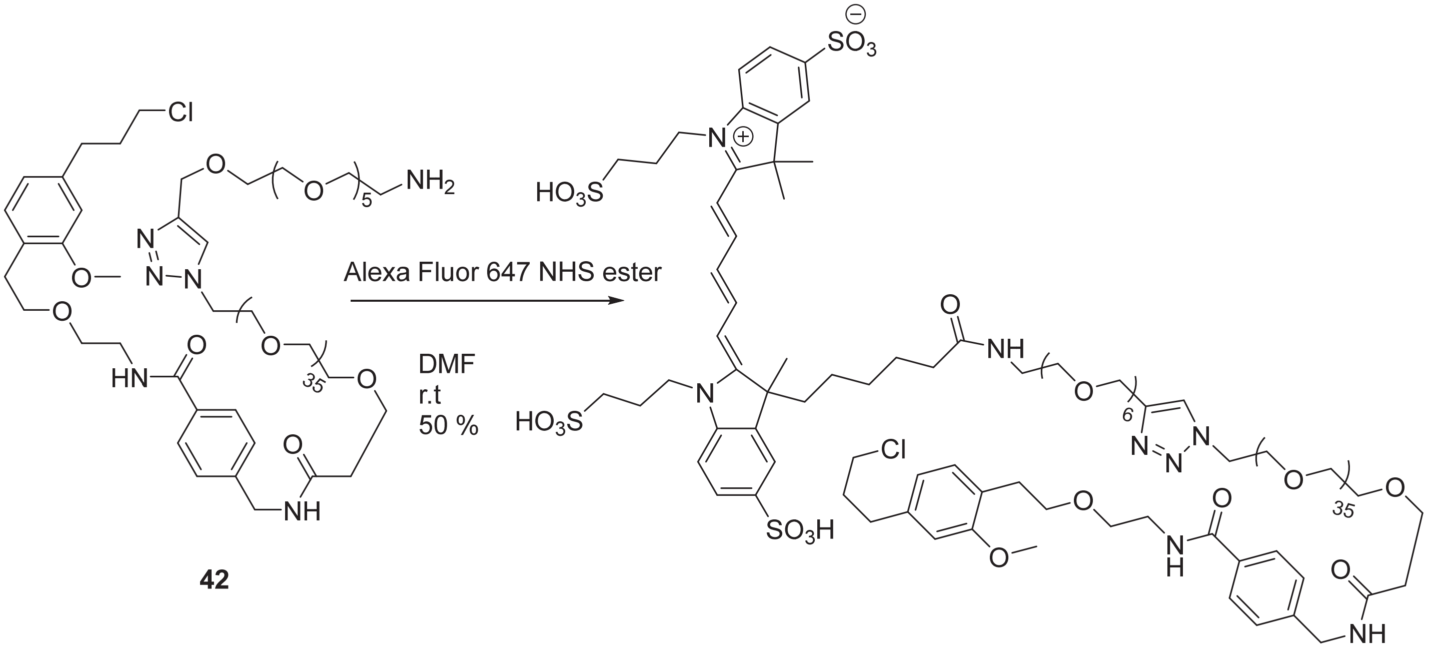


**42**

**Alexa Fluor 647.1$^{DART.2}$ (R11)**

To 1-(4-(19-amino-2,5,8,11,14,17-hexaoxanonadecyl)-1H-1,2,3-triazol-1-yl)-N-(4-((2-(4-(3-chloropropyl)-2-methoxyphenethoxy)ethyl)carbamoyl)benzyl)-3,6,9,12,15,18,21,24,27,30,33,36,39,42,45,48,51,54,57,60,63,66,69,72,75,78,81,84,87,90,93,96,99,102,105,108-hexatriacontaoxaundecahectan-111-amide**, 42** (25 mg, 10.4 µmol) was added DMF (1 mL) followed by AlexaFluor 647 NHS ester (20 mg, 20.8 µmol, ThermoFisher Scientific Catalog No. A37566). The mixture was purged with nitrogen, covered in aluminum foil and stirred overnight after which the reaction mixture was purified by reverse phase prep HPLC eluting with 30 to 60 % acetonitrile in water (0.1 % TFA, 647 nm wavelength for collection and other conditions as described in the general chemical synthesis method) in several injections and concentrated under a stream of nitrogen in the dark to give 2-((1E,3E)-5-(€-3-(1-(1-(1-(4-((2-(4-(3-chloropropyl)-2-methoxyphenethoxy)ethyl)carbamoyl)phenyl)-3-oxo-6,9,12,15,18,21,24,27,30,33,36,39,42,45,48,51,54,57,60,63,66,69,72,75,78,81,84,87,90,93,96,99,102,105,108,111-hexatriacontaoxa-2-azatridecahectan-113-yl)-1H-1,2,3-triazol-4-yl)-21-oxo-2,5,8,11,14,17-hexaoxa-20-azahexacosan-26-yl)-3-methyl-5-sulfo-1-(3-sulfopropyl)indolin-2-ylidene)penta-1,3-dien-1-yl)-3,3-dimethyl-1-(3-sulfopropyl)-3H-1l4-indole-5-sulfonic acid, **Alexa Fluor-647.1$^{DART.2}$ (R11)** (17 mg, 50 %) as a deep blue oil which was stored in a vial covered with aluminum foil. $^1$H NMR (500 MHz, $D_2O$, water suppression used- the peaks close to the water peak between 4.25 and 4.5 ppm were partially suppressed) δ 8.00 (td, *J* = 13.2, 5.4 Hz, 2H), 7.92 (s, 1H), 7.73 – 7.66 (m, 4H), 7.49 – 7.45 (m, 2H), 7.24 (dt, *J* = 8.5, 2.1 Hz, 4H), 6.89 (d, *J* = 7.6 Hz, 1H), 6.61 (d, *J* = 1.6 Hz, 1H), 6.55 (t, *J* = 12.5 Hz, 1H), 6.38 (dd, *J* = 7.6, 1.6 Hz, 1H), 6.30 (d, *J* = 13.5Hz, 1H), 6.36 (d, *J* = 13.5Hz, 1H), 4.53 (s, 2H), 4.44 (t, *J*= 5.0 Hz, 2H), 4.31 (s, 2H), 4.09 (q, *J* = 8.1 Hz, 4H), 3.77

(dd, $J$ = 5.5, 4.4 Hz, 2H), 3.65 (d, $J$ = 5.9 Hz, 2H), 3.59 – 3.39 (m, 162H), 3.34 (ddt, $J$ = 10.4, 6.9, 3.2 Hz, 6H), 3.10 (t, $J$ = 5.3 Hz, 2H), 2.87 (dt, $J$ = 10.9, 7.1 Hz, 4H), 2.63 (t, $J$ = 6.3 Hz, 2H), 2.47 – 2.38 (m, 4H), 2.18 (t, $J$ = 12.3 Hz, 1H), 2.07 (ddt, $J$ = 14.2, 10.5, 6.0 Hz, 5H), 1.89 (t, $J$ = 7.3 Hz, 2H), 1.75 (dq, $J$ = 8.5, 6.5 Hz, 2H), 1.54 – 1.49 (m, 9H), 1.22 (q, $J$ = 7.4 Hz, 2H), 0.95 (p, $J$ = 7.5 Hz, 2H), 0.67 (d, $J$ = 11.6 Hz, 1H), 0.41 (dd, $J$ = 12.3, 6.9 Hz, 1H). Note: the structure of AlexaFluor 647 NHS ester was assumed to be consistent with that reported in reference 7. (ES-LCMS) $m/z$ 813.5 $(M+4H)^{4+}$; Compound decomposed on HRMS

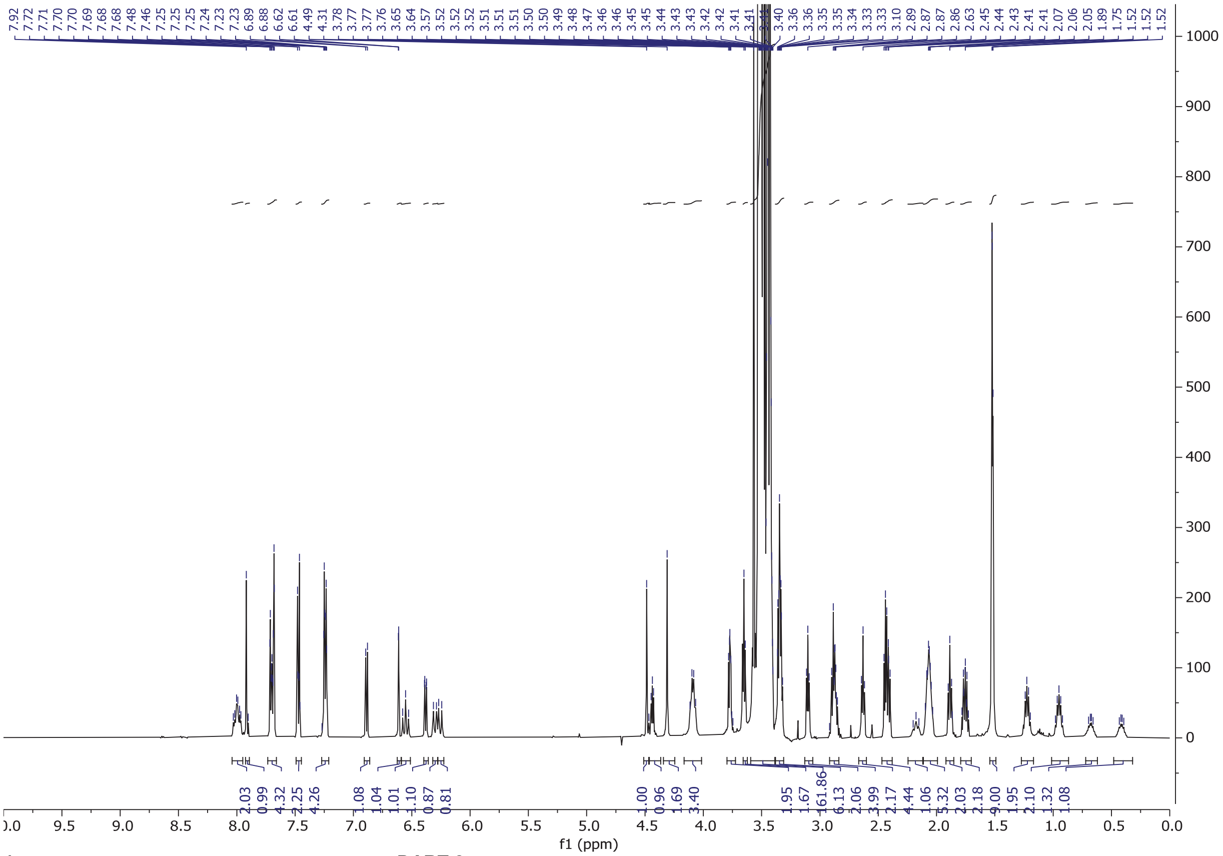

**$^1$H NMR of Alexa Fluor-647.1$^{DART.2}$ (R11) in $D_2O$ (water suppression)**

### Synthesis of R12–blank.1$^{DART.2}$

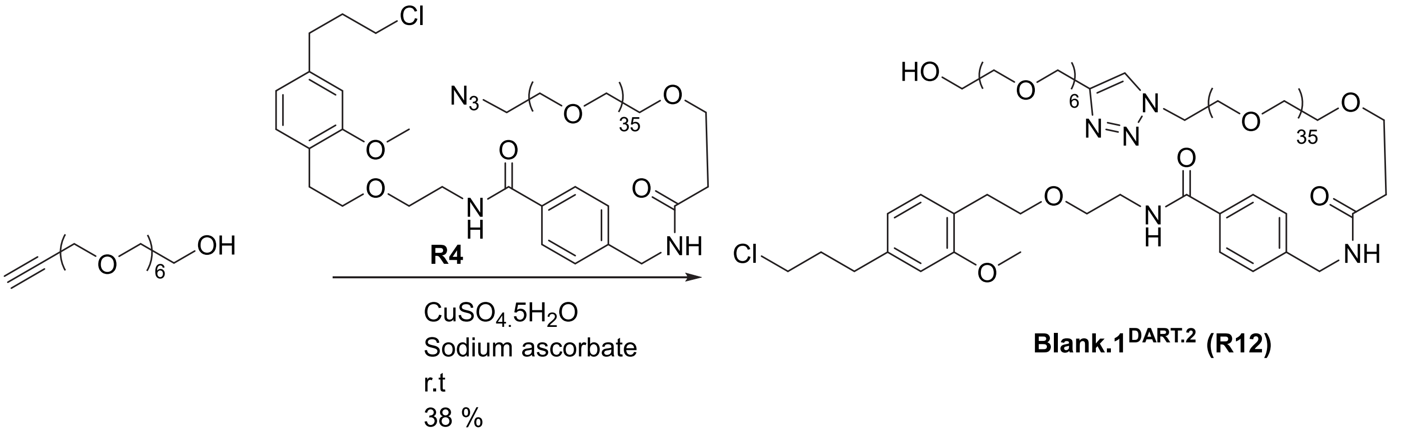


**Blank.1$^{DART.2}$ (R12)**

To 3,6,9,12,15,18-hexaoxahenicos-20-yn-1-ol (4.0 mg, 12.5 µmol) was added EtOH:t-BuOH:water (2:1:1, 0.8 mL) followed by azido-PEG36-HTL2.0, **R4** (26.1 mg, 12.5 µmol). Then, copper (II) sulfate pentahydrate (0.16 mg, 0.62 µmol) and sodium ascorbate (0.2 mg, 1.25 µmol) were added and the reaction was stirred overnight at room temperature after which it was purified by reverse phase prep HPLC eluting with 25 to 90 % acetonitrile in water (0.1 % formic acid, 220 nm wavelength for collection and the parameters as described in the general chemical synthesis section). The fraction was lyophilized to give **Blank.1$^{DART.2}$, R12** (11.4 mg, 38 %). $^1$H NMR (500 MHz, $CDCl_3$) δ 7.73 (br.s, 1H), 7.69-7.66 (m, 2H), 7.34(d, *J*= 7.9Hz, 2H), 7.09 (br.s, 1H), 7.03(d, *J*= 7.8Hz, 1H), 6.71 – 6.61 (m, 2H), 6.51 (br, 1H), 4.66 (s, 2H), 4.52 (t, *J* = 5.0 Hz, 2H), 4.48 (dd, *J* = 5.0 Hz, 2H), 3.86 (t, *J* = 4.9 Hz, 2H), 3.79 – 3.47 (m, 170H), 2.88 (t, *J*= 7.0Hz, 2H), 2.71 (t, *J* = 7.4 Hz, 2H), 2.53 (t, *J* = 5.6 Hz, 2H), 2.03 (dq, *J* = 8.3, 6.6 Hz, 2H). HRMS (ESI$^+$): *m/z* calcd for $C_{112}H_{205}ClN_5NaO_{47}$: 1215.6707 $[M+Na+H]^{2+}$; found: 1215.6749, $[M+Na+H]^{2+}$

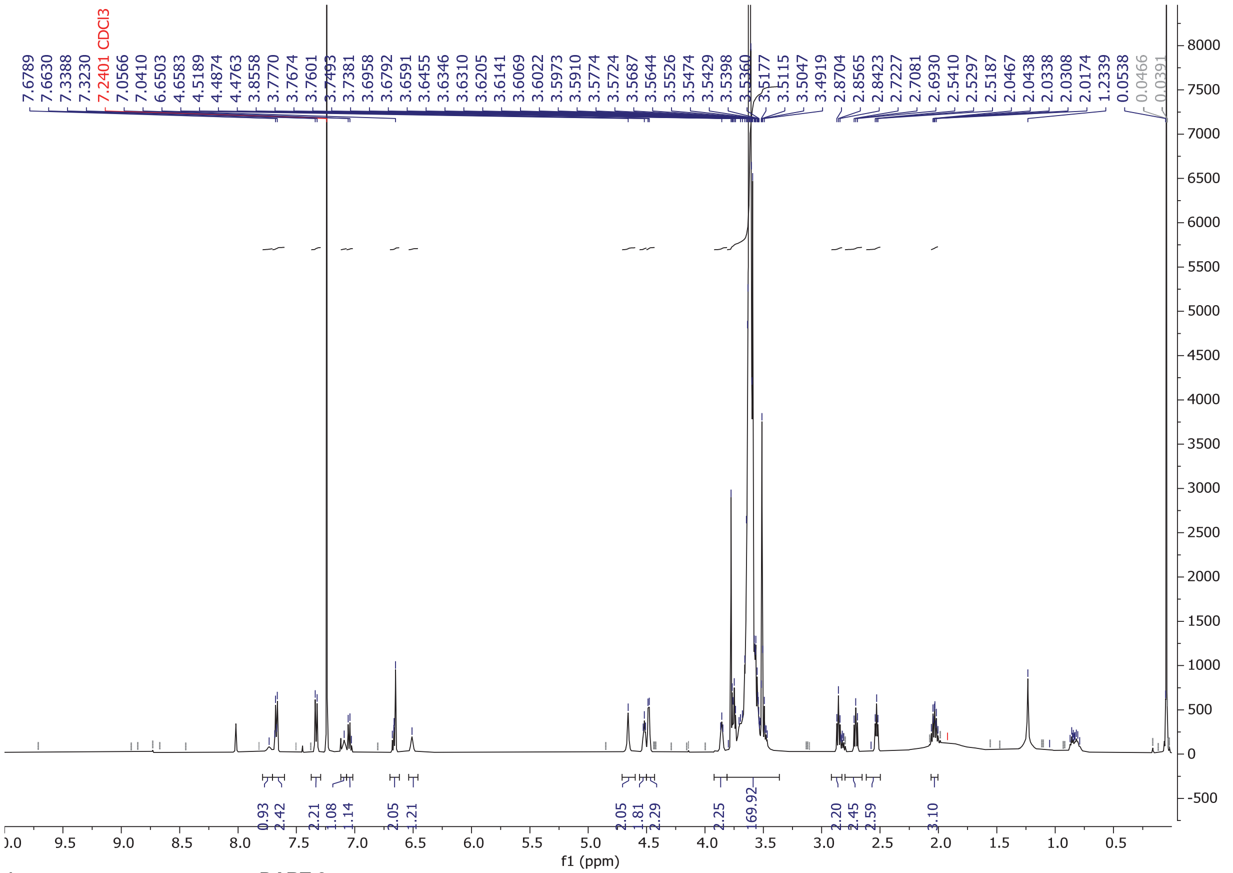


**$^1$H NMR of Blank.1$^{DART.2}$ (R12) in $CDCl_3$**

### Synthesis of R13–diazepam.1$^{DART.2}$

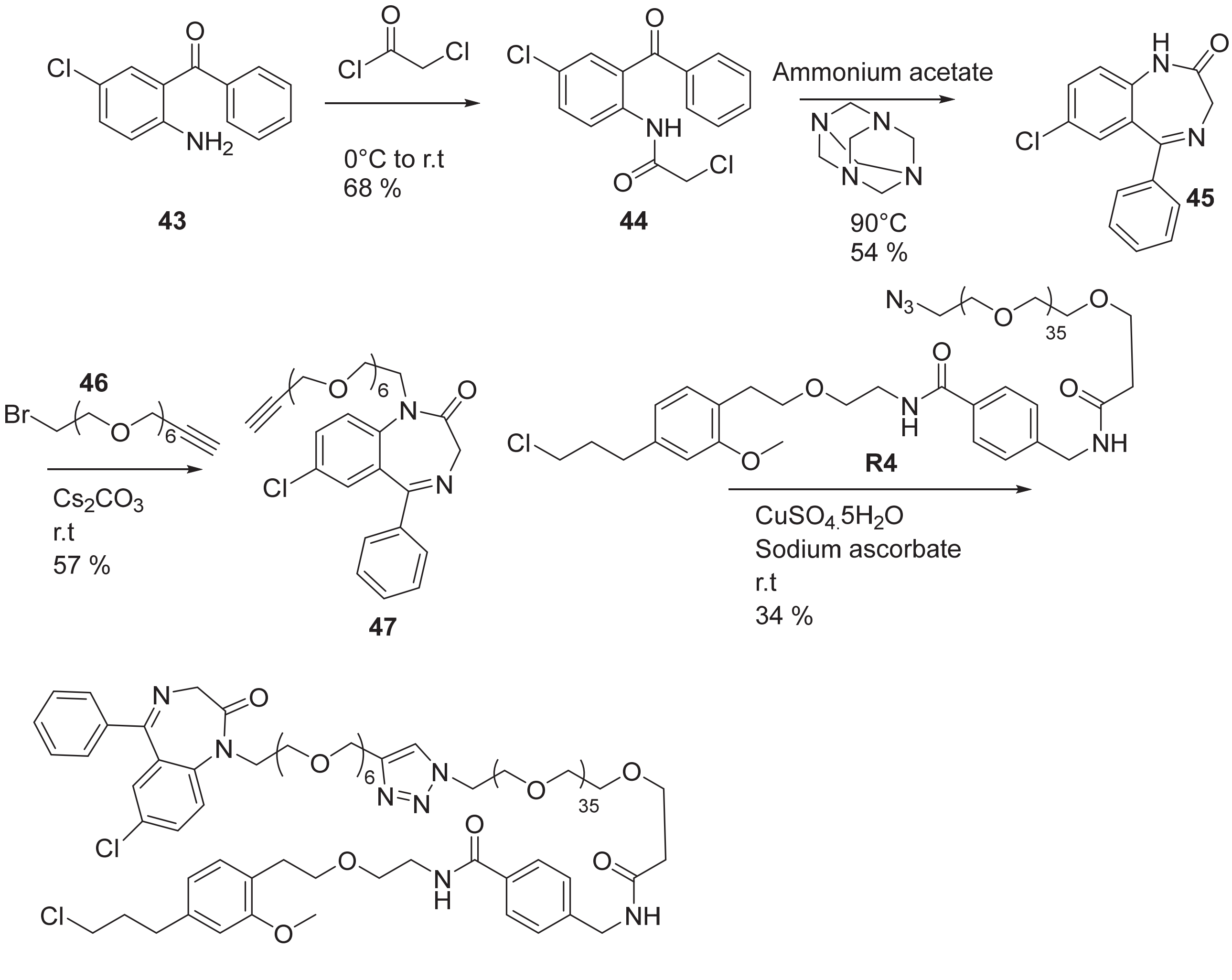


**Diazepam.1$^{DART.2}$ (R13)**

Synthesis of 1-bromo-3,6,9,12,15,18-hexaoxahenicos-20-yne, **46**

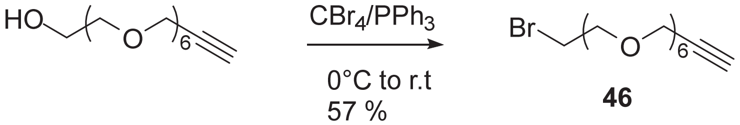


To 3,6,9,12,15,18-hexaoxahenicos-20-yn-1-ol (1.00 g, 1 equiv., 3.12 mmol) in THF (20 mL) in a cooled ice/water bath was added $CBr_4$ (1.55 g, 1.50 equiv., 4.68 mmol) and triphenylphosphine (1.23 g, 1.50 equiv., 4.68 mmol) and the reaction mixture was stirred as the ice bath expired for 4 h after which hexanes (15 mL) were added. The mixture was filtered and the solids were discarded. The filtrate was concentrated, triturated in a mixture of hexanes/THF (~ 1:1), filtered and concentrated again to remove more triphenylphosphine oxide. The yellow oil was purified by silica gel chromatography eluting with 0 to 100 % ethyl acetate in hexanes to give 1-bromo-3,6,9,12,15,18-hexaoxahenicos-20-yne, **46** (688 mg, 1.80 mmol, 57.5 %) as a yellow oil. $^1$H NMR (500 MHz, $CDCl_3$, contains a small amount of $PPh_3$=O) δ 4.19 (d, *J* = 2.4 Hz, 2H), 3.79 (t, *J* = 6.4 Hz, 2H), 3.70 – 3.59 (m, 20H), 3.45 (t, *J* = 6.4 Hz, 2H), 2.41 (t, *J* = 2.4 Hz, 1H). (ES-LCMS) *m/z* 385.3 (M+H)$^+$.

Step 1: Synthesis of N-(2-benzoyl-4-chlorophenyl)-2-chloroacetamide, **44** (reported in reference 5)

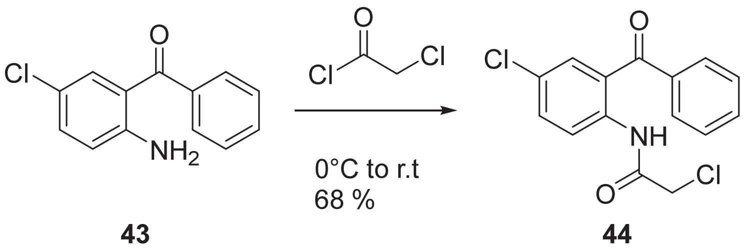

To a cooled (ice/water bath) solution of (2-amino-5-chlorophenyl)(phenyl)methanone, **43** (1 g, 4.32 mmol) in dichloromethane (14.39 mL) under nitrogen flow was added triethylamine (0.902 ml, 6.47 mmol) and DMAP (5.27 mg, 0.043 mmol) after which chloroacetyl chloride (0.380 ml, 4.75 mmol) was added dropwise. Additional chloroacetyl chloride (0.2 ml, 2.497 mmol) was added and the ice bath was removed. The reaction was stirred for another 2 h at room temperature after which the reaction was filtered to remove solids, which were washed with DCM. The organic layer was washed with saturated $NaHCO_3$ solution (2 x 5 mL), water (5 mL), brine (5 mL), and filtered through an isolute phase separator. The filtrate was concentrated to a brown solid which was crystallized from EtOH to give N-(2-benzoyl-4-chlorophenyl)-2-chloroacetamide, **44** (0.9 g, 2.92 mmol, 67.7 % yield) as a light brown/light yellow solid. $^1H$ NMR (500 MHz, $CDCl_3$) δ 11.44 (s, 1H), 8.61 – 8.55 (m, 1H), 7.76 – 7.67 (m, 2H), 7.66 – 7.58 (m, 1H), 7.58 – 7.46 (m, 4H), 4.17 (s, 2H). (ES-LCMS) *m*/*z* 310.1 $(M+H)^+$.

Step 2: Synthesis of 7-chloro-5-phenyl-1H-benzo[e][1,4]diazepin-2(3H)-one, **45** (reported in references 5 and 6)

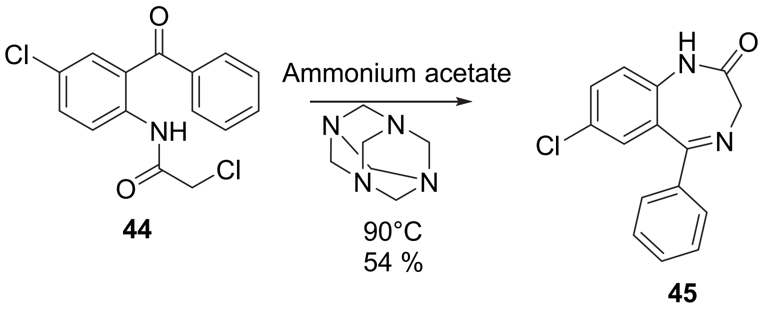

To N-(2-benzoyl-4-chlorophenyl)-2-chloroacetamide, **44** (0.9 g, 2.92 mmol) in ethanol (58.4 mL) was added ammonium acetate (0.45 g, 5.84 mmol) and hexamethylenetetramine (0.82 g, 5.84 mmol) after which the reaction was heated to 90°C for 1 h. Additional ammonium acetate (120 mg, 1.56 mmol) and hexamethylenetetramine (240 mg, 1.7 mmol) were added and the reaction was heated at 75°C overnight after which the reaction was diluted with water (240 mL). The suspension was extracted with ethyl acetate (2 x 100 mL), brine (30 mL), and filtered through an isolute phase separator after which the filtrate was concentrated. The residue was triturated in hexanes/acetone and filtered to give 7-chloro-5-phenyl-1H-benzo[e][1,4]diazepin-2(3H)-one, **45** (0.43 g, 1.588 mmol, 54.4 % yield) as a tan solid. $^1H$ NMR (500 MHz, Chloroform-*d*) δ 9.05 (s, 1H), 7.54 – 7.49 (m, 2H), 7.49 – 7.35 (m, 4H), 7.29 (d, *J* = 2.4 Hz, 1H), 7.10 (d, *J* = 8.7 Hz, 1H), 4.57 – 4.02 (m, 2H). (ES-LCMS) *m*/*z* 271.2 $(M+H)^+$.

Step 3: Synthesis of 7-chloro-1-(3,6,9,12,15,18-hexaoxahenicos-20-yn-1-yl)-5-phenyl-1H-benzo[e][1,4]diazepin-2(3H)-one

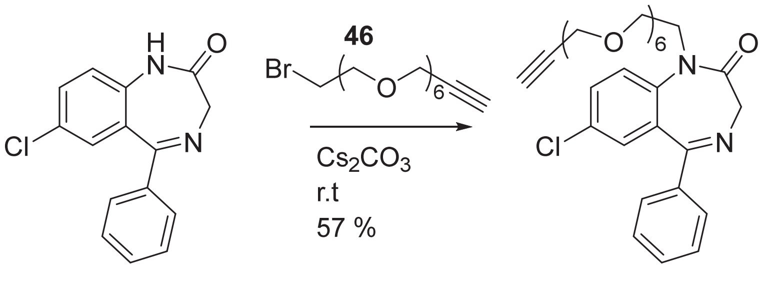


To a vial was added 7-chloro-5-phenyl-1,3-dihydro-2H-benzo[e][1,4]diazepin-2-one, **45** (100 mg, 1 equiv., 369 µmol) and DMF (3 mL) after which 1-bromo-3,6,9,12,15,18-hexaoxahenicos-20-yne, **46** (142 mg, 1 equiv., 369 µmol) was added followed by cesium carbonate (181 mg, 1.5 Eq, 554 µmol) and the reaction mixture was allowed to stir overnight. The reaction mixture was diluted with water (5 mL) and ethyl acetate (20 mL). The aqueous layer was extracted once with ethyl acetate (20 mL). Combined organic layers were washed with brine, filtered through an isolute phase separator, concentrated and purified by reverse phase prep HPLC eluting with 20 to 80 % acetonitrile in water (0.1 % formic acid and other conditions as reported in the general chemical synthesis section) to give 7-chloro-1-(3,6,9,12,15,18-hexaoxahenicos-20-yn-1-yl)-5-phenyl-1,3-dihydro-2H-benzo[e][1,4]diazepin-2-one, **47** (121 mg, 211 µmol, 57.2 %) as a yellow oil. $^1$H NMR (500 MHz, $CDCl_3$) δ 7.63 (d, *J* = 8.8 Hz, 1H), 7.61 – 7.57 (m, 2H), 7.49 – 7.43 (m, 2H), 7.42 – 7.36 (m, 2H), 7.22 (d, *J* = 2.5 Hz, 1H), 4.77 (d, *J* = 10.4 Hz, 1H), 4.17 (d, *J* = 2.4 Hz, 2H), 4.16 – 4.09 (m, 1H), 3.89 (ddd, *J* = 14.3, 5.4, 4.0 Hz, 1H), 3.79 – 3.71 (m, 2H), 3.70 – 3.38 (m, 21H), 2.40 (t, *J* = 2.4 Hz, 1H). (ES-LCMS) *m/z* 573.4 $(M+H)^+$.

Step 4: Synthesis of **Diazepam.1$^{DART.2}$ (R13)**

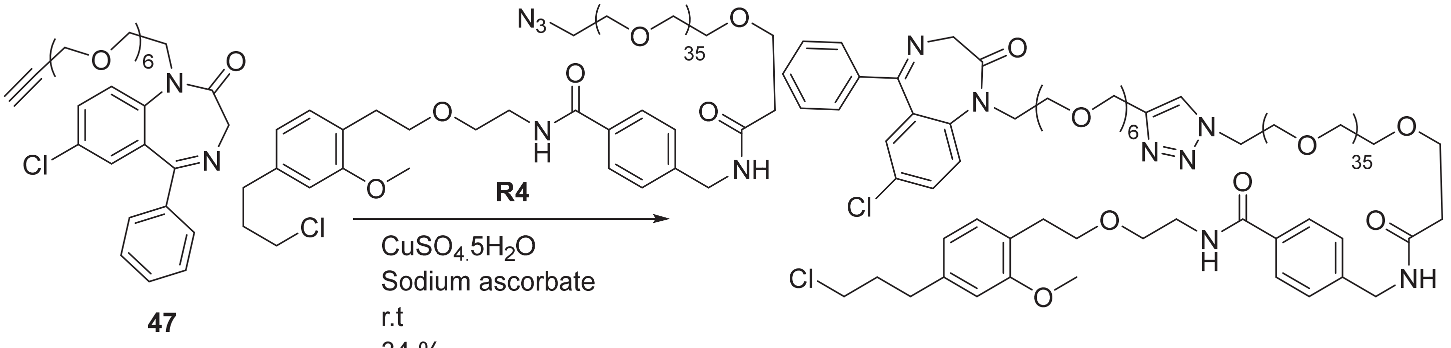


To -chloro-1-(3,6,9,12,15,18-hexaoxahenicos-20-yn-1-yl)-5-phenyl-1,3-dihydro-2H-benzo[e][1,4]diazepin-2-one (5 mg, 8.7 µmol), **47** was added a solution azido-PEG36-HTL2.0, **R4** (18 mg, 8.7 µmol) in EtOH (0.4 mL)/t-BuOH (0.2 mL)/Water (0.2 mL). Then, copper (II) sulfate pentahydrate (0.1 mg, 0.44 µmol) and sodium ascorbate (0.15 mg, 0.87 µmol) were added and the reaction mixture was stirred overnight after which it was purified by reverse phase prep HPLC eluting with 5 to 95 % acetonitrile in water (0.1 % formic acid, 220 nm wavelength for collection and other conditions as reported in the general chemical synthesis methods section) to give 1-(4-(19-(7-chloro-2-oxo-5-phenyl-2,3-dihydro-1H-benzo[e][1,4]diazepin-1-yl)-2,5,8,11,14,17-hexaoxanonadecyl)-1H-1,2,3-triazol-1-yl)-N-(4-((2-(4-(3-chloropropyl)-2-methoxyphenethoxy)ethyl)carbamoyl)benzyl)-3,6,9,12,15,18,21,24,27,30,33,36,39,42,45,48,51,54,57,60,63,66,69,72,75,78,81,84,87,90,93,96,99,102,105,108-hexatriacontaoxaundecahectan-111-amide, **Diazepam.1$^{DART.2}$, R13** (8 mg, 34 %). $^1$H NMR (500 MHz, $CDCl_3$) δ 7.71 (s, 1H), 7.69-7.66 (m, 2H), 7.63 (d, *J* = 8.9 Hz, 1H), 7.60 – 7.57 (m, 2H), 7.48 – 7.44 (m, 2H), 7.41-7.37 (m, 2H), 7.33 (d, *J* = 8.2 Hz, 2H), 7.22 (d, *J* = 2.5 Hz, 1H), 7.10 (t, *J* = 6.4 Hz, 1H), 7.05 (d, *J* = 7.8

Hz, 1H), 6.66 (d, *J* = 6.4 Hz, 2H), 6.52 (br.s, 1H), 4.77 (d, *J* = 10.5 Hz, 1H), 4.65 (s, 2H), 4.51 (t, *J*=5.1 Hz, 2H), 4.48 (d, J= 6.0Hz, 2H), 4.16 – 4.12 (m, 1H), 3.91 – 3.42 (m, 184H), 2.85 (t, *J* = 7.0 Hz, 2H), 2.71 (t, *J* = 7.4 Hz, 2H), 2.53 (t, *J* = 5.6 Hz, 2H), 2.03 (dq, *J* = 8.2, 6.6 Hz, 2H).
HRMS (ESI+): m/z calcd for $C_{127}H_{214}Cl_2N_7NaO_{47}$: 1341.6934 $[M+Na+H]^{2+}$; found: 1341.6982 $[M+Na+H]^{2+}$

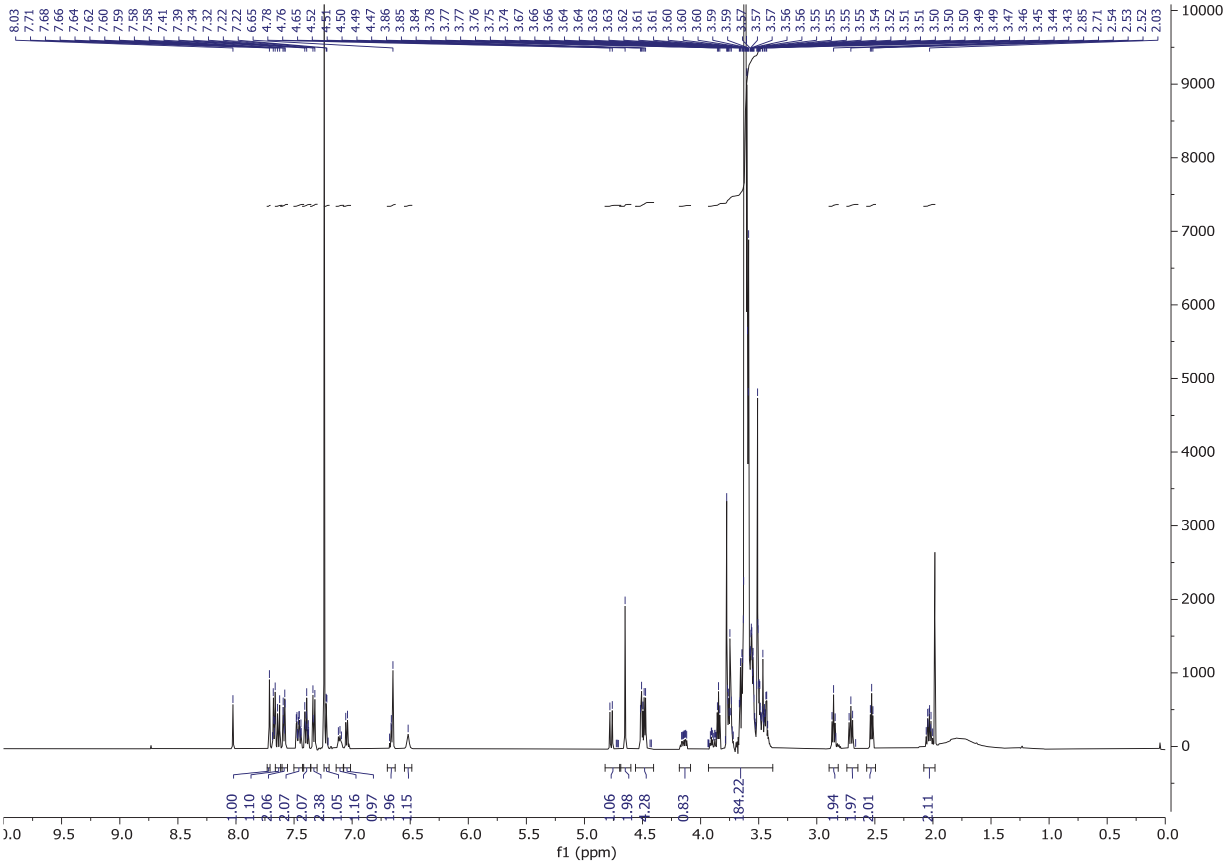

**$^1$H NMR of Diazepam.1$^{DART.2}$ (R13) in $CDCl_3$**

### Synthesis of R14–flumazenil.1DART.2

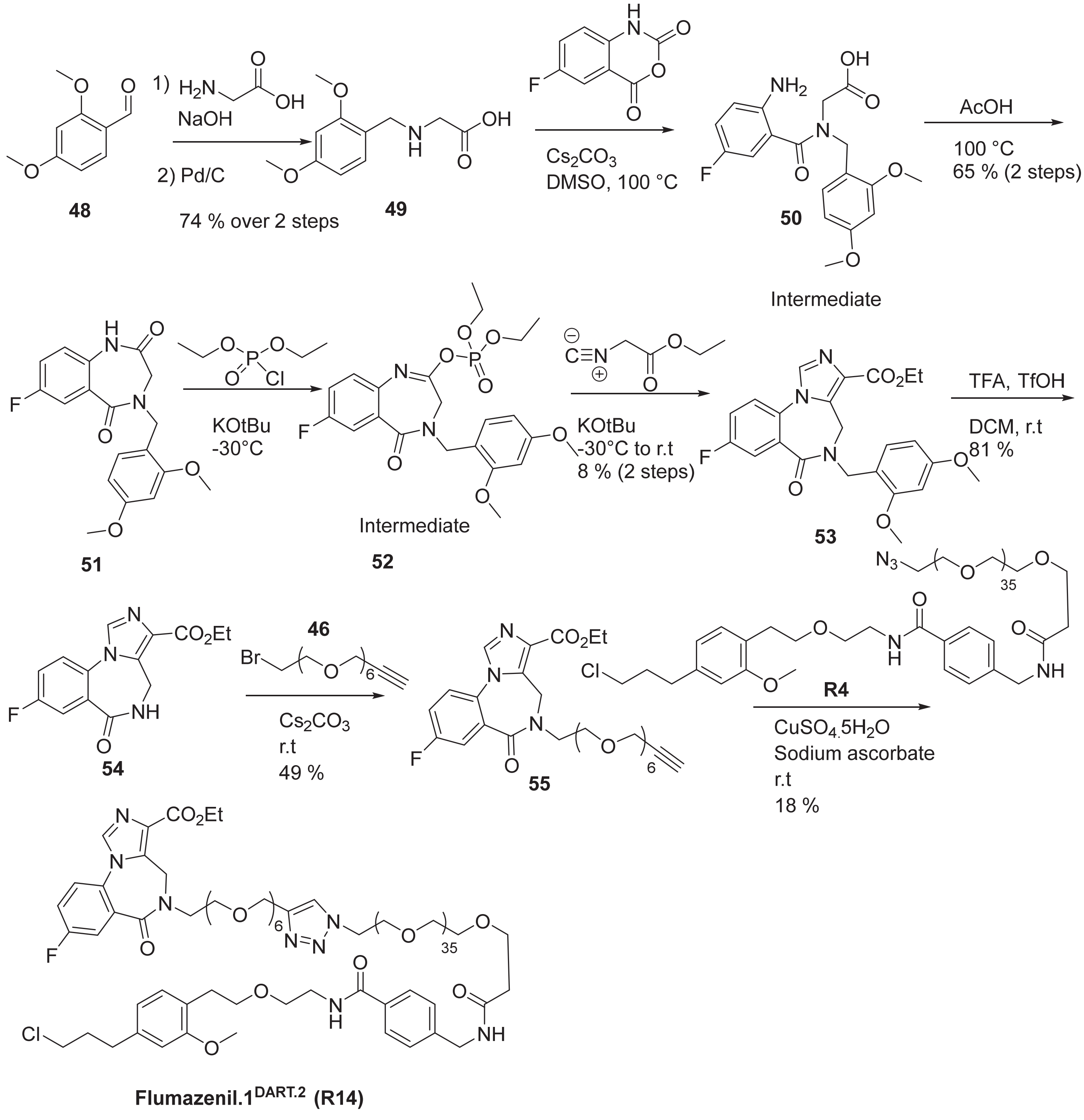


**Flumazenil.1DART.2 (R14)**

Step 1: Synthesis of 2-((2,4-dimethoxybenzyl)amino)acetic acid, **49**

See reference 8 for the reported synthesis until the debenzylated analog.

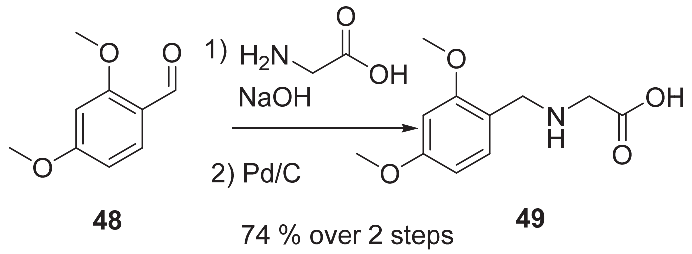


To a round bottom flask was added glycine (2 g, 26.6 mmol) to which 1M NaOH (32 mL) was added. In a separate vial was added 2,4-dimethoxybenzaldehyde, **48** (3.98 g, 23.98 mmol) in methanol (16 mL) and this stirred suspension was added portionwise to the round bottom flask containing glycine/NaOH. A white solid precipitated and the suspension was stirred for 10 min at room temperature after which Pd/C (0.8 g, 0.752 mmol) was added under a stream of nitrogen. The reaction was stirred overnight under an atmosphere of hydrogen after which it was filtered through a pad of celite, washing with hot methanol. The filtrate was concentrated on the rotary evaporator to give a yellow aqueous solution which was cooled in an ice bath. The mixture was acidified (3 M HCl) until pH = 4. The mixture was concentrated to remove water, taken up in methanol and filtered. The filtrate was concentrated then triturated from acetone to give 2-((2,4-dimethoxybenzyl)amino)acetic acid, **49** (4.45 g, 19.76 mmol, 74.2 % yield) as a white solid. $^1$H NMR (500 MHz, DMSO-$d_6$) δ 13.71 (br.s, 1H), 9.26 (br.s, 2H), 7.36 (d, *J* = 8.4 Hz, 1H), 6.62 (d, *J* = 2.4 Hz, 1H), 6.57 (dd, *J* = 8.4, 2.4 Hz, 1H), 4.08 (s, 2H), 3.82 (s, 3H), 3.79 (s, 3H), 3.72 (s, 2H). (ES-LCMS) *m*/*z* 226.1 (M+H)$^+$.

Step 2/3: Synthesis of 4-(2,4-dimethoxybenzyl)-7-fluoro-3,4-dihydro-1H-benzo[e][1,4]diazepine-2,5-dione, **51**

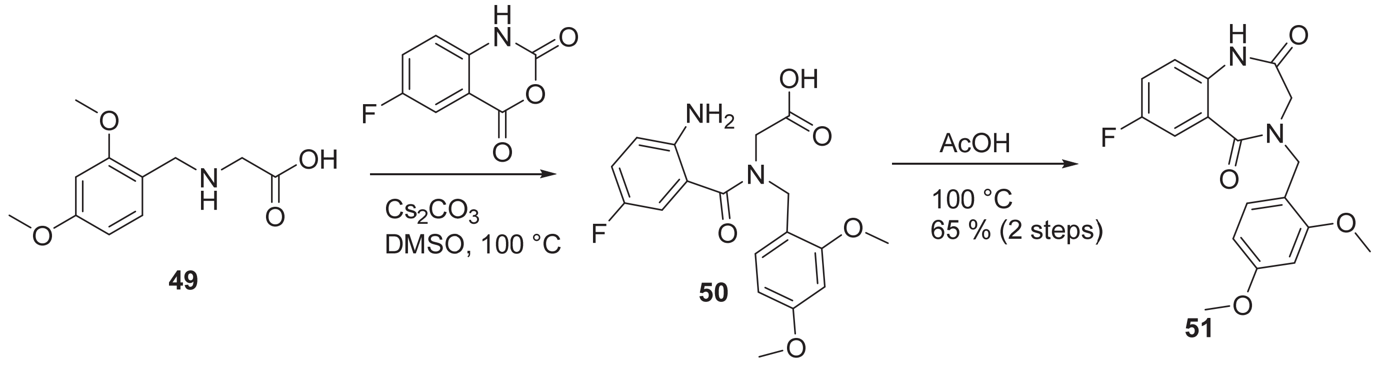


To a vial was added 2-((2,4-dimethoxybenzyl)amino)acetic acid, **49** (2 g, 8.88 mmol) to which DMSO (11 mL) was added. Then, 6-fluoro-1H-benzo[d][1,3]oxazine-2,4-dione (1.59 g, 8.78 mmol) was added and the mixture was stirred until all solids dissolved. Cesium carbonate (2.89 g, 8.88 mmol) was added and the reaction mixture was heated at 100 °C for 10 min after the intermediate **50** was observed on LCMS. Pressure buildup was observed during this reaction. Acetic acid (2.033 ml, 35.5 mmol) was added and the reaction mixture was heated at 100°C overnight. To this mixture was added water and the suspension was thoroughly stirred, then filtered. The collected tan solid was washed generously with water and dried overnight to give 4-(2,4-dimethoxybenzyl)-7-fluoro-3,4-dihydro-1H-benzo[e][1,4]diazepine-2,5-dione, **51** (2 g, 5.81 mmol, 65.4 % yield) as a tan solid. $^1$H NMR (500 MHz, Chloroform-d) δ 8.38 (s, 1H), 7.67 (dd, J = 9.0, 3.1 Hz, 1H), 7.27 (d, J = 8.1 Hz, 1H), 7.15 (ddd, J = 8.8, 7.3, 3.1 Hz, 1H), 6.90 (dd, J = 8.8, 4.6 Hz, 1H), 6.44 (d, J = 7.6 Hz, 2H), 4.77 (s, 2H), 3.88 (s, 2H), 3.80 (s, 3H), 3.77 (s, 3H). (ES-LCMS) *m*/*z* 345.0 (M+H)$^+$.

Step 4/5: Synthesis of ethyl 5-(2,4-dimethoxybenzyl)-8-fluoro-6-oxo-5,6-dihydro-4H-benzo[f]imidazo[1,5-a][1,4]diazepine-3-carboxylate, **53**

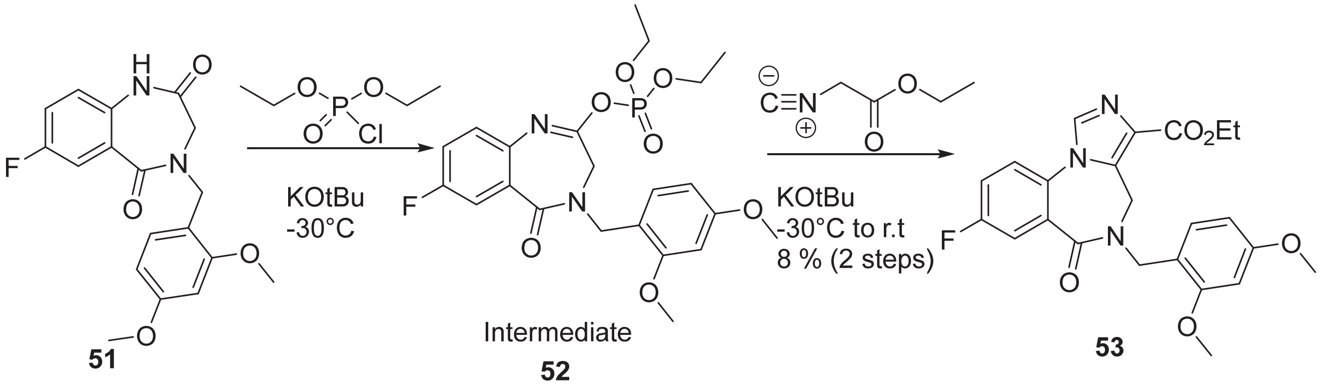

4-(2,4-dimethoxybenzyl)-7-fluoro-3,4-dihydro-1H-benzo[e][1,4]diazepine-2,5-dione, **51** (2 g, 5.81 mmol) was added to a vial to which dry DMF (8 mL) was added followed by potassium tert-butoxide (0.782 g, 6.97 mmol). After stirring for 15 min under nitrogen, the reaction was placed in an acetonitrile/dry ice bath at -30 °C and diethyl chlorophosphate (0.919 ml, 6.39 mmol) was added dropwise over 2 minutes. Upon addition, the reaction was stirred for 20 min and maintained in the cold bath during this time.

In a separate vial, potassium tert-butoxide (0.782 g, 6.97 mmol) was dissolved in anhydrous DMF (3 mL) and after stirring for 5 min at room temperature, this solution was cooled in an acetonitrile/dry ice bath at -50 °C and placed under nitrogen after which ethyl 2-isocyanoacetate (0.698 ml, 6.39 mmol) in DMF (1 mL) was added dropwise. This dark brown solution was immediately added dropwise over 5 min to the solution (which at this point has warmed to -20 °C) containing the intermediate described in the previous paragraph. The resulting mixture was stirred at room temperature for 2 hours after which it was quenched with acetic acid (0.499 mL, 8.71 mmol). Saturated aqueous $NH_4Cl$ solution (5 mL) was added. The reaction mixture was diluted with ethyl acetate (40 mL) and the aqueous layer was extracted with EtOAc (2x30 mL). Combined organic extracts were washed with brine and dried (sodium sulfate). The organic layer was decanted off and concentrated to give a black oil (~ 3 g) with a stench odor.

The mixture was purified by silica gel chromatography eluting with 5 to 50 % ethylacetate in dichloromethane to give 600 mg of a slightly unclean product. This solid was recrystallized from ethyl acetate to give desired product, **53** as a white powder (200 mg, 8 %). $^1$H NMR (500 MHz, Chloroform-d) δ 7.86 - 7.75 (m, 2H), 7.38 (dd, J = 8.8, 4.5 Hz, 1H), 7.31 (ddd, J = 8.8, 7.1, 3.0 Hz, 1H), 7.19 (d, J = 8.2 Hz, 1H), 6.52 - 6.28 (m, 2H), 5.36 (d, J = 15.9 Hz, 1H), 5.00 - 4.55 (m, 2H), 4.31 (q, J = 7.1 Hz, 2H), 4.15 (d, J = 15.9 Hz, 1H), 3.77 (d, J = 17.1 Hz, 6H), 1.37 (t, J = 7.1 Hz, 3H). (ES-LCMS) *m/z* 440.3 $(M+H)^+$.

Step 6: Synthesis of ethyl 8-fluoro-6-oxo-5,6-dihydro-4H-benzo[f]imidazo[1,5-a][1,4]diazepine-3-carboxylate, **54**

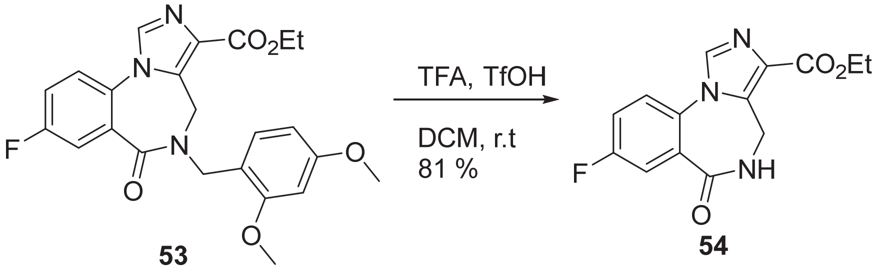

To a vial containing ethyl 5-(2,4-dimethoxybenzyl)-8-fluoro-6-oxo-5,6-dihydro-4H-benzo[f]imidazo[1,5-a][1,4]diazepine-3-carboxylate, **53** (0.15 g, 0.341 mmol) was added DCM (1 mL) and the mixture was placed in an ice/water bath after which TFA (1 mL, 12.98 mmol) and trifluoromethanesulfonic acid (69 uL, 0.777 mmol) were added. The reaction mixture turned from light pink to dark purple over the course of 90 minutes. The reaction mixture was concentrated and diluted with DCM/saturated aqueous sodium bicarbonate solution. The solution was stirred and the color changed from purple to a very light pink, and the aqueous layer was slightly basic (pH=8). The aqueous layer was extracted with DCM and all combined organic extracts were washed with

brine, filtered through an isolute phase separator, and concentrated to give a white solid which was triturated from hot ethyl
acetate and diethyl ether to give ethyl 8-fluoro-6-oxo-5,6-dihydro-4H-benzo[f]imidazo[1,5-a][1,4]diazepine-3-carboxylate, **54** (80 mg ,0.277 mmol, 81 % yield) as a white solid.
$^{1}$H NMR (500 MHz, $CDCl_3$) δ 7.85 – 7.77 (m, 2H), 7.44 (dd, *J* = 8.8, 4.5 Hz, 1H), 7.37 (ddd, *J* = 8.9, 7.0, 2.9 Hz, 1H), 6.52 (t, *J* = 6.3 Hz, 1H), 4.41 (q, *J* = 7.1 Hz, 2H), 3.84 – 3.63 (m, 1H), 1.42 (t, *J* = 7.1 Hz, 3H). (ES-LCMS) *m*/*z* 290.1 $(M+H)^+$.

Step 7: Synthesis of ethyl 8-fluoro-5-(3,6,9,12,15,18-hexaoxahenicos-20-yn-1-yl)-6-oxo-5,6-dihydro-4H-benzo[f]imidazo[1,5-a][1,4]diazepine-3-carboxylate, **55**

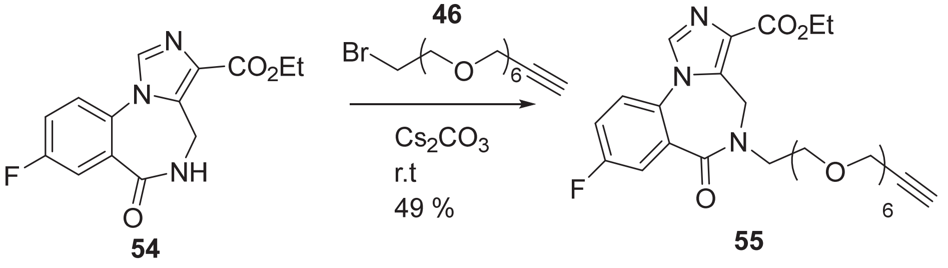


In a vial was added ethyl 8-fluoro-6-oxo-5,6-dihydro-4H-benzo[f]imidazo[1,5-a][1,4]diazepine-3-carboxylate, **54** (10 mg, 0.035 mmol) followed by anhydrous DMF (700 uL) and 1-bromo-3,6,9,12,15,18-hexaoxahenicos-20-yne, **46** (13.25 mg, 0.035 mmol) followed by anhydrous cesium carbonate (22.53 mg, 0.069 mmol) and the reaction was stirred at room temperature overnight after which additional alkyl bromide (5 mg) was added. After stirring for several hours, no change was detected by LCMS (still some SM remaining).

The reaction was diluted with ethyl acetate (20 mL) and water (5 mL). The organic layer was washed with brine (5 mL), filtered through an isolute phase separator, concentrated and purified by RP HPLC eluting with 5 to 70 % acetonitrile in water (0.1 % FA and the rest of conditions as described in the general chemistry synthesis section) to give ethyl 8-fluoro-5-(3,6,9,12,15,18-hexaoxahenicos-20-yn-1-yl)-6-oxo-5,6-dihydro-4H-benzo[f]imidazo[1,5-a][1,4]diazepine-3-carboxylate, **55** (10 mg , 0.017 mmol, 48.9 % yield) as a brown oil.
$^{1}$H NMR (500 MHz, $CDCl_3$) δ 7.85 (s, 1H), 7.77 (d, *J* = 8.8 Hz, 1H), 7.42 (dd, *J* = 9.2, 4.4 Hz, 1H), 7.33 (t, *J* = 8.2 Hz, 1H), 5.39 (d, *J* = 16.1 Hz, 1H), 4.48 – 4.23 (m, 4H), 4.17 (d, *J* = 5.7 Hz, 4H), 3.98 (d, *J* = 13.6 Hz, 1H), 3.84 – 3.37 (m, 45H), 2.40 (s, 2H), 1.42 (t, *J* = 7.2 Hz, 3H). (ES-LCMS) *m*/*z* 592.1 $(M+H)^+$.

Step 8: Synthesis of **Flumazenil.1$^{DART.2}$ (R14)**

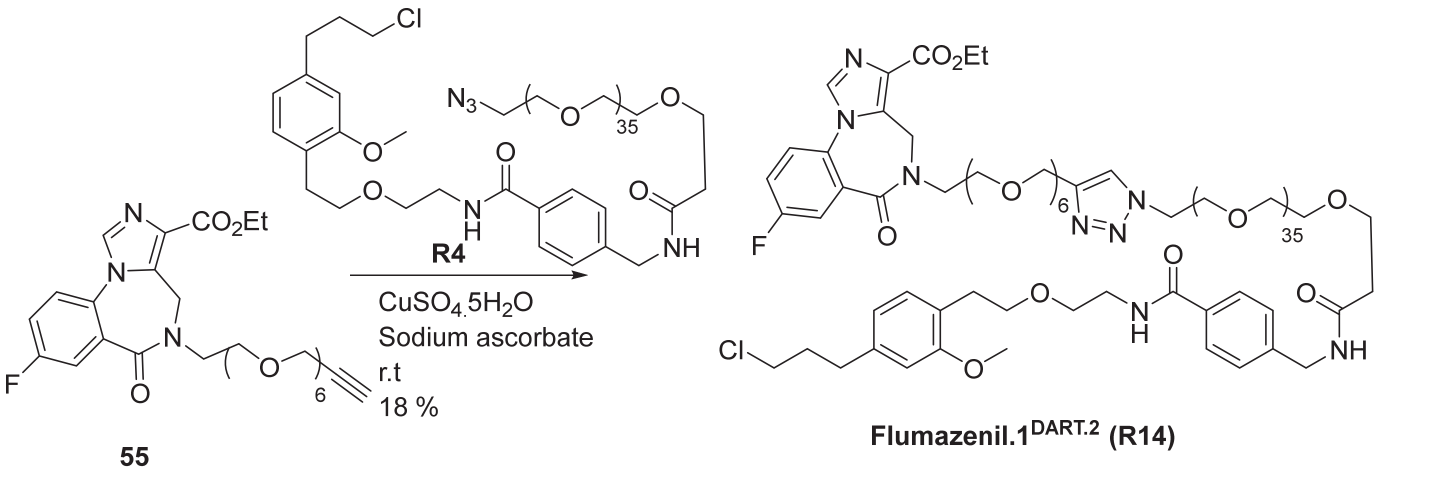


To a vial containing ethyl 8-fluoro-5-(3,6,9,12,15,18-hexaoxahenicos-20-yn-1-yl)-6-oxo-5,6-dihydro-4H-benzo[f]imidazo[1,5-a][1,4]diazepine-3-carboxylate, **55** (5 mg, 8.5 μmol) was added a solution of azido-PEG36 HTL2.0, **R4** (18 mg, 8.5 μmol) in EtOH:t-BuOH:water (2:1:1, 0.8 mL). Then, copper (II) sulfate pentahydrate (0.1 mg, 0.42 μmol) and sodium ascorbate (0.15 mg, 0.85 μmol) were added and the reaction mixture was

stirred overnight after which it was purified by RP HPLC eluting with 10 to 80 % acetonitrile in water (0.1 % FA, 220 nm wavelength) and other conditions as described in the general chemistry synthesis section to give **Flumazenil.1$^{DART.2}$, R14** (4 mg, 18 %).
$^{1}$H NMR (500 MHz, $CDCl_3$) δ 7.86 (s, 1H), 7.77 (dd, *J* = 8.8, 2.9 Hz, 1H), 7.72 – 7.63 (m, 3H), 7.43 (dt, *J* = 8.8, 4.5 Hz, 1H), 7.36 – 7.30 (m, 3H), 7.14 – 7.01 (m, 2H), 6.68 – 6.60 (m, 2H), 6.50 (s, 1H), 5.39 (d, *J* = 16.0 Hz, 1H), 4.65 (d, *J* = 5.5 Hz, 2H), 4.53 – 4.46 (m, 4H), 4.44 – 4.28 (m, 2H), 3.98 (d, *J* = 13.7 Hz, 1H), 3.85 (t, *J* = 5.2 Hz, 2H), 3.79 – 3.72 (m, 6H), 3.70 – 3.43 (m, 170H), 2.89 – 2.78 (m, 3H), 2.71 (t, *J* = 7.4 Hz, 2H), 2.53 (t, *J* = 5.6 Hz, 2H), 2.03 (dq, *J* = 8.3, 6.6 Hz, 2H), 1.42 (t, *J* = 7.1 Hz, 3H). HRMS (ESI$^{+}$): *m/z* calcd for $C_{126}H_{214}ClFN_8Na_2O_{49}$: 1362.1995 $[M+2Na]^{2+}$; found: 1362.1700 $[M+2Na]^{2+}$

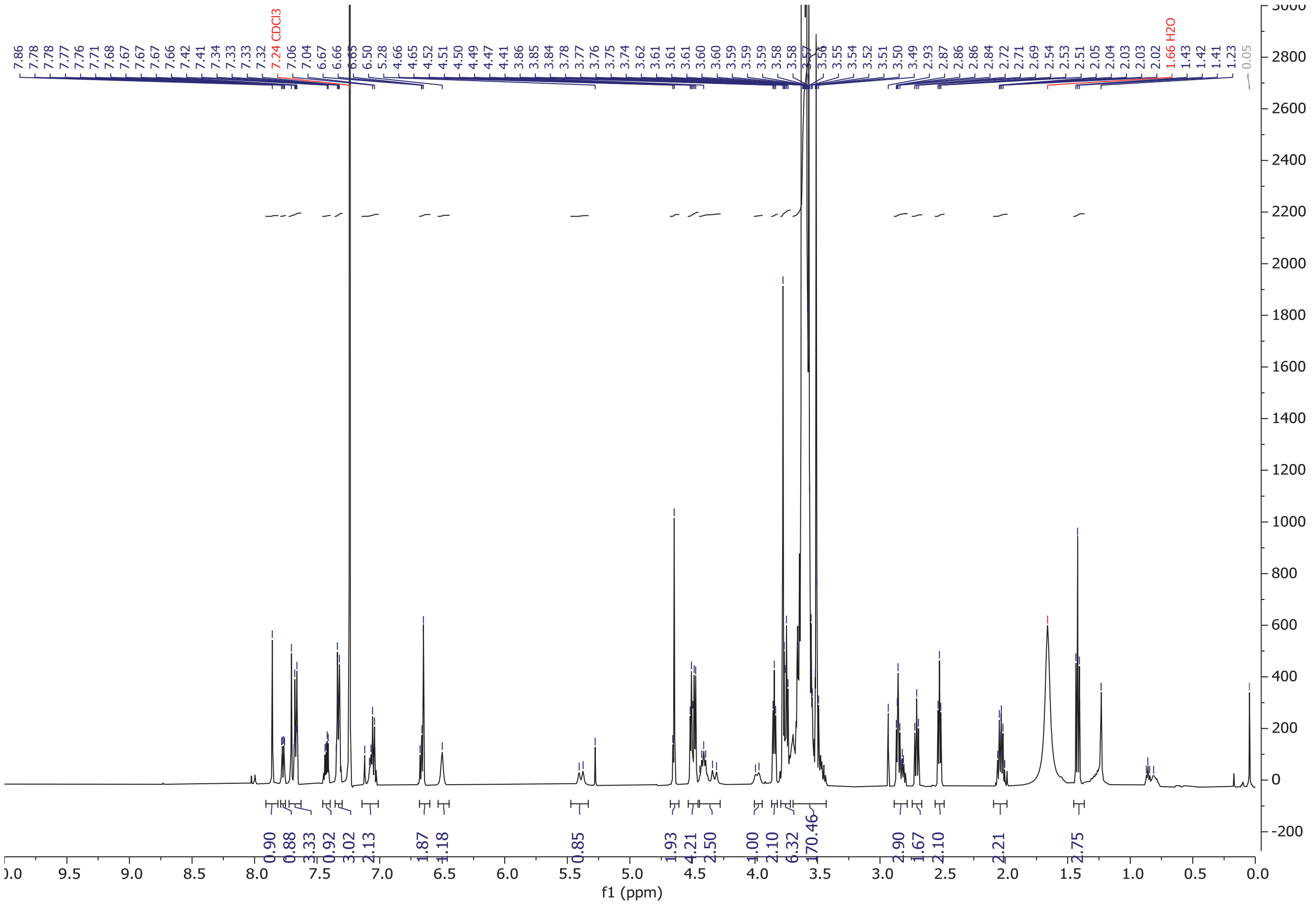

**$^{1}$H NMR of Flumazenil.1$^{DART.2}$ (R14) in $CDCl_3$**

### Synthesis of R15–CMPDA.1[DART.1]

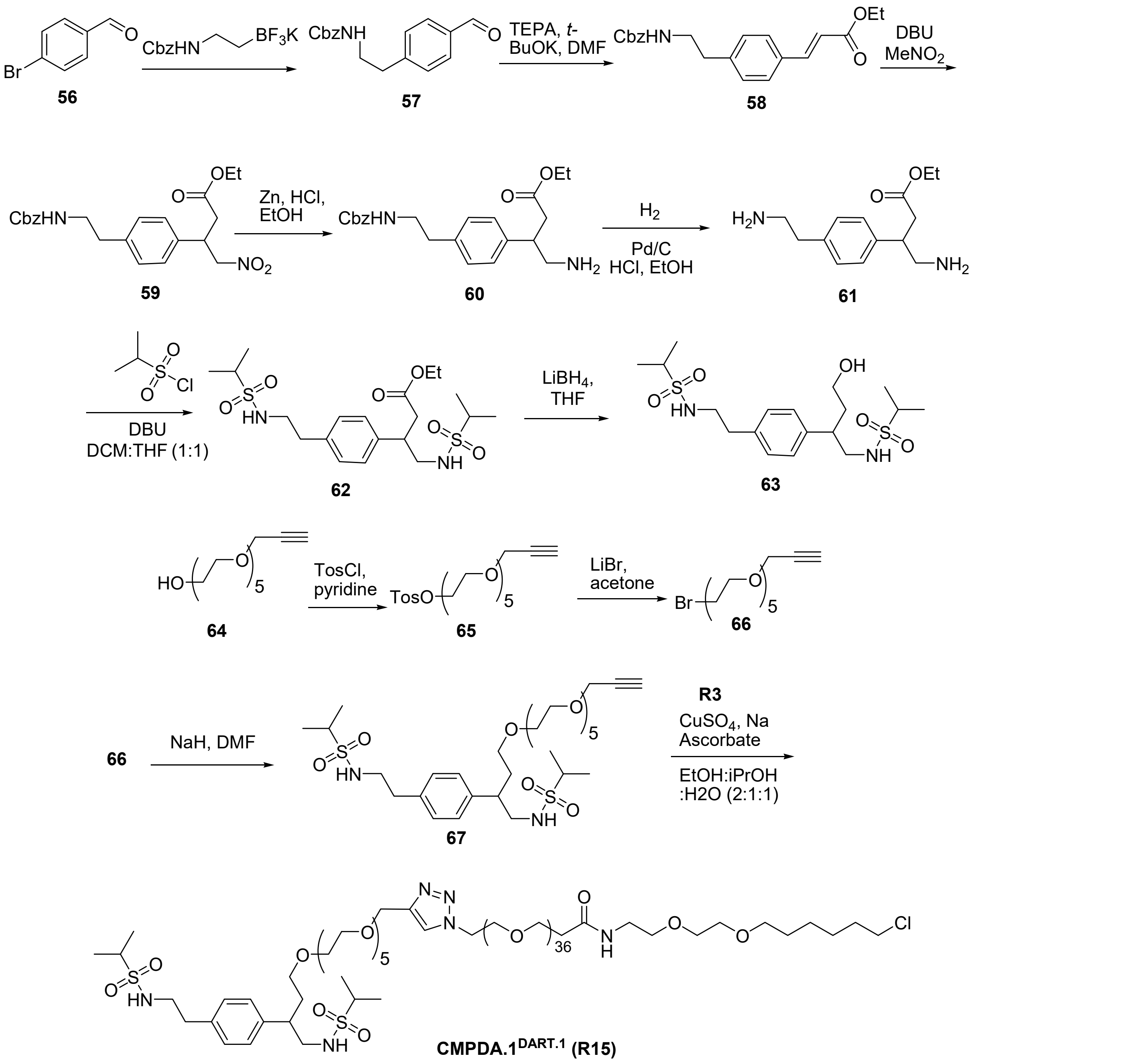


<u>Step 1: Synthesis of benzyl 4-formylphenethylcarbamate, **57**</u>

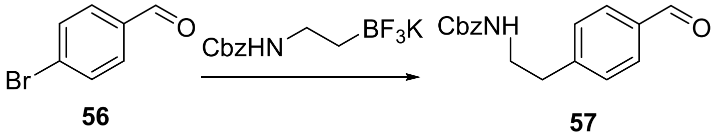


To solution of 4-bromobenzaldehyde (2.36 g, 12.7 mmol) in toluene (75 mL) in a sealed tube were added potassium potassium benzyl N-[2-(trifluoroboranuidyl)ethyl]carbamate (4.0 g, 14.0 mmol), $Cs_2CO_3$ (12.4 g, 38.2 mmol), $PdCl_2(dppf).CH_2Cl_2$ (0.52 g, 0.64 mmol) and water (25 mL). The reaction mixture was purged with nitrogen. The tube was sealed and heated to 80 °C for 18 h. The reaction was allowed to cool to room temperature and saturated solution of $NH_4Cl$ was added. The suspension was extracted with $CH_2Cl_2$ (30 mL X 3 times). The organic extracts were combined, dried ($Na_2SO_4$), and concentrated under reduced pressure to provide crude compound. Purification by flash chromatography (24g silica cartridge, EtOAc/Hexane) gave the title compound (2.0 g, 56% yield).

Step 2: Synthesis of benzyl-4-((E/Z)-2-(ethoxycarbonyl)vinyl)phenethylcarbamate, **58**

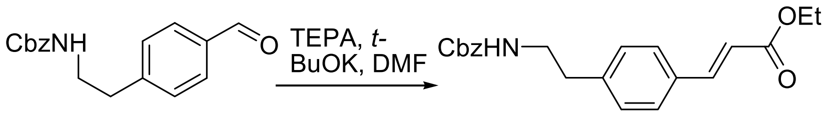


**57** **58**

Triethylphosphonoacetate (9.6 g, 2.4 mmol) and potassium t-butoxide (1.1 g, 27.1 mmol) were dissolved in DMF (10 mL) at 0°C and stirred 30 minutes. Benzyl-4-formylphenethylcarbamate (3.3 g, 13.5 mmol) dissolved in DMF (15 mL) was then added over 5 minutes. Reaction stirred overnight. The reaction was diluted with water and then extracted thrice (EtOAc). The organic extracts were combined, dried ($Na_2SO_4$), and concentrated under reduced pressure to provide crude compound. Purification by flash chromatography (24g silica cartridge, EtOAc/Hexane) gave the title compound (1.96 g, 78% yield). (ESI) *m/z* 354 $(M+1)^+$.

Step 3: Synthesis of benzyl 4-(1-(ethoxycarbonyl)-3-nitropropan-2-yl)phenethylcarbamate, **59**

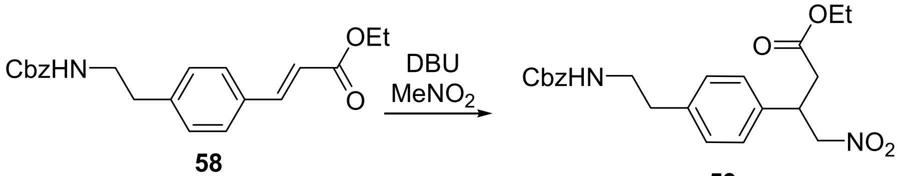


**58** **59**

The starting material benzyl-4-((E/Z)-2-(ethoxycarbonyl)vinyl)phenethylcarbamate (1.96 g, 5.55 mmol) was dissolved in nitromethane (20 mL) and 1,8-diazabicyclo 5.4.0 undec-7-ene (1.5 g, 10 mmol) was then added. The reaction mixture was stirred at room temperature for 48h. The reaction mixture was diluted with ethyl acetate and washed twice with aqueous 1N HCl. The organic layer was dried ($Na_2SO_4$), and concentrated under reduced pressure to provide crude product. Purification by flash chromatography (EtOAc/Hexane) gave the title compound (2.34 g, quant. yield). $^1$H NMR (300 MHz, CHLOROFORM-d) δ = 7.36 (s, 5H), 7.16 (s, 4H), 5.10 (s, 2H), 4.82 - 4.54 (m, 3H), 4.17 - 3.91 (m, 4H), 3.51 - 3.37 (m, 2H), 2.83 - 2.69 (m, 4H), 2.06 (s, 1H), 1.42 - 1.07 (m, 3H). (ESI) *m/z* 415 $(M+1)^+$.

Step 4: Synthesis of benzyl 4-(1-(ethoxycarbonyl)-3-aminopropan-2-yl)phenethylcarbamate, **60**

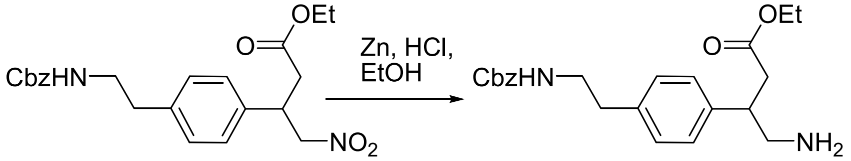


**59** **60**

To benzyl 4-(1-(ethoxycarbonyl)-3-nitropropan-2-yl)phenethylcarbamate (3.35 g, 8.1 mmol) in ethanol (100 mL) was added zinc (5.25 g, 81 mmol) and 4N HCl in dioxane (24 mL, 97 mmol). After 1 hour, the reaction was filtered through celite and the filtrate concentrated. The residue was dissolved in ethyl acetate and washed with aqueous sodium bicarbonate. The aqueous layer was extracted twice (EtOAC). The organic extracts were combined, dried ($Na_2SO_4$), and concentrated under reduced pressure to provide crude compound. Purification by flash chromatography (24g silica cartridge, EtOAc/Hexane) gave the title compound (3.11 g, quant. yield). (ESI) *m/z* 385 $(M+1)^+$.

Step 5: Synthesis of ethyl 4-amino-3-(4-(2-aminoethyl)phenyl)butanoate, **61**

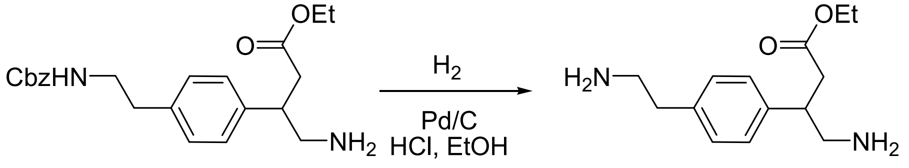


**60** **61**

To benzyl-4-(1-(ethoxycarbonyl)-3-aminopropan-2-yl)phenethylcarbamate (3.11 g, 8.1 mmol) in ethanol (100 mL) in a Parr bottle was added 10% Pd/C (350 mg) and 4N HCl in dioxane (6.1 mL). The sample was hydrogenated (50 psi) for 20 hours. Methanol was added to solubilize the product and the reaction was then filtered through celite and concentrated to crude product used without further purification. $^1$H NMR (300 MHz, CHLOROFORM-d) δ = 7.19 (br s, 4H), 4.08 (br s, 3H), 3.40 (br s, 3H), 3.10 (br d, $J$ = 5.9 Hz, 2H), 2.86 (br s, 1H), 2.81 - 2.71 (m, 1H), 2.70 - 2.59 (m, 1H), 1.39 - 1.25 (m, 5H), 1.20 (br s, 2H). (ESI) $m/z$ 251 $(M+1)^+$.

Step 6: Synthesis of ethyl 4-(propane-2-sulfonamido)-3-{4-[2-(propane-2-sulfonamido)ethyl]phenyl}butanoate, **62**

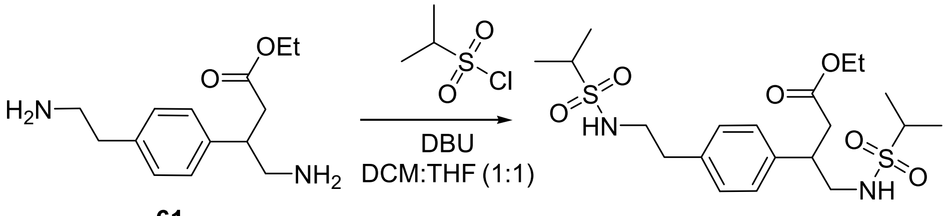


**61** **62**

Ethyl 4-amino-3-(4-(2-aminoethyl)phenyl)butanoate (2.07 g, 6.4 mmol) in $CH_2Cl_2$:THF (60 mL, 1:1) was cooled to 0 °C. 1,8-Diazabicyclo 5.4.0 undec-7-ene (5.7 mL, 38.4 mmol), and isopropyl sulfonyl chloride (2.73 g, 19.2 mmol) were then added. The reaction was stirred at room temperature for 22 h. The reaction was poured into 1N HCl and the solid collected by filtration to yield title compound (1.73 g, 58%).

Step 7: Synthesis of N-(2-{4-[4-hydroxy-1-(propane-2-sulfonamido)butan-2-yl]phenyl}ethyl)propane-2-sulfonamide, **63**

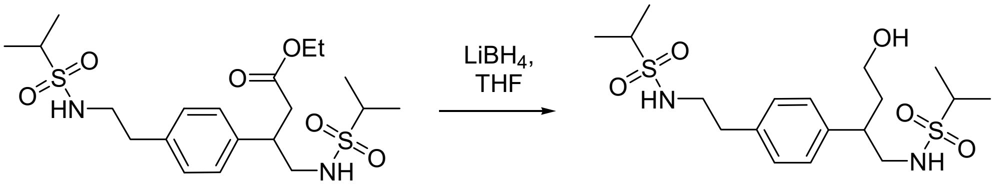


**62** **63**

To ethyl 4-(propane-2-sulfonamido)-3-{4-[2-(propane-2-sulfonamido)ethyl]phenyl}butanoate (200 mg, 0.45 mmol) in THF (15 mL) was added lithium borohydride (4M in THF, 110 □L, 0.45 mmol) and the reaction heated to reflux for 2 hours. After cooling, the reaction was poured into aqueous buffer (pH=7) and extracted thrice with ethyl acetate. The organic extracts were combined, dried ($Na_2SO_4$), and concentrated under reduced pressure. Purification by flash chromatography (EtOAc/Hexane) afforded the title compound (0.16 g, 83 yield). (ESI) $m/z$ 421 $(M+1)^+$.

Step 8: Synthesis of 2-(2-(2-(2-(2-(prop-2-ynyloxy)ethoxy)ethoxy)ethoxy)ethoxy)ethyl 4-methylbenzenesulfonate, **65**

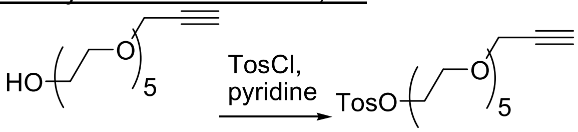


**64** **65**

To 2-(2-(2-(2-(2-(prop-2-ynyloxy)ethoxy)ethoxy)ethoxy)ethoxy)ethanol (250 mg, 0.91 mmol) in pyridine (3 mL) at 0°C was added tosyl chloride (345 mg, 1.81 mmol). The reaction warmed to room temperature while stirring overnight. The reaction was then concentrated and purified by flash chromatography (EtOAc/Hexane) to yield the title compound (57 mg, 14% yield). $^1$H NMR (300 MHz, CHLOROFORM-d) δ = 7.72 (br d, $J$ = 6.7 Hz, 2H), 7.27 - 6.99 (m, 2H), 4.24 - 3.99 (m, 4H), 3.64 - 3.50 (m, 18H), 2.38 (br s, 4H).

Step 9: Synthesis of 3-(2-(2-(2-(2-(2-bromoethoxy)ethoxy)ethoxy)ethoxy)ethoxy)prop-1-yne, **66**

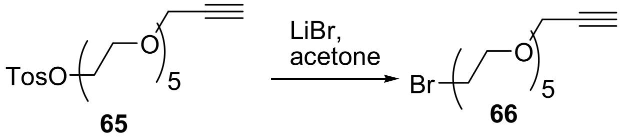


**65** **66**

2-(2-(2-(2-(2-(Prop-2-ynyloxy)ethoxy)ethoxy)ethoxy)ethoxy)ethyl 4-methylbenzenesulfonate (57 mg, 0.132 mmol), and lithium bromide (35 mg, 0.397 mmol) were refluxed in acetone (5 mL) for 3 hours. The reaction was concentrated and residue purified by flash chromatography (EtOAc/Hexane) to yield the title compound (45 mg, quant. yield). $^{1}H$ NMR (300 MHz, CHLOROFORM-d) δ = 4.21 (br s, 2H), 3.86 - 3.76 (m, 2H), 3.67 (br s, 16H), 3.59 - 3.38 (m, 2H), 2.44 (br s, 1H).

Step 10: Synthesis of N-(2-{4-[23-(propane-2-sulfonamido)-4,7,10,13,16,19-hexaoxatricos-1-yn-22-yl]phenyl}ethyl)propane-2-sulfonamide, **67**

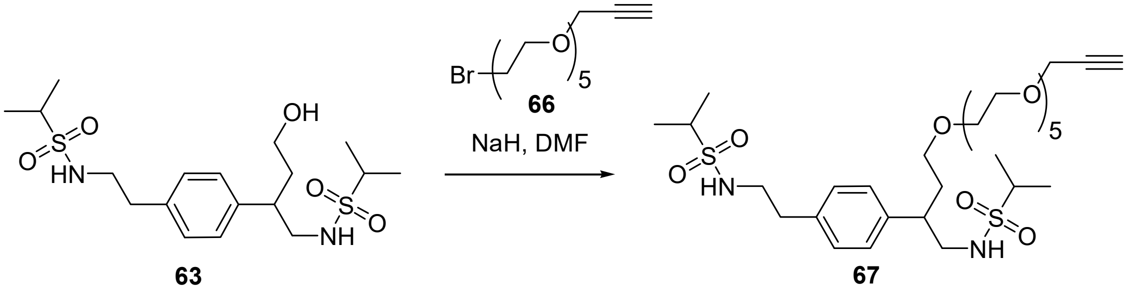


**63** **67**

To N-(2-{4-[4-hydroxy-1-(propane-2-sulfonamido)butan-2-yl]phenyl}ethyl)propane-2-sulfonamide (**63**, 51 mg, 0.12 mmol) in DMF (1 mL) was added 60% sodium hydride in mineral oil (15 mg, 0.37 mmol) and the reaction stirred at 0°C for 30 minutes. 3-(2-(2-(2-(2-(2-Bromoethoxy)ethoxy)ethoxy)ethoxy)ethoxy)prop-1-yne (**66**, 45 mg, 0.133 mmol) was added and the reaction stirred at ambient temperature overnight. The reaction was purified via reverse phase chromatography to yield the title compound (19 mg, 23%). (ESI) *m/z* 679 $(M+1)^{+}$.

Step 11: Synthesis of N-(2-{2-[(6-chlorohexyl)oxy]ethoxy}ethyl)-1-{4-[21-(propane-2-sulfonamido)-20-{4-[2-(propane-2-sulfonamido)ethyl]phenyl}-2,5,8,11,14,17-hexaoxahenicosan-1-yl]-1H-1,2,3-triazol-1-yl}-3,6,9,12,15,18,21,24,27,30,33,36,39,42,45,48,51,54,57,60,63,66,69,72,75,78,81,84,87,90,93,96,99,102,105,108-hexatriacontaoxa111n-111-amide, **CMPDA.1$^{DART.1}$**

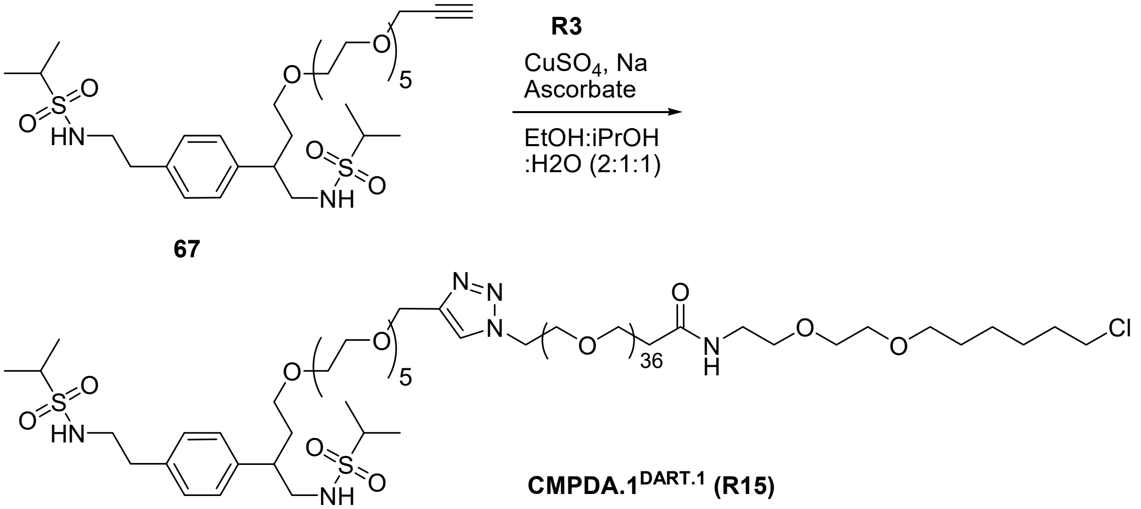


**CMPDA.1$^{DART.1}$ (R15)**

To a vial containing azido-PEG$_{36}$-HTL.1 ($^{azido}$DART.1, **R3**) (37 mg, 0.019 mmol), N-(2-{4-[23-(propane-2-sulfonamido)-4,7,10,13,16,19-hexaoxatricos-1-yn-22-yl]phenyl}ethyl)propane-2-sulfonamide (**67**, 13 mg, 0. 19 mmol), and the solvents EtOH:*i*-PrOH:$H_2O$ (2:1:1) (1 mL) was added $CuSO_4.5H_20$ (0.3 mg, 0.001 mmol) and (+)-sodium L-ascorbate (0.4 mg, 0.002 mmol) pre-dissolved in solutions of EtOH:*i*-PrOH:$H_2O$ (2:1:1) (1 mL). The reaction was stirred for 20h at room temperature. The reaction mixture was then filtered, concentrated, and purified by reverse phase HPLC to afford the title compound, **R15** (36 mg, 72% yield). (ESI) *m/z* 1293 $(^{1/2}M+1)^{+}$.

### Synthesis of R16—CMPDA.2[DART.2]

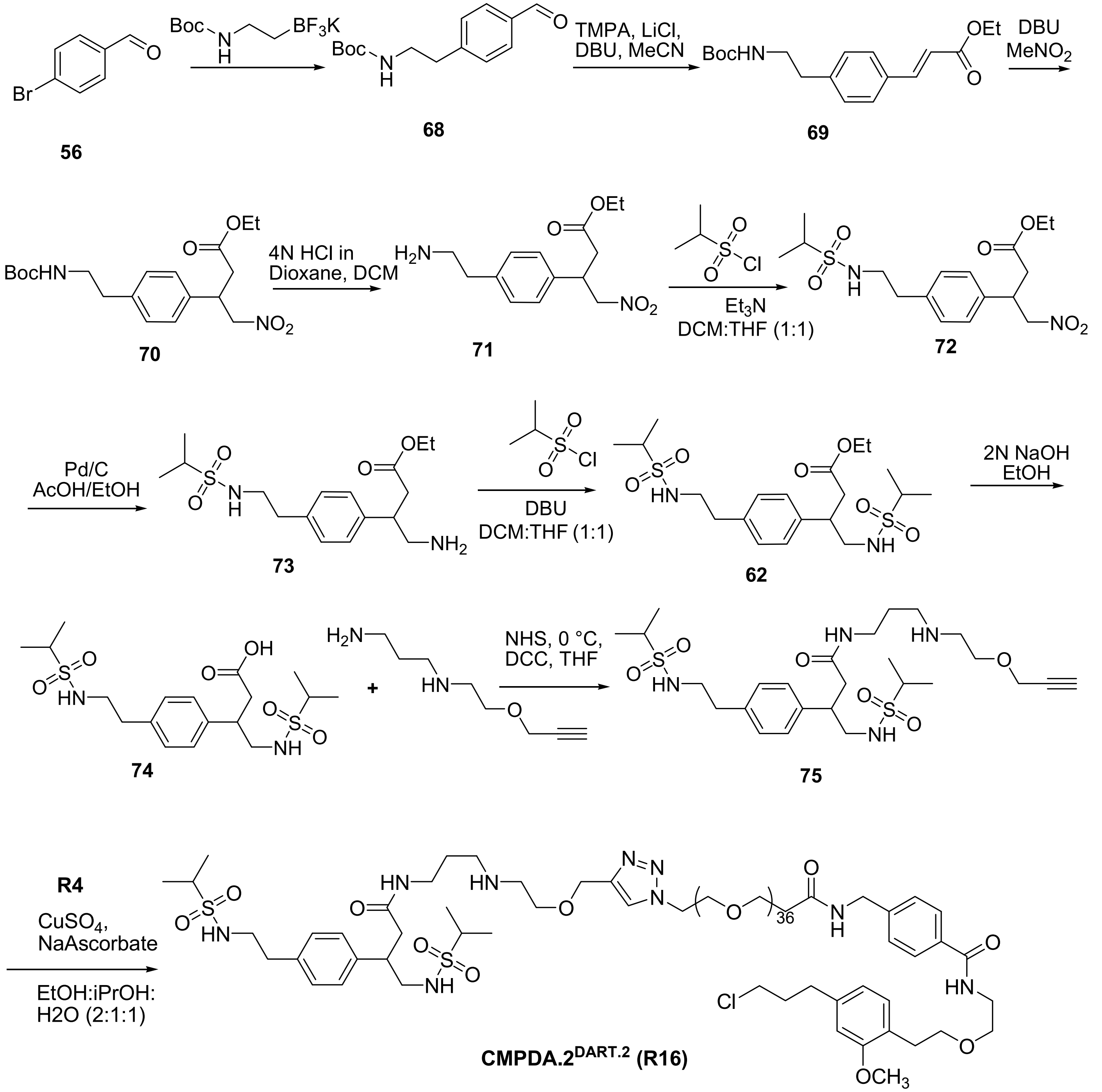

Step 1: *tert*-butyl (4-formylphenethyl)carbamate, **68**

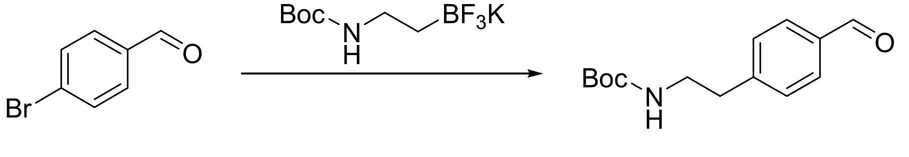


**56** **68**

To solution of 4-bromobenzaldehyde (3.0 g, 16.3 mmol) in toluene (25 mL) in a sealed tube were added potassium tert-butyl *N*-[2-(trifluoroboranuidyl)ethyl]carbamate (4.9 g, 19.6 mmol), $Cs_2CO_3$ (15.9 g, 48.9 mmol), $PdCl_2$(dppf).$CH_2Cl_2$ (2.6 g, 3.2 mmol) and water (5 mL). The rection mixture was degassed under vacuum and filled with nitrogen gas, the process repeated three times. The tube with reaction mixture was sealed and heated to 80 °C for 18 h. The reaction was allowed to cool to room temperature, saturated solution of $NH_4Cl$ (20 mL) was added, and the suspension was extracted with $CH_2Cl_2$ (30 mL X 3 times). The organic extracts were combined, dried ($Na_2SO_4$), and concentrated under reduced pressure to provide crude compound. Purification by flash chromatography (24g silica cartridge, EtOAc/Hexane) gave the title compound (3.3g, 73% yield). $^1$H NMR (CHLOROFORM-d, 300MHz): δ = 9.93 - 10.04 (m, 1 H), 7.74 - 7.90 (m, 2 H), 7.32 - 7.42 (m, 2 H), 4.39 - 4.63 (m, 1 H), 3.23 - 3.54 (m, 2 H), 2.73 - 3.02 (m, 2 H), 1.52 - 1.60 (m, 1 H), 1.43 ppm (s, 9 H). (ESI) *m*/*z* 273 (M+23)$^+$.

Step 2: Ethyl-3-(4-(2-((*tert*-butoxycarbonyl)amino)ethyl)phenyl)-(*E*/*Z*)-acrylate, **69**

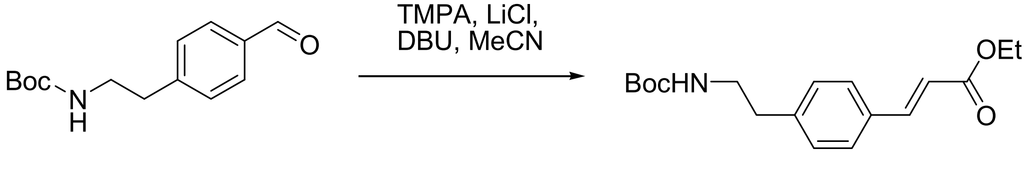


**68** **69**

*Tert*-butyl (4-formylphenethyl)carbamate (3.3 g, 13.5 mmol) was dissolved in acetonitrile (15 mL) and added ethyl 2-(diethoxyphosphoryl)acetate (9.6 g, 2.4 mmol), lithium chloride (1.1 g, 27.1 mmol), 1,8-diazabicyclo 5.4.0 undec-7-ene (1.3 g, 3.9 mmol) added, the reaction mixture was maintained at room temperature for 18 h. After completion of reaction, the reaction mixture was concentrated and added $CH_2Cl_2$ (30 mL) and $H_2O$ (30 mL) the suspension was extracted with $CH_2Cl_2$ (3 × 30 mL). The organic extracts were combined, washed with brine (20 mL), dried ($Na_2SO_4$), and concentrated under reduced pressure to provide crude compound. Purification by flash chromatography (40 g silica cartridge, EtOAc/Hexane) gave the title compound (2.6 g, 60% yield). $^1$H NMR (CHLOROFORM-d, 300MHz): δ = 7.66 (d, *J*=15.8 Hz, 4 H), 7.46 (d, *J*=8.2 Hz, 7 H), 7.17 - 7.28 (m, *J*=11.1 Hz, 9 H), 6.40 (d, *J*=15.8 Hz, 4 H), 4.26 (q, *J*=7.0 Hz, 7 H), 3.38 (q, *J*=6.2 Hz, 7 H), 2.81 (t, *J*=7.0 Hz, 7 H), 1.43 (s, 32 H), 1.33 ppm (t, *J*=7.0 Hz, 12 H). (ESI) *m*/*z* 343 (M+23)$^+$.

Step 3: Synthesis of ethyl 3-(4-(2-((*tert*-butoxycarbonyl)amino)ethyl)phenyl)-4-nitrobutanoate, **70**

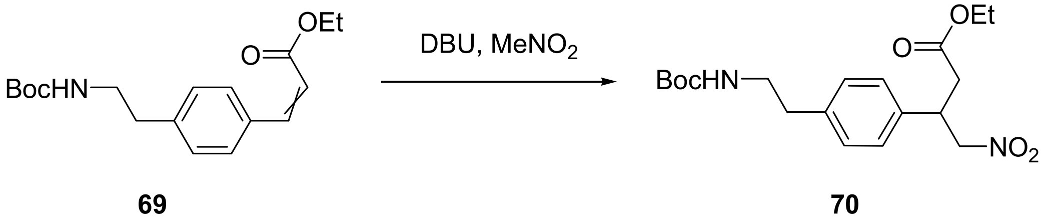


**69** **70**

The starting material ethyl-3-(4-(2-((*tert*-butoxycarbonyl)amino)ethyl)phenyl)-(*E*/*Z*)-acrylate (1.3 g, 3.9 mmol) was dissolved in nitromethane (6.5 mL) and added 1,8-diazabicyclo 5.4.0 undec-7-ene (1.6 g, 3.9 mmol), the reaction mixture was maintained at room temperature for 18 h. After completion of reaction, the reaction mixture was diluted with ethyl acetate (20 mL) and washed with 1N HCl (15 mL X 2 times) the combined aqueous layers were reextracted with ethyl acetate (15 mL). The organic extracts were combined again washed with 1N HCl (10 mL), dried ($Na_2SO_4$), and concentrated under reduced pressure to provide crude. Purification by flash chromatography (40 g silica cartridge, 30% EtOAc in Hexane) gave the title compound (0.8 g, 49% yield). (ESI) *m*/*z* 382 (M+1)$^+$.

Step 4: Synthesis of ethyl 3-(4-(2-aminoethyl)phenyl)-4-nitrobutanoate hydrochloride, **71**

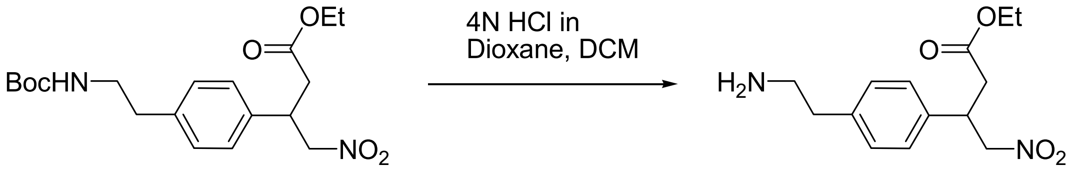


**70** **71**

The ethyl 3-(4-(2-((*tert*-butoxycarbonyl)amino)ethyl)phenyl)-4-nitrobutanoate (1.3 g, 3.4 mmol) was dissolved in dichloromethane (10 mL) and added 4N HCl (3 mL). The reaction was maintained at room temperature overnight. Reaction was monitored by LC-MS. After completion of reaction, the reaction mixture was concentrated *in vacuo* and resultant solids were washed with ether to get desired compound in quantitative yield (0.85 g, 88.7% yield). $^{1}$H NMR (METHANOL-$d_4$, 300MHz): δ = 7.16 - 7.42 (m, 4 H), 4.82 (s, 2 H), 3.83 - 4.13 (m, 2 H), 3.43 - 3.54 (m, 1 H), 3.09 - 3.23 (m, 2 H), 2.88 - 2.99 (m, 2 H), 2.65 - 2.87 (m, 2 H), 1.15 ppm (s, 3 H). (ESI) *m/z* 282 $(M+1)^+$.

Step 5: Synthesis of ethyl 3-(4-(2-((1-methylethyl)sulfonamido)ethyl)phenyl)-4-nitrobutanoate, **72**

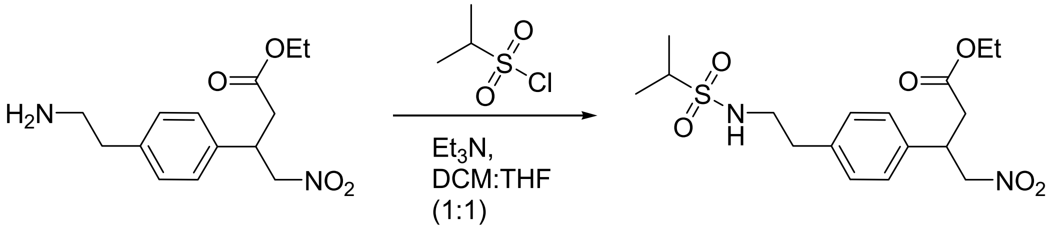


**71** **72**

The starting material ethyl 3-(4-(2-aminoethyl)phenyl)-4-nitrobutanoate·HCl (397 mg, 1.42 mmol) in $CH_2Cl_2$:THF (8 mL, 1:1) was cooled to 0 °C. To reaction mixture added triethylamine (418 uL, 3.0 mmol) and then isopropyl sulfonyl chloride (221 uL, 1.5 mmol) drop wise. The reaction mixture progress was monitored by LC-MS. The reaction was maintained at room temperature for 18 h. To reaction mixture added EtOAc and the suspension was extracted with EtOAc (30 mL X 3 times). The organic extracts were combined, dried ($Na_2SO_4$), and concentrated under reduced pressure to provide crude compound. Purification by flash chromatography (24 g silica cartridge, 10% MeOH in $CH_2Cl_2$) gave the title compound (207 mg, 45%). (ESI) *m/z* 388 $(M+1)^+$.

Step 6: Synthesis of ethyl 4-amino-3-(4-(2-((1-methylethyl)sulfonamido)ethyl)phenyl)butanoate, **73**

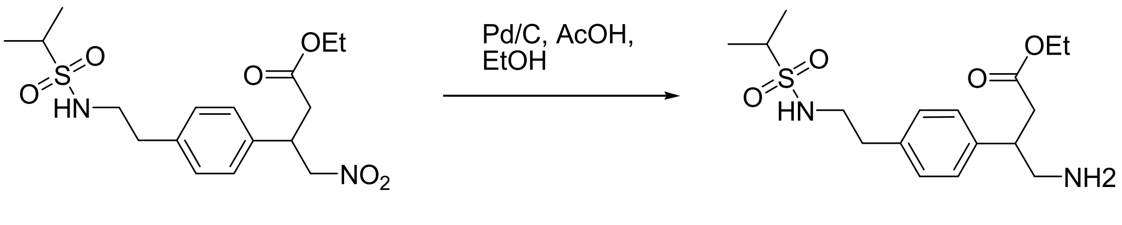


**72** **73**

To a solution of ethyl 3-(4-(2-((1-methylethyl)sulfonamido)ethyl)phenyl)-4-nitrobutanoate (200 mg, 0.51 mmol) in ethanol, under nitrogen atmosphere added 10% Pd/C (110.6 mg, 0.15 mmol) and setup was placed on par hydrogenator and maintained under hydrogen atmosphere 50psi for 18 hours. The reaction was checked by LC-MS and after complete conversion, the reaction mixture was filtered through celite and concentrated *in vacuo* to get crude compound (82 mg, 44%). The crude product was used as is without further purification. (ESI) *m/z* 358 $(M+1)^+$.

Step 7: Synthesis of ethyl 4-((1-methylethyl)sulfonamido)-3-(4-(2-((1-methylethyl)sulfonamido)ethyl)phenyl)butanoate, **62**

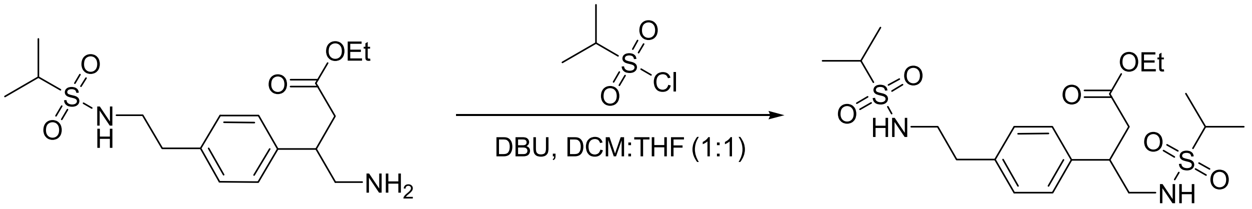

**73** **62**

The starting material ethyl 4-amino-3-(4-(2-((1-methylethyl)sulfonamido)ethyl)phenyl)butanoate (67 mg, 0.189 mmol) in $CH_2Cl_2$:THF (1 mL, 1:1) was cooled to 0 °C. 1,8-Diazabicyclo 5.4.0 undec-7-ene (68 mg, 0.45 mmol), and isopropyl sulfonyl chloride (59 mg, 0.41 mmol) were then added. The reaction was maintained at room temperature for 18 h with stirring. To the reaction mixture was added HCl (5 mL) and EtOAc (10 mL) and the suspension was extracted with EtOAc (3 × 10 mL). The organic extracts were combined, dried ($Na_2SO_4$), and concentrated under reduced pressure to provide crude compound. Purification by flash chromatography (12 g silica cartridge, 10% MeOH in $CH_2Cl_2$) to give the title compound (35 mg, 40%). $^1$H NMR (CHLOROFORM-d, 300MHz): δ = 7.17 (s, 4 H), 4.19 - 4.33 (m, 2 H), 4.00 - 4.13 (m, 2 H), 3.27 - 3.43 (m, 5 H), 2.99 - 3.18 (m, 2 H), 2.56 - 2.88 (m, 4 H), 1.30 (d, *J*=7.0 Hz, 12 H), 1.14 - 1.21 ppm (m, 3 H). (ESI) *m/z* 464 $(M+1)^+$.

Step 8: Synthesis of 4-((1-methylethyl)sulfonamido)-3-(4-(2-((1-methylethyl)sulfonamido)ethyl)phenyl)butanoic acid, **74**

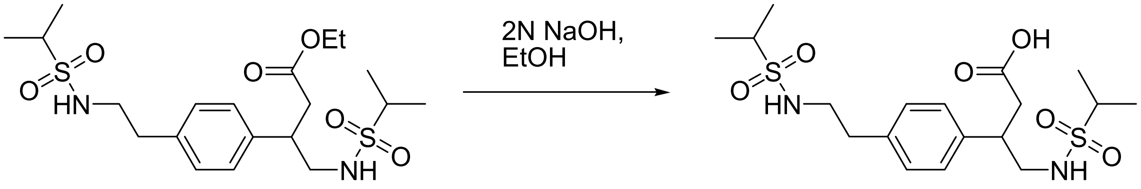

**62** **74**

The starting material ethyl 4-((1-methylethyl)sulfonamido)-3-(4-(2-((1-methylethyl)sulfonamido)ethyl)phenyl)butanoate (33 mg, 0.09 mmol) was dissolved in ethanol (0.5 mL). To this solution, sodium hydroxide (36.8 mg, 0.87 mmol) was added and reaction stirred at room temperature for 3 h. After completion of reaction, the reaction mixture was concentrated and $CH_2Cl_2$ and 1N HCl (3 mL) were added. The suspension was extracted with $CH_2Cl_2$ (3 × 5 mL). The organic extracts were combined dried ($Na_2SO_4$), and concentrated under reduced pressure to provide crude compound, which is used as is for next step. $^1$H NMR (METHANOL-d$_4$, 300MHz): δ = 7.17 - 7.31 (m, 4 H), 3.35 (s, 1 H), 3.30 (br. s., 4 H), 3.07 (dq, *J*=13.5, 6.8 Hz, 2 H), 2.78 - 2.94 (m, *J*=7.0 Hz, 3 H), 2.52 - 2.69 (m, *J*=4.7 Hz, 1 H), 1.18 - 1.34 ppm (m, 12 H). (ESI) *m/z* 436 $(M+1)^+$.

Step 9: Synthesis of 4-((1-methylethyl)sulfonamido)-3-(4-(2-((1-methylethyl)sulfonamido)ethyl)phenyl)-*N*-(3-((2-(prop-2-yn-1-yloxy)ethyl)amino)propyl)butanamide, **75**

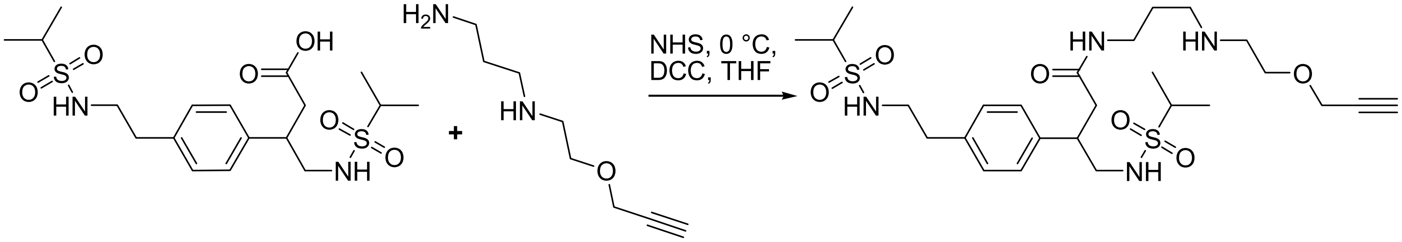

**74** **75**

A

solution of 4-((1-methylethyl)sulfonamido)-3-(4-(2-((1-methylethyl)sulfonamido)ethyl)phenyl)butanoic acid (30.2 mg, 0.07 mmol) and *N*-hydroxysuccinimide (8 mg, 0.07 mmol) in dry THF (0.3 mL) was cooled to 0 °C in an ice bath. Dicyclohexylcarbodiimide (17.9 mg, 0.09 mmol) dissolved in dry THF (0.5 mL) was added drop wise to the mixture and the reaction stirred at room temperature for 24 h. The reaction mixture was filtered to remove dicyclohexylurea by-product formed. The filtrate was concentrated under reduced pressure. The residue was dissolved in $CHCl_3$ and washed consecutively with water, a saturated solution of $NaHCO_3$ and water. The organic layer was dried ($Na_2SO_4$), and concentrated under reduced pressure to obtain the *N*-hyroxysuccinimide active ester intermediate. This intermediate (37.0 mg, 0.07 mmol) and $N^1$-(2-(prop-2-yn-1-yloxy)ethyl)propane-1,3-diamine (20.0 mg, 0.07 mmol) were dissolved in $CHCl_3$ (0.3 mL) with addition of small amount of DMF (35 uL). The reaction stirred at room temperature for 48 h. After completion of reaction, the reaction mixture was washed with $H_2O$, sat. $NaHCO_3$, and $H_2O$. filtered off the desiccant, dried over $Na_2SO_4$, concentrated and purified by reverse phase preparative HPLC (mobile phase: 0.1% TFA in acetonitrile and 0.1% TFA in water) to give pure compound 9.0 mg, 23% yield. $^1H$ NMR (METHANOL-d$_4$ ,300MHz): δ= 7.23 (s, 4 H), 4.22 - 4.30 (m, 2 H), 3.75 (t, *J*=5.0 Hz, 2 H), 3.24 (br. s., 4 H), 2.93 - 3.17 (m, 6 H), 2.71 - 2.87 (m, *J*=7.0, 7.0 Hz, 4 H), 2.35 - 2.58 (m, *J*=14.4, 9.1 Hz, 2 H), 1.57 - 1.80 (m, 2 H), 1.16 - 1.37 ppm (m, 12 H). (ESI) *m*/*z* 574 $(M+1)^+$.

Step 10: Synthesis of reagent CMPDA.$2^{DART.2}$ (R16)

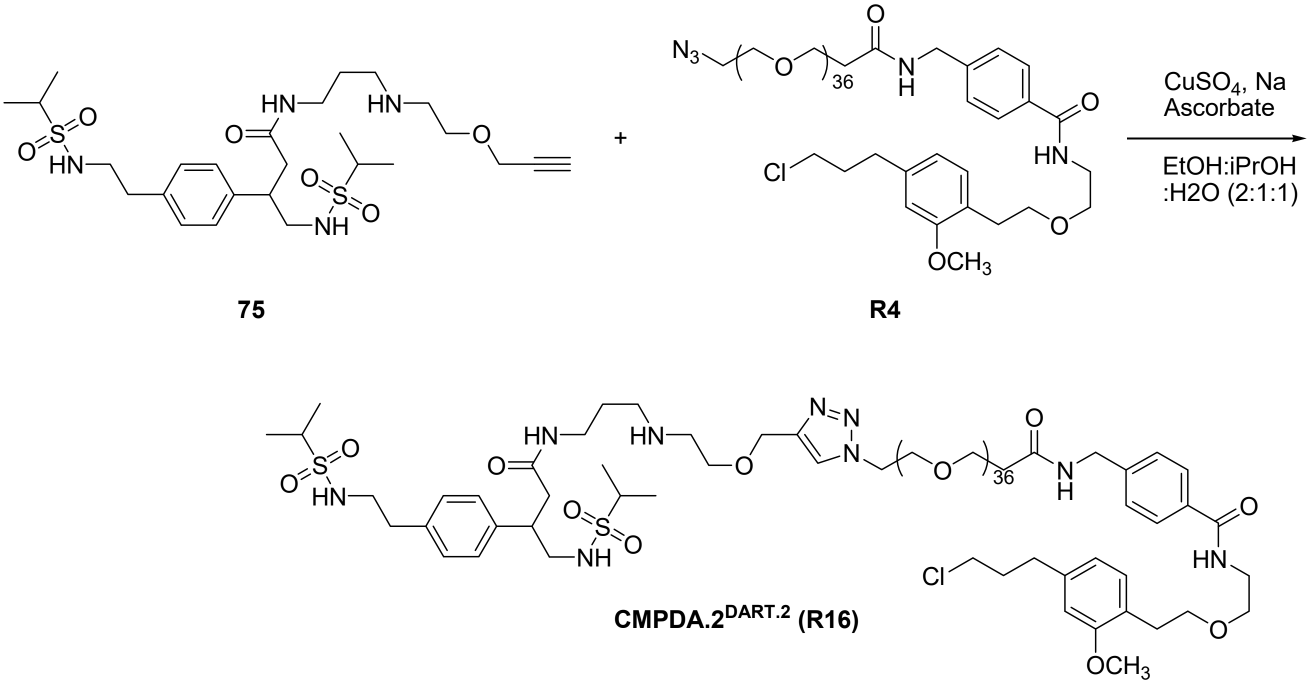


**75** **R4**

**CMPDA.$2^{DART.2}$ (R16)**

To a vial containing the azido-PEG$_{36}$-HTL.2 ($^{azido}$DART.2, **R4**) (28.4 mg, 0.013 mmol) and 4-((1-methylethyl)sulfonamido)-3-(4-(2-((1-methylethyl)sulfonamido)ethyl)phenyl)-*N*-(3-((2-(prop-2-yn-1-yloxy)ethyl)amino)propyl)butanamide (7.8 mg, 0.013 mmol), the solvent mixture EtOH:iPrOH:$H_2O$ (2:1:1) (200 uL) was added followed by $CuSO_4.5H_20$ (2.39 mg, 0.015 mmol) and (+)-sodium L-ascorbate (3.0 mg, 0.015 mmol) was pre-dissolved in solutions of EtOH:iPrOH:$H_2O$ (2:1:1) (400 uL). The reaction was stirred for 18 h at room temperature. After completion of reaction, reaction mixture was filtered, concentrated and purified by reverse phase preparative HPLC (mobile phase: 0.1% TFA in acetonitrile and 0.1% TFA in water) to get pure compound **R16** 17.0 mg, 47% yield. $^1H$ NMR (300 MHz, METHANOL-*d$_4$*) δ ppm 8.19 - 8.25 (m, 1 H) 7.84 - 7.97 (m, 2 H) 7.51 (d, *J*=7.03 Hz, 2 H) 7.33 (s, 4 H) 7.17 (d, *J*=7.03 Hz, 1 H) 6.87 (br. s., 1 H) 6.76 (d, *J*=7.62 Hz, 1 H) 4.86 (s, 1 H) 4.80 (br. s., 1 H) 4.71 (br. s., 1 H) 4.57 (br. s., 2 H) 3.99 (m, *J*=3.51 Hz, 3 H) 3.83 - 3.94 (m, 8 H) 3.57 - 3.83 (m, 158 H) 3.20 (m, *J*=11.72 Hz, 4 H) 2.88 (m, *J*=7.62 Hz, 8 H) 2.50 - 2.70 (m, 4 H) 2.14 (t, *J*=6.44 Hz, 2 H) 1.81 (br. s., 2 H) 1.37 (m, *J*=2.34 Hz, 12 H); (ESI) *m*/*z* 1332 $(M/2+1)^+$; HRMS (ESI): m/z calcd for $C_{123}H_{221}ClN_9O_{46}S_2$: 2659.4341 $[M+H]^+$; found: 2659.43551, $[M+H]^+$

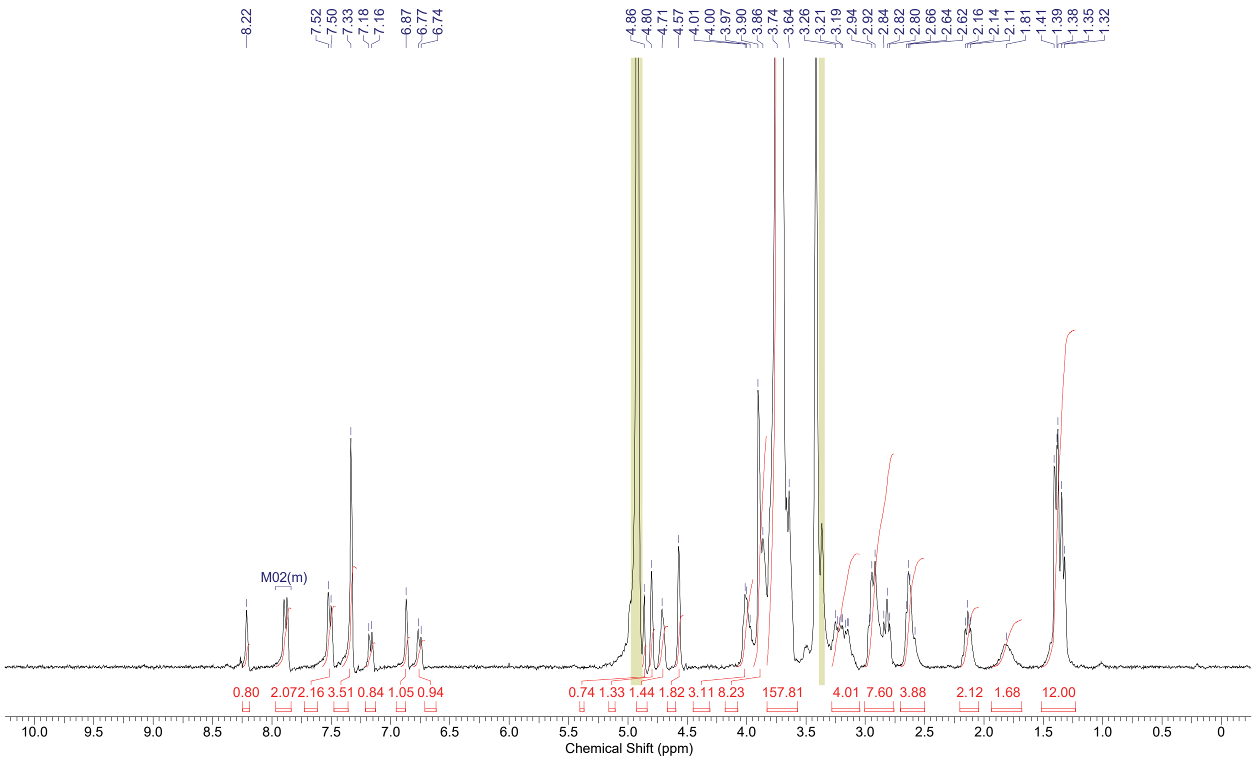

**[1]H NMR of CMPDA.2[DART.2], R16**
